# Benzylic Radical Stabilization Permits Ether Formation During Darobactin Biosynthesis

**DOI:** 10.1101/2023.11.29.569256

**Authors:** Austin M. Woodard, Francesca Peccati, Claudio D. Navo, Gonzalo Jiménez-Osés, Douglas A. Mitchell

## Abstract

The Gram-negative selective antibiotic darobactin A has attracted interest owing to its intriguing fused bicyclic structure and unique mode of action. Biosynthetic studies have revealed that darobactin is a ribosomally synthesized and post-translationally modified peptide (RiPP). During maturation, the darobactin precursor peptide (DarA) is modified by a radical *S*-adenosyl methionine (rSAM)-dependent enzyme (DarE) to contain ether and C-C crosslinks. In this work, we describe the enzymatic tolerance of DarE using a panel of DarA variants, revealing that DarE can install the ether and C-C crosslinks independently and in different locations on DarA. These efforts produced 57 darobactin variants, 50 of which were enzymatically modified. Several new variants with fused bicyclic structures were characterized, including darobactin W3Y, which replaces tryptophan with tyrosine at the twice-modified central position, and darobactin K5F, which displays a fused diether ring pattern. Three additional darobactin variants contained fused diether macrocycles, leading us to investigate the origin of ether versus C-C crosslink formation. Computational analyses found that more stable and long-lived Cβ radicals found on aromatic amino acids correlated with ether formation. Further, molecular docking and calculated transition state structures provide support for the different indole connectivity observed for ether (Trp-C7) and C-C (Trp-C6) crosslink formation. We also provide experimental evidence for a β-oxotryptophan modification, a proposed intermediate during ether crosslink formation. Finally, mutational analysis of the DarA leader region and protein structural predictions identified which residues were dispensable for processing and others that govern substrate engagement by DarE. Our work informs on darobactin scaffold engineering and sheds additional light on the underlying principles of rSAM catalysis.

## Introduction

The biosynthetic gene clusters (BGCs) for ribosomally synthesized and post-translationally modified peptides (RiPPs) encode a precursor peptide and the cognate modifying enzyme(s). Canonical precursor peptides possess an N-terminal leader region that contains a recognition sequence that is directly engaged by biosynthetic proteins and a C-terminal core region that receives all enzymatic modifications.^1^ Occasionally, RiPP precursor peptides display a C-terminal follower sequence, which, like the leader region, is not part of the final molecule. Some RiPP BGCs encode peptidases to release unmodified portion(s) of the substrate peptide, while others co-opt proteases encoded elsewhere in the genome.^1–3^ Similarly, RiPP BGCs may encode a dedicated transporter for export of the final product.

Numerous advances pertaining to RiPP genome-mining, natural product discovery, and enzyme mechanism have been recently reported. Structurally intriguing products are routinely identified as being formed by radical *S*-adenosylmethionine (rSAM) enzymes, which constitute one of the largest and most diverse enzyme superfamilies (PF04055).^4–6^ Within the rSAM active site, a [4Fe-4S] center reductively cleaves SAM to produce L-methionine and 5’-deoxyadenosyl radical (5’-dAdo•).^7^ Hydrogen atom abstraction from the substrate by 5’-dAdo• propagates a radical cascade from which different reaction pathways are available to generate the final product(s).^4,8^ As compiled by the web-based repository www.radicalsam.org, ∼100,000 rSAM enzymes contain one or more additional [Fe-S] centers and are classified as members of the SPASM/twitch group.^5,9,10^ These auxiliary [Fe-S] centers are proposed to aid in substrate alignment and facilitate electron transfer.^11–13^ Within RiPP biosynthesis, rSAM enzymes catalyze numerous modifications, including methylation, carbon-carbon and carbon-sulfur crosslinking, epimerization, and more.^1,14–25^

Darobactin A is a twice-cyclized rSAM-modified RiPP (Figure 1).^26^ The precursor peptide DarA (WP_152962143.1) possesses a heptapeptide core region (_1_WNWSKSF_7_) that is modified with two crosslinks: *(i)* an ether (C-O-C) between Trp1-C7 and Trp3-Cβ and *(ii)* a C-C crosslink between Trp3-C6 and Lys5-Cβ. Biological mode of action studies identified that darobactin targets BamA of the β-barrel assembly machinery (BAM) complex, which folds and inserts outer membrane proteins (OMPs) in Pseudomonadota (formerly Proteobacteria).^26–29^ Structural data show that darobactin acts as a β-strand mimetic that binds to the lateral gate of BamA analogously to incoming nascent OMPs.^30–34^ The tight binding of darobactin to BamA prevents OMP folding and leads to cell-membrane disruption and bacterial lysis.^26,30^

**Figure 1:**
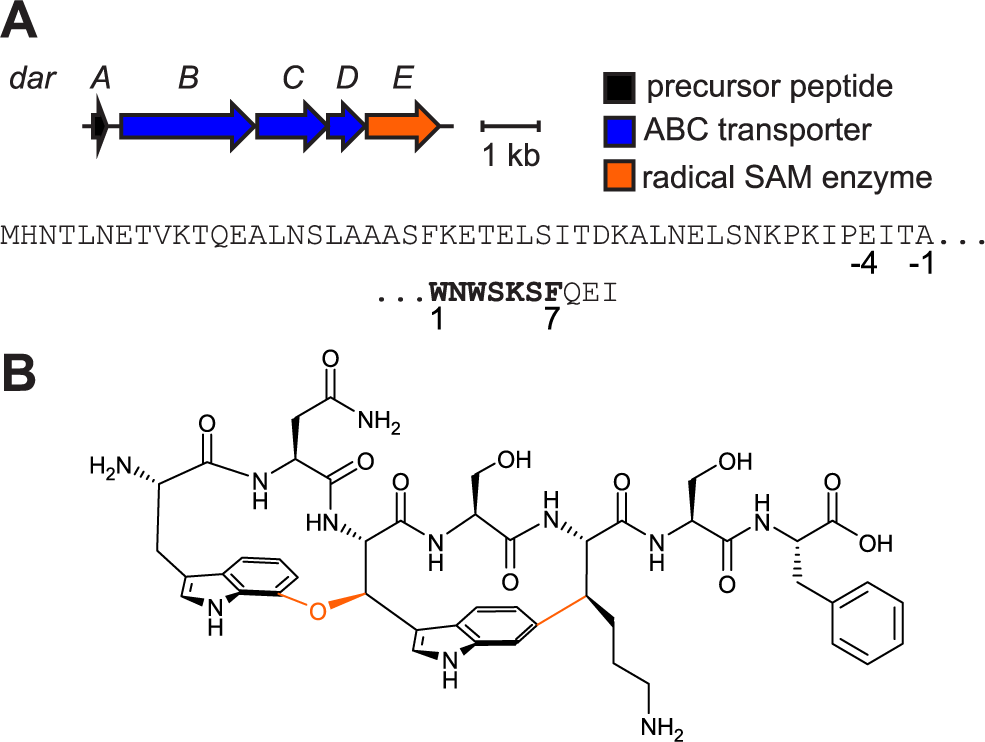
Darobactin BGC and structure. (A) Darobactin biosynthetic gene cluster (BGC) diagram and DarA sequence. *Top*, sequence of leader region; *bottom*, sequence of heptapeptide core (bold) and Gln-Glu-Ile tripeptide follower sequence. Glu(-4) provides a convenient site for endoproteinase GluC cleavage. (B) Structure of darobactin. Orange bonds and atoms indicate DarE modifications (i.e., Trp1-C7 to Trp3 - Cβ ether and Trp3-C6 to Lys5-Cβ C-C).

DarE (WP_152962147.1) installs both crosslinks onto DarA.^35,36^ While precedented, it is rare for a single active site enzyme to catalyze two distinct reaction outcomes on the same substrate.^37,38^ Recent mechanistic work has begun to elucidate crosslink formation by DarE.^39^ In one proposal, hydrogen abstraction by 5’-dAdo• generates a Trp3-Cβ radical on DarA, which further reacts with molecular oxygen to form a hydroperoxide intermediate. The ether crosslink is then formed by either reaction of the Trp1 indole-NH with the hydroperoxide or homolytic cleavage of the hydroperoxide and addition into the Trp1 indole.^39^ An alternative mechanism where hydroxylation of Trp1 precedes ether crosslinking has also been proposed.

Formation of the C-C crosslink has been proposed to be similar to SuiB, a rSAM enzyme responsible for streptide maturation.^13,39^ Lys5-CβH abstraction by 5’-dAdo• produces a Lys5-Cβ radical that reacts with the Trp3 indole to form a crosslinked Trp-Lys radical. Radical quenching, potentially via hydrogen atom abstraction and electron transfer to an auxiliary [Fe-S] center, completes the reaction.^13^ Alternatively, nucleophilic attack by the Trp3 indole side chain onto α,β-dehydrogenated Lys5, formed after Lys5-CβH abstraction, may produce the C-C crosslink.^39^

Many naturally occurring and engineered darobactin variants have been characterized, which have probed the substrate tolerance of DarE. Most of the reported variants involved substitution of the non-modified residues of the DarA core region (i.e., positions 2, 4, 6, 7), which collectively showed minimal impact on forming the fused bicyclic product.^31,34,36,40^ Limited analysis of the modified core residues (i.e., positions 1, 3, and 5) has been performed. DarE tolerated substitutions of Lys5 with either Arg or Ala to produce darobactins D and 15, respectively.^31,36,40^ Until very recently, no experimental data were available regarding the impact of substitutions to DarA core positions 1 and 3. The single tested variant at core position 1 (variant W1F) did not produce either crosslink; however, when core position 3 was changed to Tyr, His, or Phe, an ether crosslink at core positions 1 and 3 was formed. Similarly, a C-C crosslink formed between core positions 1 and 3 on variant W3K, suggesting that the crosslink chemistry depended on the residue identity.^40,41^ However, no bicyclic variants of darobactin have been reported that contain a residue other than Trp at core positions 1 and 3. These previous studies provide initial insight into crosslink formation but raise other questions about how and why a specific crosslink forms between particular residues.

Here, we describe a combination of experimental and computational analyses to investigate the molecular principles of ether vs. C-C crosslink formation by DarE. Single site-saturation mutagenesis of the three modified core positions indicated that each crosslink can be formed independently of the other, and the location is contingent on the properties of the modified residue. Ether crosslinks form between two aromatic amino acids, while C-C crosslinks form between aromatic and aliphatic amino acids. Immature products with β-oxo amino acids were also characterized, implicating oxygen addition to aromatic residues during ether crosslink formation. Computational assessment suggests that the crosslink chemistry is governed by the enhanced longevity of aromatic Cβ radicals over aliphatic Cβ radicals, permitting reaction with molecular oxygen. New darobactin variants with two fused ether crosslinks and one with a non-Trp central residue were characterized. Although these variants lacked antibiotic activity, structural analysis identified key geometric features for engineering darobactins to maintain BamA inhibition. Finally, we determined the positions of the DarA leader region that are vital for substrate processing.

## Results

### DarE Forms the C-C and Ether Crosslinks Independently

Previous *in vivo* biosynthetic analyses of darobactin relied on proteolysis and export of mature darobactins by heterologous or native hosts; however, a limitation of most *in vivo* studies are the challenges in detecting biosynthetic intermediates and shunt products.^31,34,36,42^ To identify and analyze biosynthetically immature peptides, we devised a workflow to reduce endogenous proteolysis of expressed peptides (Figure S1). A co-expression plasmid containing a maltose-binding protein (MBP) fusion to DarA with the Gln-Glu-Ile follower region removed [MBP-DarA(Q8*)] and untagged DarE was constructed. We then performed site-saturation mutagenesis on the three positions modified in DarA (Trp1, Trp3, Lys5) to investigate the substrate tolerance of DarE. After expression in BL21(DE3) *E. coli*, the DarA variants were purified and analyzed by proteolytic digestion and matrix-assisted laser desorption/ionization time-of-flight mass spectrometry (MALDI-TOF-MS) (Figures S2-S4). Glu(-4) of wild-type DarA provided a convenient endoproteinase GluC cut site that excludes all but three residues of the leader region (Ile-Thr-Ala) and was thus used throughout this study. The Ile-Thr-Ala tripeptide was readily removed from variant darobactins by replacing GluC with proteinase K, as previously described.^39^ More detailed structural analysis was performed using high-resolution and tandem mass spectrometry (HR-MS/MS), MALDI-LIFT-TOF/TOF-MS (hereafter, LIFT-MS/MS) and multidimensional NMR spectroscopy.^43^

The most prevalent DarE-installed modification among the DarA variants was a loss of 2 Da (Figures S2-S4). HR-MS/MS analysis of the -2 Da species of multiple DarA variants (e.g., W1D, W1H, W1N, K5Q, K5S, and K5Y) localized the mass loss to Trp3, consistent with previous studies showing DarE can form α,β-dehydrotryptophan (dhTrp) (Figures S5-S10).^39,40^ However, some Trp1 variants (e.g., W1N and W1D) gave very low-intensity daughter ions between Trp3 and Lys5, which in mature darobactin are C-C linked. Given that C-C formation also yields a -2 Da mass loss with hindered MS/MS fragmentation, we performed ultra high-performance liquid chromatography (UHPLC) of GluC-digested DarA-W1N and -W1D variants. In both cases, the -2 Da species eluted into two distinct fractions (Figures S11-S12). The peptides were analyzed by HR-MS/MS and site-specific proteolytic digestion to differentiate dhTrp and C-C crosslink formation at core position 3. Trypsin does not cut after crosslinked Lys5 in darobactin A, and similarly, chymotrypsin does not cut after crosslinked Trp (Figure S13).^40^ The earlier eluting -2 Da species from the DarA-W1N and -W1D variants lacked HR-MS/MS fragmentation within the Trp3-Lys5 motif; both were also recalcitrant to trypsin and chymotrypsin digestion, supporting C-C crosslink formation. HR-MS/MS of the later eluting species localized the modification to Trp3, and these peptides were susceptible to trypsin digestion at Lys5, supporting the absence of the C-C crosslink. We also noticed that variants with a dhTrp modification were not chymotryptic sites (Figures S14-S21).

Formation of a single C-C crosslink was not exclusive to Trp1 variants. Indeed, DarA-W3K, -W3R, -W3N, and -W3Q displayed -2 Da modifications that lacked HR-MS/MS fragmentation between core positions 1 and 3 (Figures S22-S25). The -2 Da modified DarA-W3R variant, for instance, was not cleaved after Trp1 by chymotrypsin or Arg3 by trypsin (Figure 2). To confirm or refute C-C crosslink formation, we produced a larger quantity of the GluC-digested W3R variant. The resulting peptide, which retains Ile-Thr-Ala from the leader region (Figure 1), was purified by UHPLC and subjected to HRMS analysis (C_55_H_84_N_16_O_15_^2+^, *calc*. [M+H]^2+^: 604.3146; *obs*. [M+H]^+^: 604.3143; error: 0.5 ppm) (Figure S26). Subsequent NMR spectroscopic analysis, specifically ^1^H-^1^H TOCSY and ^1^H-^13^C HSQC experiments, yielded diagnostic correlations indicating that Arg3-Cβ was monosubstituted (Figures S27-S31). Similarly, Trp1 was assigned as monosubstituted at C6 via ^1^H-^1^H TOCSY, ^1^H-^13^C HSQC, and ^1^H-^1^H NOESY cross peaks. The NOE correlations between Trp1-C7H with Arg3-CαH/CγH/CδH and Arg3-CβH with Trp1-C5H established a Trp1-C6 and Arg3-Cβ connectivity for the C-C crosslink and suggested an (*S*) configuration for the new stereocenter at Arg3-Cβ (Figure 2). These data show C-C crosslink formation where the ether crosslink is naturally found in darobactin.

**Figure 2:**
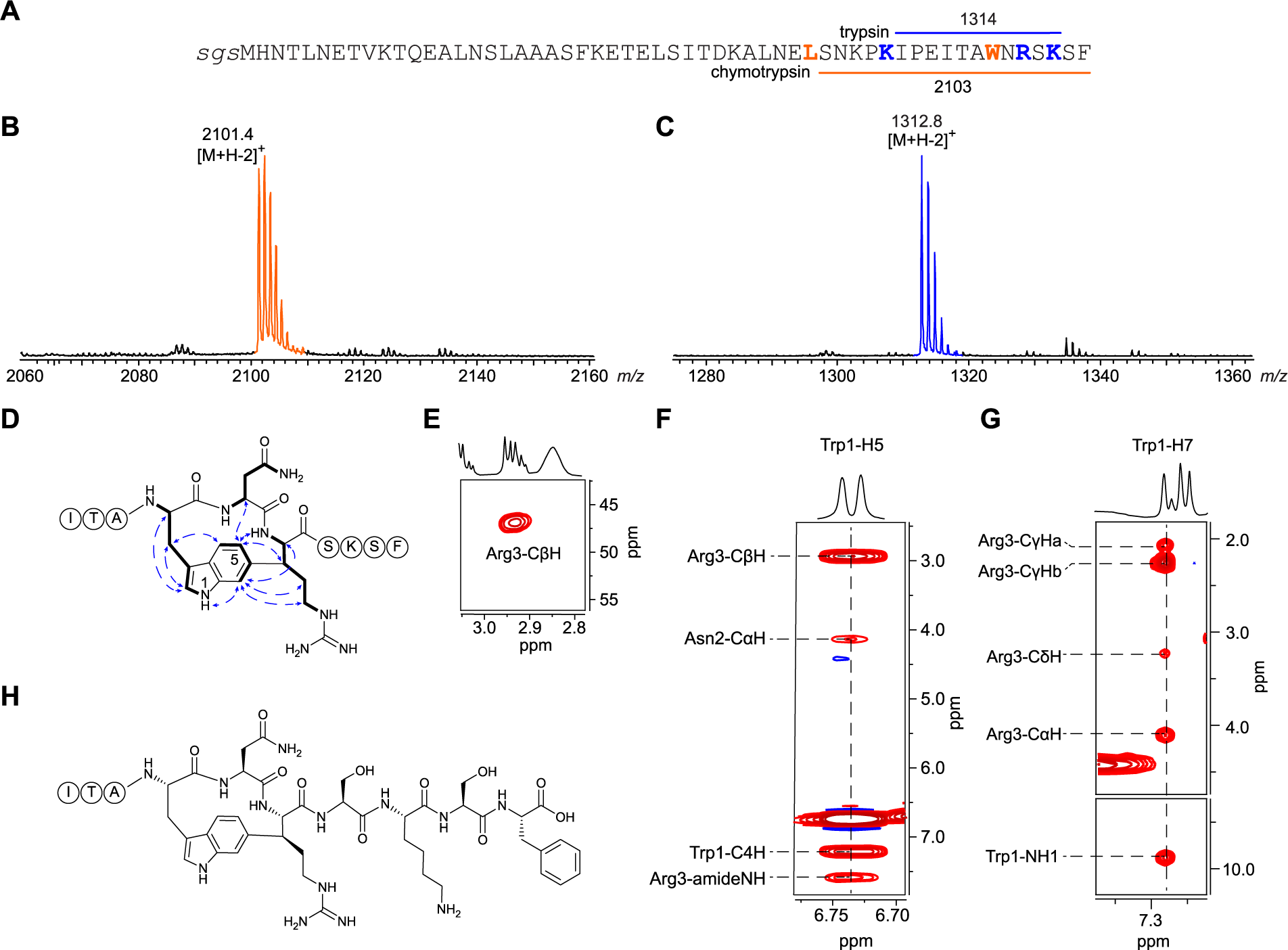
The DarA-W3R variant forms a single Trp1-Arg3 C-C crosslink. (A) Sequence of TEV protease-cleaved MBP-DarA-W3R variant with the pertinent chymotryptic (orange) and tryptic (blue) peptide fragments and masses indicated. (B-C) MALDI-TOF-MS results of (B) chymotryptic and (C) tryptic digests of DarA-W3R. (D) Diagnostic NMR correlations for the W3R variant confirming a Trp1-Arg3 macrocycle (^1^H-^1^H NOESY: blue arrows; ^1^H-^1^H TOCSY: thick black lines). (E) ^1^H-^13^C HSQC cross peak for Arg3-CβH. (F) Trp1-H5 and (G) Trp1-H7 NOE correlations that establish connectivity and stereochemistry. (H) W3R variant structure with stereochemical configurations shown. Complete structural assignment is provided in the supplemental information (Figures S27-S31, Table S2).

Another prevalent modification of DarE-processed DarA variants was a +14 Da mass shift that matched the expected deviation for ether (C-O-C) crosslink formation (Figures S2-S4).^39–41^ Previous work has shown that collision-induced dissociation (CID) of the ether crosslink produces distinguishing daughter ions.^31^ HR-MS/MS spectra consistent with monoether formation at core positions 1 and 3 were observed for DarA variants W3F, W3H, W3Y, K5F, K5H, K5W, and K5Y (Figures S32-S38). However, the CID spectra for variants K5H, K5Y, and K5W were consistent with an isobaric mixture of monoether formation at core positions 3 and 5. UHPLC and HRMS/MS analysis of modified K5H confirmed formation of Trp1-Trp3 (monoether) and Trp3-His5 (monoether) (Figure S36). Analogous to C-C instead of ether installation at core position 1 and 3, an ether can be installed at position 3 and 5 where the C-C crosslink is naturally found in darobactin A.

With these insights into the flexibility of macrocycle linkage type and location, we next investigated if the macrocycle size was adjustable. For macrocycle expansion, 1-2 Gly residues were inserted within each three-residue motif of the DarA core region (i.e., WN**G**WSKSF, WN**GG**WSKSF, WNWS**G**KSF, and WNWS**GG**KSF). For macrocycle contraction, either DarA-Asn2 or -Ser4 were individually deleted. After co-expression with DarE, no variant produced an appreciable extent of modification except for WNWS**G**KSF. While this variant displayed an ether between Trp1 and Trp3, there was no evidence of C-C bond formation, indicating DarE was intolerant of these substrate alterations (Figures S39-S40).

### DarE Generates β-oxo Amino Acids and Formylglycine

Instead of a -2 Da mass shift resulting from C-C crosslink formation or Trp dehydrogenation, HR-MS/MS of DarA-W3S and -K5S localized a -2 Da modification to Ser3 and Ser5, respectively (Figures S41-S42). Given that DarE is retrieved by TIGR03942, it is homologous to anaerobic sulfatase-maturing enzyme and bears 48% sequence similarity to the enzyme anSMEcpe (BAB80341.1).^44–47^ Thus, we suspected that DarE may have generated formylglycine at the newly introduced Ser positions. To explore this possibility, DarE-modified and purified DarA-W3S was reacted with *o*-benzylhydroxylamine. Oxime formation at core position 3 was confirmed by LIFT-MS/MS. (Figures S43-S44). Similar results were obtained with variant K5S; however, the low signal intensity of the oxime adduct prevented MS/MS-based confirmation.

Similarly, HR-MS/MS analysis of multiple Trp1 variants with a +14 Da modification localized the mass increase to Trp3 (Figures S45-S48). Instead of ether formation, we suspected conversion to β-oxotryptophan. Labeling of DarA-W1Y with *o*-benzylhydroxylamine was unsuccessful, potentially owing to reduced electrophilicity of the benzylic position. Treatment of DarE-modified DarA-W1Y with 2,4-dinitrophenylhydrazine (2,4-DNPH), however, resulted in hydrazone formation with LIFT-MS/MS localizing the adduct to core position 3 (Figure S49).^43,48,49^ DarA variants W3N, K5H, K5R, K5W, and K5Y also contained a β-oxotryptophan modification. While β-oxotryptophan resided at core position 3 for variants K5H, K5R, K5W, and K5Y, DarE processing of variant W3N converted Trp1 to β-oxotryptophan (Figures S50-S54). Using HR-MS/MS and reactivity towards 2,4-DNPH, we detected formation of β-oxotyrosine and β-oxohistidine at core position 3 on variants W3Y and W3H (Figures 3, S55-S57, Table S3).^43^ Further, variant K5H displayed β-oxohistidine at position 5.

**Figure 3:**
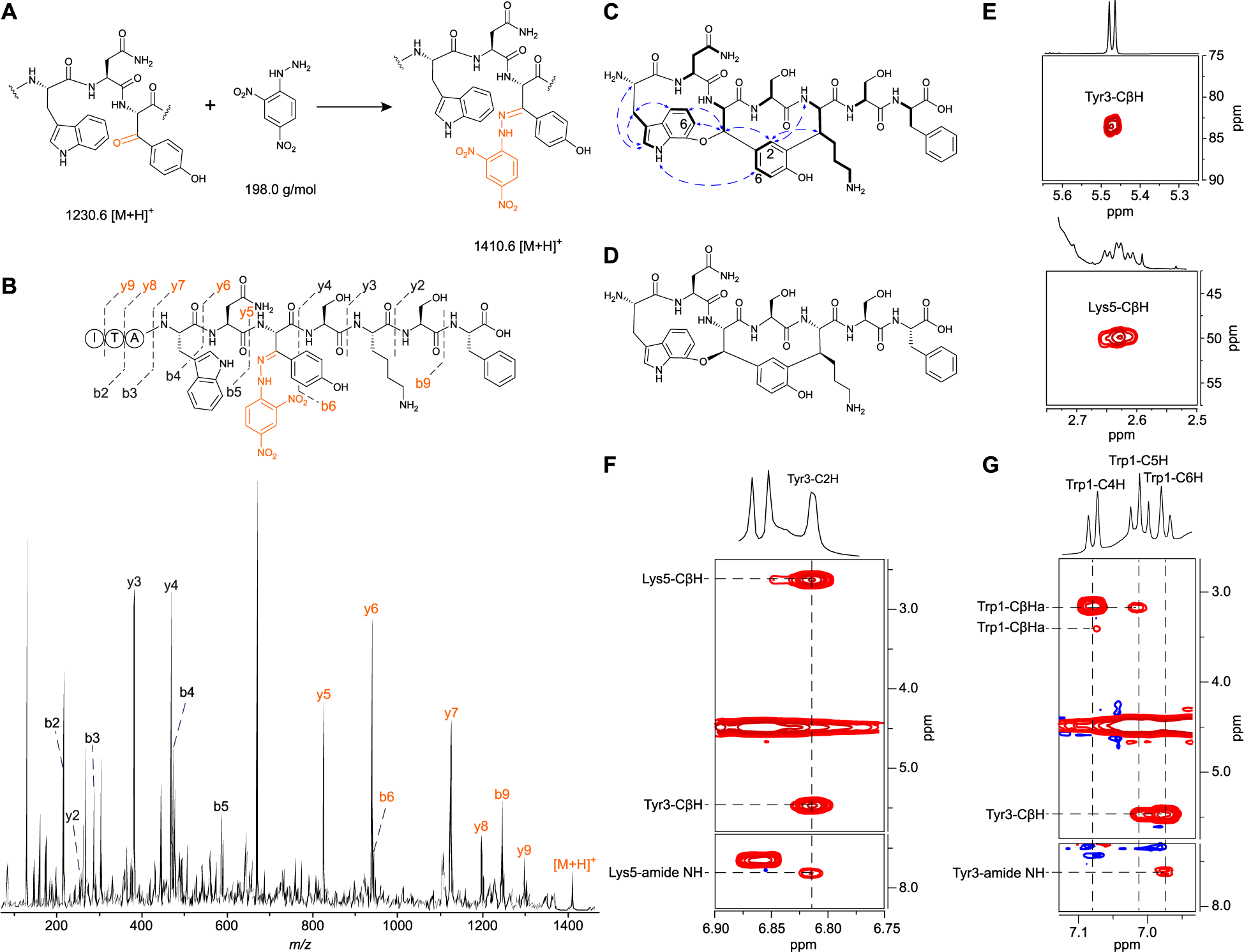
Analysis of a β-oxotyrosine-containing peptide and darobactin W3Y. (A) Hydrazone formation from reaction of β-oxotyrosine with 2,4-DNPH. (B) LIFT-MS/MS analysis of the W3Y hydrazone (*m/z* 1410.7) with observed daughter ions indicated. Orange, ions harboring the hydrazone. (C) Structure of darobactin W3Y (^1^H-^1^H NOESY: blue arrows; ^1^H-^1^H TOCSY: thick black lines). D) Structure of darobactin W3Y with elucidated stereochemical information. (E) ^1^H-^13^C HSQC cross peak for Tyr3- and Lys5-CβH. (F) Tyr3 and (G) Trp1 ^1^H-^1^H NOESY correlations that establish crosslink connectivity and stereochemistry. Complete structural assignment i*s* provided in the supplemental information (Figures S60-S64, Table S4).

### Variant W3Y Contains Ether and C-C Crosslinks

Eight DarA variants exhibited +12 Da modifications, consistent with ether (+14 Da) and C-C (-2 Da) crosslink formation (Figures S2-S4). While no DarA-Trp1 variant gave evidence of ether and C-C crosslink formation, variant W3Y yielded a +12 Da product after co-expression with DarE (HRMS: C_58_H_79_N_13_O_17_^2+^, *calc*. [M+H]^+^: 614.7853; *obs*. [M+H]^+^: 614.7839; error: 2.3 ppm) (Figure S58). The DarA-W3Y product was thus scaled up, digested with proteinase K, purified, and structurally characterized by NMR spectroscopy (Figures 3, S60-S64, Table S4). Using ^1^H-^1^H TOCSY and ^1^H-^13^C HSQC couplings, the characteristic downfield signal, *δ*_CH_ 5.47/83.6, was assigned to Tyr3-CβH and hence was expected to be involved in an ether crosslink. Similarly, Trp1-C7 was found to be monosubstituted, as evidenced by the absence of ^1^H-^1^H TOCSY and ^1^H-^1^H NOESY cross peaks. In addition, the observed NOE correlations between *(i)* Trp1-C5H/C6H and Tyr3-CβH as well as *(ii)* Trp1-NH1 and Tyr3-C6H indicated that the ether formed between Trp1-C7 and Tyr3-Cβ (*R* stereochemistry) similar to other darobactins.^31^ Analogously, ^1^H-^1^H TOCSY and ^1^H-^13^C HSQC resonances highlighted that Tyr3 featured a *meta* and *para*-disubstituted phenyl substructure (*δ*_H6_ 6.81, d ≈ 2.3 Hz) and Lys5-Cβ was monosubstituted (Figures 3, S64-S68). The observation of NOE correlations from Tyr3-C2H to Lys5-CβH/amide NH suggested that the C-C crosslink was between Tyr3-C3 and Lys5-Cβ with (*S*) stereochemistry at Lys5-CβH.

### DarE Installs Fused Ether and C-C Crosslinks on Multiple Lys5 Variants

DarE processing of the Lys5 panel yielded seven DarA variants (i.e., K5R, K5A, K5L, K5Q, K5N, K5F, and K5W) with a +12 Da mass shift (Figure S4). Previous research characterized variants K5R (darobactin D) and K5A (darobactin 16) as having ether and C-C crosslinks akin to darobactin A.^31,36^ HR-MS/MS of variants K5L, K5N, and K5Q identified diagnostic ether crosslink daughter ions and no fragmentation within the _3_WSX_5_ motif, providing evidence for ether and C-C crosslink formation (Figures S65-S67). The yields for darobactins K5L, K5N, and K5Q were too low for structural confirmation by NMR; however, we hypothesize that the ether and C-C crosslinks likely formed with the same connectivity and stereochemistry as in darobactin A.^26^ HR-MS/MS of DarA-K5W (+12 Da) indicated Trp1-Trp3 ether formation but was inconsistent with C-C crosslink formation. Rather, the -2 Da modification was localized onto Trp5, consistent with dhTrp (Figure S68).

### Aromatic Residues at Core Position 5 Yield Two Ether Crosslinks

While the +12 Da species from variant K5F gave very low ion intensity, a robust +28 Da modification was observed suggesting a diether product (Figure S4). Analysis of variant K5F (+28 Da) supported a molecular formula with net addition of two oxygens and net loss of four hydrogens (HRMS: C_63_H_76_N_13_O_17_^+^, *calc*. [M+H]^+^: 1286.5477; *obs*. [M+H]^+^: 1286.5459; error: 1.4 ppm) as well as daughter ions consistent with a fused diether product (Figure S69). Subsequent NMR analysis of proteinase K-digested and UHPLC-purified darobactin K5F confirmed the structure. (Figures 4, S70-S75, Table S5).

**Figure 4:**
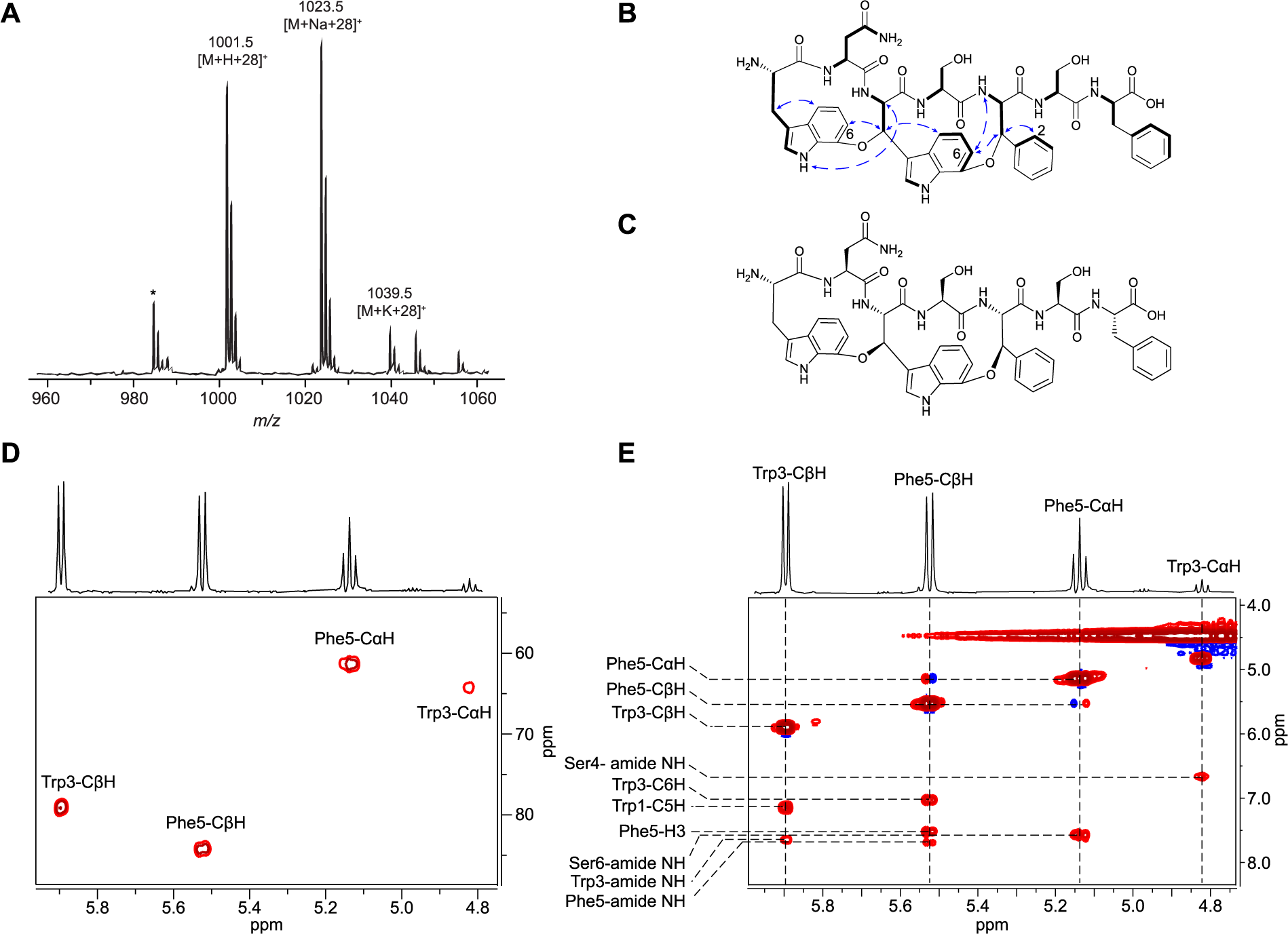
Structural characterization of darobactin K5F. (A) MALDI-TOF mass spectrum of proteinase K-digested DarA-K5F. * indicates laser-induced deamination. (B) Structure of darobactin K5F (^1^H-^1^H NOESY: blue arrows; ^1^H-^1^H TOCSY: thick black lines). (C) Structure of darobactin K5F with elucidated stereochemical information. (D) ^1^H-^13^C HSQC cross peaks for the Trp3 and Phe5 CαH and CβH. (E) ^1^H-^1^H NOESY correlations that established linkage connectivity and stereochemistry. Complete structural assignment is provided in the supplemental information (Figure S71-75, Table S5).

Analysis of the ^1^H-^13^C HSQC spectrum revealed two discriminative doublet resonances (*δ*_CH_ 5.9/79.2, 5.52/84.2), assigned as Trp3-CβH and Phe5-CβH, respectively. Assisted by ^1^H-^1^H TOCSY and ^1^H-^1^H NOESY, the C7 positions of Trp1 and Trp3 were implicated in ether crosslink formation. Specifically, NOE correlations from Trp3-CβH to Trp1-C6H and Phe5-CβH to Trp3-C6H established the locations of the two ether crosslinks. Ultimately, the observed NOEs between Trp1-NH1 and Trp3-CαH alongside Trp3-C6H and Lys5-amide NH elucidated both new stereocenters as having an (*R*) configuration (Figures 4, S72-S75).

HR-MS/MS confirmed that variants K5H, K5W, and K5Y similarly displayed two ether crosslinks after DarE processing (Figures S76-S78). Notably, a variant with a single ether crosslink and a β-oxo amino acid would be isobaric (+28 Da) with the same peptide containing two ether crosslinks. For variants K5W and K5Y, evidence of these isobaric species was observed by HR-MS/MS (Figures S77-S78). In both cases, the fragmentation data always supported ether formation at Trp1-Trp3 but only partial ether formation at core position 3-5. The remainder of the daughter ions assigned the +14 Da modification to core position 5, consistent with β-oxotryptophan and β-oxotyrosine for variants K5W and K5Y, respectively.

### Computational Assessment of Ether Versus C-C Crosslinking

Armed with DarA variant processing data, we next investigated the factors behind DarE crosslink differentiation. The intrinsic preferences towards ether versus C-C formation were studied quantum mechanically on abbreviated models of unmodified DarA. The models (**A_Me_H** and **A_Ph_H**) feature trimmed tripeptides mimicking the amino acid sequences for the second macrocycle (WSX motif): an indole-3-propionic acid, mimicking Trp3; an Ala, mimicking Ser4; and 2-methyl- or 2-phenylethanamine as mimics for β-alkyl (i.e., Lys) and β-aryl (i.e., Phe) amino acids. To eliminate the influence of conformational effects, which are presumably minimal when bound to the DarE active site, models were considered to be in a near-attack conformation similar to that found in atomic-resolution structures of bound darobactin.^30^ For transition state calculations, only the reaction leading to the experimentally observed stereochemistry for each crosslink was considered.

First, we evaluated the stability of Cβ-centered radicals as a function of the substituent (alkyl or aryl) on DarA core position 5. Amino acid radical stability is commonly calculated from X-H bond dissociation energies (BDE, X = C, N, O, S) using the isodesmic reaction method.^50,51^ Following this approach, we analyzed the thermodynamics of the equilibria between the neutral peptide models and the 5’-dAdo• cofactor (**A_R_H** + 5’-dAdo• ⇆ **A_R_**• + 5’-dAdoH), which was exergonic in all cases. The radical **A_Ph_**• was calculated to be ∼9 kcal mol^-1^ more stable than **A_Me_**• (ΔG[**A_Ph_**•] = - 13.7 kcal mol^-1^ vs. ΔG[**A_Me_**•] = -4.5 kcal mol^-1^), resulting in an activation energy for the nucleophilic addition of the radical to Trp3-C6 that is ∼5 kcal mol^-1^ higher (ΔG^‡^[**A_Ph_**•**_TS_6_**] = 19.8 kcal mol^-1^ vs. ΔG^‡^[**A_Me_**•**_TS_6_**] = 14.9 kcal mol^-1^). This energy difference was reflected in the later transition state character of **A_Ph_**•**_TS_6_** (C-C forming bond distance 2.09 Å) compared to **A_Me_**•**_TS_6_** (C–C forming bond distance 2.16 Å). Translation of activation energies into radical half-lives (t_1/2_) at 25 °C yields t_1/2_[**A_Ph_**•] = 37 s, whereas t_1/2_[**A_Me_**•] = 0.009 s (Figure 5). These results suggest that alkyl Cβ-centered radicals rapidly react to form C-C crosslinks. In contrast, aryl Cβ-centered radicals may live long enough to react with molecular oxygen, as proposed for ether crosslink formation in darobactin.^39^

**Figure 5:**
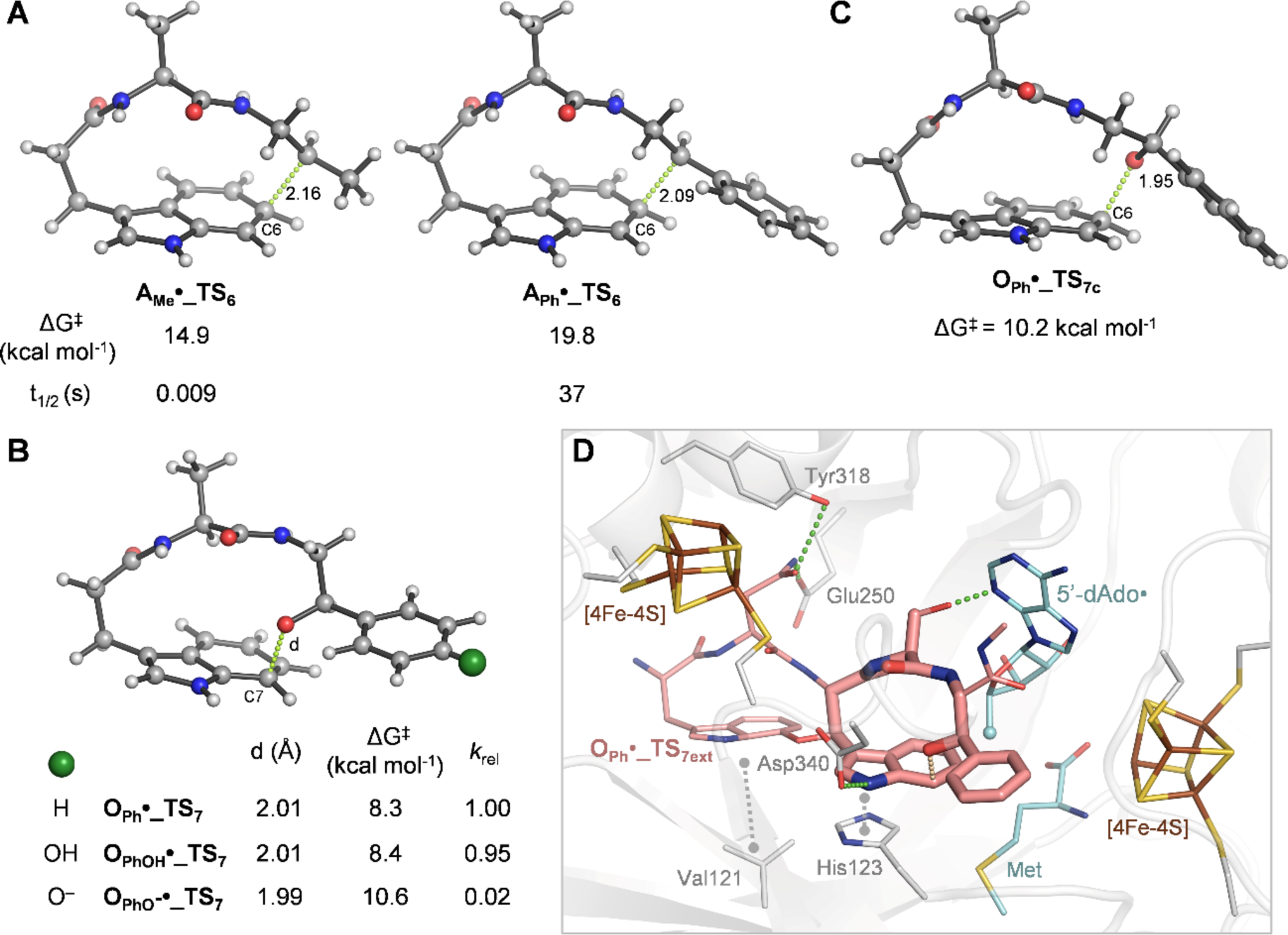
Computational Investigation of Crosslink Formation in Darobactin. (A) QM-optimized transition state structures leading to C-C bond formation at C6 from C-centered alkyl (**A_Me_•_TS_6_**) and aryl (**A_Ph_•_TS_6_**) radicals. The activation barriers (ΔG^‡^) and half-lives at 25 °C (from Eyring’s equation) are given. (B) QM-optimized transition state structures leading to ether formation at C7 from O-centered benzyl (**O_Ph_•_TS_7_**) and *p*-hydroxybenzyl (protonated and deprotonated forms, **O_PhOH_•_TS_7_** and **O_PhO_-•_TS_7_**, respectively). In addition to ΔG^‡^ and half-life values at 25 °C, distances for bond formation are given (Å) and indicated as green dashed lines. (C) Similar to panel B but for **O_Ph_•_TS_7c_**. (D) Molecular docking of a truncated model of darobactin (pink), featuring the QM-optimized transition state for ether crosslinking between Trp3-C7 and alkoxy-Phe5, bound to DarE. 5’-AdoH and Met (cyan, the reacting 5’ methyl carbon is shown as a sphere), [4Fe-4S] centers, and relevant active site residues (grey) are shown in wireframe. H-bonds and van der Waals interactions are shown as green and grey dashed lines, respectively.

Of note, the type of substituent at core position 5 does not significantly affect the stability of the radical intermediates (**A_R_**•**_Int_6_**; ΔΔG_Ph-Me_ = 0.5 kcal mol^-1^) or the final aromatic products (**A_R_**•**_Prod_6_**; ΔΔG_Ph-Me_ = 1.1 kcal mol^-1^), suggesting that the experimentally observed reactivity must be under kinetic control. Therefore, the calculated mechanism is similar to that proposed for the Minisci reaction, which involves a stepwise addition of nucleophilic *C*-centered radicals to heteroarenes to form a C-C bond. Here, we assume that the nucleophilic addition of the radical to the six-membered ring of the indole is rate-limiting.^52,53^ Since the final products are aromatic and thus highly stable (ΔG ≈ -80 kcal mol^-1^ with 5’-dAdo• as the presumed radical hydrogen atom abstractor), a small activation barrier for the subsequent hydrogen abstraction step is expected according to Hammond’s postulate; in fact, attempts to locate transition structures using oxygen or carbon radical hydrogen atom abstractors resulted in barrierless, downhill energy profiles.

We then focused on the ether crosslink, which is hypothesized to form from an *O*-centered alkoxy radical (**O_Ph_**• and **O_Me_**•) resulting from the reaction of Cβ-centered radicals (**A_Ph_**• and **A_Me_**•) with molecular oxygen followed by homolytic cleavage of the O-O bond.^39^ In both cases, the calculated intrinsic activation barriers were significantly lower for aryl over alkyl-substituted alkoxy radicals (ΔG^‡^[**O_Ph_**•**_TS_7_**] = 8.3 kcal mol^-1^ vs. ΔG^‡^[**O_Me_**•**_TS_7_**] = 9.7 kcal mol^-1^) leading to half-lives around 0.1–1 μs (Figures 5, S79). Hence, the fact that ether formation is intrinsically faster than C-C bond formation suggests that the chemoselectivity observed for aromatic amino acids at core position 5 to form ether crosslinks must arise from the higher stability of their initial Cβ-centered radicals.

Ether crosslink formation was also analyzed using phenol and phenoxide substituents to mimic neutral and deprotonated Tyr5 (**O_PhOH_**• and **O_PhO_**-•, respectively). The activation barrier increases for the phenoxide surrogate due to the higher delocalization of the spin density (ΔG^‡^[**O_PhO_-**•**_TS_7_**] = 10.6 kcal mol^-1^) (Figures 5, S80). This increase in the activation barrier is even more dramatic when the environmental dielectric constant in the calculations is changed from water (ε=78) to lower values often used to mimic the polarity inside the active sites of typical enzymes (ε=4), where the basic *O*-centered radical can deprotonate the indole NH (ΔG^‡^[**O_PhO_-**•**_TS_7_’**] = 35.1 kcal mol^-1^) (Figure S80).^54,55^ Altogether, these results support the experimentally observed formation of β-oxotyrosine as a competitive pathway to ether crosslinking.

Finally, the observed change in the regioselectivity of ether versus C-C crosslinking to give reactions at C6 or C7 of Trp, respectively, was also calculated for **O_Ph_**•. In the case of the experimentally observed addition to C7, the backbone displayed an extended β-sheet-like conformation in the transition state (**O_Ph_**•**_TS_7_**). However, for addition to C6, up to three different transition state conformations with different backbone geometries (polyproline-II, β-sheet, and inverse γ-turn) but similar energies were calculated (**O_Ph_**•**_TS_6a-c_**; ΔΔG^‡^_TS6-TS7_ ≈ -0.4 to +1.3 kcal mol^-1^ assuming Curtin-Hammet conditions). These three conformationally distinct pathways converge to the same aromatic product after hydrogen atom abstraction (Figures S81-S86, Table S6-S9). Although the similar energies of the regiodivergent transition states could make the formation of both products competitive, molecular docking calculations show significant differences in the preferred binding poses of those transition states to DarE (computational details provided in methods). **O_Ph_**•**_TS_7ext_** binds to DarE with the highest score and places the alkoxy radical closest to the 5’-dAdoH cofactor. This binding mode is stabilized by multiple interactions, including a hydrogen bond between DarA-Ser4 side chain and the cofactor adenine (Figure 5). Additionally, numerous polar residues of DarE (i.e., Glu250, Arg298, Asp316, and Tyr318) engage the DarA-Asn2 side chain while DarE-Asp340 and -His123 stabilize the DarA-Trp3 side chain through hydrogen bonding and π-stacking. These results suggest that DarE enhances the slight innate preference towards ether crosslinking at C7 by binding DarA in a near-attack, extended conformation.

### Biological Activity Assessment

None of the tested variants (i.e., W3R, darobactin W3Y, and darobactin K5F) were growth suppressive up to 32 µg/mL against *E. coli* K-12 MG1655. However, variant W3R displayed a single crosslink and retained the Ile-Thr-Ala motif from the leader region (Figure 2). Thus, the rigidity and ability to pre-organize into a β-strand for proper alignment with BamA was expectedly impacted.^30^ Conversely, darobactin W3Y contains ether and C-C crosslinks while K5F contains two ether linkages. These variants are considered mature, as W3Y and K5F were bicyclic and did not retain any leader or follower residues (Figures 3-4). We surmised that the altered residues and linkages caused an unfavorable distortion of the β-strand-like structure preventing proper alignment with BamA. To investigate this possibility, the atomic-resolution structure of darobactin A bound to BamA was compared with computationally designed darobactins A, W3Y, and K5F (Figures S87). We focused on the individual and overall macrocyclic sizes which define the locations of atoms involved in H-bonding between darobactin and BamA. Measured from darobactin A bound to BamA (PDB: 7NRE), the Trp1-Cα to Trp3-Cα (**D1**), Trp3-Cα to Lys5-Cα (**D2**), and Trp1-Cα to Lys5-Cα (**D3**) distances are 6.78 Å, 6.81 Å, and 13.52 Å, respectively.^30^ Since we lack atomic-resolution structures of the new darobactin variants bound to BamA, we drew darobactin A *in silico* and performed energy minimization using the Molecular Operating Environment (MOE) software package. The obtained values for **D1-D3** were 6.99 Å, 7.06 Å, and 13.92 Å, respectively, and permit a more direct comparison to the same measurements on the new variants.

Accordingly, darobactins W3Y, and K5F, were drawn *in silico* and energy minimized in MOE based using NMR-determined stereochemical information. While **D1** for darobactin W3Y (6.99 Å) was identical to darobactin A, the **D2** and **D3** values were significantly shorter (6.26 Å and 13.22 Å, respectfully), likely preventing an optimal geometry for interaction with BamA (Figure S87, Table S10). The **D1** and **D2** values for darobactin K5F were in closer agreement with darobactin A (6.87 Å and 6.79 Å); however, **D3** was again significantly shorter (13.37 Å). Another consideration for BamA engagement would be maintenance of the ideal deviation from a fully extended conformation. Thus, we measured a “ruffling angle” that is formed by the imaginary **D1** and **D2** lines at the Cα of core position 3. The crystallographically- and *in silico*-derived darobactin A structures have ruffling angles of 168.4° and 164.4°, respectively. Thus, we interpret similar angular values as being optimal for BamA inhibition. The ruffling angles for the inactive darobactins W3Y and K5F are 172.3° and 156.3°, respectively, which both deviate from the optimal *in silico* angle by 8°. As additional support, darobactin B bound to BamA (PDB: 7P1C) has a crystallographically determined ruffling angle of 166.9° (Table S10). While more data points could strengthen these trends, we encourage the use of these measurements in future darobactin engineering efforts.

### Assessment of the DarA Leader Identifies Critical and Dispensable Regions

While previous work evaluated the mechanism and substrate scope of DarE, the leader region of DarA has not yet been investigated. The darobactin BGC lacks a RiPP Recognition Element (RRE) commonly used for substrate binding; however, RRE-independent RiPPs are well known.^1,56,57^ DarE has an N-terminal extension of ∼60 residues before the rSAM domain, which could serve the same purpose as an RRE. AlphaFold multimer structures composed of multiple darobactin maturases show that the cognate precursor peptides wrap around the N-terminal lobe of the darobactin synthase before entering the active site (Figure S88, Supporting Dataset 1).^58^ Removal of residues 1-60 from DarE abolished processing (Figure S89).

To evaluate the contribution of the leader region to DarE processing, upwards of 30 residues were removed from the N-terminus of DarA with a truncation step size of 5 amino acids. These truncated constructs were then co-expressed with DarE (Figure 6). Unmodified DarA was highly susceptible to endogenous proteolytic degradation in *E. coli*, especially at elevated expression temperatures. To reduce proteolysis, the series of DarA leader region truncations were induced at colder temperatures and slower shaking. Further, we found that retaining the wild-type Gln-Glu-Ile tripeptide follower sequence of DarA aided in detection of unmodified peptides. Removal of the first 5 and 10 residues of DarA had little effect on processing by DarE while removal of the first 15 residues led to a more pronounced processing deficit, as assessed by MS (observation of the +12 Da species). DarE processing of DarA was nearly abolished after removal of the first 20 residues of DarA. This consequential region of DarA contains multiple highly conserved residues, which may be recognized by DarE and thus explain the loss of processing (Figure 6, Supporting Dataset 2). To evaluate the contribution of conserved residues of the DarA leader region, we prepared six individual Ala-substituted variants and performed co-expression with DarE in *E. coli*. Variants S(-32)A, L(-31)A, and F(-27)A displayed significantly reduced processing while the L(- 34)A, S(-28)A, and K(-26)A exhibited much more minor effects on DarA processing (Figure 6). The AlphaFold multimer predicted structure of DarA and DarE places Leu(-31) and Phe(-27) into conserved hydrophobic pockets of DarE while Ser(-32) may engage DarE through polar interactions (Figure S90, Supporting Dataset 1).^58–60^

**Figure 6:**
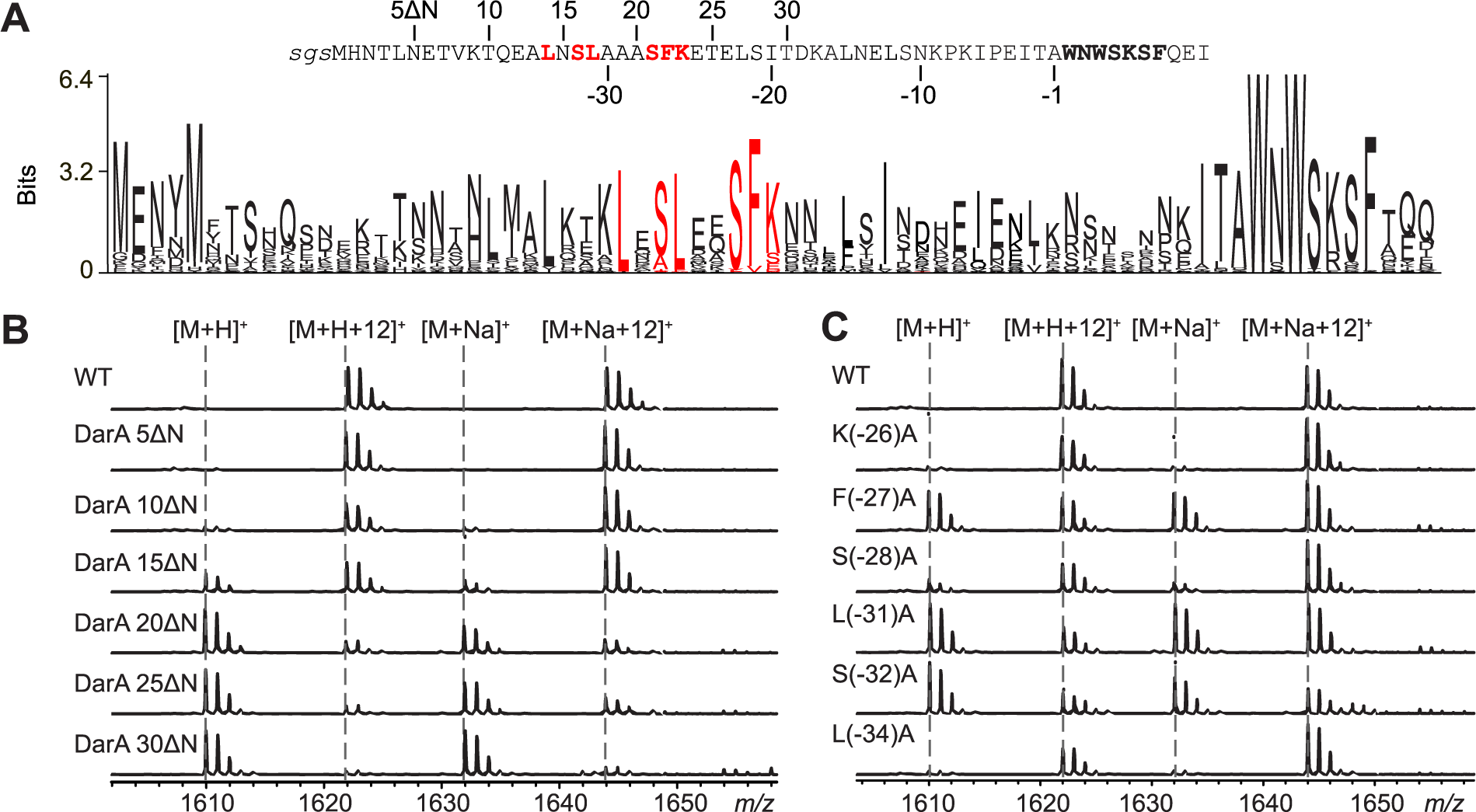
*In vivo* Characterization of DarA Processing with Leader Variants. (A) Sequence of TEV-protease cleaved DarA with truncation numbering from N-terminus (above sequence) and the conventional negative numbering for leader regions (below sequence) starting from the core sequence (bold). A sequence logo of bioinformatically identified darobactin precursor peptides (*n* = 148) is shown. Red, conserved positions evaluated by Ala substitution. (B) MALDI-TOF mass spectra of GluC-digested DarA leader truncations after co-expression with DarE. *m/z* 1610 corresponds to unmodified DarA that includes Ile-Thr-Ala and Qln-Glu-Ile from the leader and follower regions, respectively. M+12 ions correspond to the same sequence that includes the ether and C-C crosslinks of mature darobacin. (C) Same as panel B but for Ala-substituted leader variants after co-expression with DarE.

As illustrated by the sequence logo, the next best conserved portion of the DarA leader region is the Ile-Thr-Ala(-1) motif (Figure 6). To assess the relative importance of this portion of DarA, we prepared several constructs, co-expressed the DarA variants with DarE, and evaluated processing by MALDI-TOF-MS. While the I(-3)A variant resulted in minimal change to DarA processing, the T(-2)A variant was significantly decreased in processing (Figure S91). The T(-2)V variant was fully processed while the T(-2)S variant was diminished, supporting a hydrophobic interaction with DarE at this leader position (Figure S92).^58^ We thus hypothesize that Thr(-2) plays a pivotal role in correctly orientating the DarA core region in the DarE active site. To evaluate this idea, we shifted the register of Thr(-2) by C-terminal insertion of Ala (IT**A**AWNWSKSF), which was devoid of processing after co-expression with DarE. In contrast, insertion of Ala on the N-terminal side of Thr(-2) retains the register (I**A**TAWNWSKSF) and had little to no impact on DarE processing. While atomic-resolution validation will require additional investigation, this work delineates the specific positions of the DarA leader region that engender processing by DarE.

## Discussion

The rSAM enzyme DarE uniquely installs ether (C-O-C) and C-C crosslinks on DarA (Figure 1). To investigate the substrate tolerance and gain insights into the mechanism of DarE and structure-activity relationship of BamA inhibition, we generated 57 variants of DarA with the majority directly targeting the crosslinking sites. Among these variants, 50 contained at least one DarE-dependent modification while 10 variants possessed a bicyclic structure analogous to mature darobactin. A noticeable facet of the different variants we analyzed is the installation of ether crosslinks between aromatic amino acids, while C-C crosslinks formed between an aromatic and a non-aromatic amino acid. Further, the processing outcomes of the DarA variants disclosed here firmly demonstrate that neither crosslink is a prerequisite to form the other, and that crosslinks can be formed at non-canonical core positions (i.e., C-C at positions 1-3 and ether at positions 3-5). Taken together with the results of site-saturation mutagenesis panels at core positions 1, 3, and 5, as well as formylglycine generation in certain variants, we conclude that the DarE-catalyzed reaction is under substrate control, akin to other RiPP-processing enzymes^61–68^ and recently published research on the darobactin family of natural products.^41^

While several DarA variants produced in our work resulted in a fused diether product, the Zhang group identified a natural daropeptide from *Rhizobium sullae* displaying two discrete ether crosslinks.^41^ Under our co-expression conditions, DarE did not tolerate DarA variants that would form a discrete pattern of ether crosslinks, underscoring the nuances of coevolution of darobactin synthases and their cognate substrate peptides. It remains possible that natural darobactin-like compounds possess fused diether rings, given the genomic identification of core sequences that contain aromatic residues at core position 5 (e.g., WNWS**F**RF and WRWS**W**PF).^25^ Sequence analysis of DarA-like core sequences show position 5 is less conserved, and of the 10 bicyclic darobactin variants produced in this study, 8 contained substitutions at position 5 (Figures 6).

In an effort to better understand ether versus C-C crosslink formation, we performed extensive quantum mechanical calculations and enzyme docking studies. Previous theoretical analyses have investigated the rSAM enzyme mechanism.^69–71^ However, our studies illustrate how substrate radical stability and transition state structure correlates with the DarE reaction outcome. These insights should allow the rational engineering of darobactin variants, and perhaps any rSAM-installed cyclophane, to contain a desired crosslink type. Such knowledge could be crucial, given that subtle differences in macrocycle diameter and geometry can vastly perturb the resulting biological activity.

Many natural and artificial darobactin analogs have been reported. These include an intriguing case where Trp1 is naturally brominated, which displays improved activity towards clinically relevant pathogens.^42^ Although none of the variants we tested were growth-suppressive to *E. coli*, the perturbations to the macrocyclic structure of darobactin A provide orthogonal data on the requisite features of high affinity binding to BamA. This includes individual macrocycle diameters of ∼7.0 Å and overall fused macrocycle width of just under 14.0 Å when energy minimized. A peptide backbone “ruffling angle” was also considered to evaluate the β-strand conformation of darobactin to allow optimal engagement with BamA. Our analysis indicated angles near 165° are ideal. Future efforts may consider producing an N-terminal extension of darobactin that would enter the lumen of BamA (akin to dynobactin) while maintaining a fused ring pattern to retain conformational rigidity and thus high affinity for the lateral gate.^25^

Another benefit resulting from our DarA mutational analysis is additional information on the enzymatic mechanism of ether and C-C crosslink formation. Specifically, our workflow was designed to permit the characterization of biosynthetically immature products. In a previously proposed ether crosslink mechanism, an alkoxy radical intermediate on Trp3-Cβ is postulated to form.^39^ When conditions are inadequate for ether formation, the alkoxy radical may form β-oxotryptophan. We observed multiple DarA variants containing β-oxotryptophan, β-oxotyrosine, or β oxohistidine, supporting oxygen addition to Cβ of the C-terminal residue undergoing crosslink formation. A particularly informative DarA variant was W3Y, which formed three distinct products upon co-expression with DarE: *(i)* Trp1-Tyr3 ether and Tyr3-Lys5 C-C (fused bicyclic), *(ii)* Trp1-Tyr3 ether (single macrocycle), and *(iii)* β-oxotyrosine at position 3 (no macrocycle). Our interpretation is that each product originated from an initial alkoxy radical that proceeded down diverging pathways to what was ultimately detected. Clearly, DarE struggles to process the W3Y variant, and the presumed reduced rate of processing led to enhanced detection of biosynthetically immature and/or shunt products. On one hand, these results suggest that the ether ring may form first; however, the fact that DarE can install the crosslinks independently prevents a firm conclusion from being made at the present time. Irrespective of ring installation order, our data oppose a mechanism for DarE where Trp1 is first hydroxylated.^39^ Instead, our data align well with alkoxy radical formation at Trp3 during ether crosslink mechanism.

While the above analysis provides new insights into the core region of DarA, the processing of canonical RiPP precursor peptides is driven by interaction with the leader region. While DarE does not possess an RRE, the first ∼60 amino acids of DarE were predicted to be involved in precursor binding according to AlphaFold multimer predictions and lack of DarA processing upon ablation of DarE(1-60).^56,58^ The DarA leader peptide was subjected to a series of truncations, which found that the first 15 amino acids of DarA are largely dispensable for DarE processing. Mutational analysis revealed several residues in the central region of the leader, specifically positions (-32), (-31), and (-27) that were crucial for processing. Similarly, Thr(-2) was identified as a key residue for DarA processing. Intriguingly, lasso peptides, an unrelated RiPP class, also have a conserved Thr(-2) residue that is highly conserved and critical for biosynthetic maturation.^72^ This may point to a broader trend of β-branched aliphatic residues immediately before the core region to facilitate active site engagement. For this trend to be confirmed or refuted, additional investigation is required.

## Conclusions

Research into the mechanisms and capabilities of rSAM enzymes is rapidly progressing. In this study, we uncover the fundamental principles of crosslink formation by the darobactin synthase DarE. Like other RiPP biosynthetic enzymes, DarE is under substrate control with ether and C-C formation determined by the identity of the residues being crosslinked. The computational aspects of this study determined that aromatic residues enable ether over C-C crosslink formation due to increased radical stability and longevity that permits reaction with molecular oxygen. Further, molecular docking and transition state structural calculations provide rationale for the different connectivity for the ether (Trp-C7) and C-C (Trp-C6) crosslinks. Characterized immature products provide further mechanistic insight into DarE and directly implicate oxygen addition at the Cβ position of residue 3 in the macrocycle forming motif. Along with the mechanism, substrate-enzyme binding was investigated. DarE binds to DarA in an RRE-independent manner, and our work uncovers multiple residues critically involved in the interaction. In summary, the combined experimental and computational analyses disclosed within explain the principles underlying crosslink differentiation and should enhance rational engineering efforts on darobactin-like scaffolds.

## Supporting Information

Experimental methods and supporting figures (S1-S94) and tables (S1-S10) (PDF). Supplemental Dataset 1 contains AlphaFold multimer models (ZIP of PDB coordinate files) Supplemental Dataset 2 contains a list of bioinformatically identified darobactin precursor sequences (XSLX).

## Author Information

## Corresponding Author

Douglas A. Mitchell - Carl R. Woese Institute for Genomic Biology, University of Illinois at Urbana-Champaign, Urbana, Illinois 61801, USA; Department of Chemistry and Department of Microbiology, University of Illinois at Urbana-Champaign, Urbana, Illinois 61801, USA. https://orcid.org/0000-0002-9564-0953 Email

## Authors

**Austin M. Woodard -** Carl R. Woese Institute for Genomic Biology, University of Illinois at Urbana-Champaign, Urbana, Illinois 61801, USA

**Francesca Peccati** – Center for Cooperative Research in Biosciences (CIC bioGUNE), Basque Research and Technology Alliance (BRTA), Bizkaia Technology Park, Building 801A, 48160 Derio, Spain

**Claudio D. Navo** – Center for Cooperative Research in Biosciences (CIC bioGUNE), Basque Research and Technology Alliance (BRTA), Bizkaia Technology Park, Building 801A, 48160 Derio, Spain

**Gonzalo Jiménez-Osés** – Center for Cooperative Research in Biosciences (CIC bioGUNE), Basque Research and Technology Alliance (BRTA), Bizkaia Technology Park, Building 801A, 48160 Derio, Spain; Ikerbasque, Basque Foundation for Science, 48013 Bilbao, Spain

## Notes

The authors declare no competing financial interest.

## Acknowledgments

We thank Sangeetha Ramesh and Timothy Precord for HR-MS/MS data acquisition and Mayuresh Gadgil and Hamada Saad for UHPLC and NMR assistance. We further acknowledge helpful discussions with Prof. David Sarlah, Taras Pogorelov, and Lingyang Zhu. Shravan Dommaraju and Hamada Saad assisted in manuscript proofreading. This work was supported in part by the National Institutes of Health (R01GM123998 to DAM) and by MCIN/AEI/10.13039/501100011033 (PID2021-125946OB-I00 to G.J.O., CEX2021-001136-S to CIC bioGUNE and IJC2020-045506-I to F.P.)

