## Supplementary material for "Benzylic Radical Stabilization Permits Ether Formation During Darobactin Biosynthesis": SI document

### Materials and Methods

**General materials and methods.** Unless otherwise noted, chemical reagents were purchased from Sigma-Aldrich, Gold Biotechnology, or Thermo Fisher Scientific. Reagents for molecular biology experiments, including plasmid cloning and manipulation, were purchased from New England Biolabs. Oligonucleotide primers were purchased from Integrated DNA Technologies (IDT). Synthetic genes were purchased from Twist Bioscience. DNA Spin Columns were purchased from Epoch Life Sciences. *E. coli* DH5α and BL21(DE3) were used for plasmid maintenance and protein overexpression, respectively. An Avanti JXN-26 centrifuge (Beckman Coulter) was used for cell harvesting, and a Sorvall RC6 Plus centrifuge (Thermo Fisher Scientific) was used for lysate centrifugation during large-scale protein purifications. Sanger DNA sequencing was performed at the UIUC Core Sequencing Facility in the Roy J. Carver Biotechnology Center (University of Illinois at Urbana-Champaign). Matrix-assisted laser desorption ionization time-of-flight mass spectrometry (MALDI-TOF-MS, and MALDI-LIFT-TOF-TOF-MS) data were collected with an UltrafleXtreme instrument (Bruker Daltonics) at the Mass Spectrometry Lab (University of Illinois Urbana-Champaign) while high-resolution and tandem mass spectrometry (HR-MS/MS) data were collected on an Orbitrap Fusion ESI-MS using an Advion Triversa Nanomate 100 at the Proteomic Core Facility in the Roy J. Carver Biotechnology Center (University of Illinois Urbana-Champaign).

**Molecular biology techniques.** A modified pET28b plasmid (pET28-MBP) providing fusion of a maltose-binding protein (MBP) tag to the N-terminus of the protein of interest was used. This construct also provides a tobacco etch virus (TEV) protease cut site C-terminal of the MBP tag to enable facile removal of MBP. *E. coli* codon-optimized genes for both *darA* (precursor peptide) and *darE* (rSAM enzyme) (Twist Bioscience) were cloned into pET28-MBP by Gibson assembly resulting in MBP-*darA* and untagged *darE*. The *darA* gene was further modified to alter the first codon of the follower into a stop codon (DarA(Q8*)E). Mutants of *darA* and *darE* were introduced using the QuikChange method using the listed primers (**Table S1**). Unmodified DarA was obtained by BamHI and HindIII restriction cloning of the *darA* gene from the pET28-MBP-DarA(Q8*)E plasmid to a pET28-MBP plasmid without DarE. DarA leader peptide truncations were performed by Gibson assembly in the pET28-MBP-DarAE plasmid. All plasmid mutations were confirmed by Sanger sequencing.

**NNK mutagenesis panel generation.** Resulting nicked plasmids from site-directed mutagenesis with NNK primers (**Table S1**) were transformed into DH5α, plated (LB, 50 μg/mL kanamycin), and grown at 37 °C for 20-24 h. All colonies were transferred into an overnight culture of 5 mL LB (50 μg/mL kanamycin) and grown at 37 °C for 14-16 h using a tube roller. The polyclonal plasmids were then purified by miniprep (QIAprep Spin Miniprep Kit, Qiagen).

**Small scale protein overexpression and purification.** Polyclonal NNK panel for the codons corresponding to Trp1, Trp3, and Lys5 of DarA were each transformed into BL21(DE3). From this plate, 96 different colonies were inoculated into 10 mL of LB (50 μg/mL kanamycin) at 37 °C while in a rotary tube roller. Once the cultures reached $OD_{600}$ 0.4-0.8, they were placed into the cold room for 15-20 min, and protein expression was induced by adding 10 μL 0.5 M Isopropyl β-D-1-thiogalactopyranoside (IPTG) for 16-18 h at 31 °C in a rotary tube roller.

After overexpression, the cells of each culture were harvested by centrifugation at 4000 × g (Beckman-Coulter Allegra X-15R Centrifuge) and washed with 300 μL of phosphate buffered saline (PBS). The resulting cell pellets were resuspended in 350 μL of Ni-NTA lysis buffer (6 M guanidinium hydrochloride (Gu-HCl), 50 mM Tris, 0.5 M NaCl, 15 mM imidazole, 2.5 % (v/v) glycerol, 0.1 % (v/v) Triton X-100, pH 7.5) supplemented with 4 mg/mL lysozyme, 2 μM benamidine HCl, 2 μM leupeptin, 2 μM E64, and 0.1 mM phenylmethylsulfonyl fluoride and equilibrated on ice for 15 min. The resuspended cells were then flash frozen in liquid nitrogen for 2 min and then placed into a water bath for 10 min to thaw. This process was repeated twice more, and after the final freeze-thaw cycle, the lysate was processed by centrifugation at 18,200 × g for 20 min at 4 °C in a tabletop microcentrifuge (Eppendorf 5430R). The resulting supernatant was transferred to a clean 96-well plate on ice.

For protein purification, a HisPur Ni-NTA 96-well spin plate was used according to the manufacturer's instructions (Thermo Fisher). All steps were performed at 4 °C unless otherwise stated. The Sorvall ST 40R was used for all centrifugation steps. Briefly, the plate assembly was subjected to centrifugation for 1 min at 1000 × g to remove the storage solution. Each well was equilibrated with 200 μL Ni-NTA lysis buffer (containing 6 M Gu-HCl) and subjected to centrifugation for 1 min at 1000 × g. The flowthrough was discarded, and this step was repeated twice more. Clarified lysate from each culture was loaded into the different wells of the spin plate, and the plate was equilibrated at 4 °C for 15 min. The plate assembly was again subjected to centrifugation for 1 min at 1000 × g, and the flow through was discarded. The resin was washed consecutively with 1 x 200 µL/well of Ni-NTA lysis buffer (with 6 M Gu-HCl), 1 x 200 µL/well of Ni-NTA lysis buffer (no Gu-HCl), and 6 x 200 µL/well of wash buffer (50 mM Tris, 0.5 M NaCl, 50 mM imidazole, pH 7.5) with centrifugation at 1000 × g for 1 min in between each step. Flow through was discarded after each centrifugation step. Protein bound to the resin was eluted by the addition of 2 x 100 µL Ni-NTA elution buffer (50 mM Tris, 0.3 M NaCl, 250 mM imidazole, pH 7.5) to each well. The plate assembly was placed at room temperature for 2 min then processed by centrifugation. A clean 96-well plate was used to collect the elution fractions.

**NNK panel modification analysis.** Eluted protein was used immediately in protease digests. TEV protease was added to a final ratio of approximately 1:10 TEV: MBP-DarA and allowed to proceed at room temperature for 18-20 h. After TEV digestion, all samples were transferred to a new PCR-capable 96 well plate and heated at 80 °C for 10 min, then processed by centrifugation at 4700 × g for 15 min to precipitate TEV protease and cleaved MBP. The supernatant was transferred to 0.6 mL tubes, flash frozen in liquid nitrogen, and lyophilized to dryness. Samples were individually resuspended in 100 µL of deionized water, from which a portion was diluted with acetonitrile to a final concentration of 50%. Samples were mixed with sinapinic acid (SA) matrix for MALDI-TOF-MS to determine the identity of the DarA variant in each sample. To validate DarA identities, colony PCR (primers listed in **Table S1**) was used to amplify the *darAE* genes from the colonies used in the expressions. The resulting DNA fragments were sequenced by Sanger sequencing.

Samples containing unique DarA variants were further digested with endoproteinase GluC (1:10 ratio of enzyme: substrate) according to the manufacturer's specifications (NEB). Samples were then desalted with a C18 Ziptip (EMD Millipore), eluted into 60% acetonitrile, and mixed with α-cyano-4-hydroxycinnamic acid (CHCA) or super DHB (SDHB) matrix for MALDI-TOF-MS analysis.

**HR-ESI-MS/MS analysis of modified peptides.** GluC-digested samples, prepared as described above, were submitted directly for HR-ESI-MS/MS analysis. The samples were diluted 1:4 with an 80% acetonitrile / 1% acetic acid solution and directly infused into a ThermoFisher Scientific Orbitrap Fusion ESI-MS using an Advion TriVersa Nanomate 100. The MS was calibrated and tuned with Pierce LTQ Velos ESI Positive Ion Calibration Solution (ThermoFisher). The MS was operated using the following parameters: mass range, 100-2000 m/z; resolution, 120,000; isolation width (MS/MS), 0.5-1 m/z; normalized collision energy (MS/MS), 35; activation q value

(MS/MS), 0.4; activation time (MS/MS), 30 ms. Fragmentation was performed using CID at 30% or 50%. Data analysis was conducted using the Qualbrowser application of Xcalibur software (ThermoFisher Scientific) and IPSA.[1]

**Generation of DarA variants absent from the initial NNK panels.** Variants not identified in the initial NNK panels were generated by site-directed mutagenesis as described earlier (primers listed in **Table S1**). Sanger sequencing verified the mutant constructs. Small-scale expression and purification of these variants followed identically to that described above for the NNK panel.

**Large-scale darobactin variant overexpression and purification.** Plasmids containing the DarA variants of interest (W3R, K5F, and W3Y) were transformed into chemically competent BL21(DE3) *E. coli* cells and grown at 37 °C for 20-24 hours after plating the outgrowth. Single colonies were inoculated into 10 mL LB (50 μg/mL kanamycin) and grown overnight at 37 °C in a rotary tube roller. Large cultures (LB, 50 μg/mL kanamycin) were inoculated 1:200 with overnight culture and were placed into the incubator shaker at 180 rpm, 37 °C, until an $OD_{600}$ between 0.5-0.6 was reached. At that point, the cultures were moved into the cold room for 30 min then induced with 0.5 mM IPTG overnight (180 rpm, 30 °C).

Following overexpression, cells were harvested by centrifugation at 4000 × g for 20 min at 4 °C. The cells were then washed with PBS then resuspended in 30 mL Ni-NTA lysis buffer (15 mM imidazole, 0.5 M NaCl, 2.5% (v/v) glycerol, 0.1% (v/v) Triton-100, 50 mM Tris, pH 7.5), 4 mg/mL lysozyme, 2 μM benzamidine HCl, 2 μM leupeptin, 2 μM E64, and 0.1 mM phenylmethylsulfonyl fluoride. All samples were equilibrated at 4 °C for 20 min, with agitation, then sonicated for 1 min total (2 s on, 8 s off x 30 cycles) while on ice. After sonication, the samples were equilibrated at 4 °C under agitation for at least 10 min.

Lysed and homogenized samples were processed by centrifugation via a Sorvall RC6 Plus (Thermo Electron Corporation) at 39,000 × g at 4 °C for 60 min to remove cellular debris and other insoluble material. The resulting supernatant was loaded onto an His-Pur Ni-NTA resin (2 mL resin/L culture, pre-equilibrated with Ni-NTA lysis buffer) and equilibrated at 4 °C for 45 min under slight agitation. The lysate was allowed to flow through the column by gravity. The column was washed with 4 x 5 column volumes of Ni-NTA wash buffer (30 mM imidazole, 0.5 M NaCl, 50 mM Tris, pH 7.5). The purified peptide was eluted by gravity with 100 mL Ni-NTA elution buffer (250 mM imidazole, 0.3 M NaCl, 50 mM Tris, pH 7.5) 10 min after adding the Ni-NTA elution buffer to the column. Eluted peptide was concentrated and buffer exchanged into protein storage buffer [50 mM 4-(2-hydroxyethyl)-1-piperazineethanesulfonic acid (HEPES), 300 mM NaCl, 2.5% (v/v) glycerol, 0.5 mM tris(2-carboxyethyl)phosphine (TCEP)] with Amicon Ultra-15 10 kDa molecular weight cut-off (MWCO) centrifugal filters (Millipore-Sigma). Purified and buffer exchanged peptide was quantified by A280 on a Nanodrop 2000 (Thermo Scientific), aliquoted, flash frozen with liquid nitrogen, and stored at -80 °C until later use.

To remove the MBP tag on purified DarA variants, TEV protease was added to a final ratio of approximately 1:100 enzyme: protein and allowed to proceed at room temperature for 20-24 h. After TEV digestion, the samples were heated at 80 °C for 10 min while shaking in a ThermoMixer C (Eppendorf), then subjected to centrifugation in a tabletop centrifuge at 16,100 × g for 10 min to ppt and pellet TEV and cleaved MBP (Centrifuge 5430R; Eppendorf). The supernatant was transferred to a new tube where either endoproteinase GluC (~1:1000 ratio of enzyme: DarA) or proteinase K (1:20 enzyme: DarA) was directly added. The digests were allowed to proceed for at least 24 h at 37 °C. In preparation for Ultra High-Performance Liquid Chromatography (UHPLC), the GluC and proteinase K digests were concentrated to dryness (SpeedVac SPD120, Thermo Scientific).

**Expression and purification of DarA leader peptide variants.** Plasmids containing the *darA* mutants of interest were transformed into chemically competent BL21(DE3) *E. coli* cells and grown at 37 °C for 20-24 h after plating the outgrowth. Single colonies were inoculated into 5 mL LB (50 µg/mL kanamycin) and grown overnight at 37 °C in a rotary tube roller. 100 mL of LB (50 µg/mL kanamycin) was inoculated 1:250 with overnight culture and was placed into the incubator shaker at 180 rpm, 37 °C, until an $OD_{600}$ between 0.5-0.6 was reached. At that point, the cultures were moved into the cold room for 20 min then induced 0.5 mM IPTG overnight (80 rpm, 18 °C).

The purification of the leader peptide variants was identical to that of the other variants except amylose resin (NEB) was used instead of Ni-NTA. MBP lysis buffer (50 mM tris, 0.5 M NaCl, 2.5% (v/v) glycerol, 0.1% Triton x-100, pH 7.5) was used for cell pellet resuspension. For the amylose purifications, the resulting supernatant after lysis, sonication, and centrifugation, was loaded onto amylose resin (5 mL resin/L culture, pre-equilibrated with MBP lysis buffer). The lysate was allowed to flow through the column by gravity. The column was washed with 10 column volumes MBP wash buffer A (50 mM tris, 0.5 M NaCl, 2.5% (v/v) glycerol, 0.1% Triton x-100, pH 7.5) and 5 column volumes MBP wash buffer B (50 mM tris, 0.5 M NaCl, 2.5% (v/v) glycerol, pH 7.5) The purified peptide was eluted by gravity with 15 mL MBP elution buffer (50 mM tris, 0.3 M NaCl, 10 mM maltose, 2.5% (v/v) glycerol, pH 7.5).

Eluted peptide was concentrated, and buffer exchanged into protein storage buffer (50 mM HEPES, 300 mM NaCl, 2.5% (v/v) glycerol, 0.5 mM TCEP) with Amicon Ultra-15 30 kDA MWCO centrifugal filters (Millipore-Sigma). Purified and buffer exchanged peptide was quantified by A280 on a Nanodrop 2000 (Thermo Scientific), aliquoted, flash frozen with liquid nitrogen, and stored at -80 °C until later use.

**HPLC methods.** All HPLC methods were performed using a Vanquish UHPLC System (ThermoFisher Scientific). Either a Hypersil C18 semipreparative column (250 × 10 mm), an Accucore C18 analytical column (150 × 4.6 mm), or a pentyl-fluorophenyl (PFP) analytical column (150 x 4.6 mm) was used for all purifications. Mobile phases of A: $H_2O$ + 0.1% formic acid (FA) and B: acetonitrile + 0.1% FA were used unless otherwise indicated. UV-absorbances from 220-400 nm were monitored over the full course of all methods. Flow rates were 3 mL/min for the semipreparative column or 1.5 mL/min for the analytical columns unless otherwise specified. Any insoluble material after sample resuspension was removed by centrifugation at 17,000 × g for 3 min.

*DarA-W3R*

Lyophilized GluC-digested DarA-W3R was resuspended in an appropriate volume of 0.1% formic acid and injected onto the semipreparative column. The following gradient was used for purification: 5% B at $t$ = 0 min, 5% B at $t$ = 5 min, 30% B at $t$ = 40 min, 90% B at $t$ = 45 min, 90% B at $t$ = 50 min, 5% B at $t$ = 51 min. The peptide eluted fractions containing the GluC-digested modified W3R peptide were pooled and concentrated by to dryness (SpeedVac SPD120, Thermo Scientific). This initial purified sample was resuspended with an appropriate volume of 0.1% formic acid and injected onto an Accucore C18 analytical column for a second-stage purification. Multiple injections and fraction collection were both automated. The following gradient was used for purification: 5% B at $t$ = 0 min, 5% B at $t$ = 3 min, 12% B at $t$ = 18.4 min, 95% B at $t$ = 20 min, 95% B at $t$ = 25 min, 5% B at $t$ = 27 min. Fractions containing the modified W3R peptide were combined and lyophilized to dryness for NMR analysis.

*Darobactin W3Y*

Lyophilized proteinase K-digested DarA-W3Y was resuspended in an appropriate volume of 0.1% formic acid and was purified with the semipreparative column. The following gradient was used for purification: 5% B at $t$ = 0 min, 5% B at $t$ = 5 min, 20% B at $t$ = 25 min, 95% B at $t$ = 30 min, 95% B at $t$ = 37 min, 5% B at $t$ = 39 min. Fractions containing darobactin W3Y were pooled and concentrated to dryness (SpeedVac SPD120, Thermo Scientific). This initial purified sample was resuspended with an appropriate volume of 0.1% formic acid and injected onto the Accucore C18 analytical column at a flow rate of 2.0 mL/min for second-stage purification. Multiple injections and fraction collection were both automated. The following gradient was used for purification: 7% B at $t$ = 0 min, 7% B at $t$ = 3 min, 12.5% B at $t$ = 30 min, 95% B at $t$ = 31 min, 95% B at $t$ = 35 min, 5% B at $t$ = 36 min. Fractions containing darobactin W3Y were combined and lyophilized to dryness for NMR analysis.

*Darobactin K5F*

Lyophilized proteinase K-digested DarA-K5F was resuspended in an appropriate volume of 0.1% formic acid and injected onto the semipreparative column. The following gradient was used for purification: 10% B at $t$ = 0 min, 10% B at $t$ = 5 min, 45% B at $t$ = 40 min, 90% B at $t$ = 42 min, 90% B at $t$ = 47 min, 5% B at $t$ = 49 min. Fractions containing darobactin K5F were pooled and concentrated to dryness (SpeedVac SPD120, Thermo Scientific). This initial purified sample was again resuspended with an appropriate volume of 0.1% formic acid and injected onto the PFP analytical column for a second stage of purification. Multiple injections and fraction collection were both automated. The following gradient was used for purification: 25% B at $t$ = 0 min, 25% B at $t$ = 3 min, 45% B at $t$ = 19 min, 95% B at $t$ = 21 min, 95% B at $t$ = 26 min, 5% B at $t$ = 28 min. Fractions containing darobactin K5F were combined and lyophilized to dryness for NMR analysis.

*DarA-W1N and -W1D*

Acetonitrile was added to the DarA-W1N and -W1D GluC digests to a final concentration of 50% and was processed by centrifugation at 17,000 × g for 10 min to remove insoluble material. The sample was concentrated to 250 μL and injected onto an Accucore C18 analytical column. Fraction collection was automated. The following gradient was used for purification: 5% B at $t$ = 0 min, 5% B at $t$ = 5 min, 20% B at $t$ = 30 min, 30% B at $t$ = 35 min, 95% B at $t$ = 36 min, 95% B at $t$ = 41 min, 5% B at $t$ = 42 min. Fractions containing the modified peptides were collected.

*DarA-K5H*

Purified DarA-K5H (co-expressed with DarE) was digested with TEV and GluC as described above. Acetonitrile was added to the DarA-K5H GluC digest to a final concentration of 40% and was processed by centrifugation at 17,000 × g for 10 min to remove insoluble material. The entire sample was injected onto an Accucore C18 analytical column. Fraction collection was automated. The following gradient was used for purification: 15% B at *t* = 0 min, 15% B at *t* = 3 min, 50% B at *t* = 40 min, 95% B at *t* = 41 min, 95% B at *t* = 46 min, 15% B at *t* = 47 min. Fractions containing the modified peptides were collected.

**NMR methods.** All NMR experiments were conducted on a Bruker NEO 600 MHz spectrometer with a BBO Prodigy Probe. Purified darobactin variants were resuspended in 700 µL of a 66% Mili-Q $H_2O$: 32% $CD_3CN$: 2% $DCO_2D$ solution. Water suppression (WET) methods were used for all NMR experiments. One-dimensional $^1$H spectra, and two-dimensional $^1$H-$^1$H TOCSY, $^1$H-$^1$H NOESY, and $^1$H-$^{13}$C HSQC spectra were collected. Data was analyzed with MestReNova version 14.3.0.

**Bioactivity testing.** Microbroth dilution assays were performed to determine the minimum inhibitory concentration (MIC) of isolated darobactin variants against *E. coli* MG1655 and *E. coli* MG1655 BamA mutants 2 and 3.[2] Single colonies of each *E. coli* strain were inoculated into 5 mL Mueller-Hinton II media (MHII) and grown overnight at 37 °C. These cultures were diluted 100-fold into 100 mL MHII media and grown to the mid-exponential phase ($OD_{600}$ = 0.3-0.5). At that point, the cultures were diluted to $OD_{600}$ = 0.001, and 98 µL of each culture was added into round bottom 96-well plates for bioactivity testing. Darobactin variant stock solutions (W3R, W3Y, K5F; 1.6 mg/mL) were serially diluted (two-fold), and 2 µL was added into the corresponding wells for final concentrations ranging from 32 µg/mL to 0.5 µg/mL. Native darobactin A and kanamycin were used as positive controls with the same preparation described above. Each molecule was tested in duplicate. The plate was sealed and equilibrated at 37 °C without shaking for 24 h. MICs are reported as the concentration at which no visible bacterial growth was observed.

**Reactive carbonyl labeling.** A 2.4 µg/µL stock solution of 2,4-dinitrophenylhydrazine (2,4-DNPH) in 50% methanol: 50% phosphoric acid was prepared. GluC-digested W1Y and W3Y peptides were resuspended in 100 µL 60% Acetonitrile. For the reactions, 5 µL of peptide (W1Y or W3Y) was mixed with 4 µL methanol and 1 µL 2,4-DNPH (0.124 mM, large molar excess). In the negative control reaction, the 5 µL of peptide was replaced with 5 µL of 60% acetonitrile. After mixing, the solutions were placed into a 50 °C water bath for 4 h. From each reaction, 1 µL was mixed with CHCA matrix for MALDI-TOF-MS and MALDI-LIFT-TOF-TOF-MS analysis.

For formylglycine labeling, DarE processed and GluC-digested DarA-W3S and -K5S were concentrated and desalted by C18 Ziptip into a 60% acetonitrile, 0.1% formic acid solution. In a separate tube, ~5 µg of GluC-digested peptide was mixed with 10-fold molar excess of *o*-benzylhydroxylamine (*o*-BHA) in 0.1% formic acid, and 30 mM PIPES pH 6.5 with a final pH of reaction at pH 4. The labeling reaction proceeded for 16 h at room temperature, after which the reactions were transferred to low binding microfuge tubes (Eppendorf). Acetonitrile was added to each reaction, and the samples were concentrated by a SpeedVac SPD120 (Thermo Scientific) to dryness. The resulting pellet was resuspended in 100 mM 1,2-piperazinediethanesulfonic acid (PIPES) pH 6.5, and the samples were concentrated and desalted by C18 Ziptip into 60% acetonitrile. For MALDI-TOF-MS and MALDI-LIFT-TOF-TOF-MS analysis, 1 µL of each sample was mixed with either CHCA (variant W3S) or SDHB (variant K5S) matrix.

**Darobactin computational modeling.** Darobactin A, darobactin W3Y, and darobactin K5F were designed *in silico* based on stereochemical assignment from NMR and crystal structures, and energy minimized in MOE 2020.0901. Energy minimization parameters: AMBER14:EHT, Born Illicit solvent model, rigid water molecules, planar systems are rigid bodies, and RMS gradient 0.0001 kcal/mol.$Å^2$. Darobactins co-crystallized with BamA (PDB codes: 7NRE, 7NRF, 7P1C) were analyzed without energy minimization. All darobactins were visualized in Chimera.[3,4] DarA: DarE multimer predicted structures were generated with AlphaFold 2.1.2, and the highest confidence structures were selected for analysis.[15]

**Quantum mechanical (QM) calculations**. Full geometry optimizations were carried out with Gaussian 16[5] using the ωB97X-D[6] hybrid functional and 6-311G(d,p) basis set with ultrafine integration grids. Bulk solvent effects in water (ε=78) and diethyl ether (ε=4), the latter used in the calculations to mimic the polarity inside active sites of typical enzymes, were considered implicitly through the IEF-PCM polarizable continuum model.[7–9] Frequency analyses were carried out at the same level used in the geometry optimizations, and the nature of the stationary points was determined in each case according to the appropriate number of imaginary eigenvalues of the Hessian matrix. The quasiharmonic approximation reported by Truhlar et al. was used to replace the harmonic oscillator approximation for the calculation of the vibrational contribution to entropy.[10] Scaled frequencies were not considered. Mass-weighted intrinsic reaction coordinate

(IRC) calculations were carried out by using the Hratchian and Schlegel algorithm[11,12] in order to ensure that the TSs indeed connected the appropriate reactants and products. Gibbs free energies (ΔG) were used for the discussion on the relative stabilities of the considered structures. Electronic energies, entropies, enthalpies, Gibbs free energies, and lowest frequencies of the lowest energy calculated structures are summarized in Tables S5 and S6. Cartesian coordinates of the lowest energy structures calculated with PCM($H_2O$)/ωB97X-D/6-311G(d,p) and PCM($Et_2O$)/ωB97X-D/6-311G(d,p) are shown in Table S7 and S8, respectively.

**DarE model generation**. A full-atom model of *apo* DarE was generated using AlphaFold 2.3.2.[13] One hundred structure prediction runs were performed for each of the five neural network models, and the highest confidence structure (pLDDT score = 89.99) was selected. Cofactors, i. e. an *S*-adenosylmethionine (SAM) and three [$Fe_4S_4$] iron-sulfur clusters were modeled using the AlphaFill[14] online tool (https://alphafill.eu) using an anaerobic sulfatase-maturating enzyme as a template (PDB ID: 4K36). The resulting *holo* model was subjected to relaxation through a short molecular dynamics simulation.

**Molecular dynamics (MD) simulations**. MD simulations were carried out with AMBER 22[16] suite using the *ff19SB*[17] force field for the protein and *gaff2*[18] for the SAM cofactor. Specific parameters for iron-sulfur clusters were taken from the literature.[19] The *holo* enzyme model was immersed in a water box with an 8 Å buffer of OPC3[20] water molecules and neutralized by adding explicit $Na^+$ counterions. A two-stage geometry optimization approach was implemented. The first stage minimizes only the positions of solvent molecules and ions, and the second stage is an unrestrained minimization of all the atoms in the simulation cell. The systems were then heated by incrementing the temperature from 0 to 300 K under a constant pressure of 1 atm and periodic boundary conditions. Harmonic restraints of 10 kcal $mol^{-1}$ $Å^{-2}$ were applied to the solute, and the Andersen temperature coupling scheme[21,22] was used to control and equalize the temperature. The time step was kept at 1 fs during the heating stages, allowing potential inhomogeneities to self-adjust. Once equilibrated, the system was subjected to a 2 ns constant volume molecular dynamics simulation at 300 K using the SHAKE[23] algorithm and a 2 fs time step. Long-range electrostatic effects were modeled using the particle mesh Ewald method.[24] A cutoff of 8 Å was applied to Lennard-Jones interactions. Harmonic restraints of 10 kcal $mol^{-1}$ $Å^{-2}$ were applied to the cofactors at this stage to prevent large shifts from their initial placement. One hundred structures were sampled with an even stride from the 2 ns simulation and used for molecular docking.

**Molecular docking simulations**. Extended ether crosslinking transition state structures for a truncated darobactin model encompassing residues 1-5 (**OPh•_TS$_{7ext}$** and **OPh•_TS$_{6a-cext}$**) were generated using as starting point the coordinates deposited in the X-ray structure with PDB code 7NRI. These models have a free N-terminus at residue Trp1 and are capped with a NHMe group at residue Phe5. Geometric matching of the abbreviated QM transition state geometries was achieved by adjusting the internal coordinates (distances, angles, and torsions) of the truncated models only for those atoms which form part of the QM model (**OPh•_TS$_{7}$** and **OPh•_TS$_{6a-c}$**), and leaving the remaining atomic coordinates fixed at the crystallographic values. These structures were used as ligands for molecular docking. Docking calculations were performed using GOLD (CCDC Discovery 2020)[25] and the ChemScore fitness function.[26] The docking cavity was centered on the C6 atom of the SAM cofactor and allowed to extend in a spherical surrounding region with a 15 Å radius. For each ligand, the number of genetic algorithm runs was set to 30 and the 6 highest score solutions were saved. All rotatable bonds of the QM optimized geometry were kept frozen to their initial values. Five active site side chains were set as flexible in the docking procedure: Lys74, His87, Ser88, His123, and Asp340. Docking poses were collected and filtered based on the distance between the transition states and the SAM cofactor. For each docking pose, the sum of the two following distances was computed: 1) the distance between the C6 of the *S*-adenosylmethionine cofactor and the Cβ of Trp3, and 2) the distance between the C6 of the SAM cofactor and the Cβ of Phe5. The poses that minimize the sum of these distances were considered for analysis. After identifying the structure that yield the best binding pose for each transition state, the SAM cofactor transplanted through AlphaFill was changed to 5-dAdoH and methionine preserving the orientation of 5-dAdoH as in the parent SAM cofactor and rotating the side chain of the methionine to minimize steric clashes, and docking was repeated with the same parameters and fixing the protein's geometry to provide mechanistically coherent binding poses.

**Table S1: Oligonucleotide Primers Used in this Study.** F and R indicate the forward and reverse primers respectively. Lowercase letters represent the mutated codons. For the NNK and MNN codons: N = A, C, G, or T; K = G or T; M = C or A. Stop codons are indicated by *. Capitalization indicates the portion of the sequence that anneals to the gene while lower case indicates the site of mutation or an off-gene sequence.

| Primer Name | Primer Sequence (5' to 3') |
|---|---|
| DaroAE_MBP_F | CCTGTACTTCCAATCCGGATCCATGCATAACACATTGAATGAAA |
| DaroAE_MBP_R | GCCGCAAGCTTGTCGACTCGAGTTAGGCAGCAATGGTTTGCTTG |
| MBPDarAE_ColonyPCR_F | GCTATCATGCCATACCGCGAA |
| MBPDarAE_ColonyPCR_R | AACCGTCTATCAGGGCGATG |
| DarA(Q8*)_F | CTAAGTCCTTTtaaGAAATCTAAAAGGAGATATACCATGGA |
| DarA(Q8*)_R | TTTTAGATTTCttaAAAGGACTTAGACCAATTCCAAGCAGT |
| DarA_W1NNK_F | AAATTACTGCTnnkAATTGGTCTAAGTCCTTTTAAAAGGAG |
| DarA_W1NNK_R | TTAGACCAATTmnnAGCAGTAATTTCTGGGATCTTTGGCTT |
| DarA_W3NNK_F | CTGCTTGGAATnnkTCTAAGTCCTTTTAAAAGGAGATATAC |
| DarA_W3NNK_R | AAGGACTTAGAmnnATTCCAAGCAGTAATTTCTGGGATCTT |
| DarA(Q8*)_K5NNK_F | GGAATTGGTCTnnkTCCTTTTAAGAAATCTAAAAGGAGATA |
| DarA(Q8*)_K5NNK_R | TCTTAAAAGGAmnnAGACCAATTCCAAGCAGTAATTTCTGG |
| DarA(Q8*)_W1C_F | AAATTACTGCTtgcAATTGGTCTAAGTCCTTTTAAGAAATC |
| DarA(Q8*)_W1C_R | TTAGACCAATTgcaAGCAGTAATTTCTGGGATCTTTGGCTT |
| DarA(Q8*)_W1D_F | AAATTACTGCTgatAATTGGTCTAAGTCCTTTTAAGAAATC |
| DarA(Q8*)_W1D_R | TTAGACCAATTatcAGCAGTAATTTCTGGGATCTTTGGCTT |
| DarA(Q8*)_W1F_F | AAATTACTGCTtttAATTGGTCTAAGTCCTTTTAAGAAATC |
| DarA(Q8*)_W1F_R | TTAGACCAATTaaaAGCAGTAATTTCTGGGATCTTTGGCTT |
| DarA(Q8*)_W1K_F | AAATTACTGCTaaaAATTGGTCTAAGTCCTTTTAAGAAATC |
| DarA(Q8*)_W1K_R | TTAGACCAATTtttAGCAGTAATTTCTGGGATCTTTGGCTT |
| DarA(Q8*)_W1P_F | AAATTACTGCTccgAATTGGTCTAAGTCCTTTTAAGAAATC |
| DarA(Q8*)_W1P_R | TTAGACCAATTcggAGCAGTAATTTCTGGGATCTTTGGCTT |
| DarA(Q8*)_W1I_F | AAATTACTGCTattAATTGGTCTAAGTCCTTTTAAGAAATC |
| DarA(Q8*)_W1I_R | TTAGACCAATTaatAGCAGTAATTTCTGGGATCTTTGGCTT |
| DarA(Q8*)_W1M_F | AAATTACTGCTatgAATTGGTCTAAGTCCTTTTAAGAAATC |
| DarA(Q8*)_W1M_R | TTAGACCAATTcatAGCAGTAATTTCTGGGATCTTTGGCTT |
| DarA(Q8*)_W1N_F | AAATTACTGCTaacAATTGGTCTAAGTCCTTTTAAGAAATC |
| DarA(Q8*)_W1N_R | TTAGACCAATTgttAGCAGTAATTTCTGGGATCTTTGGCTT |
| DarA(Q8*)_W1Q_F | AAATTACTGCTcagAATTGGTCTAAGTCCTTTTAAGAAATC |
| DarA(Q8*)_W1Q_R | TTAGACCAATTctgAGCAGTAATTTCTGGGATCTTTGGCTT |
| DarA(Q8*)_W1S_F | AAATTACTGCTagtAATTGGTCTAAGTCCTTTTAAGAAATC |
| DarA(Q8*)_W1S_R | TTAGACCAATTactAGCAGTAATTTCTGGGATCTTTGGCTT |
| DarA(Q8*)_W1T_F | AAATTACTGCTaccAATTGGTCTAAGTCCTTTTAAGAAATC |
| DarA(Q8*)_W1T_R | TTAGACCAATTggtAGCAGTAATTTCTGGGATCTTTGGCTT |
| DarA(Q8*)_W1Y_F | AAATTACTGCTtatAATTGGTCTAAGTCCTT |
| DarA(Q8*)_W1Y_R | AAGGACTTAGACCAATTataAGCAGTAATTT |
| DarA(Q8*)_W3C_F | CTGCTTGGAATtgcTCTAAGTCCTTTTAAGAAATCTAAAAG |

| | |
|---|---|
| DarA(Q8*)_W3C_R | TCTAAGTCCTTgcaAAGATCCCAGAAATTACTGCTTGGAAT |
| DarA(Q8*)_W3D_F | CTGCTTGGAATgatTCTAAGTCCTTTTAAGAAATCTAAAAG |
| DarA(Q8*)_W3D_R | TCTAAGTCCTTatcAAGATCCCAGAAATTACTGCTTGGAAT |
| DarA(Q8*)_W3F_F | CTGCTTGGAATtttTCTAAGTCCTTTTAAGAAATCTAAAAG |
| DarA(Q8*)_W3F_R | TCTAAGTCCTTaaaAAGATCCCAGAAATTACTGCTTGGAAT |
| DarA(Q8*)_W3H_F | CTGCTTGGAATcatTCTAAGTCCTTTTAAGAAATCTAAAAG |
| DarA(Q8*)_W3H_R | AAGGACTTAGAatgATTCCAAGCAGTAATTTCTGGGATCTT |
| DarA(Q8*)_W3I_F | CTGCTTGGAATattTCTAAGTCCTTTTAAGAAATCTAAAAG |
| DarA(Q8*)_W3I_R | AAGGACTTAGAaatATTCCAAGCAGTAATTTCTGGGATCTT |
| DarA(Q8*)_W3K_F | CTGCTTGGAATaaaTCTAAGTCCTTTTAAGAAATCTAAAAG |
| DarA(Q8*)_W3K_R | AAGGACTTAGAtttATTCCAAGCAGTAATTTCTGGGATCTT |
| DarA(Q8*)_W3M_F | CTGCTTGGAATatgTCTAAGTCCTTTTAAGAAATCTAAAAG |
| DarA(Q8*)_W3M_R | AAGGACTTAGAcatATTCCAAGCAGTAATTTCTGGGATCTT |
| DarA(Q8*)_W3N_F | CTGCTTGGAATaacTCTAAGTCCTTTTAAGAAATCTAAAAG |
| DarA(Q8*)_W3N_R | AAGGACTTAGAgttATTCCAAGCAGTAATTTCTGGGATCTT |
| DarA(Q8*)_W3R_F | CTGCTTGGAATcgtTCTAAGTCCTTTTAAGAAATCTAAAAG |
| DarA(Q8*)_W3R_R | AAGGACTTAGAacgATTCCAAGCAGTAATTTCTGGGATCTT |
| DarA(Q8*)_W3Y_F | CTGCTTGGAATtatTCTAAGTCCTTTTAAGAAATCTAAAAG |
| DarA(Q8*)_W3Y_R | TCTAAGTCCTTataAAGATCCCAGAAATTACTGCTTGGAAT |
| DarA(Q8*)_K5C_F | GGAATTGGTCTtgcTCCTTTTAAGAAATCTAAAAGGAGATA |
| DarA(Q8*)_K5C_R | TCTTAAAAGGAgcaAGACCAATTCCAAGCAGTAATTTCTGG |
| DarA(Q8*)_K5F_F | GGAATTGGTCTtttTCCTTTTAAGAAATCTAAAAGGAGATA |
| DarA(Q8*)_K5F_R | TCTTAAAAGGAaaaAGACCAATTCCAAGCAGTAATTTCTGG |
| DarA(Q8*)_K5H_F | GGAATTGGTCTcatTCCTTTTAAGAAATCTAAAAGGAGATA |
| DarA(Q8*)_K5H_R | TCTTAAAAGGAatgAGACCAATTCCAAGCAGTAATTTCTGG |
| DarA(Q8*)_K5I_F | GGAATTGGTCTattTCCTTTTAAGAAATCTAAAAGGAGATA |
| DarA(Q8*)_K5I_R | TCTTAAAAGGAaatAGACCAATTCCAAGCAGTAATTTCTGG |
| DarA(Q8*)_K5N_F | GGAATTGGTCTaacTCCTTTTAAGAAATCTAAAAGGAGATA |
| DarA(Q8*)_K5N_R | TCTTAAAAGGAgttAGACCAATTCCAAGCAGTAATTTCTGG |
| DarA(Q8*)_K5T_F | GGAATTGGTCTaccTCCTTTTAAGAAATCTAAAAGGAGATA |
| DarA(Q8*)_K5T_R | TCTTAAAAGGAggtAGACCAATTCCAAGCAGTAATTTCTGG |
| DarA(Q8*)_K5W_F | GGAATTGGTCTtggTCCTTTTAAGAAATCTAAAAGGAGATA |
| DarA(Q8*)_K5W_R | TCTTAAAAGGAccaAGACCAATTCCAAGCAGTAATTTCTGG |
| DarA_T(-2A)_F | TCCCAGAAATTgcgGCTTGGAATTGGTCTAAGTCCTTTCAG |
| DarA_T(-2A)_R | CAATTCCAAGCcgcAATTTCTGGGATCTTTGGCTTATTGGA |
| DarA_I(-3A)_F | AGATCCCAGAAgcgACTGCTTGGAATTGGTCTAAGTCCTTT |
| DarA_I(-3A)_R | TTCCAAGCAGTcgcTTCTGGGATCTTTGGCTTATTGGACAA |
| DarA_L(-12)A_F | CATTAAACGAAgcgTCCAATAAGCCAAAGATCCCAGAAATT |
| DarA_L(-12)A_R | GGCTTATTGGAcgcTTCGTTTAATGCTTTGTCCGTAATTGA |
| DarA_E(-13)A_F | AAGCATTAAACgcgTTGTCCAATAAGCCAAAGATCCCAGAA |
| DarA_E(-13)A_R | TTATTGGACAAcgcGTTTAATGCTTTGTCCGTAATTGATAA |
| DarA_L(-15)A_F | CGGACAAAGCAgcgAACGAATTGTCCAATAAGCCAAAGATC |
| DarA_L(-15)A_R | GACAATTCGTTcgcTGCTTTGTCCGTAATTGATAACTCTGT |

| | |
|---|---|
| DarA_D(-18)A_F | TATCAATTACGgcgAAAGCATTAAACGAATTGTCCAATAAG |
| DarA_D(-18)A_R | TTTAATGCTTTcgcCGTAATTGATAACTCTGTTTCCTTGAA |
| DarA_I(-20A)_F | CAGAGTTATCAgcgACGGACAAAGCATTAAACGAATTGTCC |
| DarA_I(-20A)_R | GCTTTGTCCGTcgcTGATAACTCTGTTTCCTTGAACGAGGC |
| DarA_L(-22)A_F | AGGAAACAGAGgcgTCAATTACGGACAAAGCATTAAACGAA |
| DarA_L(-22)A_R | TCCGTAATTGAcgcCTCTGTTTCCTTGAACGAGGCAGCAAG |
| DarA_E(-23)A_F | TCAAGGAAACAgcgTTATCAATTACGGACAAAGCATTAAAC |
| DarA_E(-23)A_R | GTAATTGATAAcgcTGTTTCCTTGAACGAGGCAGCAAGGGA |
| DarA_K(-26A)_F | CTGCCTCGTTCgcgGAAACAGAGTTATCAATTACGGACAAA |
| DarA_K(-26A)_R | AACTCTGTTTCcgcGAACGAGGCAGCAAGGGAATTTAAGGC |
| DarA_F(-27A)_F | TTGCTGCCTCGgcgAAGGAAACAGAGTTATCAATTACGGAC |
| DarA_F(-27A)_R | TCTGTTTCCTTcgcCGAGGCAGCAAGGGAATTTAAGGCCTC |
| DarA_S(-28A)_F | CCCTTGCTGCCgcgTTCAAGGAAACAGAGTTATCAATTACG |
| DarA_S(-28A)_R | GTTTCCTTGAAcgcGGCAGCAAGGGAATTTAAGGCCTCCTG |
| DarA_L(-31A)_F | CCTTAAATTCCgcgGCTGCCTCGTTCAAGGAAACAGAGTTA |
| DarA_L(-31A)_R | AACGAGGCAGCcgcGGAATTTAAGGCCTCCTGAGTTTTAAC |
| DarA_S(-32A)_F | AGGCCTTAAATgcgCTTGCTGCCTCGTTCAAGGAAACAGAG |
| DarA_S(-32A)_R | GAGGCAGCAAGcgcATTTAAGGCCTCCTGAGTTTTAACAGT |
| DarA_L(-34A)_F | CTCAGGAGGCCgcgAATTCCCTTGCTGCCTCGTTCAAGGAA |
| DarA_L(-34A)_R | GCAAGGGAATTcgcGGCCTCCTGAGTTTTAACAGTTTCATT |
| DarALeader_5ΔN_Ins_F | tgtacttccaatccggatccAATGAAACTGTTAAAACTCAGGAG |
| DarALeader_5ΔN_Bb_R | ggatccggattggaagtacaggttc |
| DarALeader_5ΔN_Bb_F | AAGGAGATATACCATGGATACGATTATCCC |
| DarALeader_5ΔN_Ins_R | GGATAATCGTATCCATGGTATATCTCCTTTTAGATTTCCTGAAAGGACTTAGACC |
| DarALeader_10ΔN_Ins_F | tgtacttccaatccggatccACTCAGGAGGCCTTAAATTCC |
| DarALeader_10ΔN_Bb_R | ggatccggattggaagtacaggttc |
| DarALeader_10ΔN_Bb_F | AAGGAGATATACCATGGATACGATTATCCCT |
| DarALeader_10ΔN_Ins_R | GGATAATCGTATCCATGGTATATCTCCTTTTAGATTTCCTGAAAGGACTTAGACCA |
| DarALeader_15ΔN_Ins_F | cttccaatccggatccAATTCCCTTGCTGCC |
| DarALeader_15ΔN_Bb_R | ggatccggattggaagtacaggtt |
| DarALeader_15ΔN_Bb_F | AAGGAGATATACCATGGATACGATTATCCC |
| DarALeader_15ΔN_Ins_R | GGATAATCGTATCCATGGTATATCTCCTTTTAGATTTCCTGAAAGGACTTAGACCA |
| DarALeader_20ΔN_Ins_F | gtacttccaatccggatccTCGTTCAAGGAAACAGAGTTAT |
| DarALeader_20ΔN_Bb_R | ggatccggattggaagtacaggt |
| DarALeader_20ΔN_Bb_F | GGAATTGGTCTAAGTCCTTTCAGGAAATCTAA |
| DarALeader_20ΔN_Ins_R | GGATAATCGTATCCATGGTATATCTCCTTTTAGATTTCCTGAAAGGACTTAGACCA |
| DarALeader_25ΔN_Ins_F | ctgtacttccaatccggatccGAGTTATCAATTACGGACAAAGCATT |
| DarALeader_25ΔN_Bb_R | ggatccggattggaagtacaggt |
| DarALeader_25ΔN_Bb_F | AAGGAGATATACCATGGATACGATTATCCC |
| DarALeader_25ΔN_Ins_R | GGGATAATCGTATCCATGGTATATCTCCTTTTAGATTTCCTGAAAGGACTTAGACCAA |
| DarALeader_30ΔN_Ins_F | ctgtacttccaatccggatccGACAAAGCATTAAACGAATTGTCC |
| DarALeader_30ΔN_Bb_R | ggatccggattggaagtacaggt |
| DarALeader_30ΔN_Bb_F | AAGGAGATATACCATGGATACGATTATCCC |

| | |
|---|---|
| DarALeader_30ΔN_Ins_R | GGATAATCGTATCCATGGTATATCTCCTTTTAGATTTCCTGAAAGGACTTAGACCAA |
| MBPDarA T-2A Fwd | TCCCAGAAATTgcgGCTTGGAATTGGTCTAAGTCCTTTCAG |
| MBPDarA T-2A Rev | CAATTCCAAGCcgcAATTTCTGGGATCTTTGGCTTATTGGA |
| DarAQ8*_T(-2)S_F | TCCCAGAAATTagcGCTTGGAATTGGTCTAAGTCCTTTTAA |
| DarAQ8*_T(-2)S_R | CAATTCCAAGCgctAATTTCTGGGATCTTTGGCTTATTGGA |
| DarAQ8*_T(-2)V_F | TCCCAGAAATTgtgGCTTGGAATTGGTCTAAGTCCTTTTAA |
| DarAQ8*_T(-2)V_R | CAATTCCAAGCcacAATTTCTGGGATCTTTGGCTTATTGGA |
| DarA(Q8*)_T(-2)Remove_F | TCCCAGAAATTGCTTGGAATTGGTCTAAGTCCTTTTAA |
| DarA(Q8*)_T(-2)Remove_R | CAATTCCAAGCAATTTCTGGGATCTTTGGCTTATTGGA |
| DarA(Q8*)_InsA(-2)_F | CAGAAATTACTgcgGCTTGGAATTGGTCTAAGTCCTTTTAA |
| DarA(Q8*)_InsA(-2)_R | CAATTCCAAGCcgcAGTAATTTCTGGGATCTTTGGCTTATT |
| DarA(Q8*)_InsA(-3)_F | TCCCAGAAATTgcgACTGCTTGGAATTGGTCTAAGTCCTTT |
| DarA(Q8*)_InsA(-3)_R | TTCCAAGCAGTcgcAATTTCTGGGATCTTTGGCTTATTGGA |
| DarA(Q8*)_2-3InsG_F | CTGCTTGGAATggcTGGTCTAAGTCCTTTTAAGAAATCTAA |
| DarA(Q8*)_2-3InsG_R | GACTTAGACCAgccATTCCAAGCAGTAATTTCTGGGATCTT |
| DarA(Q8*)_2-3InsGG_F | CTGCTTGGAATggcggcTGGTCTAAGTCCTTTTAAGAAATCTAA |
| DarA(Q8*)_2-3InsGG_R | GACTTAGACCAgccATTCCAAGCAGTAATTTCTGGGATCTT |
| DarA(Q8*)_4-5InsG_F | GGAATTGGTCTggcAAGTCCTTTTAAGAAATCTAAAAGGAG |
| DarA(Q8*)_4-5InsG_R | TAAAAGGACTTgccAGACCAATTCCAAGCAGTAATTTCTGG |
| DarA(Q8*)_4-5InsGG_F | GGAATTGGTCTggcggcAAGTCCTTTTAAGAAATCTAAAAGGAG |
| DarA(Q8*)_4-5InsGG_R | TAAAAGGACTTgccgccAGACCAATTCCAAGCAGTAATTTCTGG |
| DarA(Q8*)_Remove N2_F | TTACTGCTTGGTGGTCTAAGTCCTTTTAAGAAATCTAA |
| DarA(Q8*)_Remove N2_R | GACTTAGACCACCAAGCAGTAATTTCTGGGATCTTTGG |
| DarA(Q8*)_Remove S4_F | CTTGGAATTGGAAGTCCTTTTAAGAAATCTAAAAGGAG |
| DarA(Q8*)_Remove S4_R | TAAAAGGACTTCCAATTCCAAGCAGTAATTTCTGGGAT |
| DarAQ8*_3-4InsW_F | CTTGGAATTGGtggTCTAAGTCCTTTTAAGAAATCTAAAAG |
| DarAQ8*_3-4InsW_R | AAGGACTTAGAccaCCAATTCCAAGCAGTAATTTCTGGGAT |
| DarAQ8*_2-3InsK_F | CTGCTTGGAATaaaTGGTCTAAGTCCTTTTAAGAAATCTAA |
| DarAQ8*_2-3InsK_R | GACTTAGACCAtttATTCCAAGCAGTAATTTCTGGGATCTT |
| DarAQ8*_3-4InsNW_F | CTTGGAATTGGaactggTCTAAGTCCTTTTAAGAAATCTAAAAG |
| DarAQ8*_3-4InsNW_R | AAGGACTTAGAccagttCCAATTCCAAGCAGTAATTTCTGGGAT |

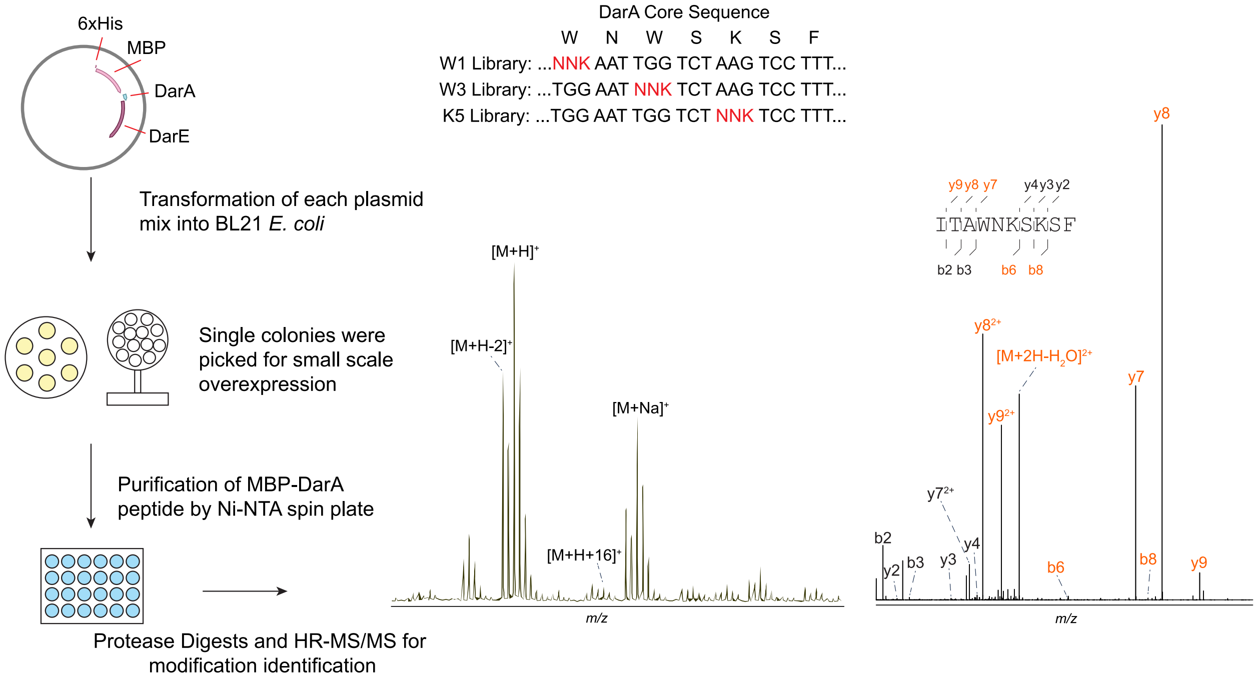

**Figure S1: Schematic overview of methods for small scale panel generation.** Three individual NNK panels were generated by PCR on the pET28-MBP-DarA(Q8*) plasmid at positions 1, 3, and 5 of the core region as indicated. This construct installs a stop codon that ablates the QEI follower sequence of DarA. The plasmid mix for each NNK panel was transformed into BL21(DE3), and single colonies were inoculated into LB for protein overexpression. MBP-DarA variants were purified by Ni-NTA chromatography and digested by both TEV and GluC for modification identification. This method was followed for each NNK panel separately.

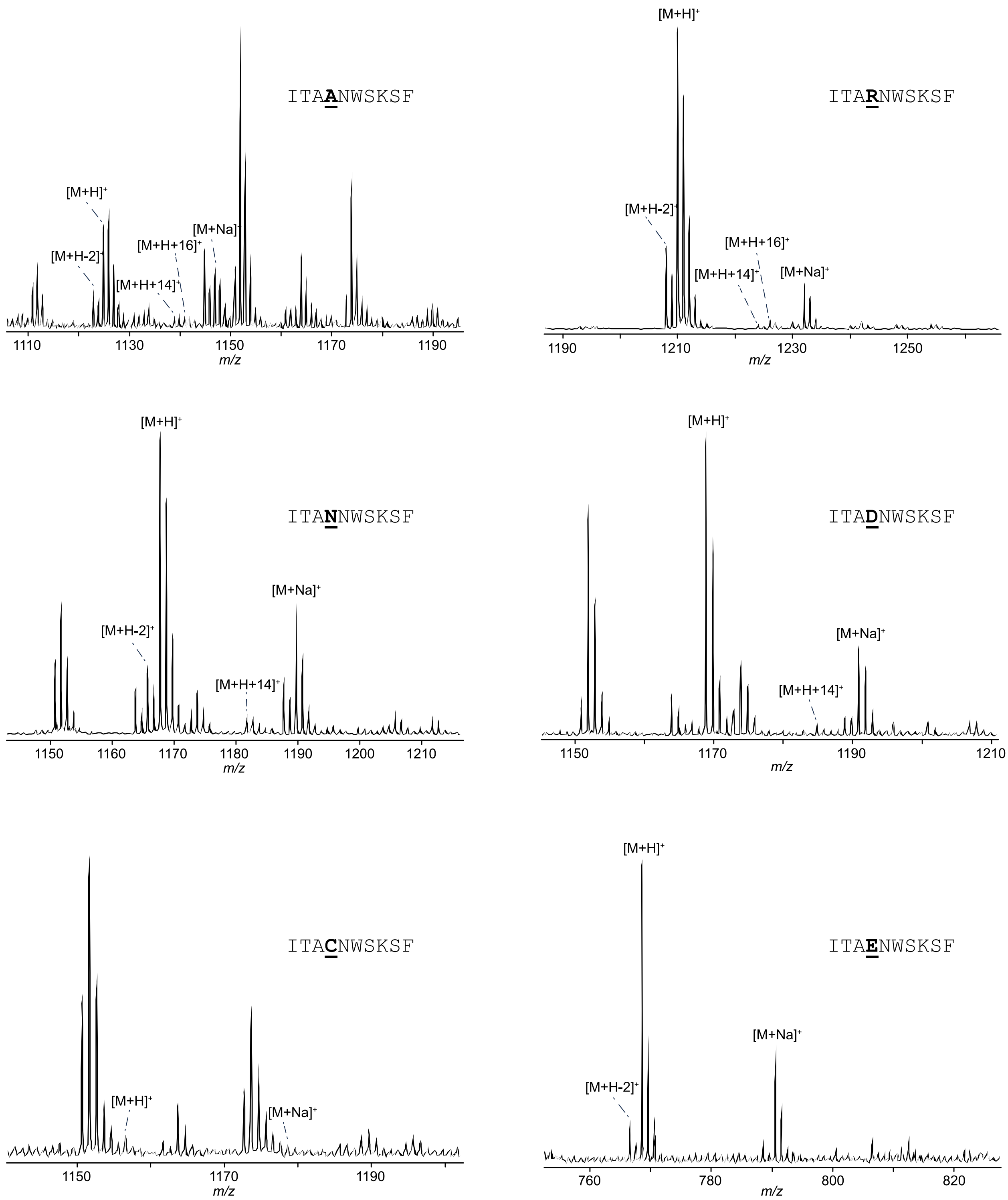

**Figure S2: MALDI-TOF-MS spectra for all DarA-Trp1 variants**

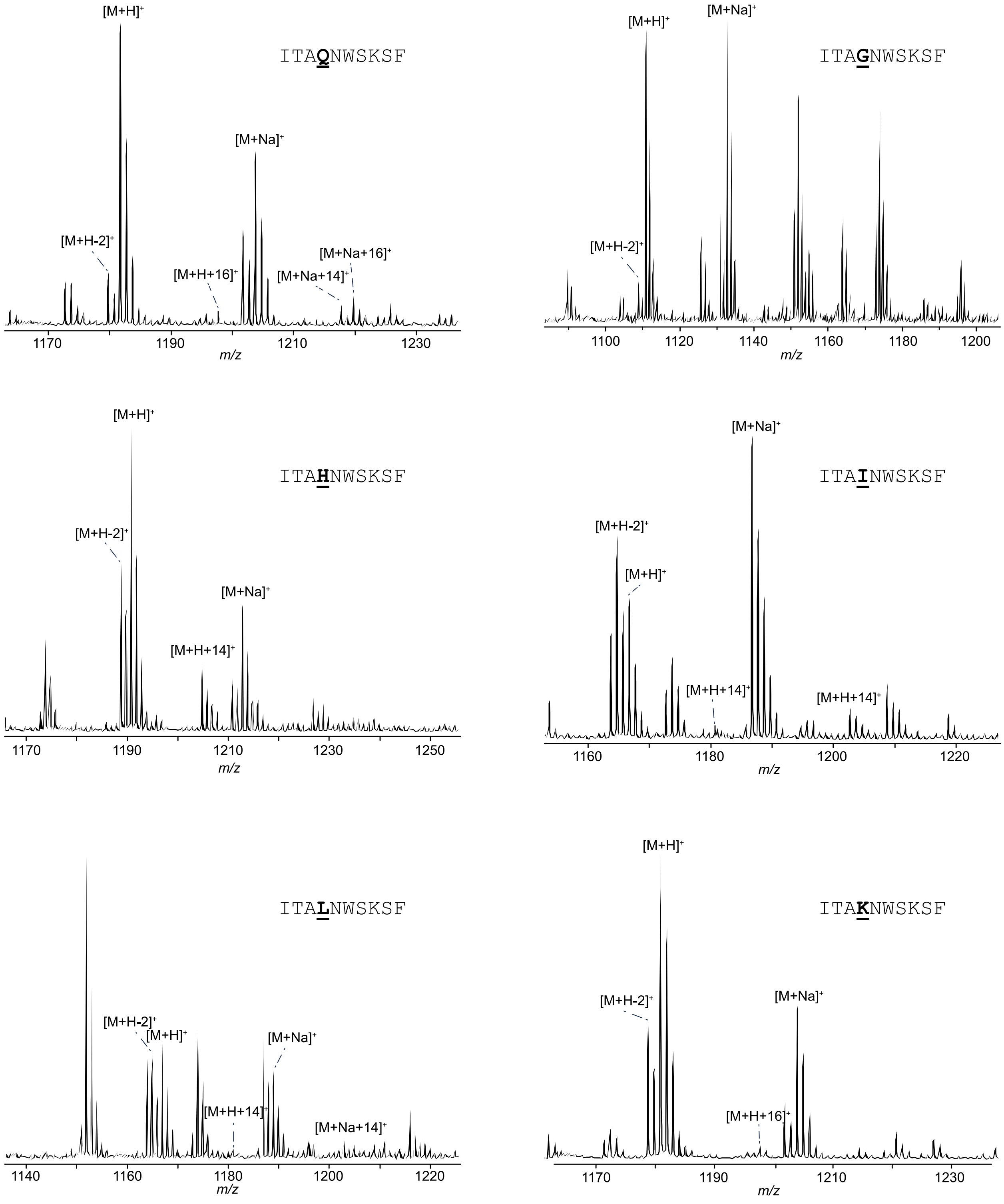

**Figure S2 cont.: MALDI-TOF-MS spectra for all DarA-Trp1 variants**

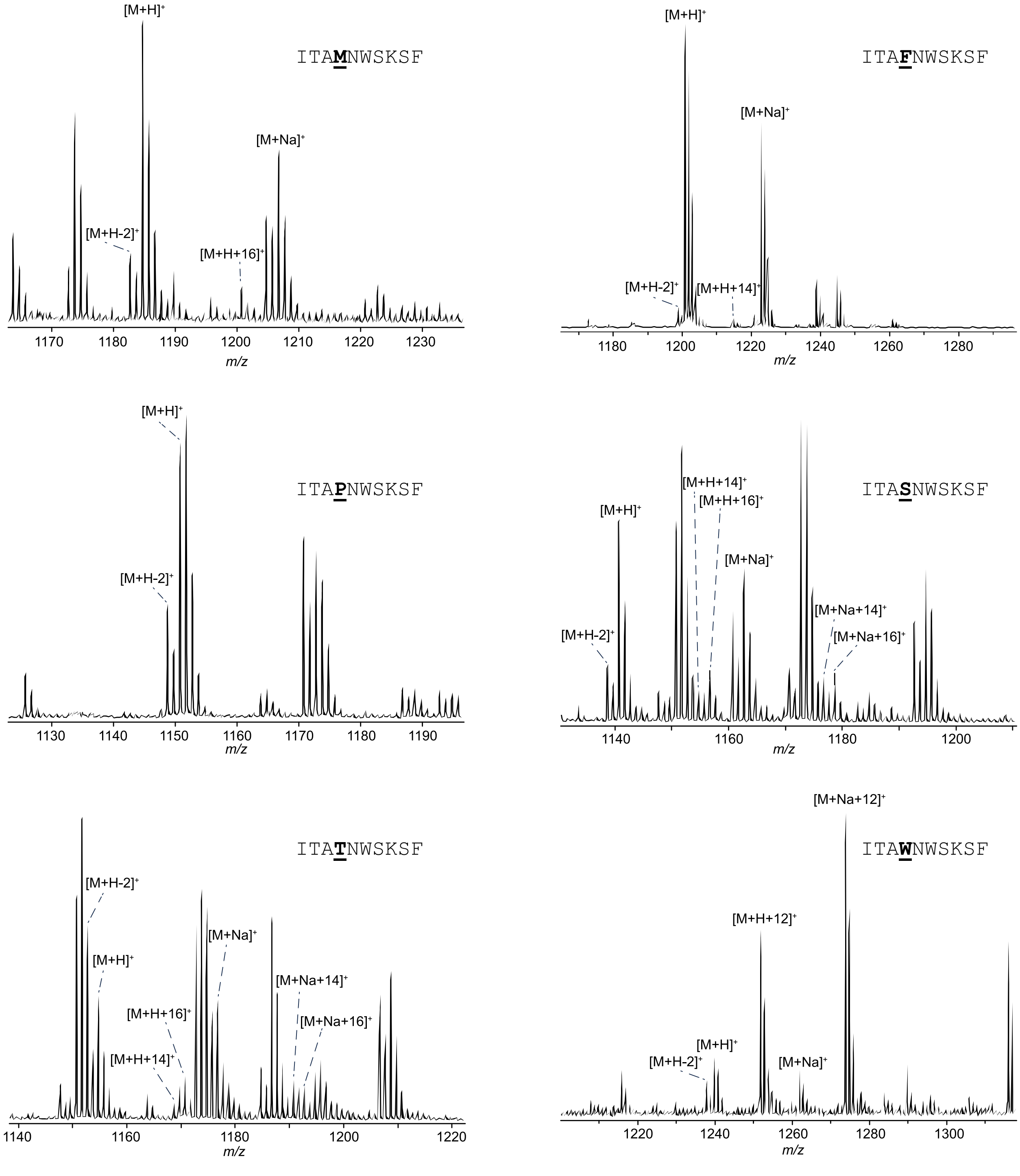

**Figure S2 cont.: MALDI-TOF-MS spectra for all DarA-Trp1 variants**

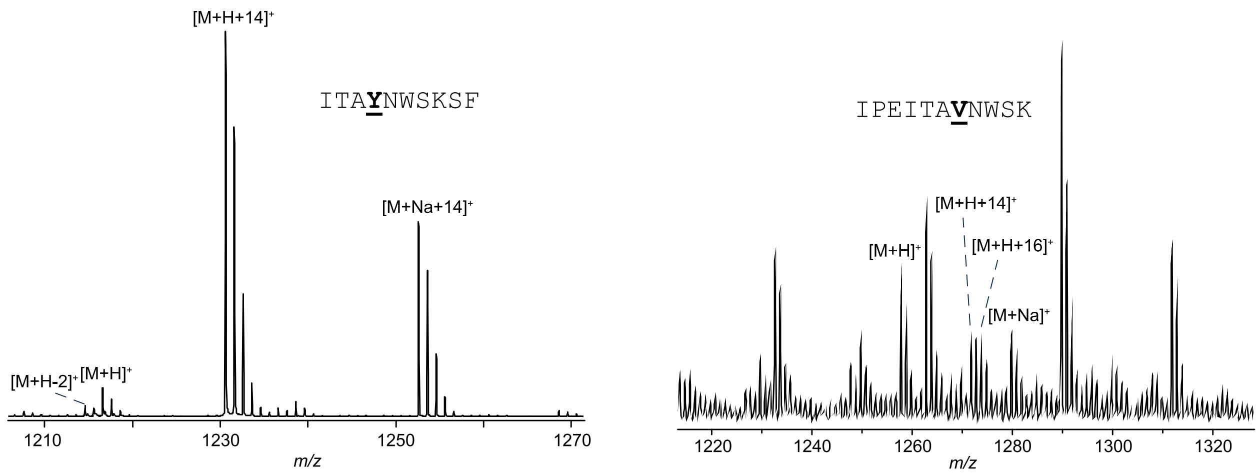

**Figure S2 cont.: MALDI-TOF-MS spectra for all DarA-Trp1 variants.** All variants were co-expressed with DarE, digested with TEV protease, and then digested with either endoproteinase GluC or trypsin. GluC digestion retains Ile-Thr-Ala from the DarA leader region while trypsin retains IPEITA. All peptides were then crystallized with CHCA or SDHB for MALDI-TOF-MS analysis. Ions of interest are indicated. $[M+H]^+$ corresponds to protonated ion of the unmodified peptide. The varied residue is bolded and underlined. Loss of 2 Da corresponds to dehydrogenation or C-C crosslink formation. Addition of 12 and 14 correspond to formation of the ether and C-C crosslinks, and a β-oxotryptophan modification, respectively. A +16 Da mass shift can result from hydroxylation.[27]

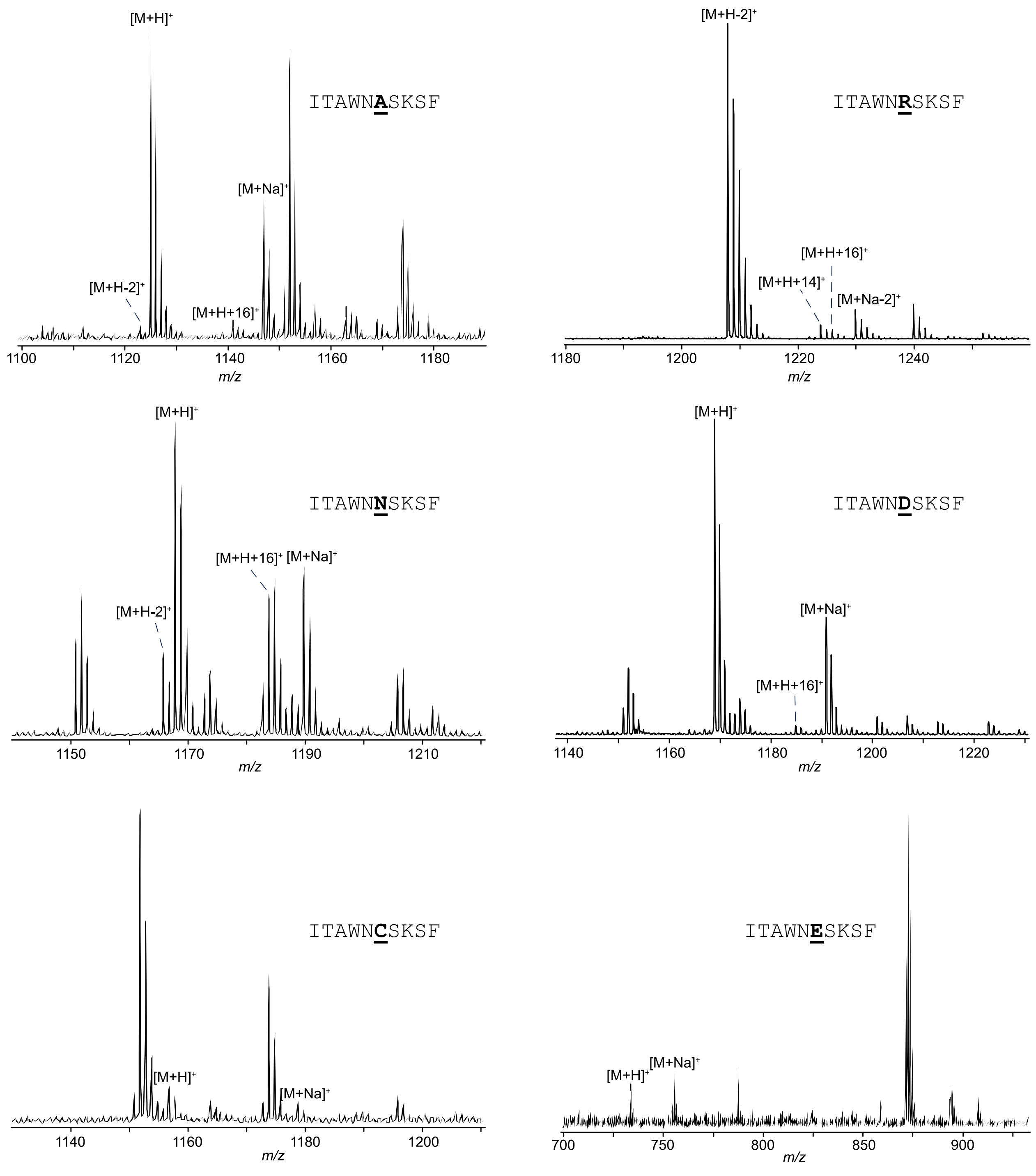

**Figure S3: MALDI-TOF-MS spectra for all DarA-Trp3 variants**

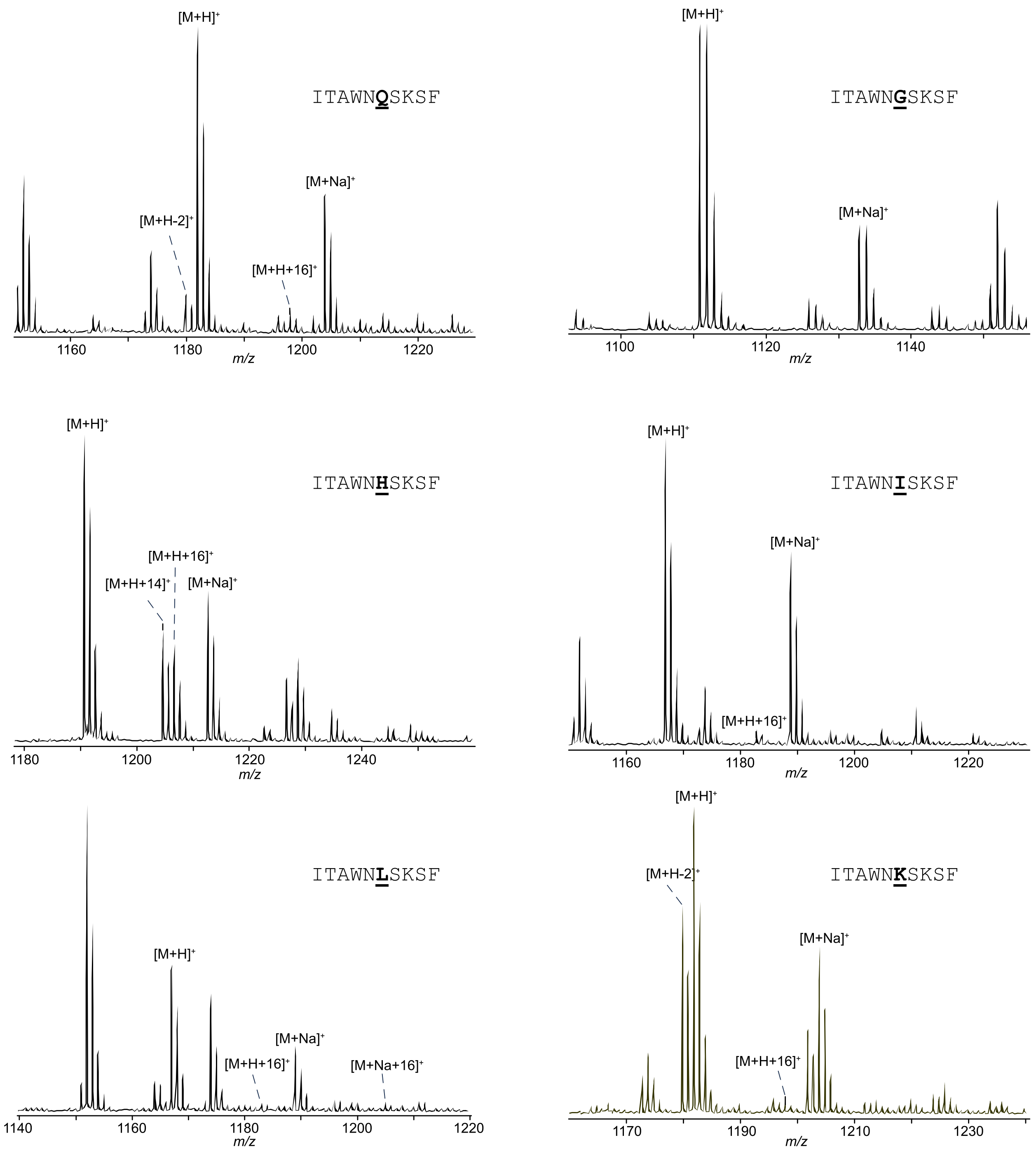

**Figure S3 cont.: MALDI-TOF-MS spectra for all DarA-Trp3 variants**

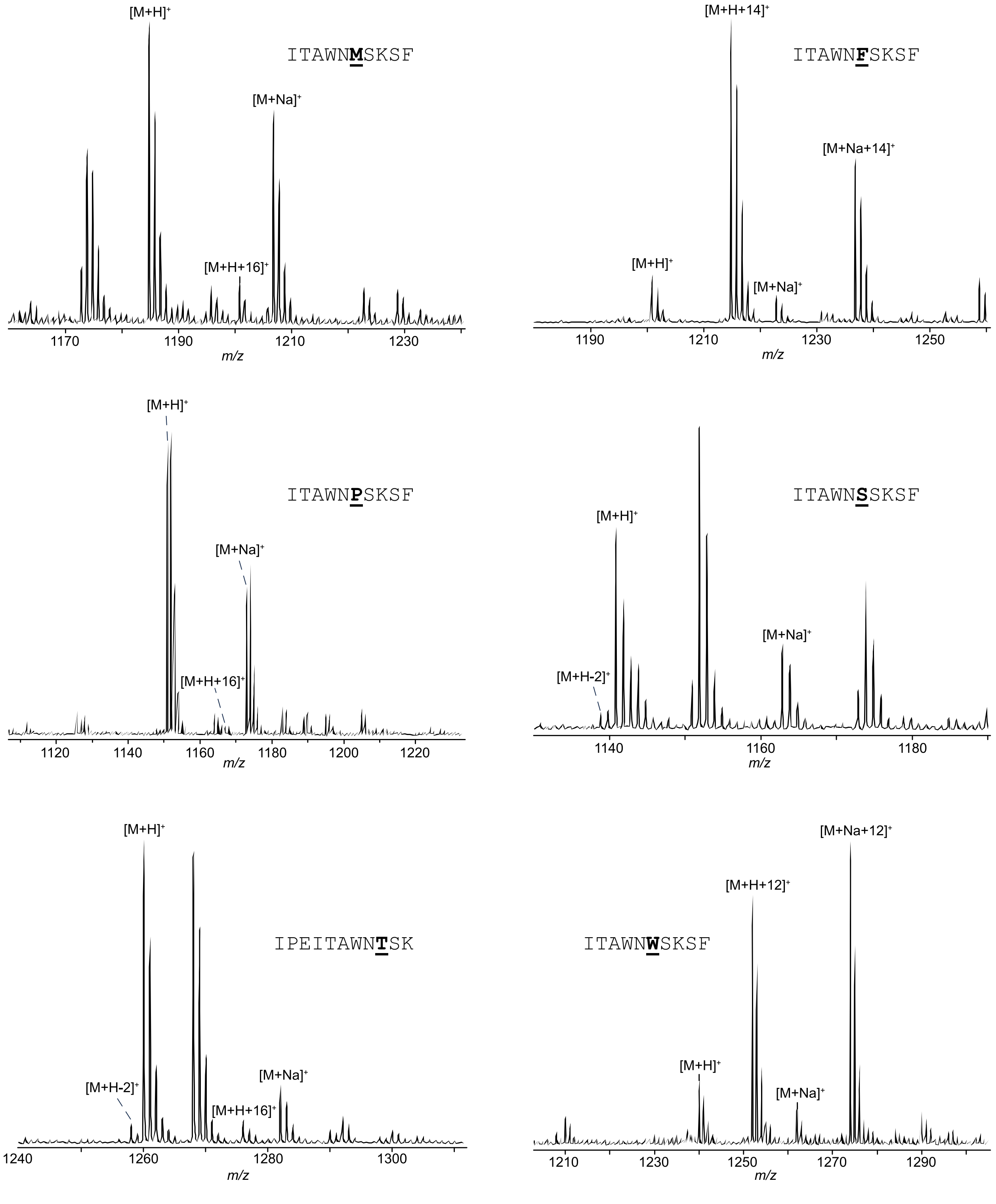

**Figure S3 cont.: MALDI-TOF-MS spectra for all DarA-Trp3 variants**

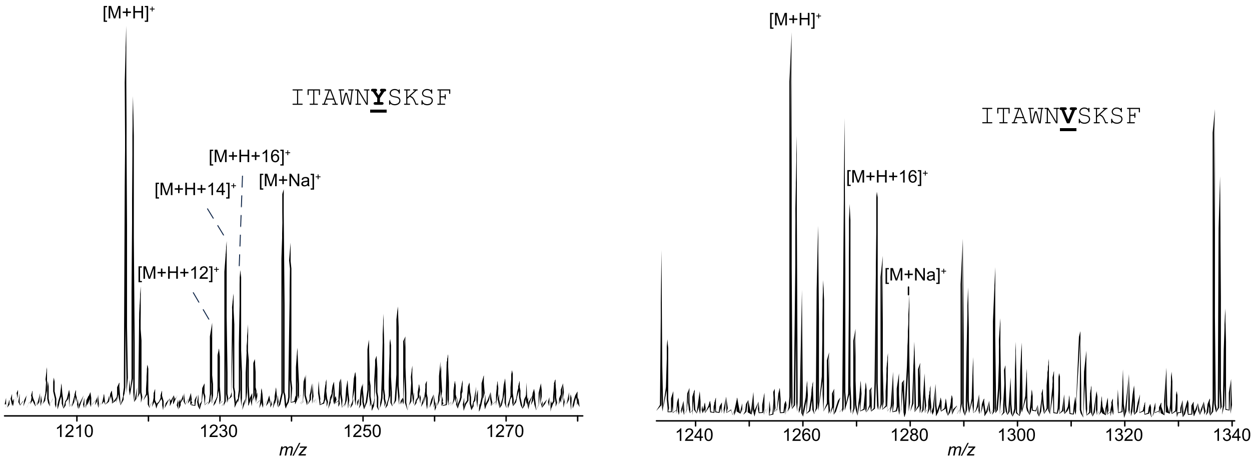

**Figure S3 cont.: MALDI-TOF-MS spectra for all DarA-Trp3 variants.** All variants were co-expressed with DarE, digested TEV protease, and then digested with either endoproteinase GluC or trypsin. GluC digestion retains Ile-Thr-Ala from the DarA leader region while trypsin retains IPEITA. All peptides were then crystallized with CHCA or SDHB for MALDI-TOF-MS analysis. Ions of interest are indicated. $[M+H]^+$ corresponds to protonated ion of the unmodified peptide. The varied residue is bolded and underlined. Loss of 2 Da corresponds to dehydrogenation, formylglycine formation, or C-C crosslink formation. Addition of 12 Da corresponds to ether and C-C crosslink formation. The +14 Da modification corresponds to either β-oxo amino acid or ether crosslink formation. A +16 Da mass shift can result from hydroxylation.[27]

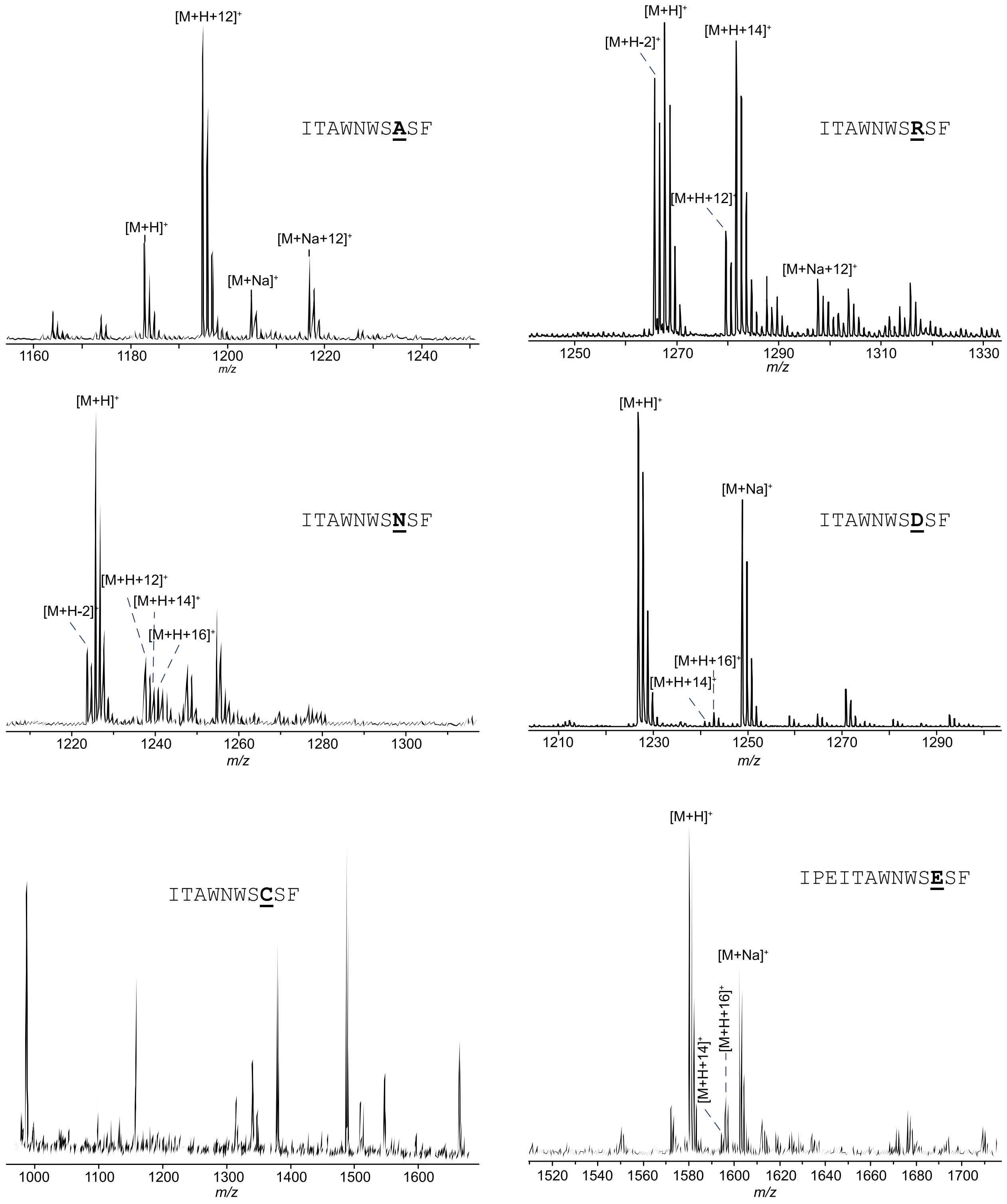

**Figure S4: MALDI-TOF-MS spectra for all DarA-Lys5 variants**

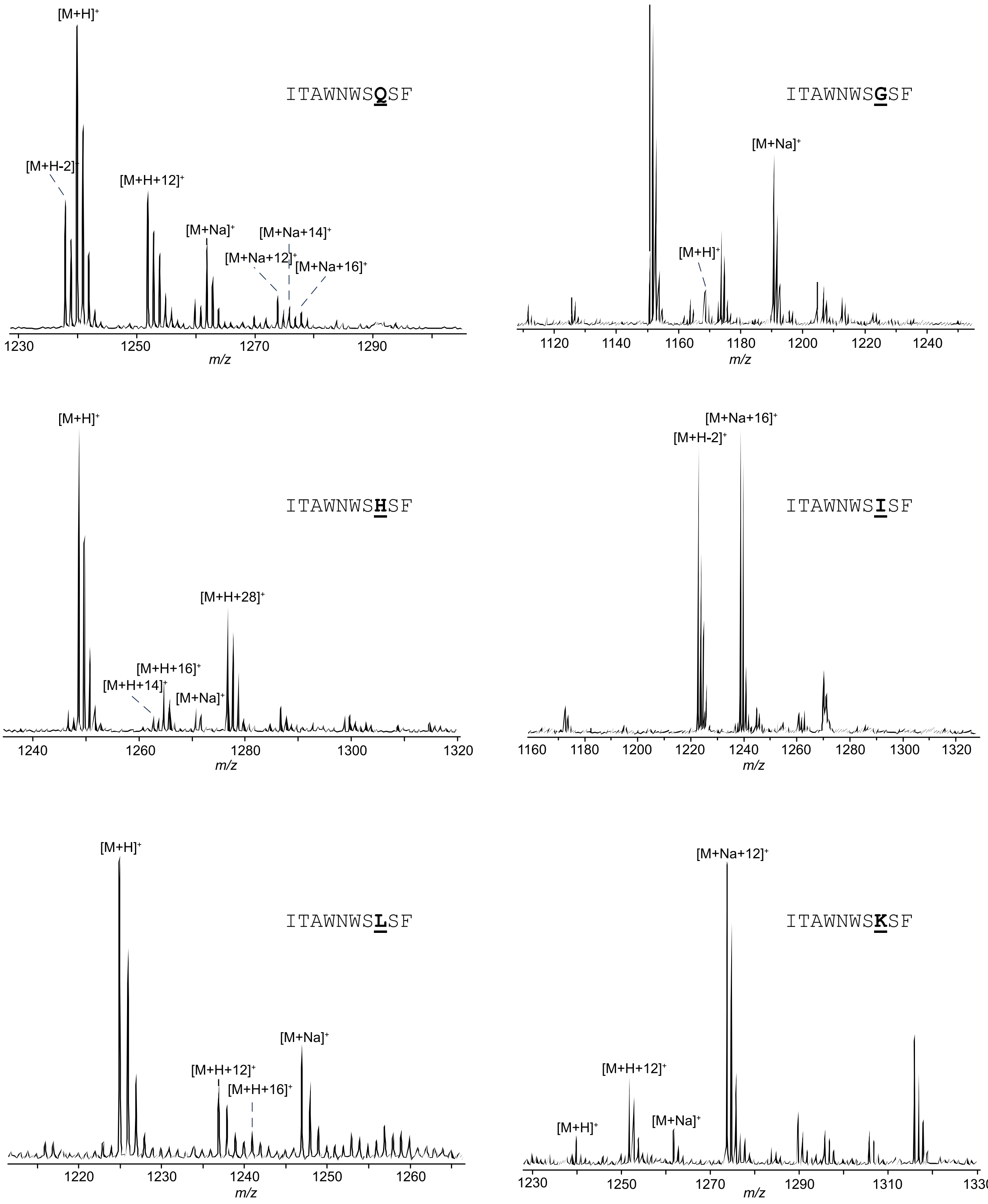

C

**Figure S4 Cont.: MALDI-TOF-MS spectra for all DarA-Lys5 variants**

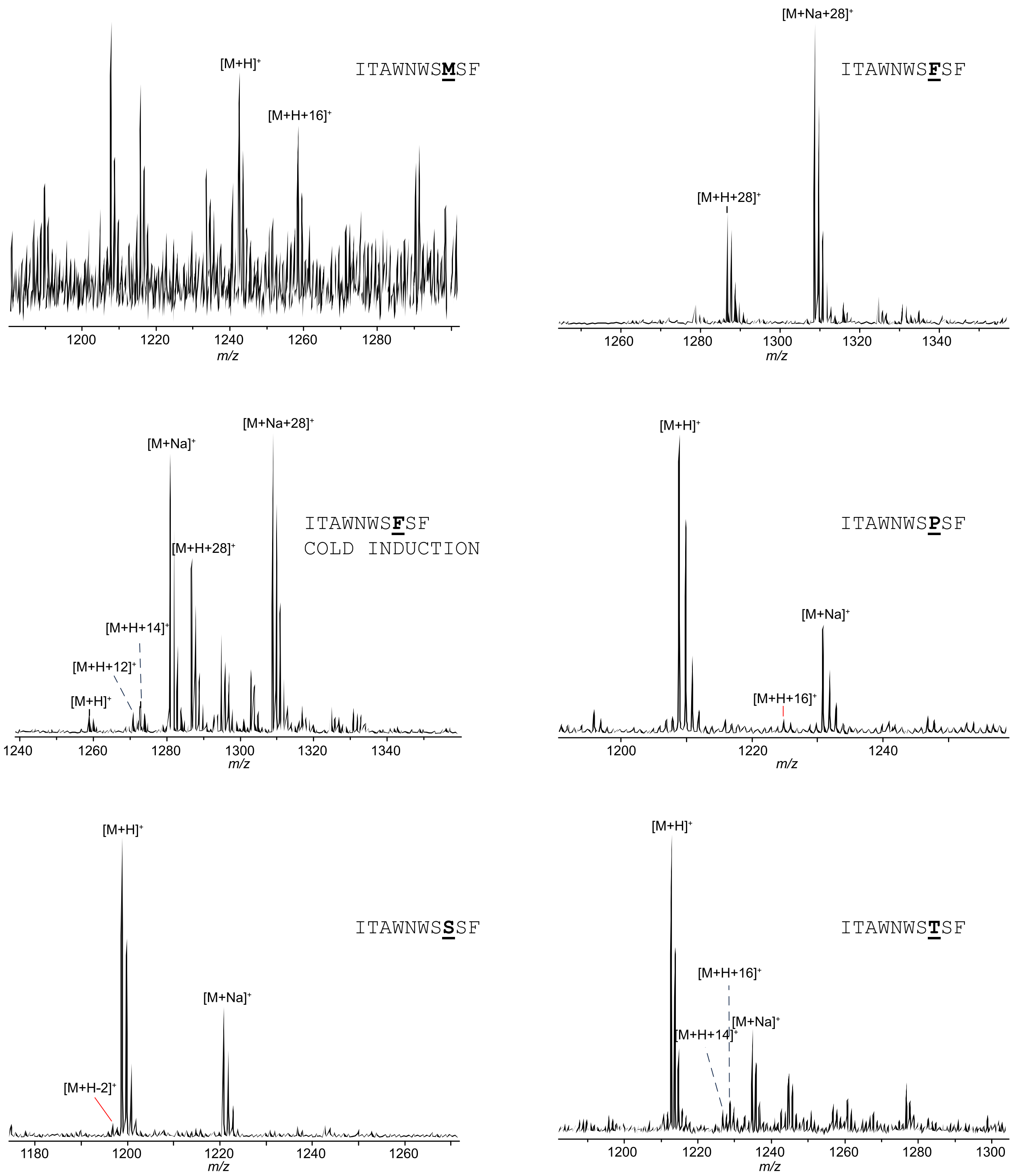

**Figure S4 cont.: MALDI-TOF-MS spectra for all DarA-Lys5 variants**

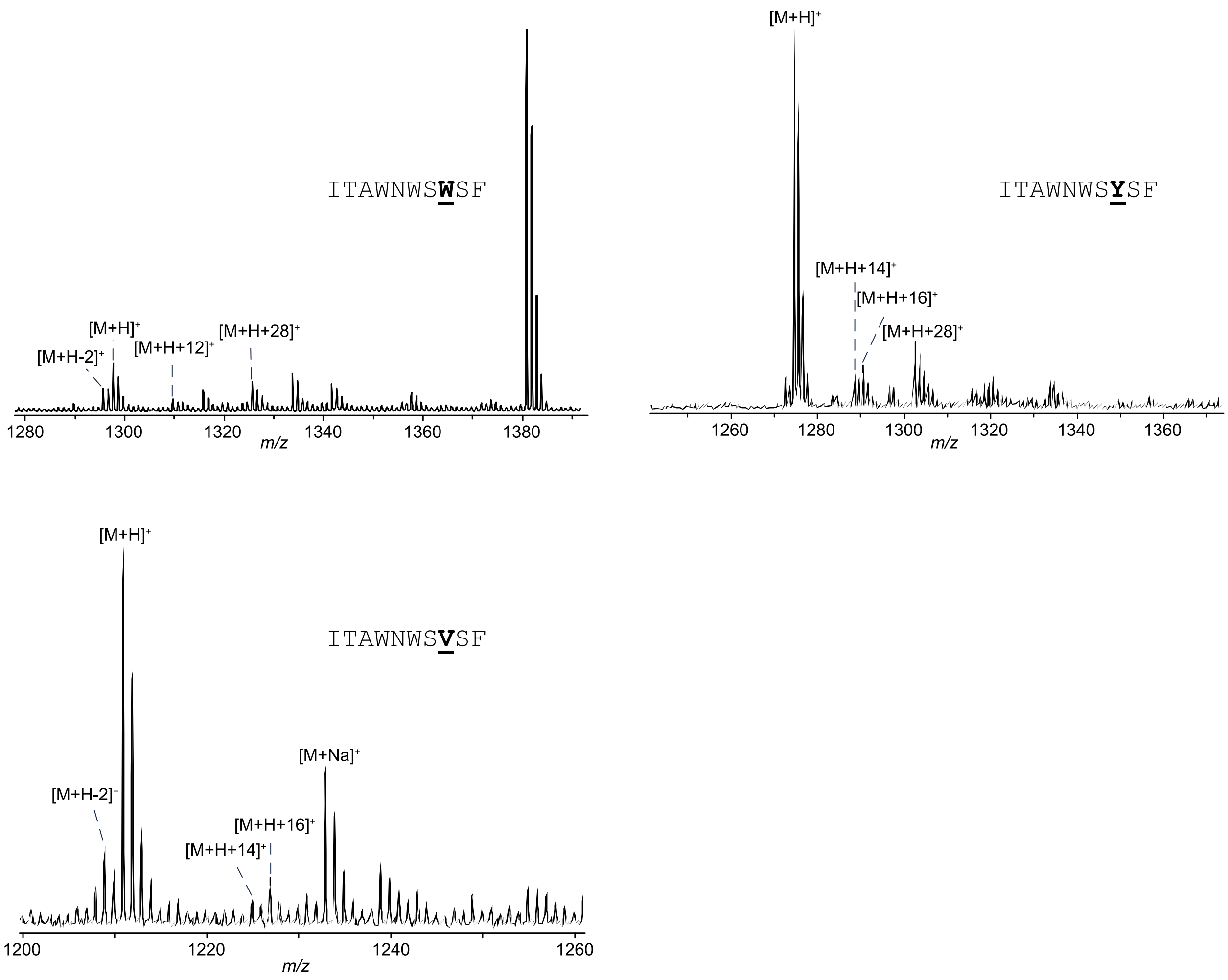

**Figure S4 cont.: MALDI-TOF-MS spectra for all DarA-Lys5 variants.** All variants were co-expressed with DarE, digested TEV protease, and either endoproteinase GluC or trypsin. GluC digestion retains Ile-Thr-Ala from the DarA leader region while trypsin retains IPEITA. All peptides were then crystallized with CHCA or SDHB for MALDI-TOF-MS analysis. Ions of interest are indicated. $[M+H]^+$ corresponds to protonated ion of the unmodified peptide. The varied residue is bolded and underlined. Loss of 2 Da corresponds to dehydrogenation, formylglycine formation, or C-C crosslink formation. Addition of 12 Da corresponds to ether and C-C crosslink formation. The +14 Da modification corresponds to either β-oxo amino acid or ether crosslink formation. A +16 Da mass shift can result from hydroxylation.[27] Addition of 28 Da corresponds to formation of two ether crosslinks or a single ether crosslink and a β-oxo amino acid modification.

A

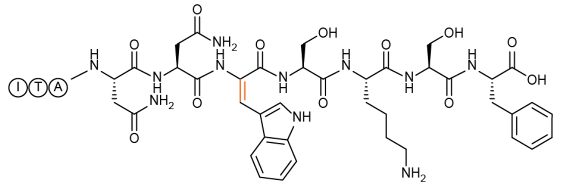

B

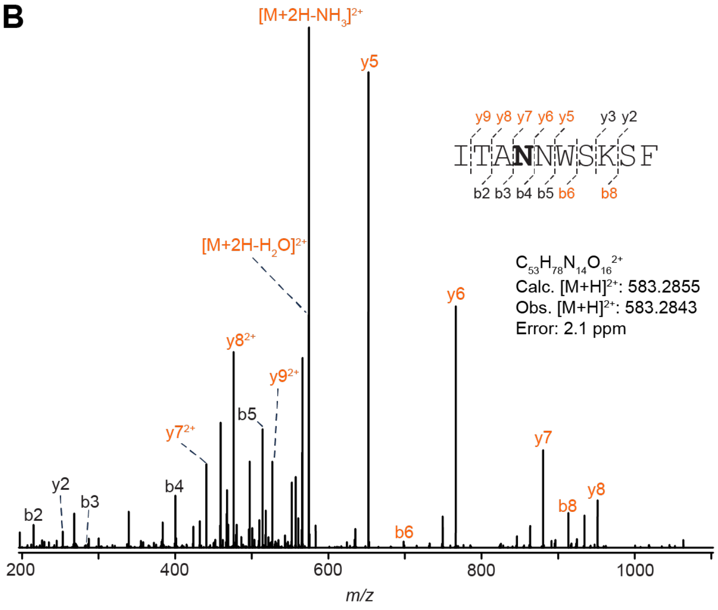

C

| Ion | Calculated Mass | Observed Mass | Error (ppm) |
|---|---|---|---|
| $[M+2H]^{2+}$ | 583.2855 | 583.2843 | 2.1 |
| $[M+2H-H_2O]^{2+}$ | 574.2803 | 574.2787 | 2.8 |
| $y2^+$ | 253.1183 | 253.1176 | 2.8 |
| $b8^+$ | 913.4527 | 913.4502 | 2.7 |
| $y3^+$ | 381.2133 | 381.2123 | 2.6 |
| $b6^+$ | 698.3257 | 698.3237 | 2.9 |
| $b5^+$ | 514.2620 | 514.2606 | 2.7 |
| $y5^+$ | 652.3090 | 652.3071 | 2.9 |
| $b4^+$ | 400.2191 | 400.2180 | 2.7 |
| $y6^+$ | 766.3520 | 766.3498 | 2.9 |
| $b3^+$ | 286.1762 | 286.1754 | 2.8 |
| $y7^+$ | 880.3949 | 880.3925 | 2.7 |
| $b2^+$ | 215.1391 | 215.1384 | 3.3 |
| $y8^+$ | 951.4320 | 951.4295 | 2.6 |
| $y8^{2+}$ | 476.2197 | 476.2183 | 2.9 |
| $y9^{2+}$ | 526.7435 | 526.7421 | 2.7 |

**Figure S5: HR-MS/MS of the DarA-W1N variant with a -2 Da modification.** (A) Hypothetical structure based on HR-MS/MS fragmentation. The dehydroTrp modification is highlighted in orange. (B) Collision-induced dissociation (CID) spectrum of the -2 Da modified variant. The sequence of the GluC-digested peptide is provided with the substitution bolded. Orange colored daughter ions include the modification. (C) Observed masses and errors from CID analysis.

A

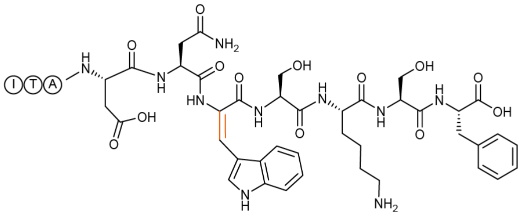

B

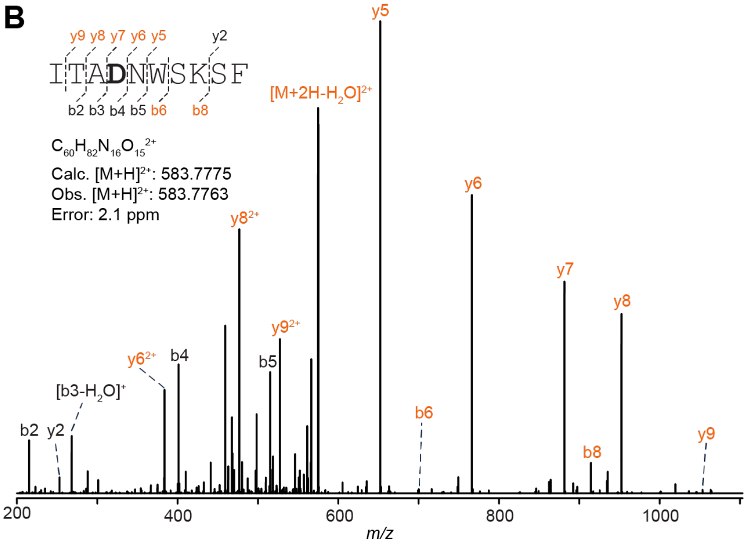

C

| Ion | Calculated Mass | Observed Mass | Error (ppm) |
|---|---|---|---|
| $[M+2H]^{2+}$ | 583.7775 | 583.7763 | 2.1 |
| $[M+2H-H_2O]^{2+}$ | 574.7722 | 574.7704 | 3.1 |
| $y2^+$ | 253.1183 | 253.1175 | 3.2 |
| $b8^+$ | 914.4367 | 914.4337 | 3.3 |
| $b6^+$ | 699.3097 | 699.3072 | 3.6 |
| $b5^+$ | 515.2461 | 515.2443 | 3.5 |
| $y5^+$ | 652.3090 | 652.3068 | 3.4 |
| $b4^+$ | 401.2031 | 401.2018 | 3.2 |
| $y6^+$ | 766.3519 | 766.3494 | 3.3 |
| $b3^+$ | 286.1762 | 286.1753 | 3.1 |
| $[b3-H_2O]^+$ | 268.1656 | 268.1648 | 3.0 |
| $y7^+$ | 881.3789 | 881.3759 | 3.4 |
| $y7^{2+}$ | 441.1931 | 441.1917 | 3.2 |
| $b2^+$ | 215.1391 | 215.1384 | 3.3 |
| $y8^+$ | 952.4160 | 952.4127 | 3.5 |
| $y8^{2+}$ | 476.7116 | 476.7101 | 3.1 |
| $y9^+$ | 1053.4636 | 1053.4595 | 3.9 |
| $y9^{2+}$ | 527.2355 | 527.2338 | 3.2 |

**Figure S6: HR-MS/MS of the DarA-W1D variant with a -2 Da modification.** (A) Hypothetical structure of the modified species based on HR-MS/MS fragmentation. The modification is highlighted in orange. (B) CID spectrum of the -2 Da modified variant. The sequence of the GluC-digested peptide is provided with the substitution bolded. Orange colored daughter ions include the modification. (C) Observed masses and errors from CID analysis.

A

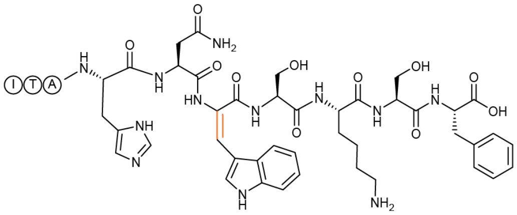

B

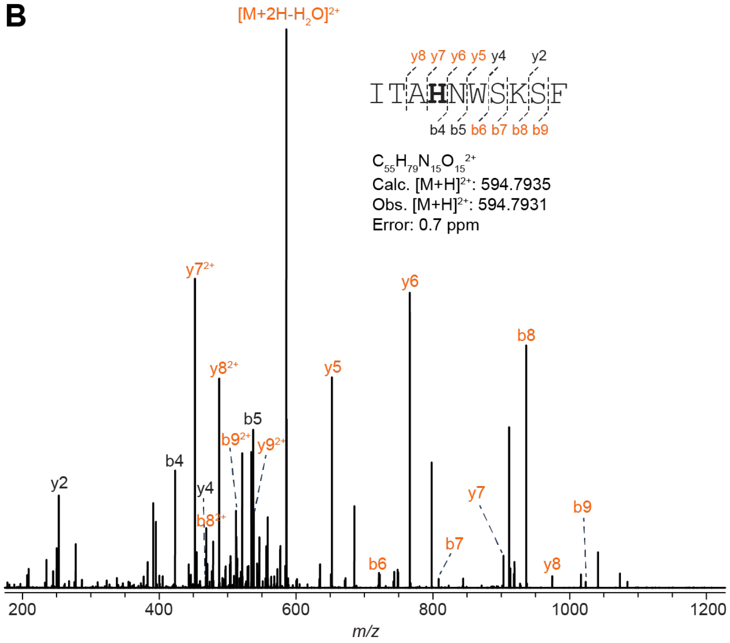

C

| Ion | Calculated Mass | Observed Mass | Error (ppm) |
|---|---|---|---|
| $[M+2H]^{2+}$ | 594.7935 | 594.7931 | 0.7 |
| $[M+2H-H_2O]^{2+}$ | 585.7882 | 585.7874 | 1.4 |
| $b9^+$ | 1023.5007 | 1023.4986 | 2.1 |
| $b9^{2+}$ | 512.2540 | 512.2529 | 2.1 |
| $y2^+$ | 253.1183 | 253.118 | 1.2 |
| $b8^+$ | 936.4687 | 936.4673 | 1.5 |
| $b8^{2+}$ | 468.7380 | 468.7374 | 1.3 |
| $b7^+$ | 808.3737 | 808.3731 | 0.7 |
| $y4^+$ | 468.2453 | 468.2448 | 1.1 |
| $b6^+$ | 721.3417 | 721.3406 | 1.5 |
| $b5^+$ | 537.2780 | 537.2772 | 1.5 |
| $y5^+$ | 652.3090 | 652.3081 | 1.4 |
| $b4^+$ | 423.2351 | 423.2345 | 1.4 |
| $y6^+$ | 766.3519 | 766.3509 | 1.3 |
| $y7^+$ | 903.4108 | 903.4096 | 1.3 |
| $y7^{2+}$ | 452.2091 | 452.2085 | 1.3 |
| $y8^+$ | 974.4479 | 974.4464 | 1.5 |
| $y8^{2+}$ | 487.7276 | 487.727 | 1.2 |
| $y9^{2+}$ | 538.2515 | 538.2508 | 1.3 |

**Figure S7: HR-MS/MS of the DarA-W1H variant with a dhTrp modification.** (A) Hypothetical structure of the modified species based on HR-MS/MS fragmentation. The modification is highlighted in orange. (B) CID spectrum of the -2 Da modified variant. The sequence of the GluC-digested peptide is provided with the substitution bolded. Orange colored daughter ions include the modification. (C) Observed masses and errors from CID analysis.

**A**

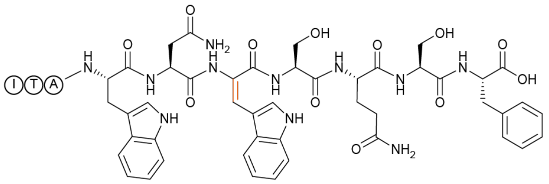

**B**

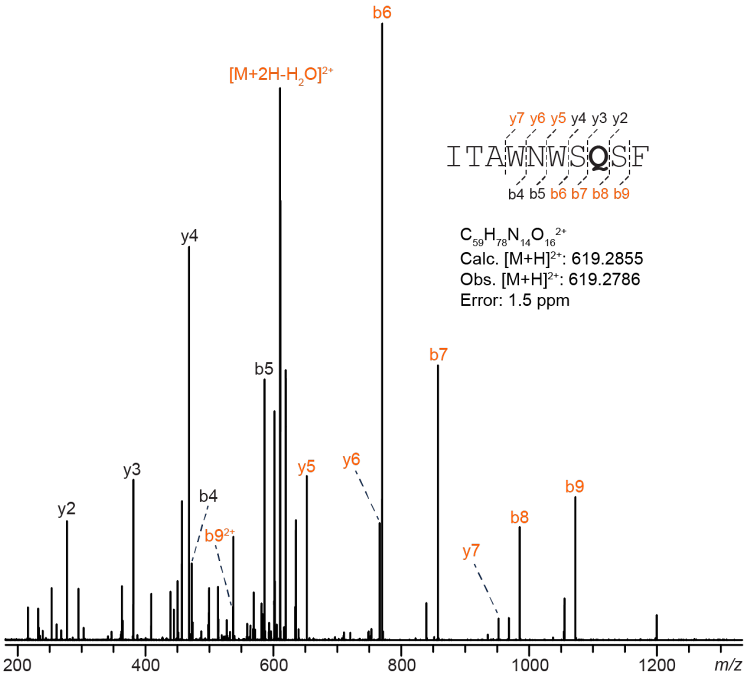

**C**

| Ion | Calculated Mass | Observed Mass | Error (ppm) |
|---|---|---|---|
| $[M+2H]^{2+}$ | 619.2855 | 619.2846 | 1.5 |
| $[M+2H-H_2O]^{2+}$ | 610.2802 | 610.2786 | 2.6 |
| $b9^+$ | 1072.4847 | 1072.4818 | 2.7 |
| $b9^{2+}$ | 536.7460 | 536.7447 | 2.4 |
| $y2^+$ | 253.1183 | 253.1177 | 2.4 |
| $b8^+$ | 985.4527 | 985.4502 | 2.5 |
| $y3^+$ | 381.1769 | 381.1760 | 2.4 |
| $b7^+$ | 857.3941 | 857.3918 | 2.7 |
| $y4^+$ | 468.2089 | 468.2078 | 2.3 |
| $b6^+$ | 770.3622 | 770.3604 | 2.3 |
| $b5^+$ | 586.2984 | 586.2970 | 2.4 |
| $y5^+$ | 652.2726 | 652.2709 | 2.6 |
| $b4^+$ | 472.2555 | 472.2544 | 2.3 |
| $y6^+$ | 766.3155 | 766.3137 | 2.3 |
| $y7^+$ | 952.3948 | 952.3924 | 2.5 |

**Figure S8: HR-MS/MS analysis of the DarA-K5Q variant with a dhTrp modification.** (A) Hypothetical structure of the modified species based on HR-MS/MS fragmentation. The modification is highlighted in orange. (B) CID spectrum of the -2 Da modified variant. The sequence of the GluC-digested peptide is provided with the substitution bolded. Orange colored daughter ions include the modification. (C) Observed masses and errors from CID analysis.

A

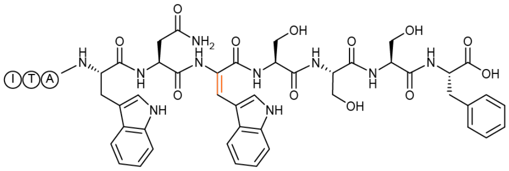

B

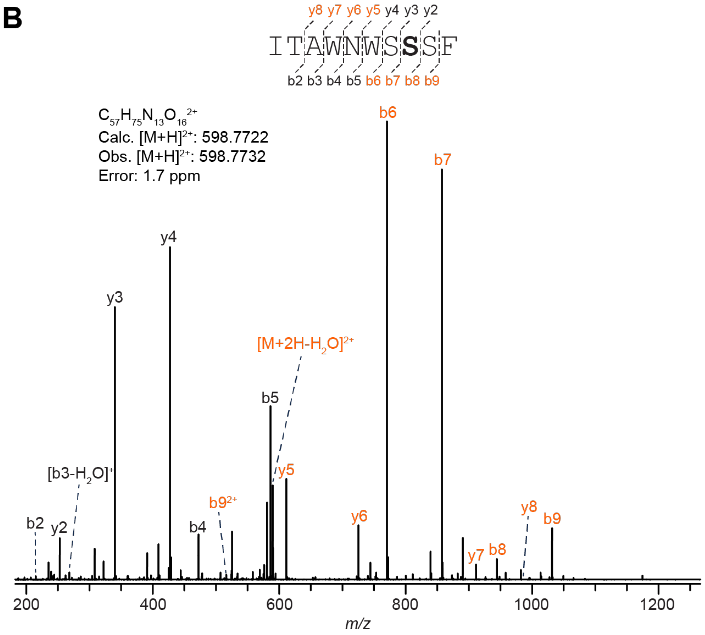

C

| Ion | Calculated Mass | Observed Mass | Error (ppm) |
|---|---|---|---|
| $[M+2H]^{2+}$ | 598.7722 | 598.7732 | 1.7 |
| $[M+2H-H_2O]^{2+}$ | 589.7670 | 589.7678 | 1.4 |
| $b9^+$ | 1031.4582 | 1031.4595 | 1.3 |
| $b9^{2+}$ | 516.2327 | 516.2334 | 1.4 |
| $y2^+$ | 253.1183 | 253.1187 | 1.6 |
| $b8^+$ | 944.4261 | 944.4274 | 1.4 |
| $y3^+$ | 340.1504 | 340.1509 | 1.5 |
| $b7^+$ | 857.3941 | 857.3950 | 1.0 |
| $y4^+$ | 427.1824 | 427.1829 | 1.2 |
| $b6^+$ | 770.3621 | 770.3630 | 1.2 |
| $b5^+$ | 586.2984 | 586.2994 | 1.7 |
| $y5^+$ | 611.2461 | 611.2469 | 1.3 |
| $b4^+$ | 472.2555 | 472.2562 | 1.5 |
| $y6^+$ | 725.2890 | 725.2901 | 1.5 |
| $b3^+$ | 286.1762 | 286.1767 | 1.7 |
| $[b3-H_2O]^+$ | 268.1656 | 268.1660 | 1.5 |
| $y7^+$ | 911.3683 | 911.3694 | 1.2 |
| $b2^+$ | 215.1391 | 215.1394 | 1.4 |
| $y8^+$ | 982.4054 | 982.4069 | 1.5 |

**Figure S9: HR-MS/MS of the DarA-K5S variant with a dhTrp modification.** (A) Hypothetical structure of the modified species based on HR-MS/MS fragmentation. The modification is highlighted in orange. (B) CID spectrum of the -2 Da modified variant. The sequence of the GluC-digested peptide is provided with the substitution bolded. Orange colored daughter ions include the modification. (C) Observed masses and errors from CID analysis.

**A**

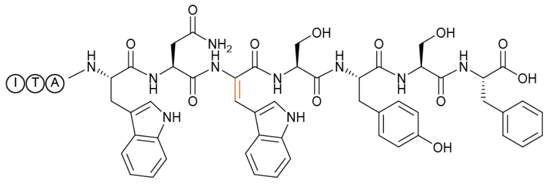

**B**

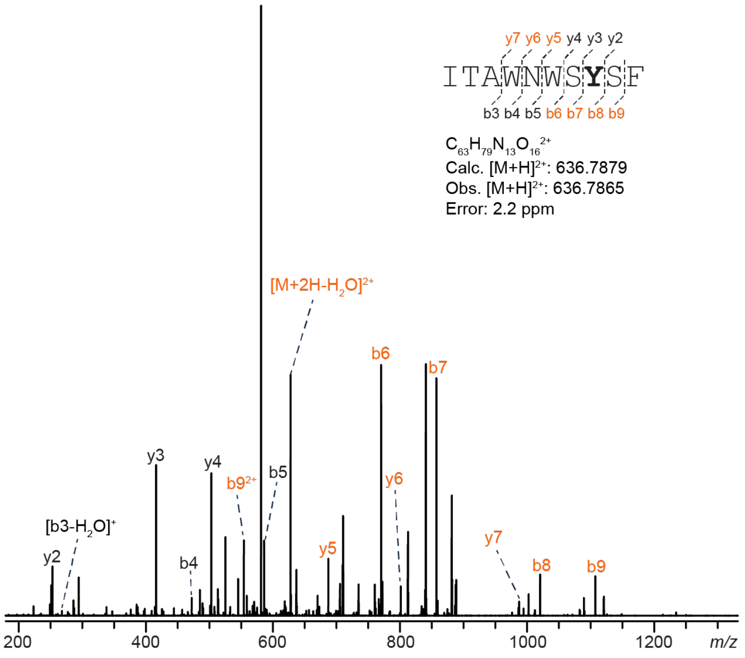

**C**

| Ion | Calculated Mass | Observed Mass | Error (ppm) |
|---|---|---|---|
| $[M+2H]^{2+}$ | 636.7879 | 636.7865 | 2.2 |
| $[M+2H-H_2O]^{2+}$ | 627.7826 | 627.7811 | 2.4 |
| $b9^+$ | 1107.4895 | 1107.4869 | 2.3 |
| $b9^{2+}$ | 554.2484 | 554.2471 | 2.3 |
| $y2^+$ | 253.1183 | 253.1178 | 2.0 |
| $b8^+$ | 1020.4574 | 1020.4553 | 2.1 |
| $y3^+$ | 416.1817 | 416.1808 | 2.2 |
| $b7^+$ | 857.3941 | 857.3922 | 2.2 |
| $y4^+$ | 503.2137 | 503.2126 | 2.2 |
| $b6^+$ | 770.3621 | 770.3606 | 1.9 |
| $b5^+$ | 586.2984 | 586.2972 | 2.0 |
| $y5^+$ | 687.2774 | 687.2758 | 2.3 |
| $b4^+$ | 472.2555 | 472.2545 | 2.1 |
| $y6^+$ | 801.3203 | 801.3187 | 2.0 |
| $[b3-H_2O]^+$ | 268.1656 | 268.165 | 2.2 |
| $y7^+$ | 987.3996 | 987.3975 | 2.1 |

**Figure S10: HR-MS/MS analysis of the DarA-K5Y variant with a dhTrp modification.** (A) Hypothetical structure of the modified species based on HR-MS/MS fragmentation. The modification is highlighted in orange. (B) CID spectrum of the -2 Da modified variant. The sequence of the GluC-digested peptide is provided with the substitution bolded. Orange colored daughter ions include the modification. (C) Observed masses and errors from CID analysis.

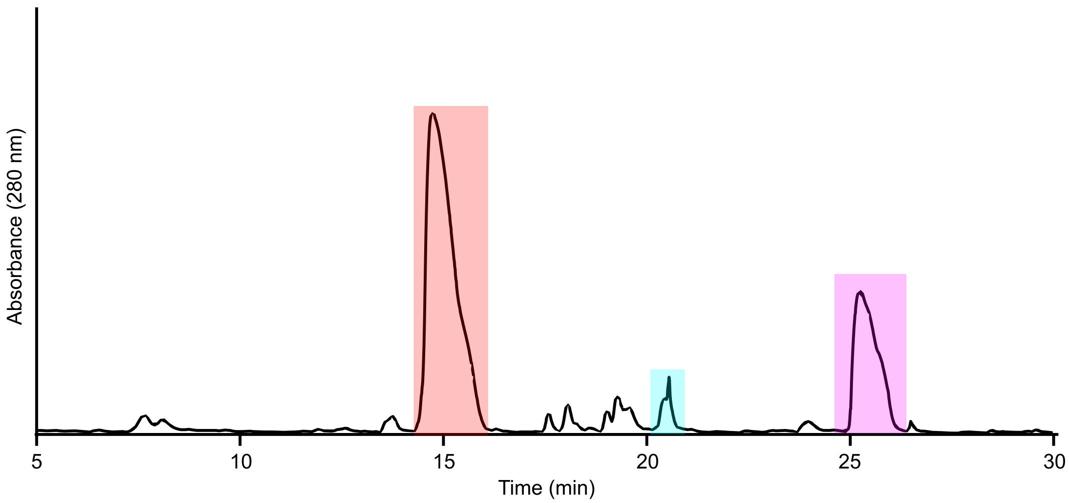

**Figure S11: UHPLC purification of the GluC-digested DarA-W1N variant.** (A) UHPLC chromatogram with the 280 nm absorbance shown. The -2 Da modified species were detected in the light red- and cyan-colored fractions at 15 and 21 min. The magenta-colored fraction at 25 min contained unmodified DarA-W1N.

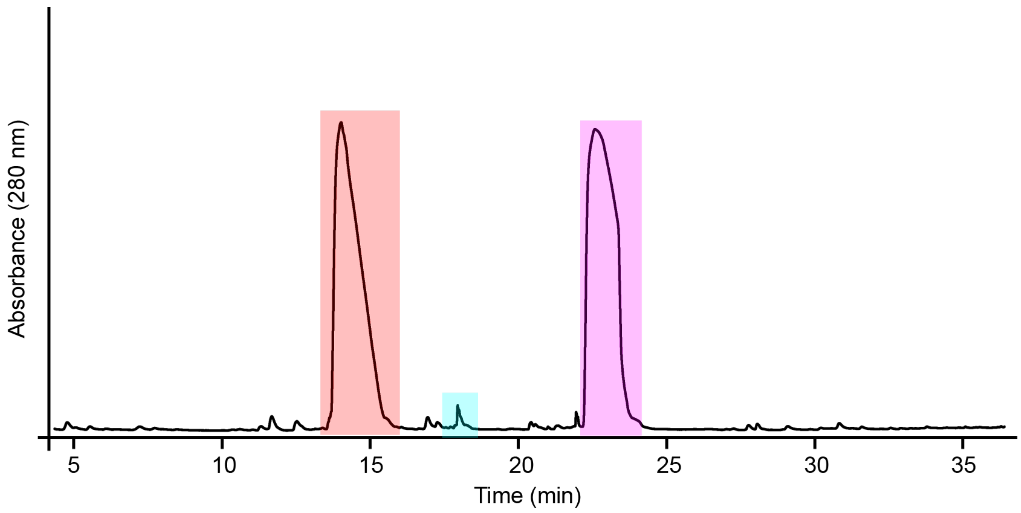

**Figure S12: UHPLC purification of the GluC-digested the DarA-W1D variant.** UHPLC chromatogram with the 280 nm absorbance is shown. The light red- and cyan-colored fractions at 15 and 18 min, respectively, contained the DarA-W1D variant with the -2 Da modification. The magenta fraction at 23 min contained unmodified DarA-W1D.

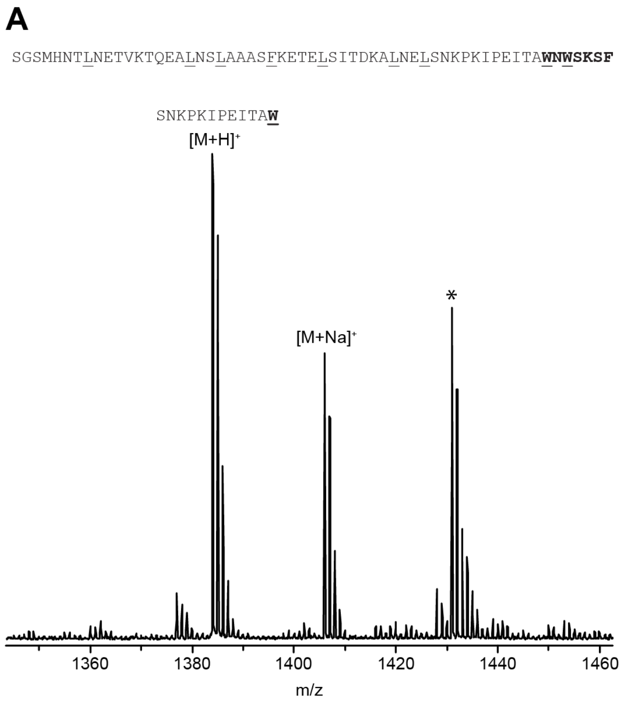

**Figure S13: Digestion of DarA by chymotrypsin.** MALDI-TOF mass spectra of (A) unmodified DarA and (B) Predarobactin (doubly modified DarA). Precursor peptide sequence after TEV protease digestion is provided above each spectrum with residues that chymotrypsin will cut after underlined. The core region is bolded. The mass increase of +12 Da and two missed chymotrypsin cleavages result from ether and C-C crosslink formation. * indicates an unidentified contaminant.

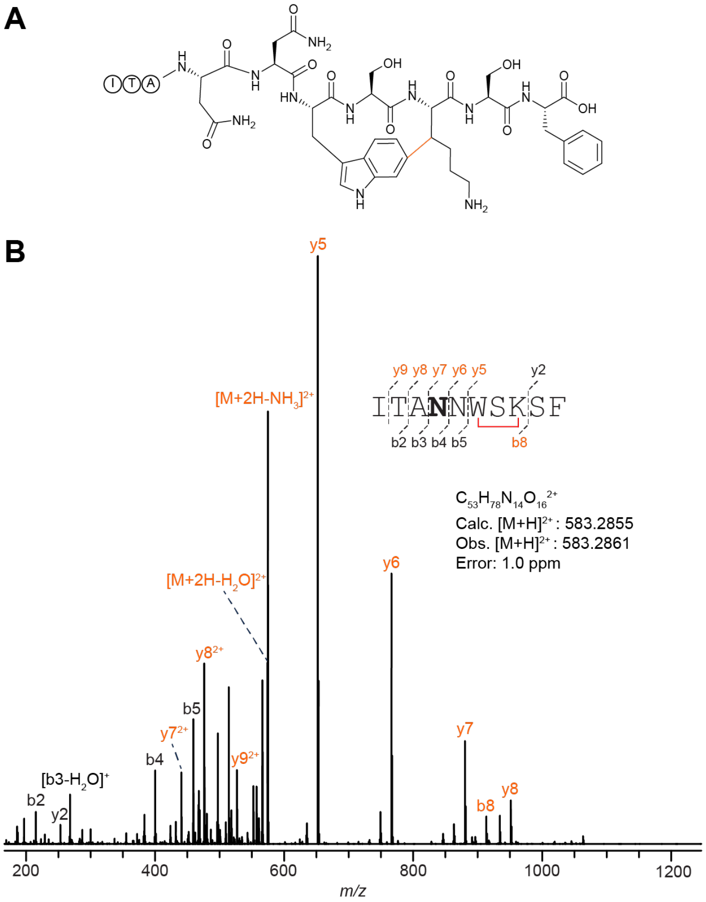

C

| Ion | Calculated Mass | Observed Mass | Error (ppm) |
|---|---|---|---|
| [M+2H]$^{2+}$ | 583.2855 | 583.2861 | 1.03 |
| [M+2H-H$_2$O]$^{2+}$ | 574.2802 | 574.2795 | 1.22 |
| [M+2H-NH$_3$]$^{2+}$ | 574.7722 | 574.7716 | 1.04 |
| y2$^+$ | 253.1183 | 253.118 | 1.19 |
| b8$^+$ | 913.4527 | 913.4512 | 1.64 |
| b5$^+$ | 514.2620 | 514.2613 | 1.36 |
| y5$^+$ | 652.3090 | 652.3078 | 1.84 |
| b4$^+$ | 400.2191 | 400.2185 | 1.50 |
| y6$^+$ | 766.3519 | 766.3506 | 1.70 |
| [b3-H$_2$O]$^+$ | 268.1656 | 268.1653 | 1.12 |
| y7$^+$ | 880.3948 | 880.3935 | 1.48 |
| y7$^{2+}$ | 440.7011 | 440.7005 | 1.36 |
| b2$^+$ | 215.1391 | 215.1388 | 1.39 |
| y8$^+$ | 951.4320 | 951.4304 | 1.68 |
| y8$^{2+}$ | 476.2196 | 476.219 | 1.26 |
| y9$^{2+}$ | 526.7435 | 526.7427 | 1.52 |

**Figure S14: HR-MS/MS analysis of the DarA-W1N variant with a C-C crosslink.** The peptide was collected from the light red-colored fraction in **Figure S11**. (A) Hypothetical structure of the modified species based on HR-MS/MS fragmentation. The connectivity of the linkage is presumed from the known structure of darobactin. The modification is highlighted in orange. (B) CID spectrum of the GluC-digested and UHPLC purified -2 Da modified variant. The sequence of the GluC-digested peptide is provided with the substitution bolded. Orange colored daughter ions include the modification. (C) Observed masses and errors from CID analysis.

A

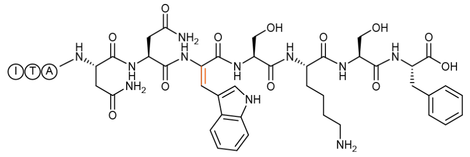

B

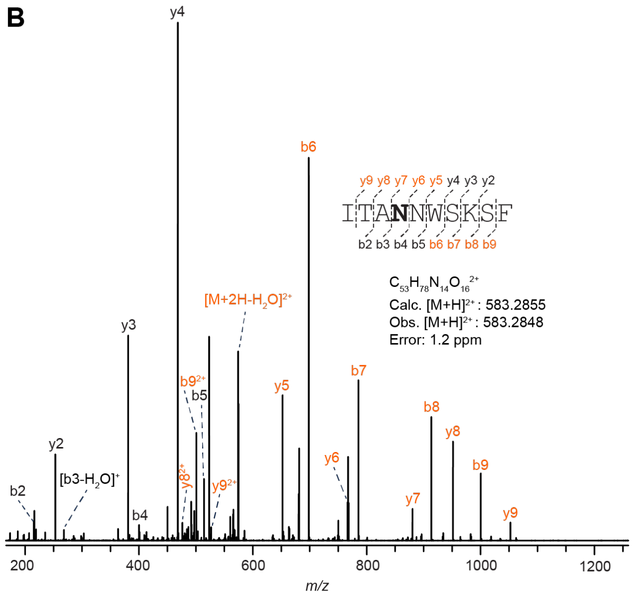

C

| Ion | Calculated Mass | Observed Mass | Error (ppm) |
|---|---|---|---|
| $[M+2H]^{2+}$ | 583.2855 | 583.2848 | 1.20 |
| $[M+2H-H_2O]^{2+}$ | 574.2802 | 574.2791 | 1.92 |
| $b9^+$ | 1000.4847 | 1000.4824 | 2.30 |
| $b9^{2+}$ | 500.7460 | 500.745 | 2.00 |
| $y2^+$ | 253.1183 | 253.1178 | 1.98 |
| $b8^+$ | 913.4527 | 913.4507 | 2.19 |
| $y3^+$ | 381.2133 | 381.2126 | 1.84 |
| $b7^+$ | 785.3577 | 785.356 | 2.16 |
| $y4^+$ | 468.2453 | 468.2443 | 2.14 |
| $b6^+$ | 698.3257 | 698.324 | 2.43 |
| $b5^+$ | 514.2620 | 514.2609 | 2.14 |
| $y5^+$ | 652.3090 | 652.3076 | 2.15 |
| $b4^+$ | 400.2191 | 400.2183 | 2.00 |
| $y6^+$ | 766.3519 | 766.3504 | 1.96 |
| $[b3-H_2O]^+$ | 268.1656 | 268.1651 | 1.86 |
| $y7^+$ | 880.3948 | 880.393 | 2.04 |
| $b2^+$ | 215.1391 | 215.1386 | 2.32 |
| $y8^+$ | 951.4320 | 951.4298 | 2.31 |
| $y8^{2+}$ | 476.2196 | 476.2186 | 2.10 |
| $y9^+$ | 1052.4796 | 1052.477 | 2.47 |
| $y9^{2+}$ | 526.7435 | 526.7424 | 2.09 |

**Figure S15: HR-MS/MS analysis of the DarA-W1N variant with a dhTrp modification.** The peptide was collected from the cyan-colored fraction in **Figure S11**. (A) Hypothetical structure of the modified species based on HR-MS/MS fragmentation. The connectivity of the linkage is presumed from the known structure of darobactin. The modification is highlighted in orange. (B) CID spectrum of the GluC-digested and UHPLC purified -2 Da modified variant. The sequence of the GluC-digested peptide is provided with the substitution bolded. Orange colored daughter ions include the modification. (C) Observed masses and errors from CID analysis.

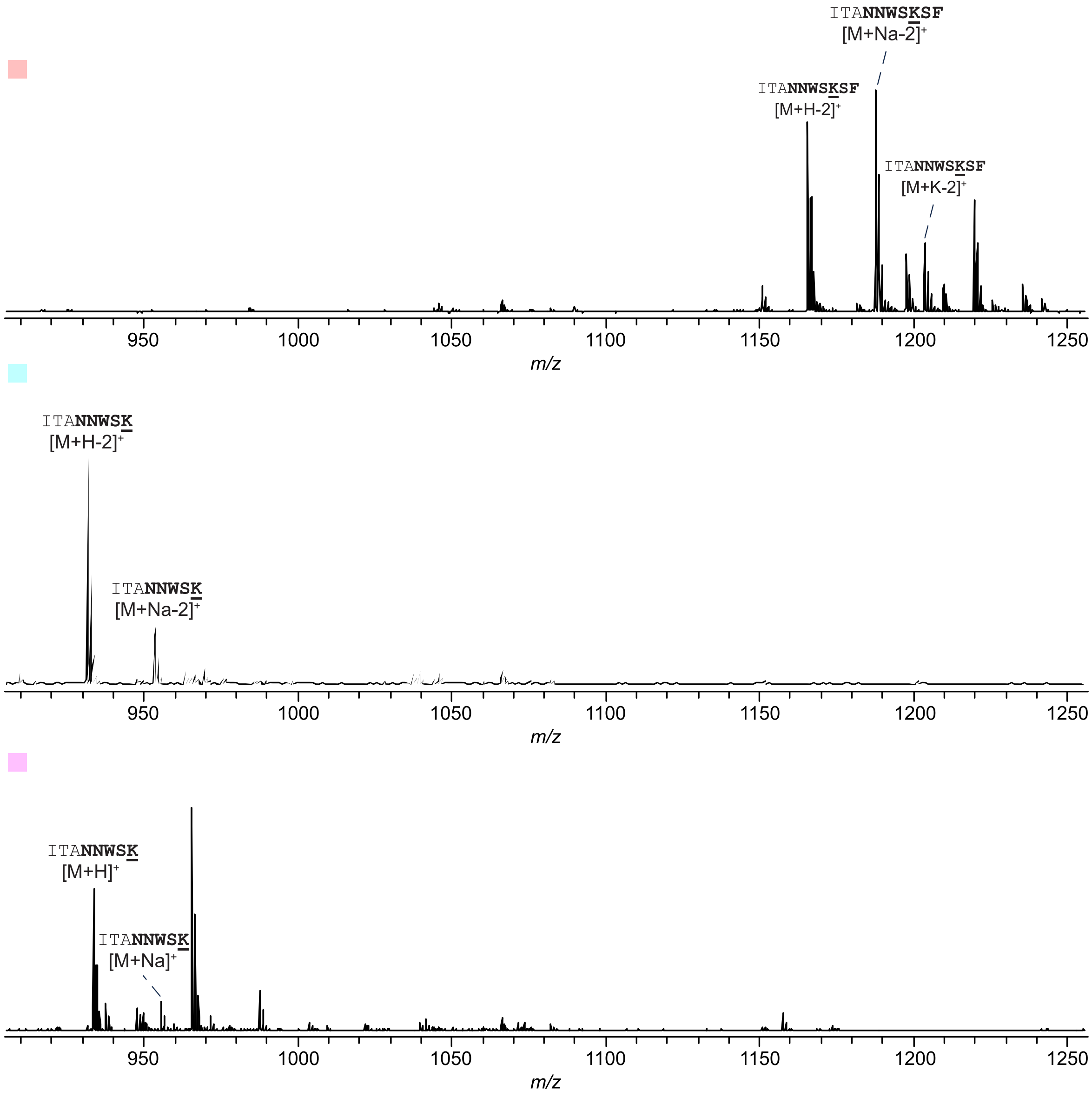

**Figure S16: Digestion of UHPLC purified DarA-W1N by trypsin.** Trypsin digests on each fraction of interest were performed to check for the presence of a C-C crosslink. The light red, cyan, and magenta squares correspond to the same fractions shown in **Figure S11**. Underlined residues indicate potential tryptic sites. The DarA core region is bolded.

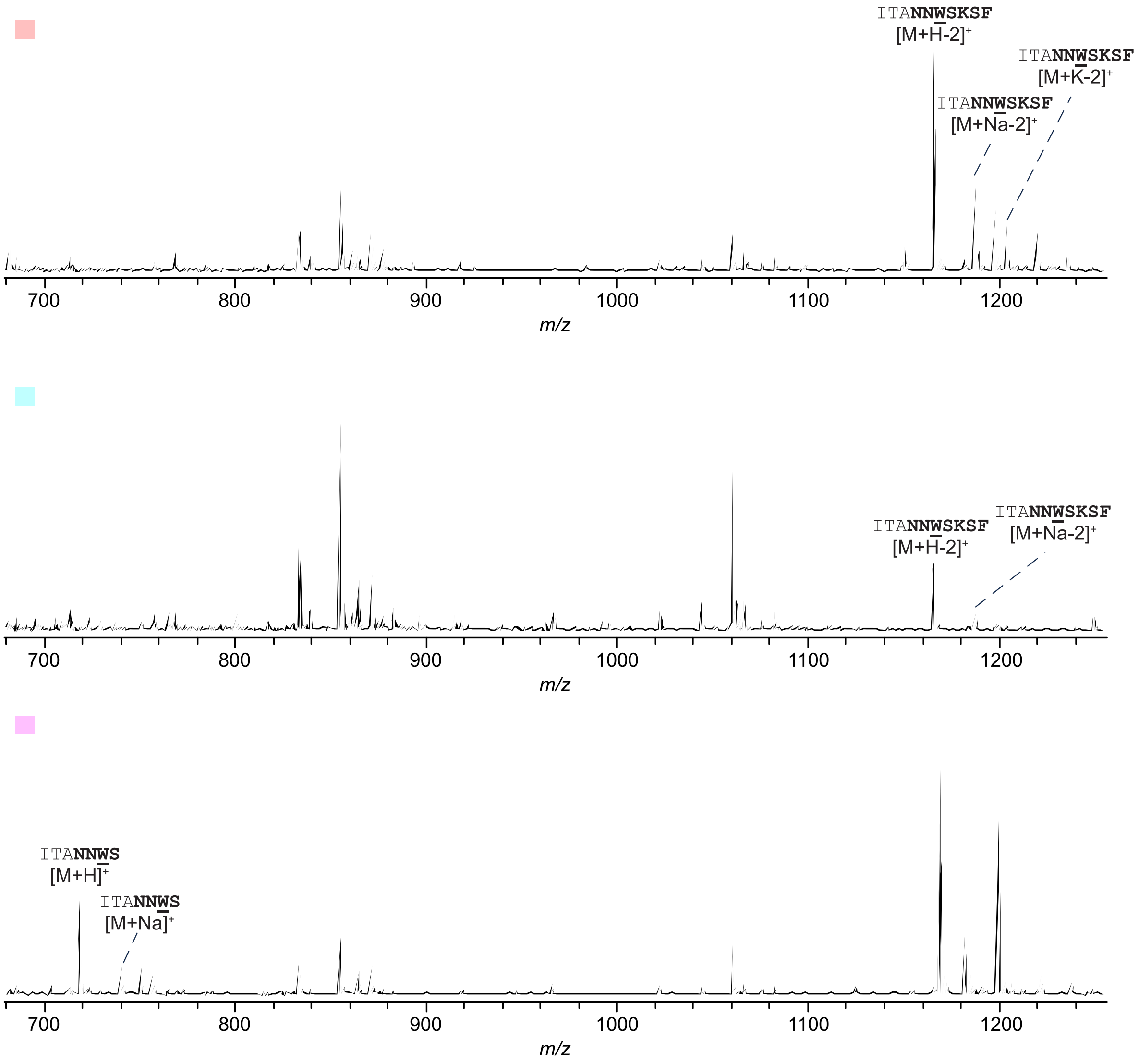

**Figure S17: Digestion of UHPLC purified DarA-W1N by chymotrypsin.** Chymotrypsin digests on each fraction of interest were performed to check for the presence of a C-C crosslink. The light red, cyan, and magenta squares correspond to the same fractions shown in **Figure S11**. Underlined residues indicate potential chymotryptic sites. The DarA core region is bolded.

**A**

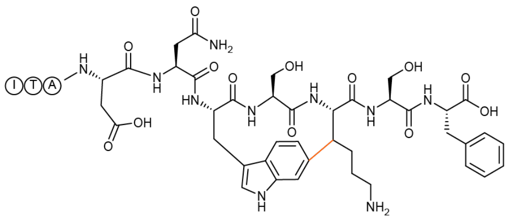

**B**

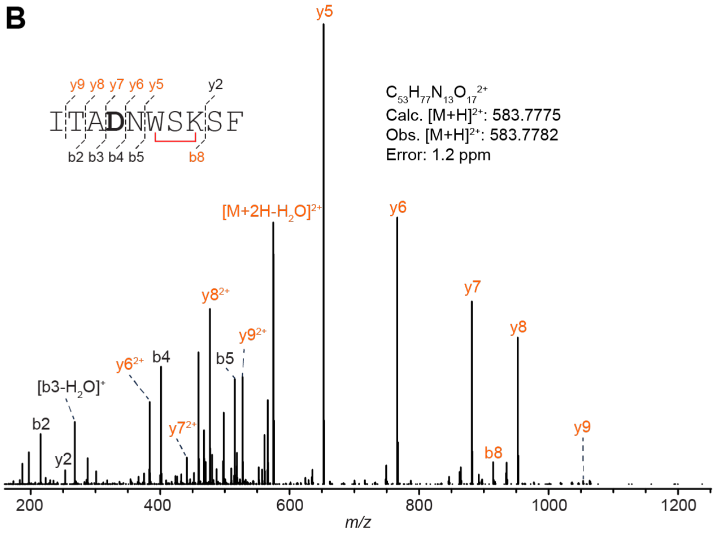

**C**

| Ion | Calculated Mass | Observed Mass | Error (ppm) |
|---|---|---|---|
| $[M+2H]^{2+}$ | 583.7775 | 583.7782 | 1.2 |
| $[M+2H-H_2O]^{2+}$ | 574.7722 | 574.7717 | 0.9 |
| $y2^{+}$ | 253.1183 | 253.1181 | 0.8 |
| $b8^{+}$ | 1001.4687 | 1001.4672 | 1.5 |
| $b5^{+}$ | 515.2461 | 515.2455 | 1.2 |
| $y5^{+}$ | 652.3090 | 652.3082 | 1.2 |
| $b4^{+}$ | 401.2031 | 401.2027 | 1.0 |
| $y6^{+}$ | 766.3519 | 766.351 | 1.2 |
| $y6^{2+}$ | 383.6796 | 383.6793 | 0.8 |
| $b3^{+}$ | 286.1762 | 286.1759 | 1.0 |
| $[b3-H_2O]^{+}$ | 268.1656 | 268.1654 | 0.7 |
| $y7^{+}$ | 881.3789 | 881.3777 | 1.4 |
| $y7^{2+}$ | 441.1931 | 441.1927 | 0.9 |
| $b2^{+}$ | 215.1391 | 215.1388 | 1.4 |
| $y8^{+}$ | 952.4160 | 952.4147 | 1.4 |
| $y8^{2+}$ | 476.7116 | 476.7112 | 0.8 |
| $y9^{+}$ | 1053.4636 | 1053.462 | 1.5 |
| $y9^{2+}$ | 527.2355 | 527.235 | 0.9 |

**Figure S18: HR-MS/MS analysis of the DarA-W1D variant with a C-C crosslink.** The peptide was collected from the light red fraction in **Figure S12**. (A) Hypothetical structure of the modified species based on HR-MS/MS fragmentation. The connectivity of the linkage is presumed from the known structure of darobactin. The modification is highlighted in orange. (B) CID spectrum of the GluC-digested and UHPLC purified -2 Da modified variant. The sequence of the GluC-digested peptide is provided with the substitution bolded. Orange colored daughter ions include the modification. (C) Observed masses and errors from CID analysis.

**A**

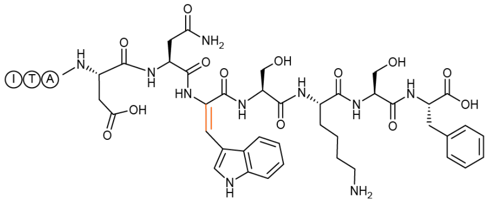

**B**

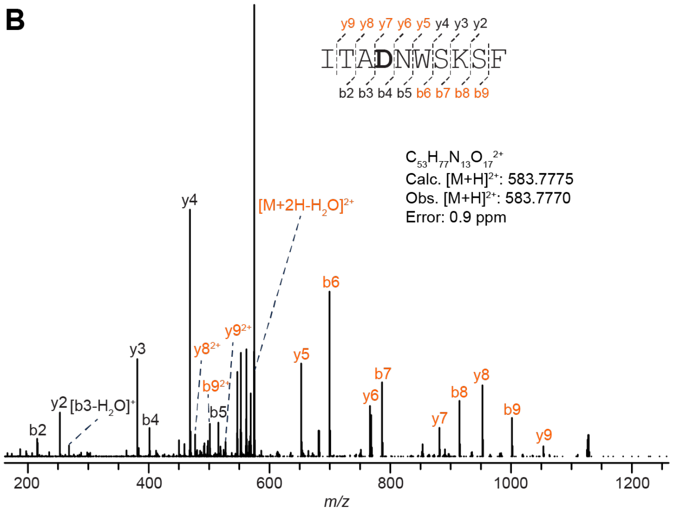

**C**

| Ion | Calculated Mass | Observed Mass | Error (ppm) |
|---|---|---|---|
| $[M+2H]^{2+}$ | 583.7775 | 583.777 | 0.9 |
| $[M+2H-H_2O]^{2+}$ | 574.7722 | 574.7713 | 1.6 |
| $b9^+$ | 1001.4687 | 1001.4681 | 0.6 |
| $b9^{2+}$ | 501.2380 | 501.2379 | 0.2 |
| $y2^+$ | 253.1183 | 253.1182 | 0.4 |
| $b8^+$ | 914.4367 | 914.4363 | 0.4 |
| $y3^+$ | 381.2133 | 381.2132 | 0.3 |
| $b7^+$ | 786.3417 | 786.3415 | 0.3 |
| $y4^+$ | 468.2453 | 468.2451 | 0.4 |
| $b6^+$ | 699.3097 | 699.3093 | 0.6 |
| $b5^+$ | 515.2461 | 515.2458 | 0.6 |
| $y5^+$ | 652.3090 | 652.3087 | 0.5 |
| $b4^+$ | 401.2031 | 401.203 | 0.2 |
| $y6^+$ | 766.3519 | 766.3517 | 0.3 |
| $y6^{2+}$ | 383.6796 | 383.6795 | 0.3 |
| $[b3-H_2O]^+$ | 268.1656 | 268.1655 | 0.4 |
| $y7^+$ | 881.3789 | 881.3785 | 0.5 |
| $b2^+$ | 215.1391 | 215.1389 | 0.9 |
| $y8^+$ | 952.4160 | 952.4153 | 0.7 |
| $y8^{2+}$ | 476.7116 | 476.7115 | 0.2 |
| $y9^+$ | 1053.4636 | 1053.463 | 0.6 |
| $y9^{2+}$ | 527.2355 | 527.2353 | 0.4 |

**Figure S19: HR-MS/MS analysis of the DarA-W1D variant with a dhTrp modification.** Peptide was collected from the cyan fraction in **Figure S12**. (A) Hypothetical structure of the modified species based on HR-MS/MS fragmentation. The modification is highlighted in orange. (B) CID spectrum of the GluC-digested and UHPLC purified -2 Da modified variant. The sequence of the GluC-digested peptide is provided with the substitution bolded. Orange colored daughter ions include the modification. (C) Observed masses and errors from CID analysis.

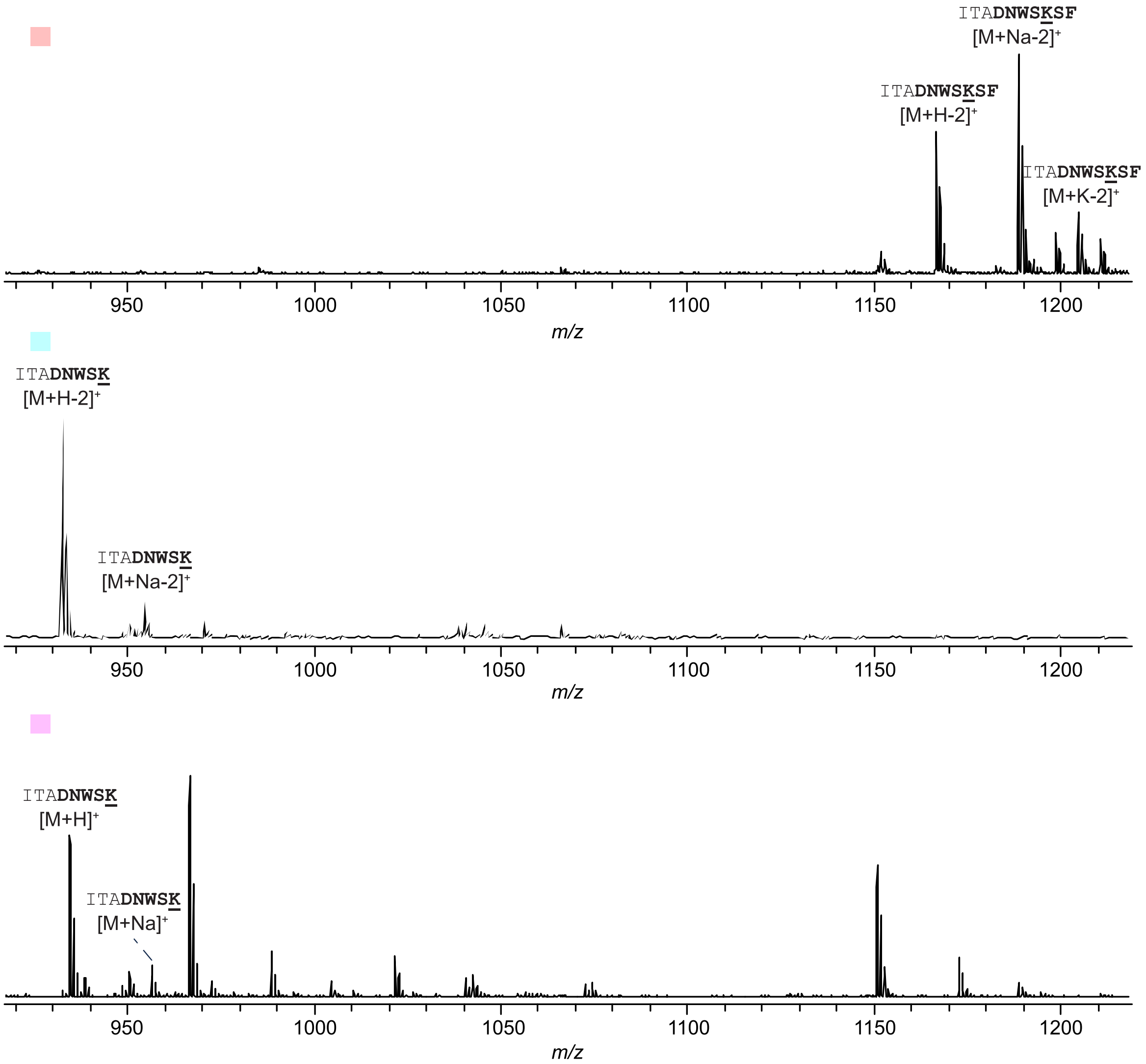

**Figure S20: Digestion of UHPLC purified DarA-W1D by trypsin.** Trypsin digests on each fraction of interest were performed to check for the presence of a C-C crosslink. The light red, cyan, and magenta squares correspond to the same fractions shown in **Figure S12**. Underlined residues indicate tryptic sites. The core region is bolded.

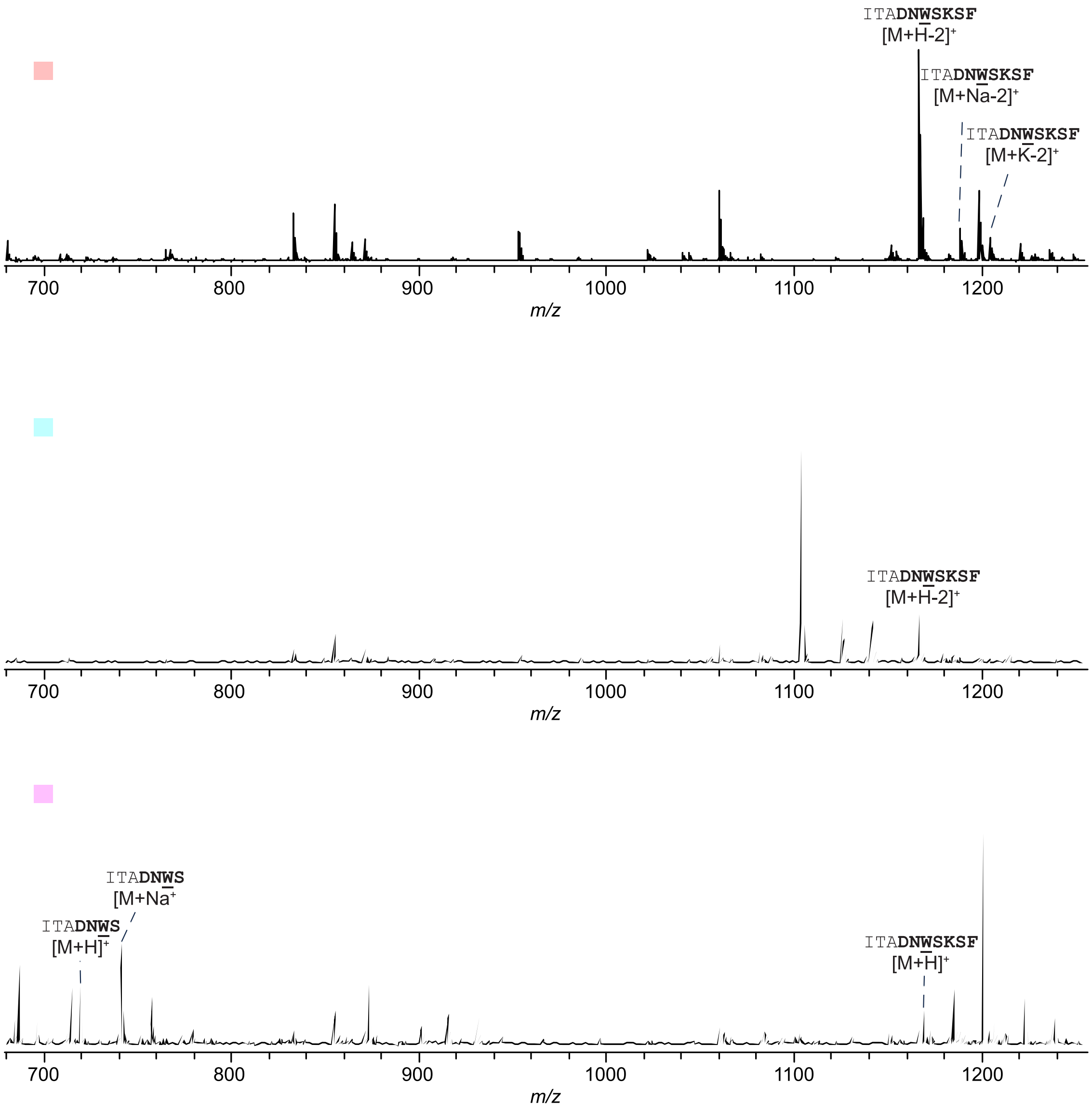

**Figure S21: Digestion of UHPLC purified DarA-W1D by chymotrypsin.** Chymotrypsin digests on each fraction of interest were performed to check for the presence of a C-C crosslink. The light red, cyan, and magenta squares correspond to the same fractions shown in **Figure S12**. Underlined residues indicate chymotryptic sites. The core region is bolded.

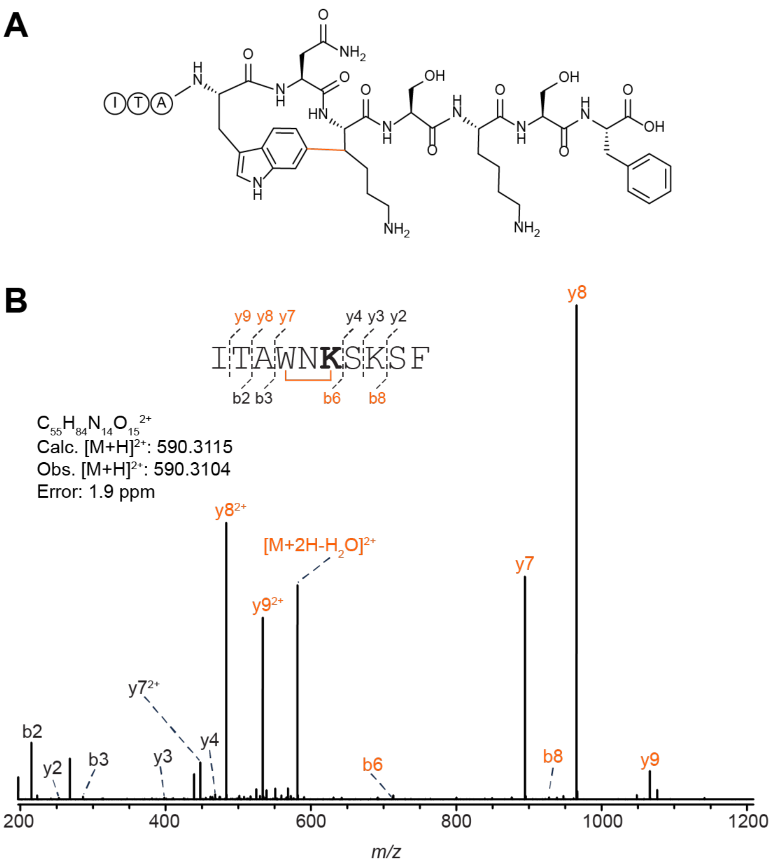

C

| Ion | Calculated Mass | Observed Mass | Error (ppm) |
|---|---|---|---|
| $[M+2H]^{2+}$ | 590.3116 | 590.3104 | 2.0 |
| $[M+2H-H_2O]^{2+}$ | 581.3063 | 581.3046 | 2.9 |
| $y2^+$ | 253.1183 | 253.1177 | 2.4 |
| $b8^+$ | 927.5048 | 927.5022 | 2.8 |
| $y3^+$ | 381.2133 | 381.2123 | 2.6 |
| $y4^+$ | 468.2453 | 468.244 | 2.8 |
| $b6^+$ | 712.3778 | 712.3756 | 3.1 |
| $b3^+$ | 286.1762 | 286.1755 | 2.4 |
| $y7^+$ | 894.4469 | 894.4441 | 3.1 |
| $b2^+$ | 215.1391 | 215.1384 | 3.3 |
| $y8^+$ | 965.484 | 965.4808 | 3.3 |
| $y8^{2+}$ | 483.2457 | 483.2443 | 2.9 |
| $y9^+$ | 1066.5317 | 1066.5286 | 2.9 |
| $y9^{2+}$ | 533.7695 | 533.768 | 2.8 |

**Figure S22: HR-MS/MS analysis of the DarA-W3K variant with a C-C crosslink. (A**) Hypothetical structure of the modified species based on HR-MS/MS fragmentation. The connectivity of the linkage is presumed from the known structure of darobactin. The modification is highlighted in orange. (B) CID spectrum of the -2 Da species modified variant. The sequence of the GluC-digested peptide is provided with the substitution bolded. Orange colored daughter ions include the modification. (C) Observed masses and errors from CID analysis.

**A**

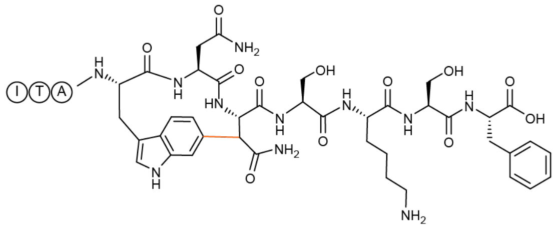

**B**

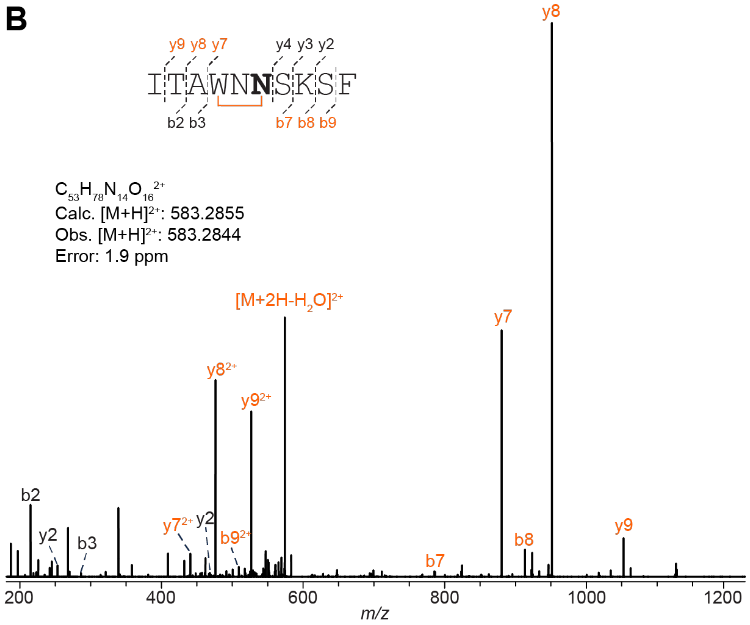

**C**

| Ion | Calculated Mass | Observed Mass | Error (ppm) |
|---|---|---|---|
| $[M+2H]^{2+}$ | 583.2855 | 583.2844 | 1.9 |
| $[M+2H-H_2O]^{2+}$ | 574.2803 | 574.2786 | 3.0 |
| $b9^{2+}$ | 500.7460 | 500.7447 | 2.6 |
| $y2^{+}$ | 253.1183 | 253.1177 | 2.4 |
| $b8^{+}$ | 913.4527 | 913.4501 | 2.8 |
| $b7^{+}$ | 785.3578 | 785.3559 | 2.4 |
| $y4^{+}$ | 468.2453 | 468.2440 | 2.8 |
| $b3^{+}$ | 286.1762 | 286.1754 | 2.8 |
| $y7^{+}$ | 880.3949 | 880.3923 | 3.0 |
| $y7^{2+}$ | 440.7011 | 440.6999 | 2.7 |
| $b2^{+}$ | 215.1391 | 215.1384 | 3.3 |
| $y8^{+}$ | 951.4320 | 951.4290 | 3.2 |
| $y8^{2+}$ | 476.2197 | 476.2183 | 2.9 |
| $y9^{+}$ | 1052.4797 | 1052.4765 | 3.0 |
| $y9^{2+}$ | 526.7435 | 526.7421 | 2.7 |

**Figure S23: HR-MS/MS analysis of the DarA-W3N variant with a C-C crosslink. (A)** Hypothetical structure of the modified species based on HR-MS/MS fragmentation. The connectivity of the linkage is presumed from the known structure of darobactin. The modification is highlighted in orange. (B) CID spectrum of the -2 Da modified variant. The sequence of the GluC-digested peptide is provided with the substitution bolded. Orange colored daughter ions include the modification. (C) Observed masses and errors from CID analysis.

A

B

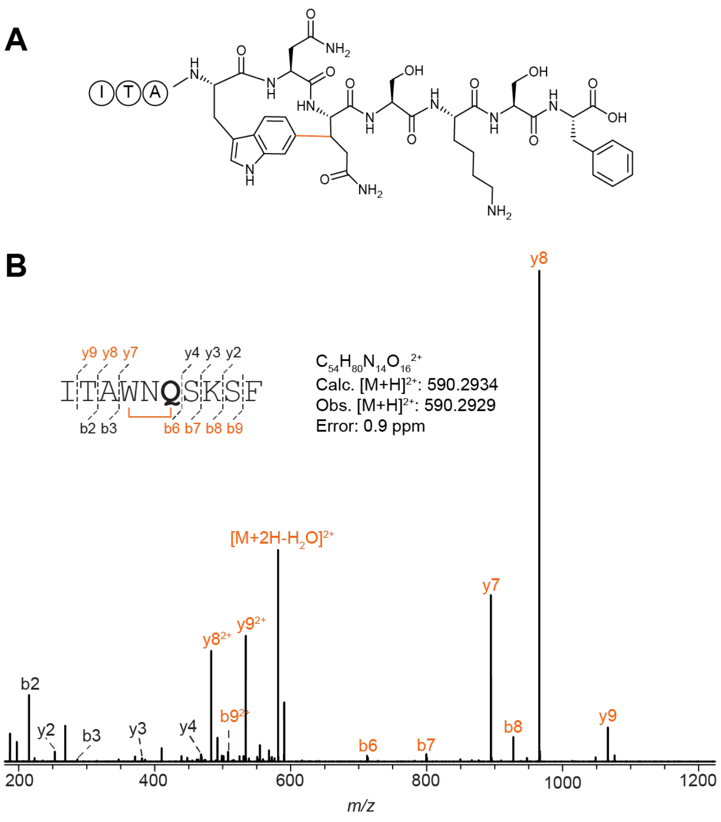

C

| Ion | Calculated Mass | Observed Mass | Error (ppm) |
|---|---|---|---|
| $[M+2H]^{2+}$ | 590.2934 | 590.2929 | 0.8 |
| $[M+2H-H_2O]^{2+}$ | 581.2881 | 581.2872 | 1.5 |
| $b9^{+}$ | 1014.5004 | 1014.4997 | 0.7 |
| $b9^{2+}$ | 507.7539 | 507.7531 | 1.6 |
| $y2^{+}$ | 253.1183 | 253.1180 | 1.2 |
| $b8^{+}$ | 927.4684 | 927.4670 | 1.5 |
| $y3^{+}$ | 381.2133 | 381.2128 | 1.3 |
| $b7^{+}$ | 799.3734 | 799.3726 | 1.0 |
| $y4^{+}$ | 468.2453 | 468.2447 | 1.3 |
| $b6^{+}$ | 712.3414 | 712.3400 | 2.0 |
| $b3^{+}$ | 286.1762 | 286.1758 | 1.4 |
| $y7^{+}$ | 894.4105 | 894.4093 | 1.3 |
| $b2^{+}$ | 215.1391 | 215.1388 | 1.4 |
| $y8^{+}$ | 965.4477 | 965.4461 | 1.7 |
| $y8^{2+}$ | 483.2275 | 483.2268 | 1.4 |
| $y9^{+}$ | 1066.4953 | 1066.4936 | 1.6 |
| $y9^{2+}$ | 533.7513 | 533.7506 | 1.3 |

**Figure S24: HR-MS/MS analysis of the DarA-W3Q variant with a C-C crosslink.** (A) Hypothetical structure of the modified species based on HR-MS/MS fragmentation. The connectivity of the linkage is presumed from the known structure of darobactin. The modification is highlighted in orange. (B) CID spectrum of the -2 Da modified variant. The sequence of the GluC-digested peptide is provided with the substitution bolded. Orange colored daughter ions include the modification. (C) Observed masses and errors from CID analysis.

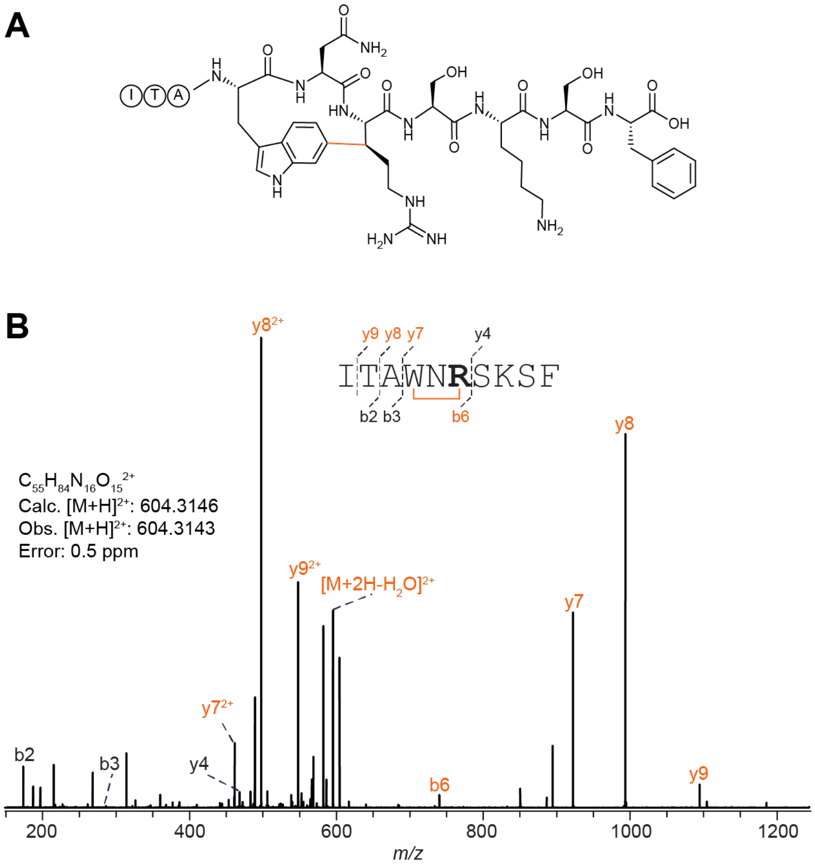

C

| Ion | Calculated Mass | Observed Mass | Error (ppm) |
|---|---|---|---|
| $[M+2H]^{2+}$ | 604.3146 | 604.3143 | 0.5 |
| $[M+2H-H_2O]^{2+}$ | 595.3093 | 595.3086 | 1.2 |
| $y4^+$ | 468.2453 | 468.2449 | 0.9 |
| $b6^+$ | 740.3830 | 740.3833 | 0.4 |
| $b4^+$ | 472.2555 | 472.2549 | 1.3 |
| $b3^+$ | 286.1762 | 286.1761 | 0.3 |
| $y7^+$ | 922.4531 | 922.4521 | 1.1 |
| $y7^{2+}$ | 461.7302 | 461.7296 | 1.3 |
| $b2^+$ | 215.1391 | 215.1388 | 1.4 |
| $y8^+$ | 993.4902 | 993.4890 | 1.2 |
| $y8^{2+}$ | 497.2488 | 497.2482 | 1.2 |
| $y9^+$ | 1094.5379 | 1094.5363 | 1.5 |
| $y9^{2+}$ | 547.7726 | 547.7719 | 1.3 |

**Figure S25: HR-MS/MS analysis of the DarA-W3R variant with a C-C crosslink.** (A) Structure of the modified species based on HR-MS/MS fragmentation. The modification is highlighted in orange (stereochemistry was later confirmed by NMR spectroscopy). (B) CID spectrum of the -2 Da modified variant. The sequence of the GluC-digested peptide is provided with the substitution bolded. Orange colored daughter ions include the modification. (C) Observed masses and errors from CID analysis.

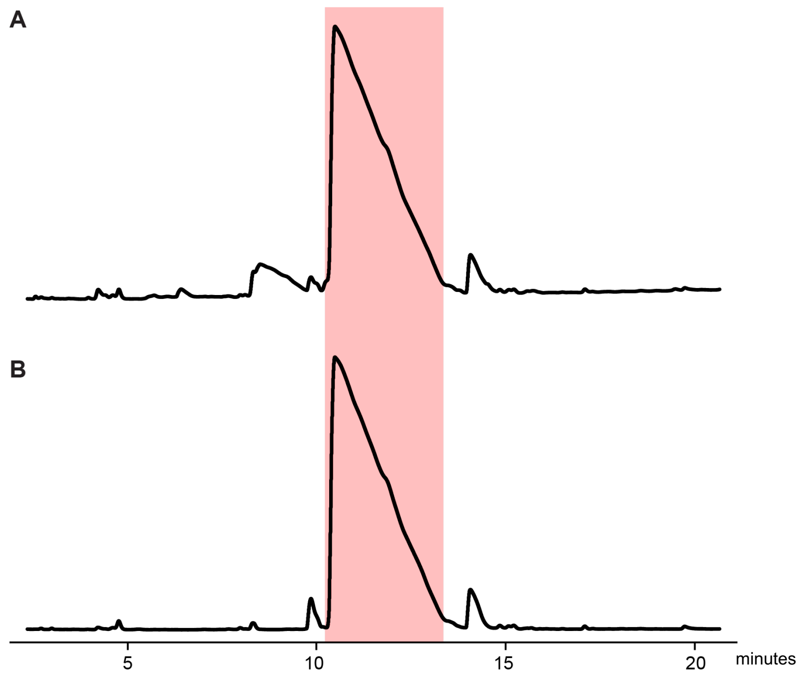

**Figure S26: UHPLC purification of the GluC-digested DarA-W3R variant.** Chromatograms with the (A) 220 nm and (B) 280 nm absorbances are shown from the C18 analytical purification. The light red-colored fraction eluting at 11 min contained the DarA-W3R variant with a -2 Da modification and was collected for NMR.

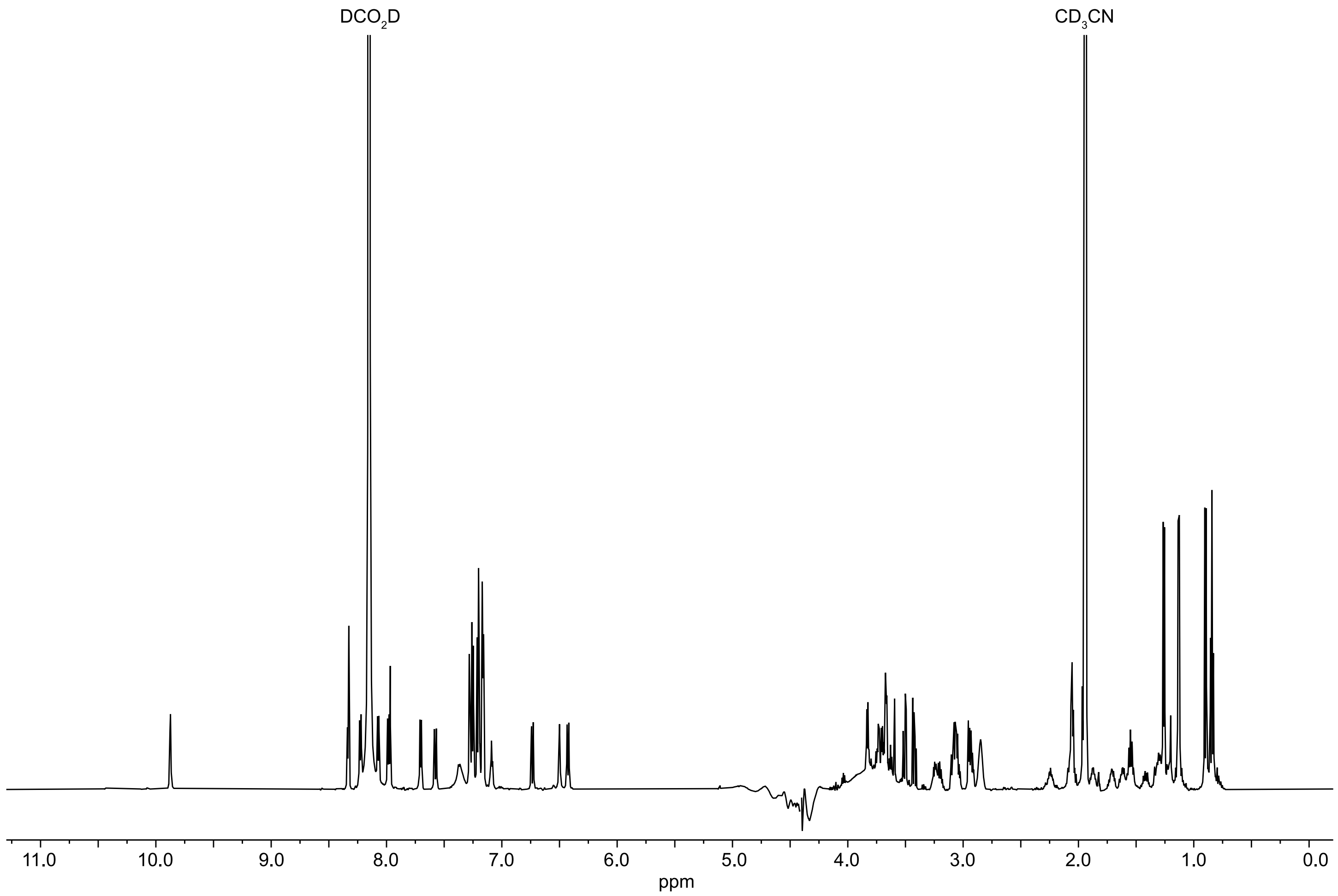

**Figure S27: [1]H NMR spectrum of the GluC-digested DarA-W3R variant.** Data were acquired in 66% $H_2O$: 32% $CD_3CN$: 2% $DCO_2D$ using a water-suppression (WET) method.

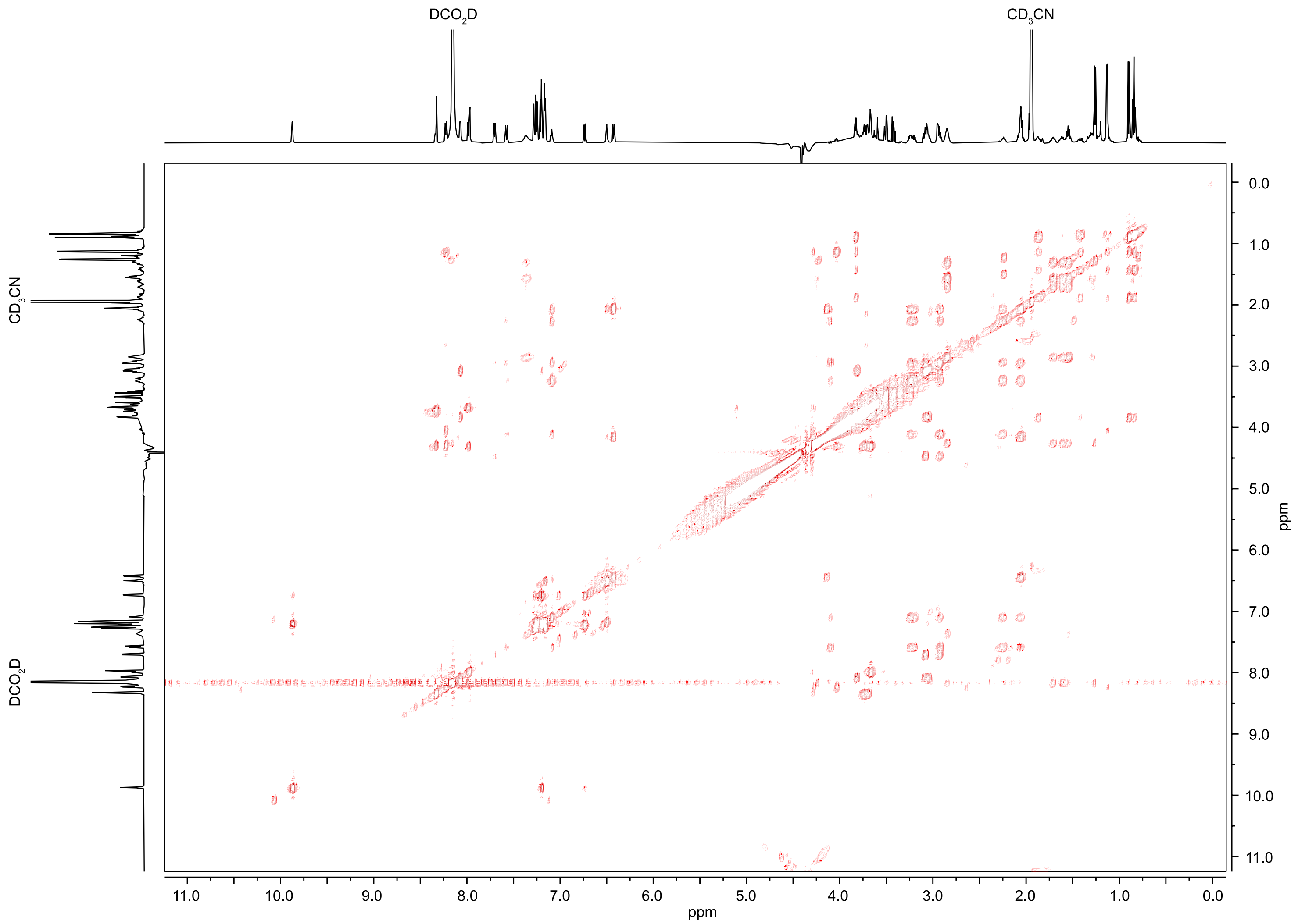

**Figure S28: $^{1}H$-$^{1}H$ TOCSY NMR spectrum of the GluC-digested DarA-W3R variant.** Data were acquired in 66% $H_2O$: 32% $CD_3CN$: 2% $DCO_2D$ using a water-suppression (WET) method.

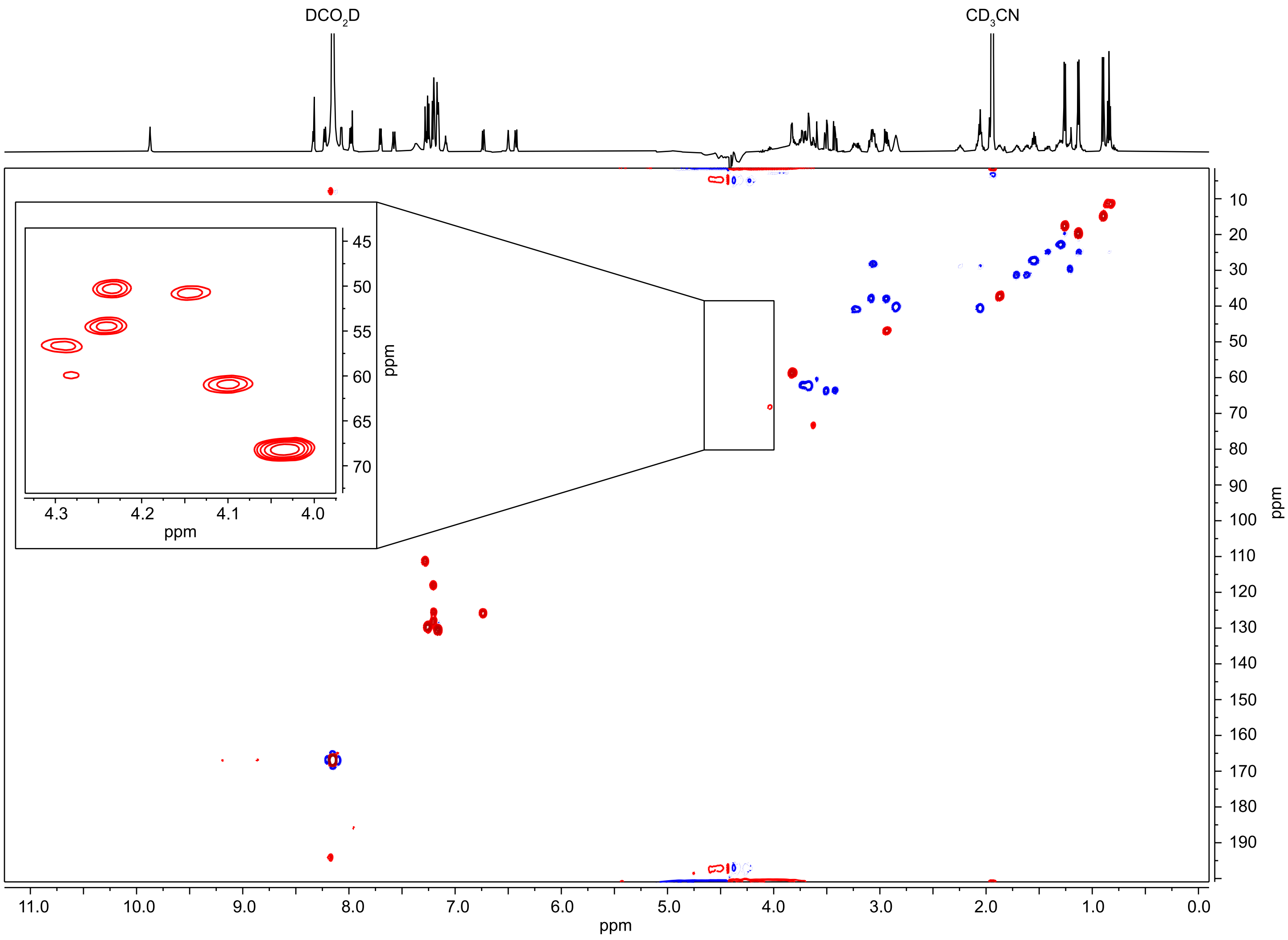

**Figure S29: $^{1}H$-$^{13}C$ HSQC NMR spectrum of the GluC digested DarA-W3R variant.** Data were acquired in 66% $H_2O$: 32% $CD_3CN$: 2% $DCO_2D$ using a water-suppression (WET) method. The CHα cross peaks are inset.

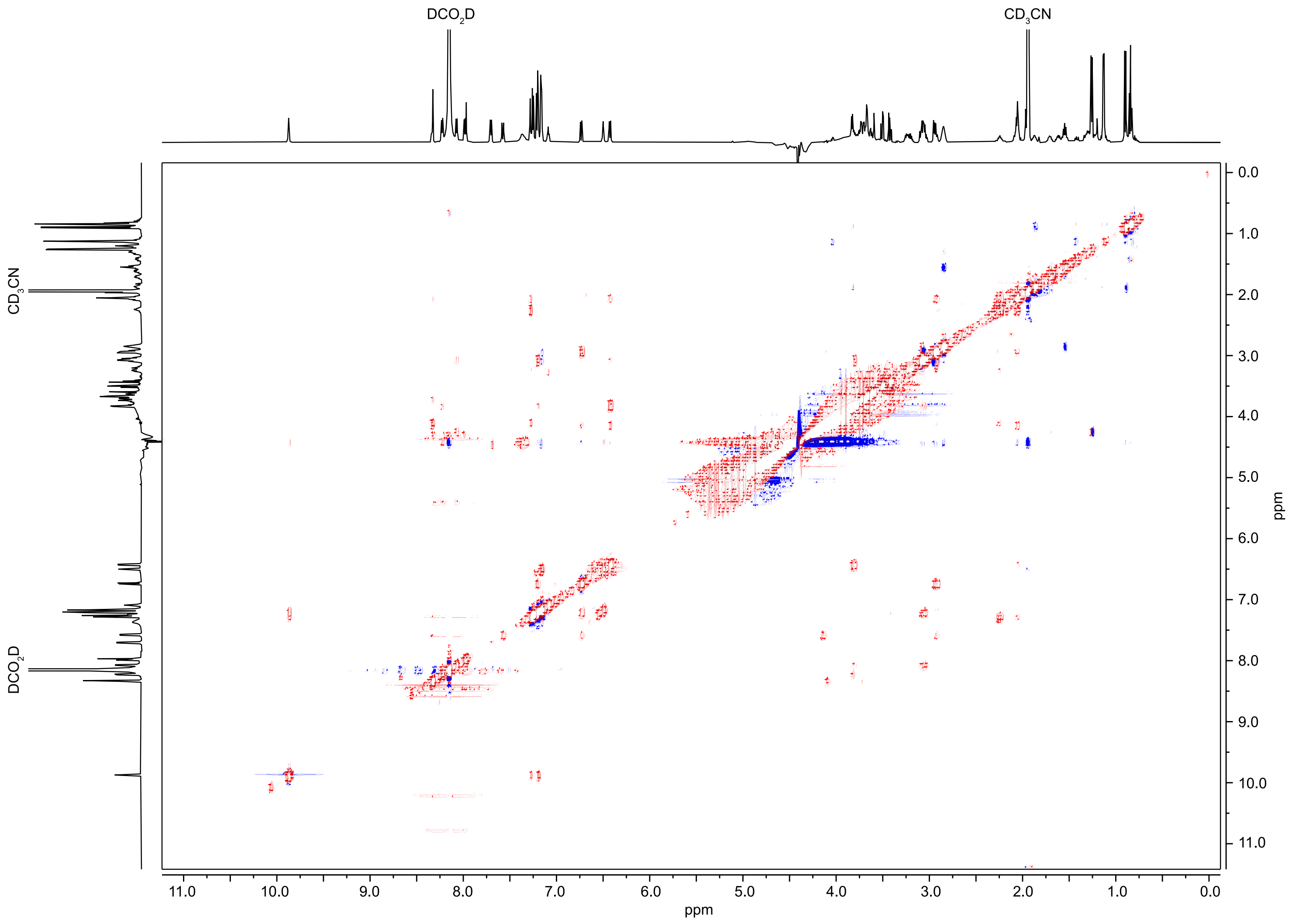

**Figure S30: [1]H-[1]H NOESY NMR spectrum of the GluC-digested DarA-W3R variant**. Data were acquired in 66% $H_2O$: 32% $CD_3CN$: 2% $DCO_2D$ using a water-suppression (WET) method.

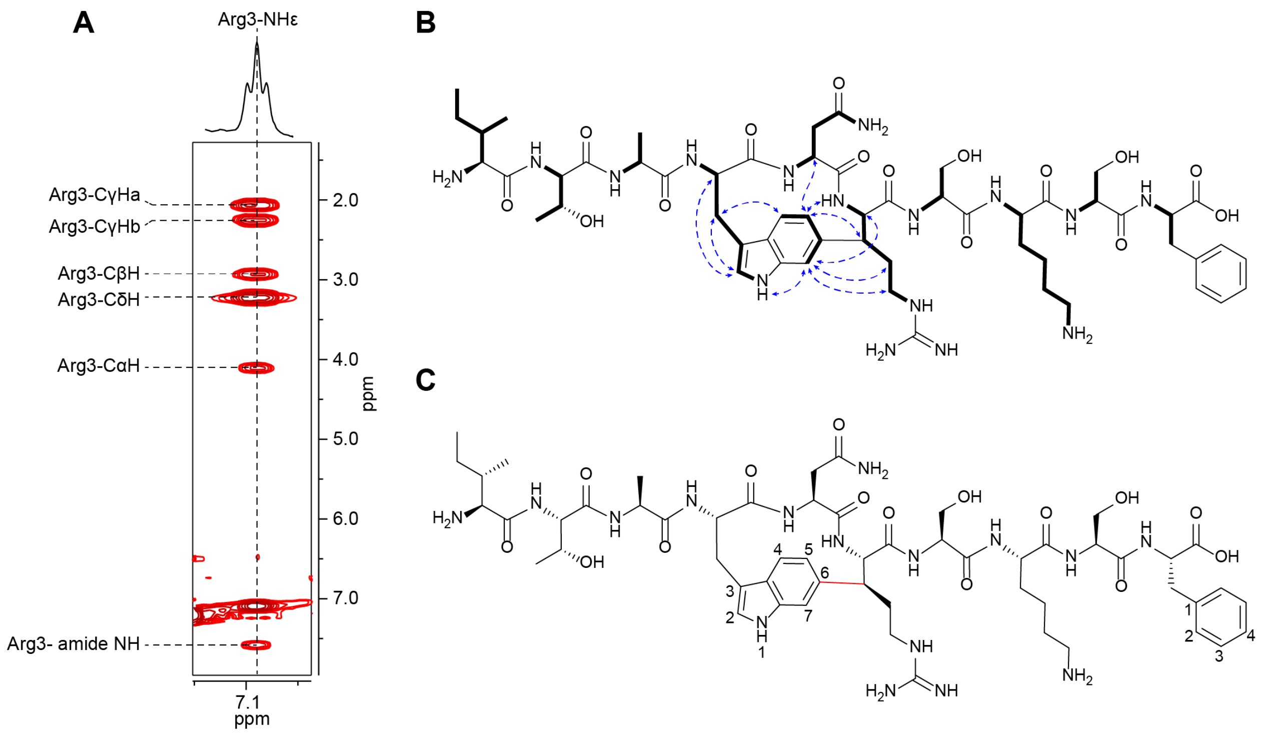

**Figure S31: Additional NMR information for the GluC-digested DarA-W3R variant.** (A) $^{1}$H-$^{1}$H TOCSY spectrum of Arg3. (B) $^{1}$H-$^{1}$H TOCSY of the full GluC-digsted W3R peptide with important $^{1}$H-$^{1}$H NOSEY correlations that enabled sterochemical determination indicated. (C) Numbered W3R structure**.** Numbering of atoms directly corresponds to those in **Table S2.**

**Table S2: NMR Chemical Shifts Assignments for the GluC-digested DarA-W3R variant.** Data were acquired in 66% $H_2O$: 32% $CD_3CN$: 2% $DCO_2D$ using a water-suppression (WET) method.

| Atom | $\delta_H$, mult. ($J$ in Hz) | $\delta_C$ | | Atom | $\delta_H$, mult. ($J$ in Hz) | $\delta_C$ |
|---|---|---|---|---|---|---|
| Ile (-3) | | | | Arg 3 | | |
| Amide NH | 8.20, overlapped | NH | | Amide NH | 7.58, d (10.3) | NH |
| α | 3.83, t (5.3) | 58.7 (CH) | | α | 4.1, t (11.2) | 61 (CH) |
| β | 1.87, ddd (10.4, 7.0, 4.0) | 37.2 (CH) | | β | 2.93, m | 46.9 (CH) |
| γ2 | 0.9, d (7.2) | 14.8 ($CH_3$) | | $γ_a$ | 2.05, dd (7.5, 5.4) | 28.7 ($CH_2$) |
| $γ1_a$ | 1.13, overlapped | 24.9 ($CH_2$) | | $γ_b$ | 2.24, dt (13.2, 6.7) | 28.7 ($CH_2$) |
| $γ1_b$ | 1.42, ddt (15.3, 11.6, 7.7) | 24.9 ($CH_2$) | | $δ_a$ | 3.23, q (5.4) | 40.9 ($CH_2$) |
| δ | 0.84, t (7.7) | 11.3 ($CH_3$) | | $δ_b$ | 3.19, dt (13.8, 7.1) | 40.9 ($CH_2$) |
| | | | | NHε | 7.09, t (5.9) | NH |
| Thr (-2) | | | | NHζ | - | - |
| Amide NH | 8.23, d (7.8) | NH | | NHζ | - | - |
| α | 4.29, overlapped | 56.6 (CH) | | | | |
| β | 4.03, quint (6.4) | 68.2 (CH) | | Ser 4 | | |
| γ | 1.13, d (6.4) | 19.7 ($CH_3$) | | Amide NH | 8.33, d (7.1) | NH |
| | | | | α | 4.28, overlapped | 59.9 (CH) |
| Ala (-1) | | | | β | 3.73, m | 62.1 ($CH_2$) |
| Amide NH | 8.16, overlapped | NH | | | | |
| α | 4.24, overlapped | 54.5 (CH) | | Lys 5 | | |
| β | 1.26, d (7.1) | 17.6 ($CH_3$) | | Amide NH | 8.16, overlapped | NH |
| | | | | α | 4.25, overlapped | (CH) |
| Trp 1 | | | | $β_a$ | 1.62, m | 31.3 ($CH_2$) |
| Amide NH | 8.07, d (6.7) | NH | | $β_b$ | 1.71, m | 31.3 ($CH_2$) |
| α | 4.23, overlapped | 50.3 (CH) | | γ | 1.30, m | 22.8 ($CH_2$) |
| $β_a$ | 3.06, m | 28.2 ($CH_2$) | | δ | 1.55, p (7.9) | 27.2 ($CH_2$) |
| $β_b$ | 3.82, m | 28.2 ($CH_2$) | | ε | 2.85, br s | 40.1 ($CH_2$) |
| 1 | 9.87, d (2.6) | NH | | NHε | 7.37, d (10.2) | NH |
| 2 | 7.2, m | 125.5 (CH) | | | | |
| 3 | - | * | | Ser 6 | | |
| 4 | 7.21, m | 118 (CH) | | Amide NH | 7.98, d (7.6) | NH |
| 5 | 6.74, d (9.1) | 125.8 (CH) | | α | 4.3, overlapped | overlapped |
| 6 | - | * | | β | 3.67, m | 62.2 ($CH_2$) |
| 7 | 7.28, s | 111.2 (CH) | | | | |
| | | | | Phe 7 | | |
| Asn 2 | | | | Amide NH | 7.7, d (7.6) | NH |
| Amide NH | 6.43, d (9.1) | NH | | α | 4.44, overlapped | overlapped |
| α | 4.15, overlapped | 50.8 (CH) | | $β_a$ | 2.94, d (7.7) | 37.9 ($CH_2$) |
| β | 2.06, dd (7.5, 5.4) | 40.8 ($CH_2$) | | $β_b$ | 3.08, dd (10.2, 4.6) | 37.9 ($CH_2$) |
| $NH_1$ | 6.5, br, s | NH | | 1 | - | * |
| $NH_2$ | 7.17, br, s | NH | | 2 | 7.16, m | 130.4 (CH) |
| | | | | 3 | 7.26, m | 129.7 (CH) |
| | | | | 4 | 7.21, m | 128.0 (CH) |

* No $^{13}C$ spectrum was recorded, thus, no $\delta_C$ was able to be assigned for the indicated carbon atoms.

**A**

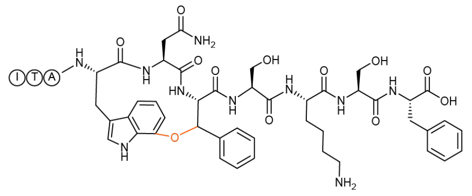

**B**

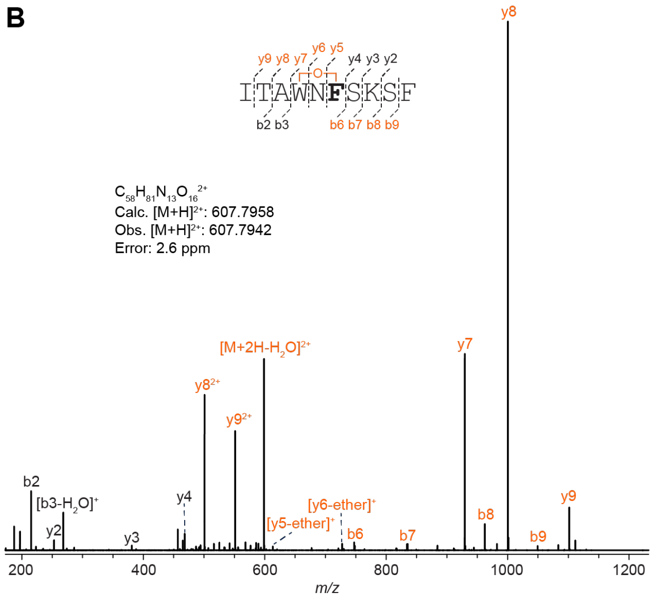

**C**

| Ion | Calculated Mass | Observed Mass | Error (ppm) |
|---|---|---|---|
| $[M+2H]^{2+}$ | 607.7958 | 607.7942 | 2.6 |
| $[M+2H-H_2O]^{2+}$ | 598.7904 | 598.7889 | 2.5 |
| $b9^+$ | 1049.5051 | 1049.5022 | 2.8 |
| $b9^{2+}$ | 525.2562 | 525.2548 | 2.7 |
| $y2^+$ | 253.1183 | 253.1177 | 2.4 |
| $b8^+$ | 962.4731 | 962.4706 | 2.6 |
| $y3^+$ | 381.2133 | 381.2124 | 2.4 |
| $b7^+$ | 834.3781 | 834.3761 | 2.4 |
| $y4^+$ | 468.2453 | 468.2441 | 2.6 |
| $b6^+$ | 747.3461 | 747.3443 | 2.4 |
| $[y5 - ether]^+$ | 613.2981 | 613.2965 | 2.6 |
| $[y6 - ether]^+$ | 727.3410 | 727.3392 | 2.5 |
| $b3^+$ | 286.1762 | 286.1755 | 2.4 |
| $[b3-H_2O]^+$ | 268.1656 | 268.165 | 2.2 |
| $y7^+$ | 929.4152 | 929.4124 | 3.0 |
| $y7^{2+}$ | 465.2113 | 465.2100 | 2.8 |
| $b2^+$ | 215.1391 | 215.1385 | 2.8 |
| $y8^+$ | 1000.4523 | 1000.4492 | 3.1 |
| $y8^{2+}$ | 500.7298 | 500.7286 | 2.4 |
| $y9^+$ | 1101.5000 | 1101.497 | 2.7 |
| $y9^{2+}$ | 551.2537 | 551.2523 | 2.5 |

**Figure S32: HR-MS/MS analysis of the DarA-W3F variant with an ether crosslink.** (A) Hypothetical structure of the modified species based on HR-MS/MS fragmentation. The connectivity of the linkage is presumed from the known structure of darobactin. The modification is highlighted in orange. (B) CID spectrum of the +14 Da modified variant. The sequence of the GluC-digested peptide is provided with the substitution bolded. Orange colored daughter ions include the modification. Distinguishing ether crosslink fragments are indicated. (C) Observed masses and errors from CID analysis.

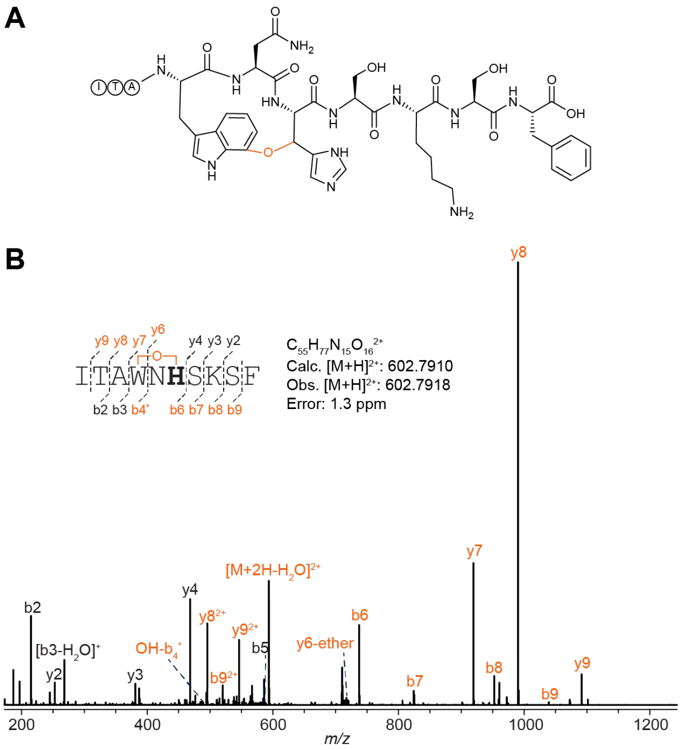

**C**

| Ion | Calculated Mass | Observed Mass | Error (ppm) |
|---|---|---|---|
| $[M+2H]^{2+}$ | 602.7910 | 602.7918 | 1.3 |
| $[M+2H-H_2O]^{2+}$ | 593.7857 | 593.7867 | 1.7 |
| $b9^+$ | 1039.4956 | 1039.4971 | 1.4 |
| $b9^{2+}$ | 520.2515 | 520.2523 | 1.5 |
| $y2^+$ | 253.1183 | 253.1187 | 1.6 |
| $b8^+$ | 952.4636 | 952.4648 | 1.3 |
| $b8^{2+}$ | 476.7355 | 476.7362 | 1.5 |
| $y3^+$ | 381.2133 | 381.2139 | 1.6 |
| $b7^+$ | 824.3686 | 824.3701 | 1.8 |
| $y4^+$ | 468.2453 | 468.246 | 1.5 |
| $b6^+$ | 737.3366 | 737.3377 | 1.5 |
| $OH\text{-}b4^*$ | 488.2504 | 488.2512 | 1.6 |
| $[y6\text{-ether}]^+$ | 717.3315 | 717.3326 | 1.5 |
| $b3^+$ | 286.1762 | 286.1766 | 1.4 |
| $[b3-H_2O]^+$ | 268.1656 | 268.1661 | 1.9 |
| $y7^+$ | 919.4057 | 919.4068 | 1.2 |
| $y7^{2+}$ | 460.2065 | 460.2072 | 1.5 |
| $b2^+$ | 215.1391 | 215.1394 | 1.4 |
| $y8^+$ | 990.4429 | 990.4439 | 1.0 |
| $y8^{2+}$ | 495.7251 | 495.7258 | 1.4 |
| $y9^+$ | 1091.4905 | 1091.4919 | 1.3 |
| $y9^{2+}$ | 546.2489 | 546.2498 | 1.6 |

**Figure S33: HR-MS/MS analysis of the DarA-W3H variant with an ether crosslink.** (A) Hypothetical structure of the modified species based on HR-MS/MS fragmentation. The connectivity of the linkage is presumed from the known structure of darobactin. The modification is highlighted in orange. (B) CID spectrum of the +14 Da species modified variant. The sequence of the GluC-digested peptide is provided with the substitution bolded. Orange colored daughter ions include the modification. Distinguishing ether crosslink fragments are indicated. (C) Observed masses and errors from CID analysis.

**A**

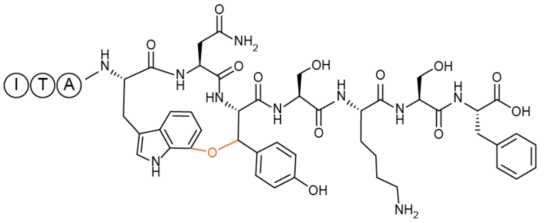

**B**

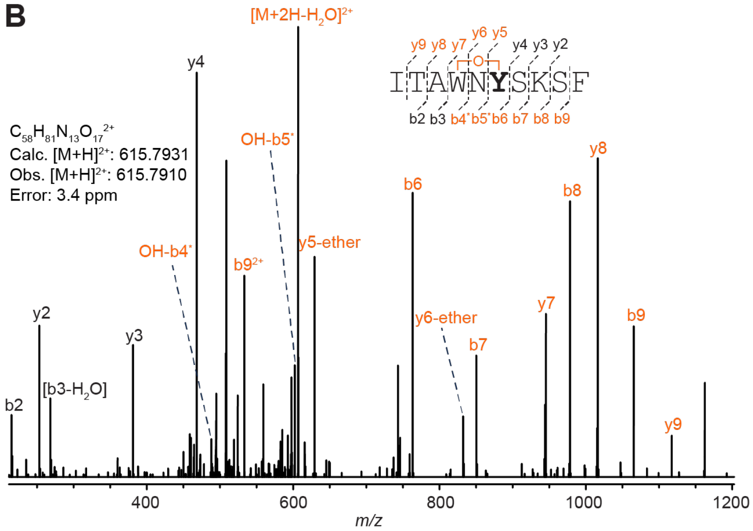

**C**

| Ion | Calculated Mass | Observed Mass | Error (ppm) |
|---|---|---|---|
| $[M+2H]^{2+}$ | 615.7931 | 615.791 | 3.4 |
| $[M+2H-H_2O]^{2+}$ | 606.7879 | 606.7853 | 4.3 |
| $b9^{+}$ | 1065.5000 | 1065.497 | 2.8 |
| $b9^{2+}$ | 533.2536 | 533.2522 | 2.6 |
| $y2^{+}$ | 253.1183 | 253.1176 | 2.8 |
| $b8^{+}$ | 978.4679 | 978.4648 | 3.2 |
| $y3^{+}$ | 381.2133 | 381.2122 | 2.9 |
| $b7^{+}$ | 850.3730 | 850.3706 | 2.8 |
| $y4^{+}$ | 468.2453 | 468.2439 | 3.0 |
| $b6^{+}$ | 763.3409 | 763.339 | 2.5 |
| $[y5 - ether]^{+}$ | 629.2930 | 629.2912 | 2.9 |
| $OH\text{-}b5^{*}$ | 602.2933 | 602.2918 | 2.5 |
| $[y6 - ether]^{+}$ | 743.3359 | 743.3341 | 2.4 |
| $OH\text{-}b4^{*}$ | 488.2504 | 488.2491 | 2.7 |
| $y7^{+}$ | 850.3730 | 850.3709 | 2.5 |
| $b3^{+}$ | 286.1762 | 286.1753 | 3.1 |
| $y8^{+}$ | 978.4680 | 978.4651 | 3.0 |
| $b2^{+}$ | 215.1391 | 215.1384 | 3.3 |
| $y9^{+}$ | 1117.4949 | 1117.4917 | 2.9 |

**Figure S34: HR-MS/MS analysis of the DarA-W3Y variant with an ether crosslink.** (A) Hypothetical structure of the modified species based on HR-MS/MS fragmentation. The connectivity of the linkage is presumed from the known structure of darobactin. The modification is highlighted in orange. (B) CID spectrum of the +14 Da modified variant. The sequence of the GluC-digested peptide is provided with the substitution bolded. Orange colored daughter ions include the modification. Distinguishing ether crosslink fragments are indicated. (C) Observed masses and errors from CID analysis.

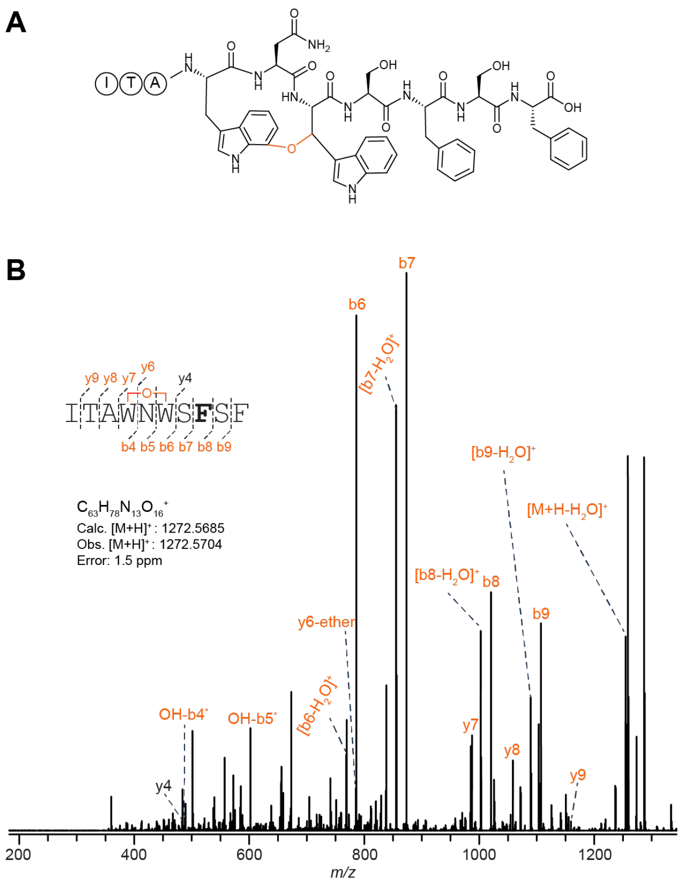

C

| Ion | Calculated Mass | Observed Mass | Error (ppm) |
|---|---|---|---|
| $[M+H]^+$ | 1272.5685 | 1272.5704 | 1.5 |
| $[M+H-H_2O]^+$ | 1254.5579 | 1254.5594 | 1.2 |
| $b9^+$ | 1107.4895 | 1107.4902 | 0.6 |
| $[b9-H_2O]^+$ | 1089.4788 | 1089.4794 | 0.6 |
| $b8^+$ | 1020.4575 | 1020.4582 | 0.7 |
| $[b8-H_2O]^+$ | 1002.4468 | 1002.4476 | 0.8 |
| $b7^+$ | 873.3891 | 873.3899 | 0.9 |
| $[b7-H_2O]^+$ | 855.3784 | 855.3793 | 1.1 |
| $y4^+$ | 487.2188 | 487.2191 | 0.6 |
| $b6^+$ | 786.3570 | 786.358 | 1.3 |
| $[b6-H_2O]^+$ | 768.3464 | 768.3475 | 1.4 |
| $OH-b5^*$ | 602.2933 | 602.294 | 1.2 |
| $OH-b4^*$ | 488.2504 | 488.2509 | 1.0 |
| $[y6 - ether]^+$ | 785.3254 | 785.3264 | 1.3 |
| $y7^+$ | 987.3996 | 987.4006 | 1.0 |
| $y8^+$ | 1058.4367 | 1058.4376 | 0.9 |
| $y9^+$ | 1159.4844 | 1159.4848 | 0.3 |

**Figure S35: HR-MS/MS analysis of the DarA-K5F variant with a single ether crosslink.** (A) Hypothetical structure of the modified species based on HR-MS/MS fragmentation. The connectivity of the linkage is presumed from the known structure of darobactin. The modification is highlighted in orange. (B) CID spectrum of the +14 Da modified variant. The sequence of the GluC-digested peptide is provided with the substitution bolded. Orange colored daughter ions include the modification. Distinguishing ether crosslink fragments are indicated. (C) Observed masses and errors from CID analysis.

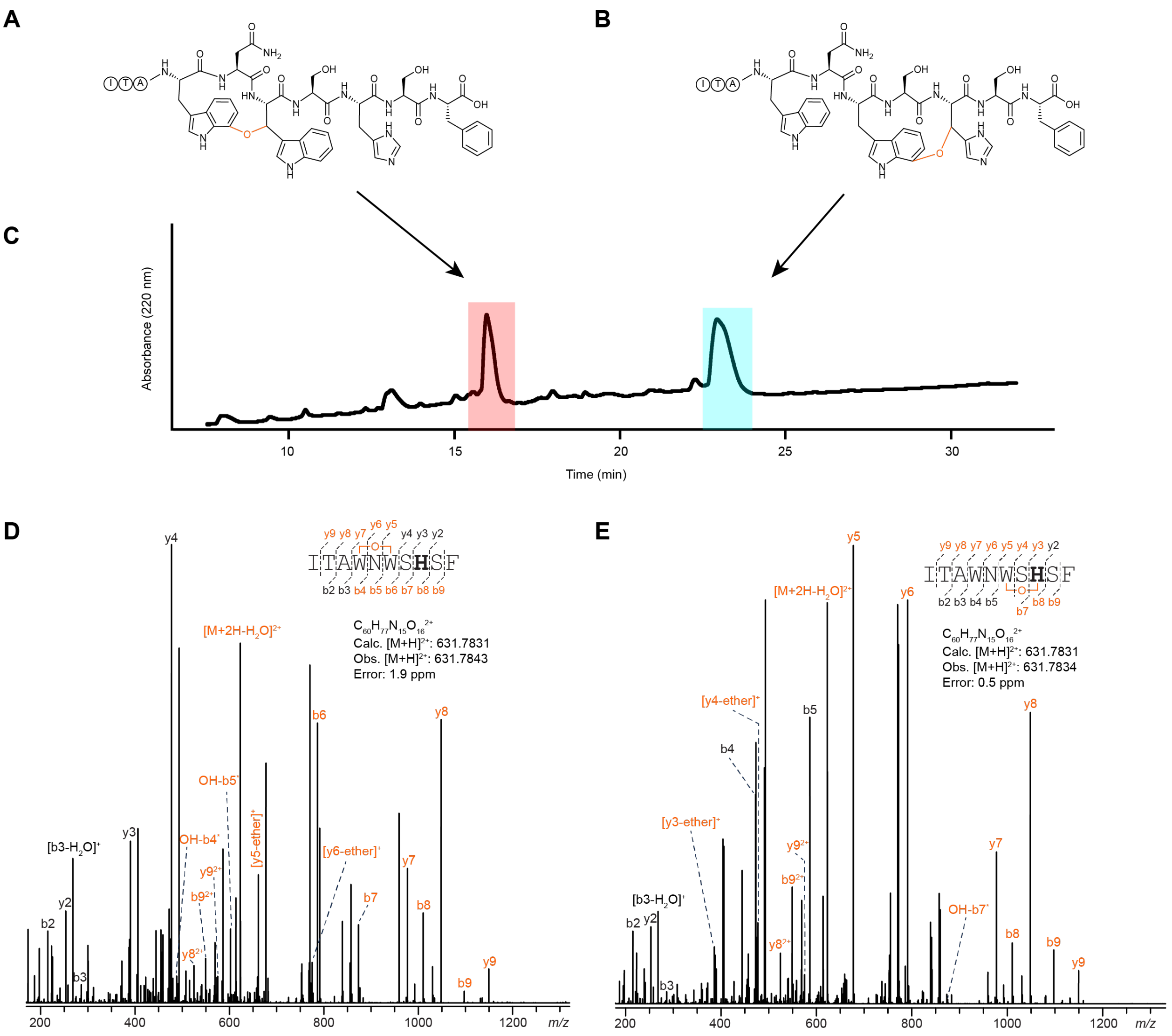

**F**

| Ion | Calculated Mass | Observed Mass | Error (ppm) |
|---|---|---|---|
| $[M+2H]^{2+}$ | 631.7831 | 631.7843 | 1.9 |
| $[M+2H-H_2O]^{2+}$ | 622.7779 | 622.7787 | 1.3 |
| $b9^{+}$ | 1097.4800 | 1097.4814 | 1.3 |
| $b9^{2+}$ | 549.2437 | 549.2444 | 1.3 |
| $y2^{+}$ | 253.1183 | 253.1187 | 1.6 |
| $b8^{+}$ | 1010.4480 | 1010.4491 | 1.1 |
| $y3^{+}$ | 390.1772 | 390.1779 | 1.8 |
| $b7^{+}$ | 873.3891 | 873.3902 | 1.3 |
| $y4^{+}$ | 477.2093 | 477.2100 | 1.5 |
| $b6^{+}$ | 786.3570 | 786.3583 | 1.7 |
| OH-$b5^{*}$ | 602.2933 | 602.2942 | 1.5 |
| $[y5\text{-ether}]^{+}$ | 661.2730 | 661.2738 | 1.2 |
| OH-$b4^{*}$ | 488.2504 | 488.2509 | 1.0 |
| $[y6\text{-ether}]^{+}$ | 775.3159 | 775.3170 | 1.4 |
| $b3^{+}$ | 286.1762 | 286.1767 | 1.7 |
| $[b3-H_2O]^{+}$ | 268.1656 | 268.1660 | 1.5 |
| $y7^{+}$ | 977.3901 | 977.3912 | 1.1 |
| $b2^{+}$ | 215.1391 | 215.1393 | 0.9 |
| $y8^{+}$ | 1048.4272 | 1048.4284 | 1.1 |
| $y8^{2+}$ | 524.7173 | 524.7178 | 1.0 |
| $y9^{+}$ | 1149.4749 | 1149.4760 | 1.0 |
| $y9^{2+}$ | 575.2411 | 575.2420 | 1.6 |

**G**

| Ion | Calculated Mass | Observed Mass | Error (ppm) |
|---|---|---|---|
| $[M+2H]^{2+}$ | 631.7831 | 631.7834 | 0.5 |
| $[M+2H-H_2O]^{2+}$ | 622.7779 | 622.7784 | 0.8 |
| $b9^{+}$ | 1097.4800 | 1097.4806 | 0.5 |
| $b9^{2+}$ | 549.2437 | 549.2443 | 1.1 |
| $y2^{+}$ | 253.1183 | 253.1186 | 1.2 |
| $b8^{+}$ | 1010.4480 | 1010.4485 | 0.5 |
| $[y3\text{-ether}]^{+}$ | 388.1616 | 388.1620 | 1.0 |
| OH-$b7^{*}$ | 875.4047 | 875.4053 | 0.7 |
| $[y4\text{-ether}]^{+}$ | 475.1937 | 475.1941 | 0.8 |
| $b5^{+}$ | 586.2984 | 586.2990 | 1.0 |
| $y5^{+}$ | 677.2679 | 677.2684 | 0.7 |
| $b4^{+}$ | 472.2555 | 472.2559 | 0.8 |
| $y6^{+}$ | 791.3108 | 791.3115 | 0.9 |
| $b3^{+}$ | 286.1762 | 286.1765 | 1.0 |
| $[b3-H_2O]^{+}$ | 268.1656 | 268.1659 | 1.1 |
| $y7^{+}$ | 977.3901 | 977.3908 | 0.7 |
| $b2^{+}$ | 215.1391 | 215.1393 | 0.9 |
| $y8^{+}$ | 1048.4272 | 1048.4279 | 0.7 |
| $y8^{2+}$ | 524.7173 | 524.7178 | 1.0 |
| $y9^{+}$ | 1149.4749 | 1149.4754 | 0.4 |
| $y9^{2+}$ | 575.2411 | 575.2417 | 1.0 |

**Figure S36 (previous page): HR-MS/MS analysis of the DarA-K5H variant with single ether crosslinks.** (A) and (B) Hypothetical structures of variant DarA-K5H with distinct monoether crosslink positions. The connectivity of the linkages are presumed from the known structure of darobactin. The modifications are highlighted in orange. (C) UHPLC chromatogram of the GluC-digested variant while monitoring the 220 nm absorbance. The light red and cyan fractions eluted at 16 min and 23 min, respectively, and both contain the +14 Da modified variant. (D) CID spectrum of the +14 Da species modified variant isolated from the light red and (E) cyan fractions as in panel C. The sequence of the GluC-digested peptide is provided with the substitution bolded. Orange colored daughter ions include the modification. Distinguishing ether crosslink fragments are indicated. (F) and (G) Observed masses and errors from CID analysis for panels D and E, respectively.

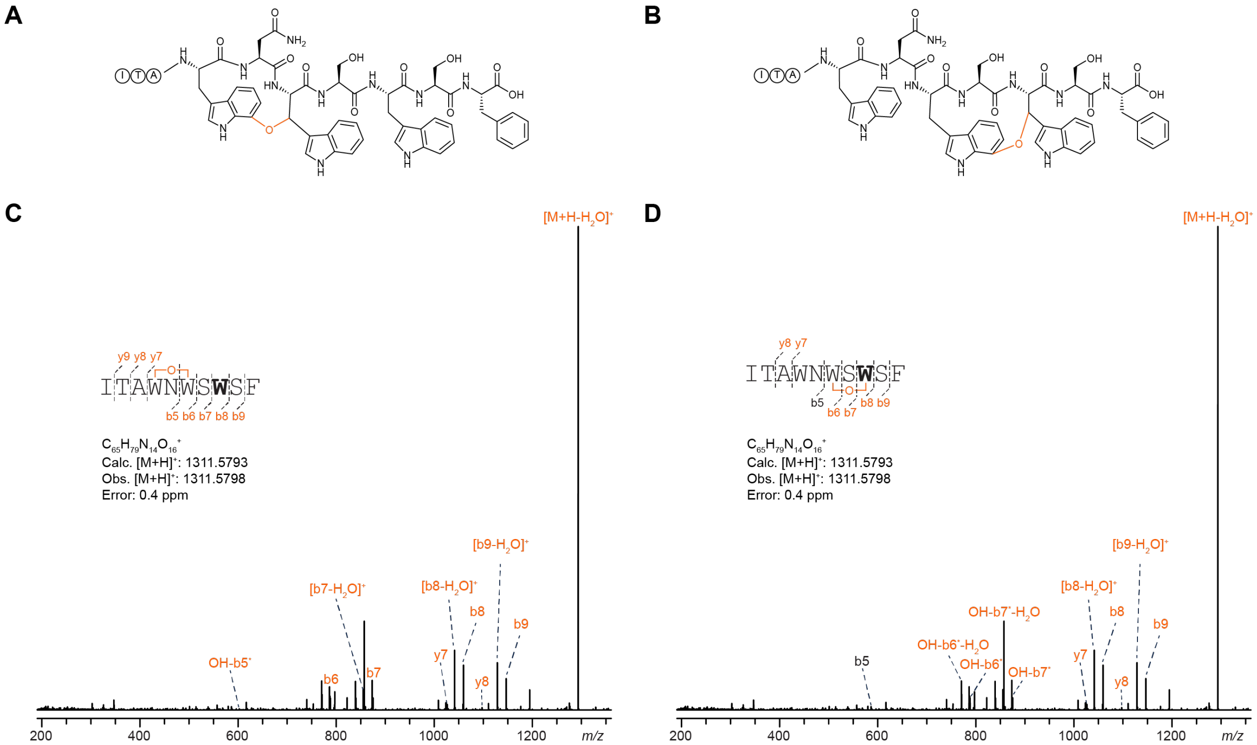

**E**

| Ion | Calculated Mass | Observed Mass | Error (ppm) |
|---|---|---|---|
| $[M+H]^+$ | 1311.5793 | 1311.5798 | 0.4 |
| $[M+H-H_2O]^+$ | 1293.5688 | 1293.5707 | 1.5 |
| $b9^+$ | 1146.5004 | 1146.4995 | 0.8 |
| $[b9-H_2O]^+$ | 1128.4897 | 1128.4904 | 0.6 |
| $b8^+$ | 1059.4683 | 1059.4682 | 0.1 |
| $[b8-H_2O]^+$ | 1041.4577 | 1041.4580 | 0.3 |
| $b7^+$ | 873.3890 | 873.3900 | 1.1 |
| $[b7-H_2O]^+$ | 855.3791 | 855.3794 | 0.4 |
| $b6^+$ | 786.3570 | 786.3583 | 1.7 |
| OH-b5* | 602.2933 | 602.2945 | 2.0 |
| $y7^+$ | 1026.4105 | 1026.4078 | 2.6 |
| $y8^+$ | 1097.4476 | 1097.4477 | 0.1 |

**F**

| Ion | Calculated Mass | Observed Mass | Error (ppm) |
|---|---|---|---|
| $[M+H]^+$ | 1311.5793 | 1311.5798 | 0.4 |
| $[M+H-H_2O]^+$ | 1293.5688 | 1293.5707 | 1.5 |
| $b9^+$ | 1146.5004 | 1146.4995 | 0.8 |
| $[b9-H_2O]^+$ | 1128.4897 | 1128.4904 | 0.6 |
| $b8^+$ | 1059.4683 | 1059.4682 | 0.1 |
| $[b8-H_2O]^+$ | 1041.4577 | 1041.4580 | 0.3 |
| $b7^+$ | 873.3890 | 873.3900 | 1.1 |
| $[b7-H_2O]^+$ | 855.3791 | 855.3794 | 0.4 |
| $b6^+$ | 786.3570 | 786.3583 | 1.7 |
| OH-b5* | 602.2933 | 602.2945 | 2.0 |
| $y7^+$ | 1026.4105 | 1026.4078 | 2.6 |
| $y8^+$ | 1097.4476 | 1097.4477 | 0.1 |

**Figure S37: HR-MS/MS analysis of the DarA-K5W variant with single ether crosslinks.** (A) and (B) Hypothetical structures of variant DarA-K5W with distinct monoether crosslink positions. The connectivity of the linkages are presumed from the known structure of darobactin. The modifications are highlighted in orange. CID spectrum of the +14 Da modified variant with a (C) Trp1-Trp3 or (D) Trp3-Trp5 ether crosslink. The sequence of the GluC-digested peptide is provided with the substitution bolded. Orange colored daughter ions include the modification. Distinguishing ether crosslink fragments are indicated. (E) and (F) Observed masses and errors from CID analysis for panels C and D, respectively.

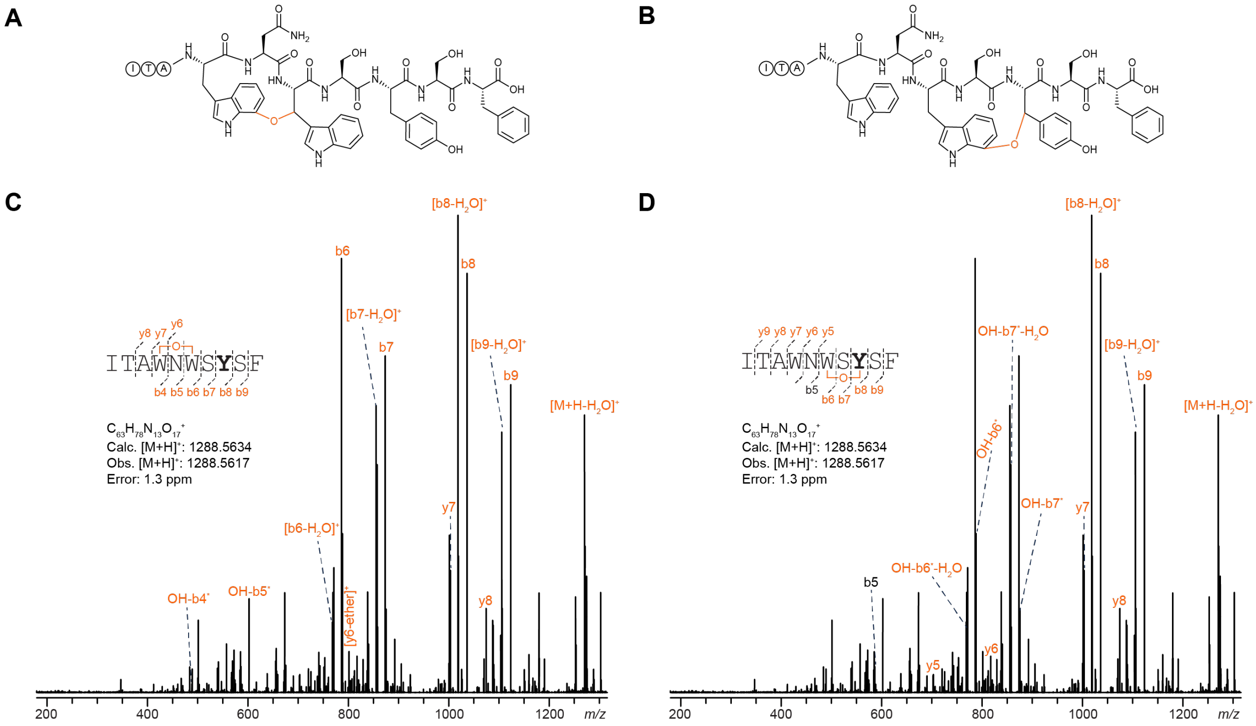

E

| Ion | Calculated Mass | Observed Mass | Error (ppm) |
|---|---|---|---|
| $[M+H]^+$ | 1288.5634 | 1288.5617 | 1.3 |
| $[M+H-H_2O]^+$ | 1270.5528 | 1270.5501 | 2.1 |
| $b9^+$ | 1123.4844 | 1123.4816 | 2.5 |
| $[b9-H_2O]^+$ | 1105.4738 | 1105.4710 | 2.5 |
| $b8^+$ | 1036.4514 | 1036.4496 | 1.7 |
| $[b8-H_2O]^+$ | 1018.4418 | 1018.4394 | 2.4 |
| $b7^+$ | 873.3890 | 873.3870 | 2.3 |
| $[b7-H_2O]^+$ | 855.3784 | 855.3763 | 2.5 |
| $b6^+$ | 786.3570 | 786.3553 | 2.2 |
| $[b6-H_2O]^+$ | 768.3464 | 768.3448 | 2.1 |
| $OH-b5^*$ | 602.2933 | 602.2918 | 2.5 |
| $OH-b4^*$ | 488.2504 | 488.2492 | 2.5 |
| $[y6 - ether]^+$ | 801.3203 | 801.3187 | 2.0 |
| $y7^+$ | 1003.3945 | 1003.3920 | 2.5 |
| $y8^+$ | 1074.4316 | 1074.4291 | 2.3 |

F

| Ion | Calculated Mass | Observed Mass | Error (ppm) |
|---|---|---|---|
| $[M+H]^+$ | 1288.5634 | 1288.5617 | 1.3 |
| $[M+H-H_2O]^+$ | 1270.5528 | 1270.5501 | 2.1 |
| $b9^+$ | 1123.4844 | 1123.4816 | 2.5 |
| $[b9-H_2O]^+$ | 1105.4738 | 1105.4710 | 2.5 |
| $b8^+$ | 1036.4524 | 1036.4496 | 2.7 |
| $[b8-H_2O]^+$ | 1018.4417 | 1018.4394 | 2.3 |
| $OH-b7^*$ | 875.4047 | 875.4024 | 2.6 |
| $[OH-b7^* - H_2O]$ | 857.3943 | 857.3919 | 2.8 |
| $OH-b6^*$ | 788.3726 | 788.3710 | 2.0 |
| $[OH-b6^* - H_2O]$ | 770.3620 | 770.3604 | 2.1 |
| $b5^+$ | 586.2984 | 586.2970 | 2.4 |
| $y5^+$ | 703.2723 | 703.2704 | 2.7 |
| $y6^+$ | 817.3152 | 817.3134 | 2.2 |
| $y7^+$ | 1003.3945 | 1003.3920 | 2.5 |
| $y8^+$ | 1074.4316 | 1074.4291 | 2.3 |

**Figure S38: HR-MS/MS analysis of the DarA-K5Y variant with a single ether crosslink.** (A) and (B) Hypothetical structures of the modified DarA-K5Y variant with distinct monoether crosslink positions. The connectivity of the linkage is presumed from the known structure of darobactin. The modification is highlighted in orange. CID spectrum of the +14 Da modified variant with a (C) Trp1-Trp3 or (D) Trp3-Tyr5 ether crosslink. The sequence of the GluC-digested peptide is provided with the substitution bolded. Orange colored daughter ions include the modification. Distinguishing ether crosslink fragments are indicated. (E) and (F) Observed masses and errors from CID analysis for panels C and D, respectively.

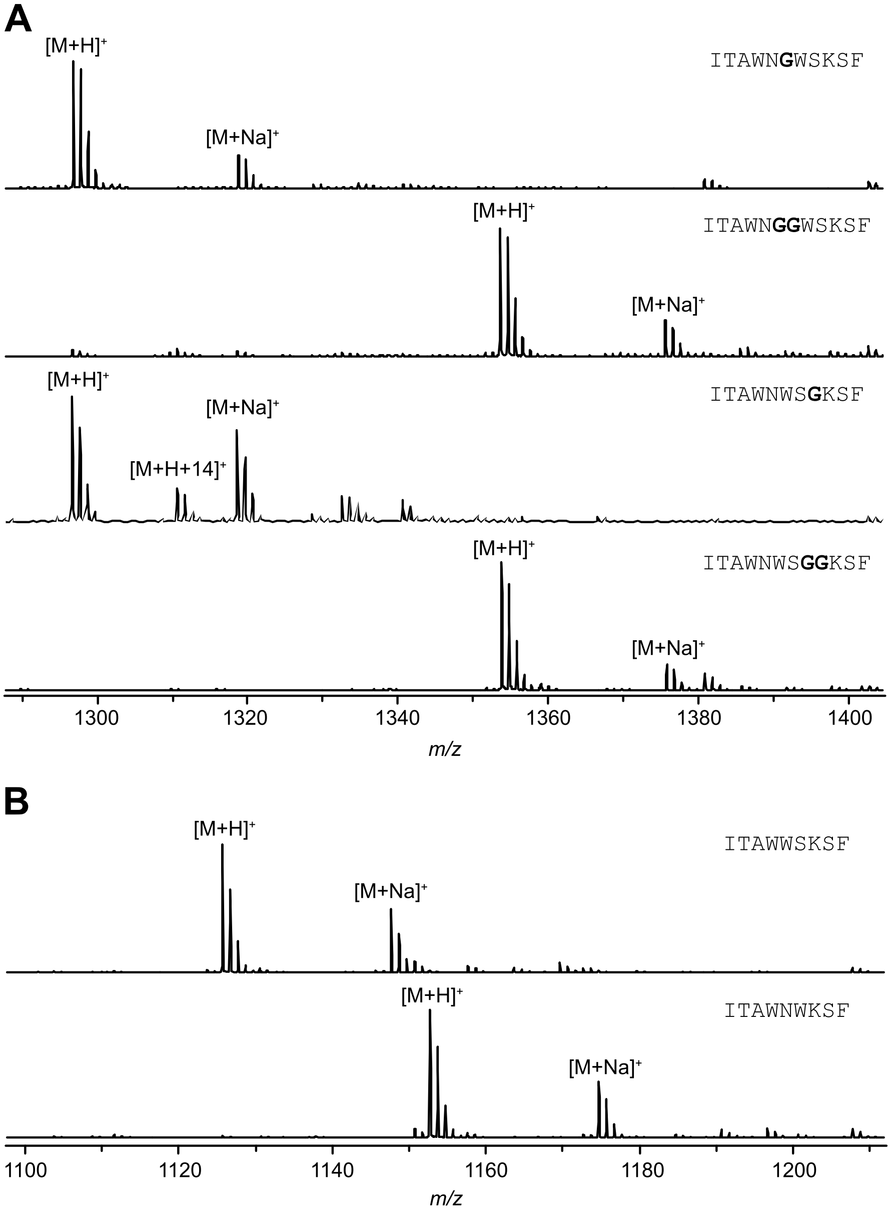

**Figure S39: MALDI-TOF mass spectra of darobactin ring expansions and contractions.** (A) GluC-digest of darobactin variants with Gly residues inserted within the ring-forming motifs. (B) GluC-digest of darobactin variants with residues removed from the ring-forming motifs. The analyzed peptide sequence is shown next to each spectrum with the added residues bolded.

**A**

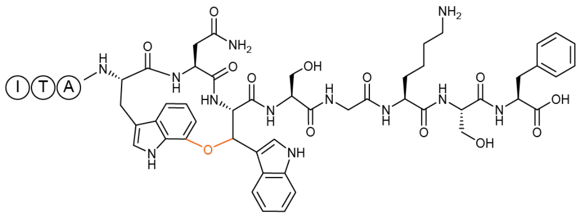

**B**

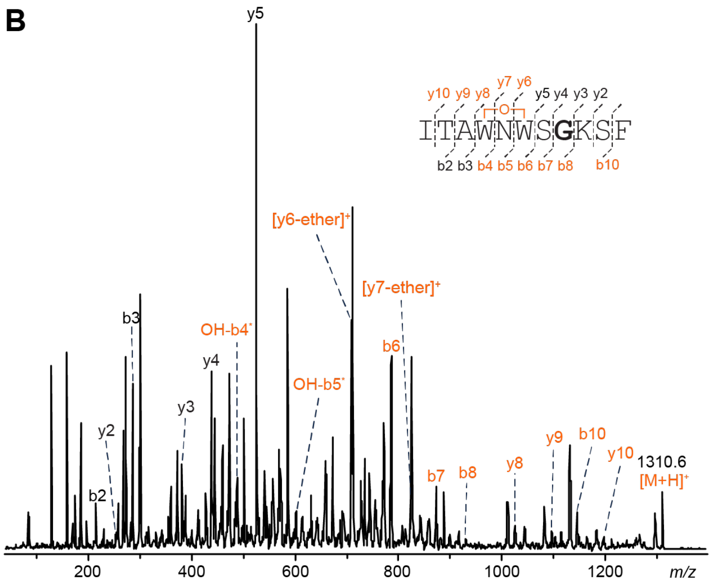

**C**

| Ion | Calculated Mass | Observed Mass |
|---|---|---|
| $[M+H]^+$ | 1310.6 | 1310.6 |
| $b10^+$ | 1145.5 | 1145.5 |
| $y2^+$ | 253.1 | 253.1 |
| $b8^+$ | 930.4 | 930.3 |
| $y3^+$ | 381.2 | 381.2 |
| $b7^+$ | 873.4 | 873.2 |
| $y4^+$ | 438.2 | 438.2 |
| $b6^+$ | 786.4 | 786.2 |
| OH-b5* | 602.3 | 602.1 |
| $y5^+$ | 525.3 | 525.1 |
| OH-b4* | 488.3 | 488.1 |
| $[y6\text{-ether}]^+$ | 709.3 | 709.1 |
| $b3^+$ | 286.2 | 286.1 |
| $[y7\text{-ether}]^+$ | 823.4 | 823.1 |
| $b2^+$ | 215.1 | 215.1 |
| $y8^+$ | 1025.4 | 1025.4 |
| $y9^+$ | 1096.5 | 1096.4 |
| $y10^+$ | 1197.5 | 1197.4 |

**Figure S40: MALDI-LIFT-TOF/TOF-MS analysis of the WNWSGSF darobactin variant.** (A) Hypothetical structure of the modified species based on HR-MS/MS fragmentation. The connectivity of the linkage is presumed from the known structure of darobactin. The modification is highlighted in orange. (B) Laser-induced dissociation (LID) spectrum of the +14 Da modified variant. The sequence of the GluC-digested peptide is provided with the substitution bolded. Orange colored daughter ions include the modification. Distinguishing ether crosslink fragments are indicated.

**A**

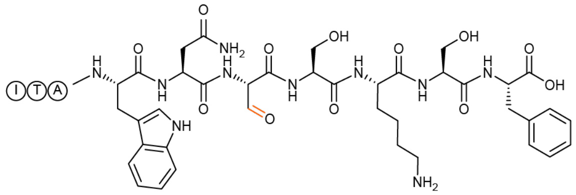

**B**

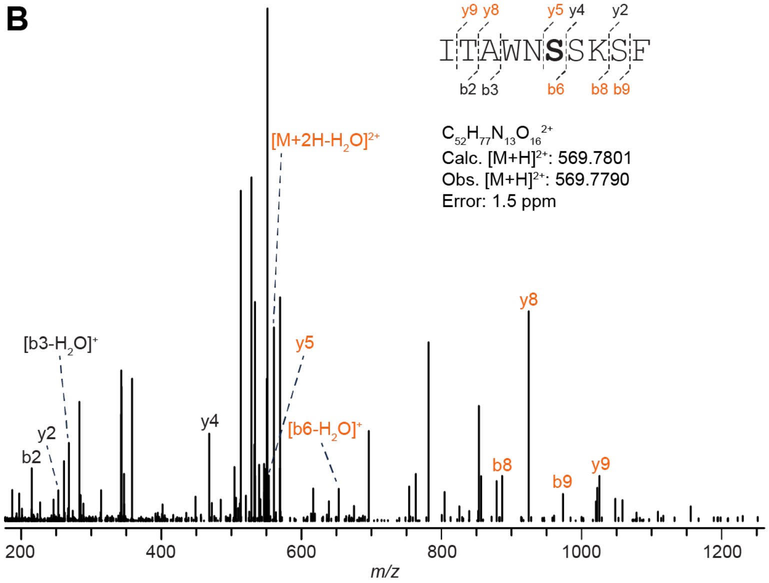

**C**

| Ion | Calculated Mass | Observed Mass | Error (ppm) |
|---|---|---|---|
| $[M+2H]^{2+}$ | 569.7801 | 569.7790 | 1.9 |
| $[M+2H-H_2O]^{2+}$ | 560.7748 | 560.7740 | 1.4 |
| $b9^+$ | 973.4738 | 973.4721 | 1.7 |
| $y2^+$ | 253.1183 | 253.1180 | 1.2 |
| $b8^+$ | 886.4418 | 886.4408 | 1.1 |
| $y4^+$ | 468.2453 | 468.2448 | 1.1 |
| $[b6-H_2O]^+$ | 653.3042 | 653.3030 | 1.8 |
| $y5^+$ | 553.2617 | 553.2625 | 1.4 |
| $b3^+$ | 286.1762 | 286.1757 | 1.7 |
| $[b3-H_2O]^+$ | 268.1656 | 268.1653 | 1.1 |
| $y7^+$ | 853.3839 | 853.3813 | 3.0 |
| $b2^+$ | 215.1391 | 215.1387 | 1.9 |
| $y8^+$ | 924.4211 | 924.4195 | 1.7 |
| $y9^+$ | 1025.4687 | 1025.4662 | 2.4 |

**Figure S41: HR-MS/MS of the DarA-W3S variant with a formylglycine modification.** (A) Hypothetical structure of the modified species based on HR-MS/MS fragmentation. The modification is highlighted in orange. (B) CID spectrum of the -2 Da modified variant. The sequence of the GluC-digested peptide is provided with the substitution bolded. Orange colored daughter ions include the modification. (C) Observed masses and errors from CID analysis.

A

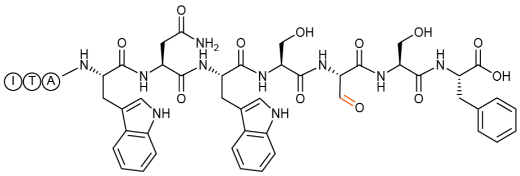

B

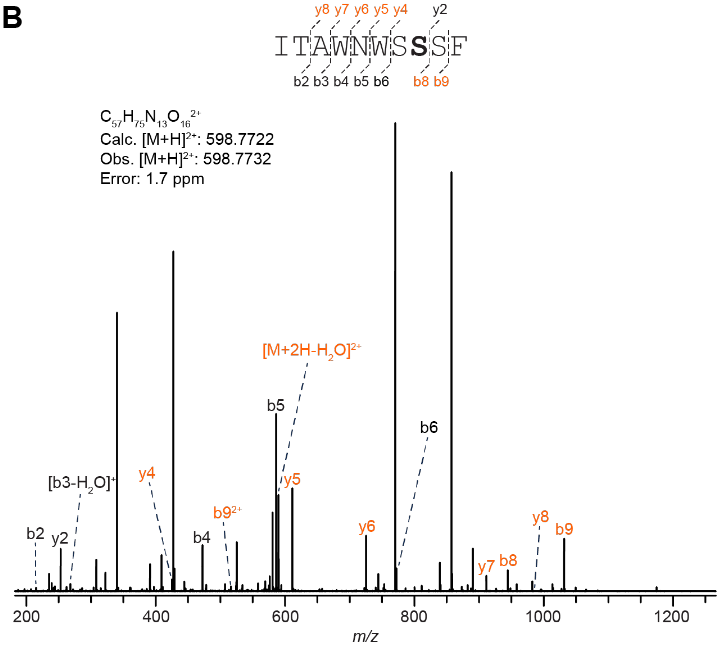

C

| Ion | Calculated Mass | Observed Mass | Error (ppm) |
|---|---|---|---|
| $[M+2H]^{2+}$ | 598.7722 | 598.7732 | 1.7 |
| $[M+2H-H_2O]^{2+}$ | 589.7670 | 589.7678 | 1.4 |
| $b9^+$ | 1031.4582 | 1031.4595 | 1.3 |
| $b9^{2+}$ | 516.2327 | 516.2334 | 1.4 |
| $y2^+$ | 253.1183 | 253.1187 | 1.6 |
| $b8^+$ | 944.4261 | 944.4274 | 1.4 |
| $y4^+$ | 425.1667 | 425.1674 | 1.6 |
| $b6^+$ | 772.3777 | 772.3789 | 1.6 |
| $b5^+$ | 586.2984 | 586.2994 | 1.7 |
| $y5^+$ | 611.2461 | 611.2469 | 1.3 |
| $b4^+$ | 472.2555 | 472.2562 | 1.5 |
| $y6^+$ | 725.2890 | 725.2901 | 1.5 |
| $b3^+$ | 286.1762 | 286.1767 | 1.7 |
| $[b3-H_2O]^+$ | 268.1656 | 268.1660 | 1.5 |
| $y7^+$ | 911.3683 | 911.3694 | 1.2 |
| $b2^+$ | 215.1391 | 215.1394 | 1.4 |
| $y8^+$ | 982.4054 | 982.4069 | 1.5 |

**Figure S42: HR-MS/MS of the DarA-K5S variant with a formylglycine modification.** (A) Hypothetical structure of the modified species based on HR-MS/MS fragmentation. The modification is highlighted in orange. (B) CID spectrum of the -2 Da modified variant. The sequence of the GluC-digested peptide is provided with the substitution bolded. Orange colored daughter ions include the modification. (C) Observed masses and errors from CID analysis.

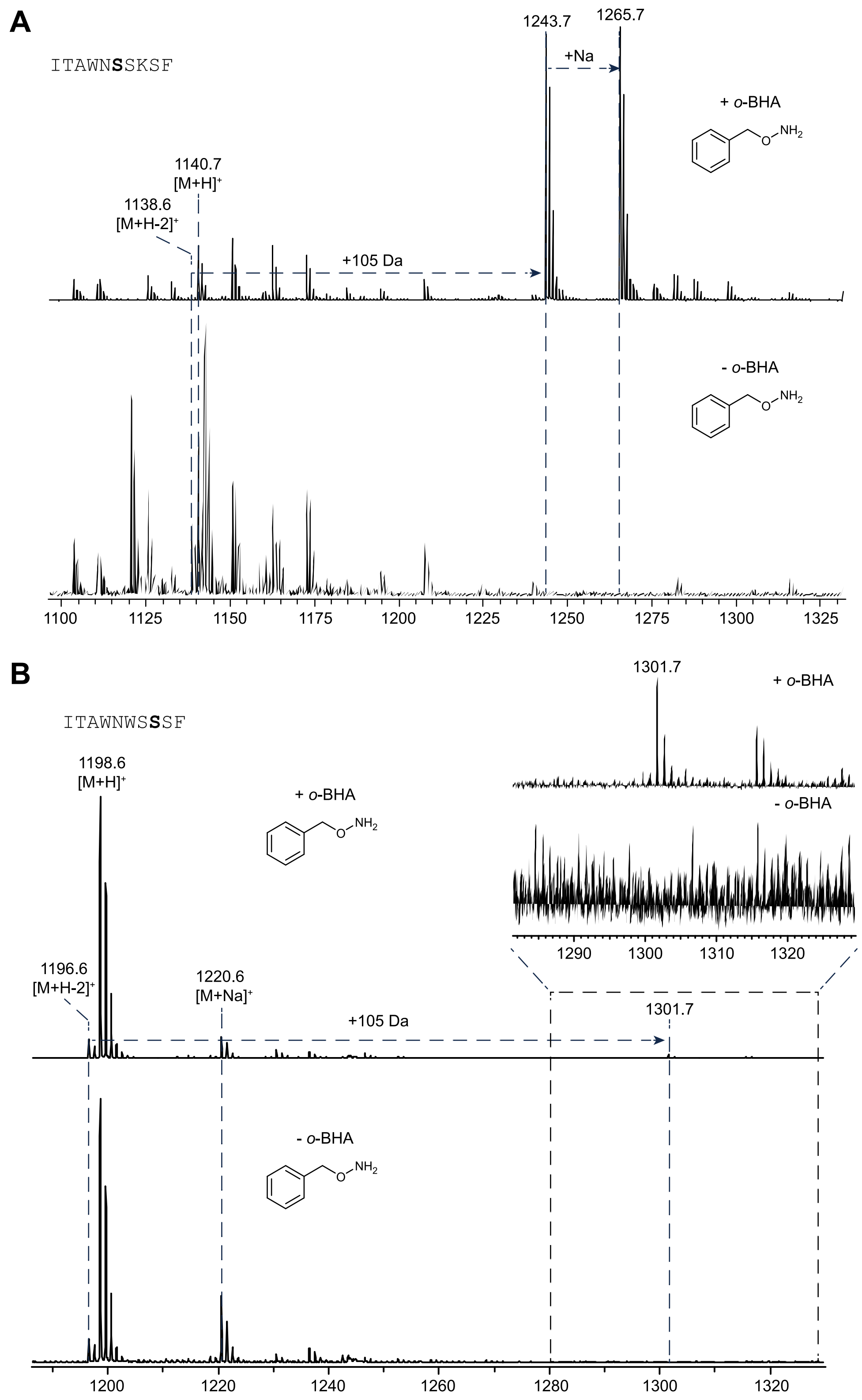

**Figure S43: Formylglycine oxime labeling reactions.** MALDI-TOF mass spectra of the GluC-digested (A) DarA-W3S and (B) the DarA-K5S variants with *o*-BHA.

**A**

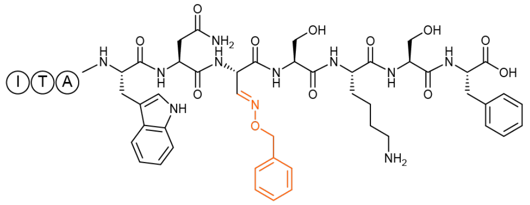

**B**

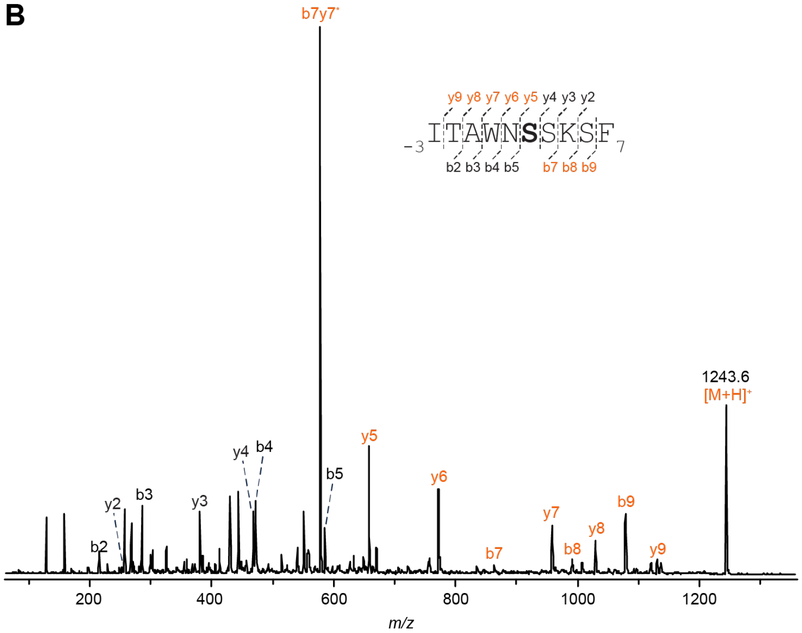

**C**

| Ion | Calculated Mass | Observed Mass |
|---|---|---|
| $[M+H]^+$ | 1243.6 | 1243.6 |
| $b9^+$ | 1078.5 | 1078.5 |
| $y2^+$ | 253.1 | 253.1 |
| $b8^+$ | 991.5 | 991.5 |
| $y3^+$ | 381.2 | 381.2 |
| $b7^+$ | 863.4 | 863.5 |
| $y4^+$ | 468.2 | 468.2 |
| $b5^+$ | 586.3 | 586.3 |
| $b7y7^+$ (internal) | 578.2 | 578.2 |
| $y5^+$ | 658.3 | 658.3 |
| $b4^+$ | 472.3 | 472.2 |
| $y6^+$ | 772.3 | 772.3 |
| $b3^+$ | 286.2 | 286.1 |
| $y7^+$ | 958.4 | 958.4 |
| $b2^+$ | 215.1 | 215.1 |
| $y8^+$ | 1029.4 | 1029.5 |
| $y9^+$ | 1130.5 | 1130.6 |

**Figure S44: MALDI-LIFT-TOF/TOF-MS analysis of the oxime labeled DarA-W3S variant.** (A) Hypothetical structure of the modified species based on MALDI-LIFT-TOF/TOF-MS fragmentation. The modification is highlighted in orange. (B) LID spectrum of the oxime labeled DarA-W3S variant. The sequence of the GluC-digested peptide is provided with the substitution bolded. Orange colored daughter ions include the modification. (C) Table of observed daughter ions.

A

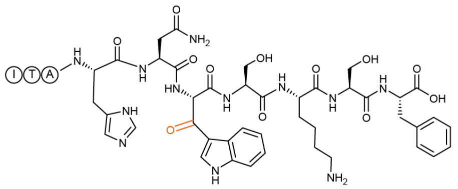

B

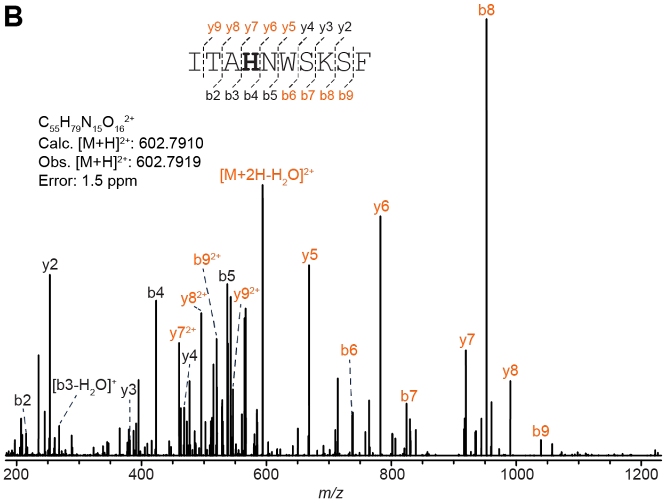

C

| Ion | Calculated Mass | Observed Mass | Error (ppm) |
|---|---|---|---|
| $[M+2H]^{2+}$ | 602.7910 | 602.7919 | 1.5 |
| $[M+2H-H_2O]^{2+}$ | 593.7857 | 593.7866 | 1.5 |
| $b9^+$ | 1039.4956 | 1039.4974 | 1.7 |
| $b9^{2+}$ | 520.2515 | 520.2522 | 1.3 |
| $y2^+$ | 253.1183 | 253.1187 | 1.6 |
| $b8^+$ | 952.4636 | 952.4648 | 1.3 |
| $b8^{2+}$ | 476.7355 | 476.7362 | 1.5 |
| $y3^+$ | 381.2133 | 381.2138 | 1.3 |
| $b7^+$ | 824.3686 | 824.3697 | 1.3 |
| $y4^+$ | 468.2453 | 468.2459 | 1.3 |
| $b6^+$ | 737.3366 | 737.3379 | 1.8 |
| $b5^+$ | 537.2780 | 537.2788 | 1.5 |
| $y5^+$ | 668.3039 | 668.3049 | 1.5 |
| $b4^+$ | 423.2351 | 423.2358 | 1.7 |
| $y6^+$ | 782.3468 | 782.3481 | 1.7 |
| $[b3-H_2O]^+$ | 268.1656 | 268.1660 | 1.5 |
| $y7^+$ | 919.4057 | 919.4068 | 1.2 |
| $y7^{2+}$ | 460.2065 | 460.2072 | 1.5 |
| $b2^+$ | 215.1391 | 215.1394 | 1.4 |
| $y8^+$ | 990.4429 | 990.4444 | 1.5 |
| $y8^{2+}$ | 495.7251 | 495.7258 | 1.4 |
| $y9^{2+}$ | 546.2489 | 546.2498 | 1.6 |

**Figure S45: HR-MS/MS of the DarA-W1H variant with a β-oxotryptophan modification.** (A) Hypothetical structure of the modified species based on HR-MS/MS fragmentation. The connectivity of the linkage is presumed from the known structure of darobactin. The modification is highlighted in orange. (B) CID spectrum of the +14 Da modified variant. The sequence of the GluC-digested peptide is provided with the substitution bolded. Orange colored daughter ions include the modification. (C) Observed masses and errors from CID analysis.

A

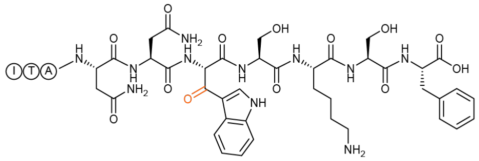

B

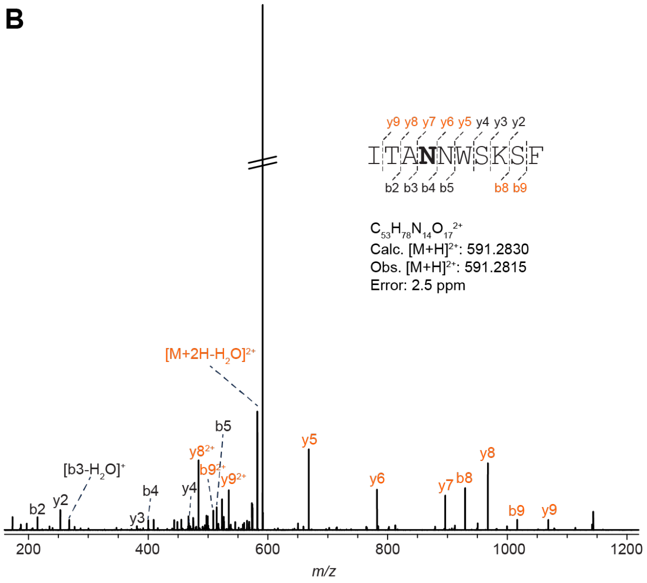

C

| Ion | Calculated Mass | Observed Mass | Error (ppm) |
|---|---|---|---|
| $[M+2H]^{2+}$ | 591.283 | 591.2815 | 2.5 |
| $[M+2H-H_2O]^{2+}$ | 582.2777 | 582.276 | 2.9 |
| $b9^+$ | 1016.4797 | 1016.4767 | 3.0 |
| $b9^{2+}$ | 508.7435 | 508.7419 | 3.1 |
| $y2^+$ | 253.1183 | 253.1176 | 2.8 |
| $b8^+$ | 929.4476 | 929.445 | 2.8 |
| $y3^+$ | 381.2133 | 381.2122 | 2.9 |
| $y4^+$ | 468.2453 | 468.2439 | 3.0 |
| $b5^+$ | 514.2620 | 514.2605 | 2.9 |
| $y5^+$ | 668.3039 | 668.3019 | 3.0 |
| $b4^+$ | 400.2191 | 400.2179 | 3.0 |
| $y6^+$ | 782.3469 | 782.3448 | 2.7 |
| $[b3-H_2O]^+$ | 268.1656 | 268.1649 | 2.6 |
| $y7^+$ | 896.3898 | 896.3873 | 2.8 |
| $y7^{2+}$ | 448.6986 | 448.6972 | 3.1 |
| $b2^+$ | 215.1391 | 215.1384 | 3.3 |
| $y8^+$ | 967.4269 | 967.424 | 3.0 |
| $y8^{2+}$ | 484.2171 | 484.2157 | 2.9 |
| $y9^+$ | 1068.4746 | 1068.4714 | 3.0 |
| $y9^{2+}$ | 534.7410 | 534.7394 | 3.0 |

**Figure S46: HR-MS/MS of the DarA-W1N variant with a β-oxotryptophan modification.** (A) Hypothetical structure of the modified species based on HR-MS/MS fragmentation. The connectivity of the linkage is presumed from the known structure of darobactin. The modification is highlighted in orange. (B) CID spectrum of the +14 Da modified variant. The sequence of the GluC-digested peptide is provided with the substitution bolded. Orange colored daughter ions include the modification. The intense ion at 591.3852 is an unrelated, singly-charged contaminant. (C) Observed masses and errors from CID analysis.

**A**

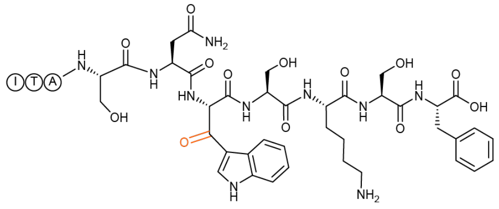

**B**

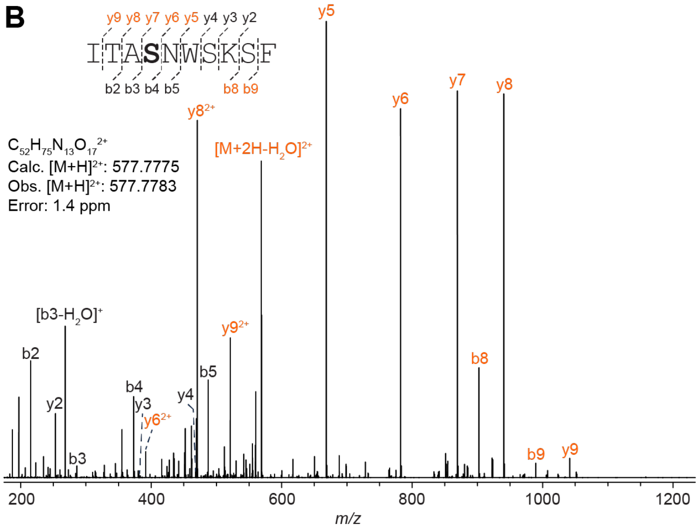

**C**

| Ion | Calculated Mass | Observed Mass | Error (ppm) |
|---|---|---|---|
| $[M+2H]^{2+}$ | 577.7775 | 577.7783 | 1.4 |
| $[M+2H-H_2O]^{2+}$ | 568.7722 | 568.7728 | 1.1 |
| $b9^+$ | 989.4687 | 989.4697 | 1.0 |
| $y2^+$ | 253.1183 | 253.1186 | 1.2 |
| $b8^+$ | 902.4367 | 902.4374 | 0.8 |
| $y3^+$ | 381.2133 | 381.2136 | 0.8 |
| $y4^+$ | 468.2453 | 468.2457 | 0.9 |
| $b5^+$ | 487.2511 | 487.2516 | 1.0 |
| $y5^+$ | 668.3039 | 668.3046 | 1.0 |
| $b4^+$ | 373.2082 | 373.2087 | 1.3 |
| $y6^+$ | 782.3468 | 782.3476 | 1.0 |
| $b3^+$ | 286.1762 | 286.1765 | 1.0 |
| $[b3- H_2O]^+$ | 268.1656 | 268.1659 | 1.1 |
| $y7^+$ | 869.3789 | 869.3795 | 0.7 |
| $b2^+$ | 215.1391 | 215.1393 | 0.9 |
| $y8^+$ | 940.4160 | 940.4168 | 0.9 |
| $y8^{2+}$ | 470.7116 | 470.7122 | 1.3 |
| $y9^+$ | 1041.4634 | 1041.4642 | 0.8 |
| $y9^{2+}$ | 521.2355 | 521.2361 | 1.2 |

**Figure S47: HR-MS/MS of the DarA-W1S variant with a β-oxotryptophan modification.** (A) Hypothetical structure of the modified species based on HR-MS/MS fragmentation. The modification is highlighted in orange. (B) CID spectrum of the +14 Da modified variant. The sequence of the GluC-digested peptide is provided with the substitution bolded. Orange colored daughter ions include the modification. (C) Observed masses and errors from CID analysis.

A

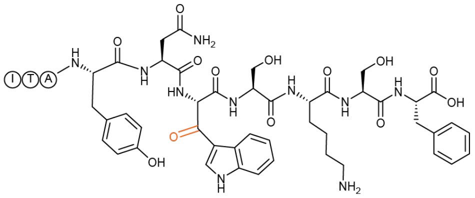

B

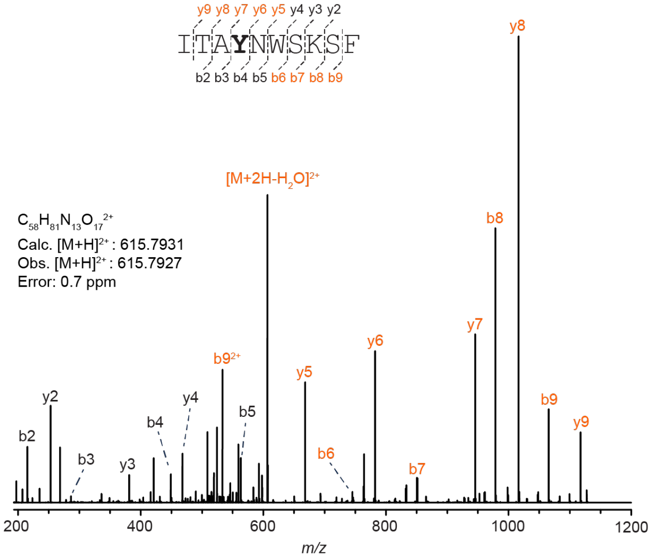

C

| Ion | Calculated Mass | Observed Mass | Error (ppm) |
|---|---|---|---|
| $[M+2H]^{2+}$ | 615.7931 | 615.7926 | 0.81 |
| $[M+2H-H_2O]^{2+}$ | 606.7879 | 606.7869 | 1.65 |
| $b9^+$ | 1065.5000 | 1065.4983 | 1.60 |
| $b9^{2+}$ | 533.2536 | 533.2529 | 1.31 |
| $y2^+$ | 253.1183 | 253.1179 | 1.58 |
| $b8^+$ | 978.4679 | 978.4664 | 1.53 |
| $y3^+$ | 381.2133 | 381.2127 | 1.57 |
| $b7^+$ | 850.3730 | 850.3719 | 1.29 |
| $y4^+$ | 468.2453 | 468.2446 | 1.49 |
| $b6^+$ | 763.3409 | 763.3401 | 1.05 |
| $b5^+$ | 563.2824 | 563.2815 | 1.60 |
| $y5^+$ | 668.3038 | 668.3027 | 1.65 |
| $b4^+$ | 449.2395 | 449.2388 | 1.56 |
| $y6^+$ | 782.3468 | 782.3457 | 1.41 |
| $b3^+$ | 286.1762 | 286.1757 | 1.75 |
| $y7^+$ | 945.4101 | 945.4086 | 1.59 |
| $b2^+$ | 215.1391 | 215.1386 | 2.32 |
| $y8^+$ | 1016.4472 | 1016.4454 | 1.77 |
| $y8^{2+}$ | 508.7273 | 508.7265 | 1.57 |
| $y9^+$ | 1117.4949 | 1117.493 | 1.70 |

**Figure S48: HR-MS/MS of the W1Y variant with a β-oxotryptophan modification.** (A) Hypothetical structure of the modified species based on HR-MS/MS fragmentation. The modification is highlighted in orange. (B) CID spectrum of the +14 Da species associated with the GluC-digested W1Y darobactin variant. The sequence of the GluC-digested peptide is provided with the substitution bolded. Orange colored daughter ions include the modification. (C) Observed masses and errors from CID analysis.

A

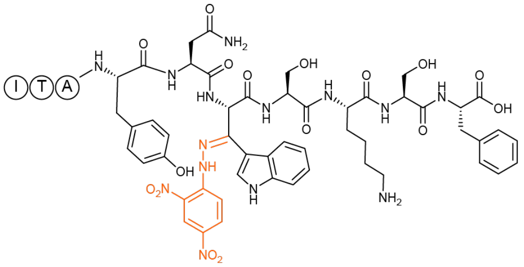

B

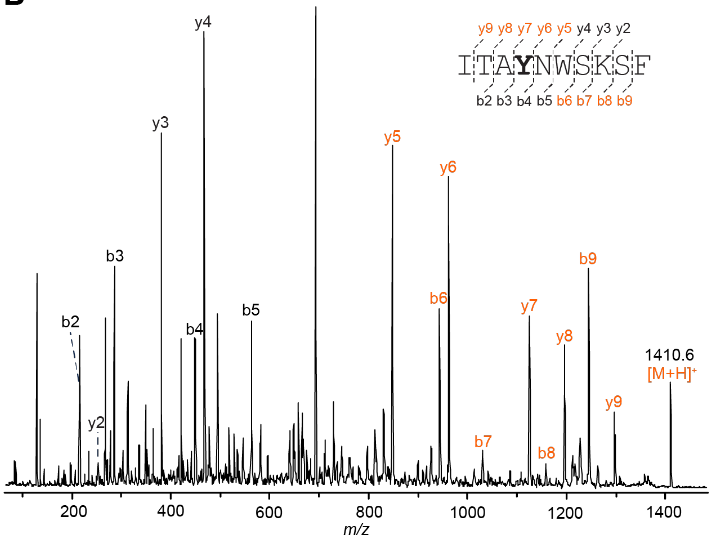

C

| Ion | Calculated Mass | Observed Mass |
|---|---|---|
| $[M+H]^+$ | 1410.6 | 1410.6 |
| $b9^+$ | 1245.5 | 1245.5 |
| $y2^+$ | 253.1 | 253.2 |
| $b8^+$ | 1158.5 | 1158.5 |
| $y3^+$ | 381.2 | 381.3 |
| $b7^+$ | 1030.4 | 1030.4 |
| $y4^+$ | 468.2 | 468.3 |
| $b6^+$ | 943.4 | 943.3 |
| $b5^+$ | 563.3 | 563.3 |
| $y5^+$ | 848.3 | 848.3 |
| $b4^+$ | 449.2 | 449.3 |
| $y6^+$ | 962.4 | 962.3 |
| $b3^+$ | 286.2 | 286.2 |
| $y7^+$ | 1125.4 | 1125.4 |
| $b2^+$ | 215.1 | 215.2 |
| $y8^+$ | 1196.5 | 1196.5 |
| $y9^+$ | 1297.5 | 1297.5 |

**Figure S49: MALDI-LIFT-TOF/TOF-MS analysis of hydrazone labeled DarA-W1Y.** (A) Hypothetical structure of the modified species based on MALDI-LIFT-TOF/TOF-MS fragmentation. The modification is highlighted in orange. (B) LID spectra of the hydrazone labeled DarA-W1Y variant. The sequence of the GluC-digested peptide is provided with the substitution bolded. Orange colored daughter ions include the modification. (C) Table of observed daughter ions.

A

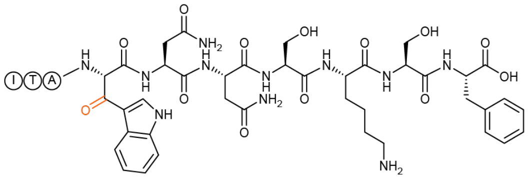

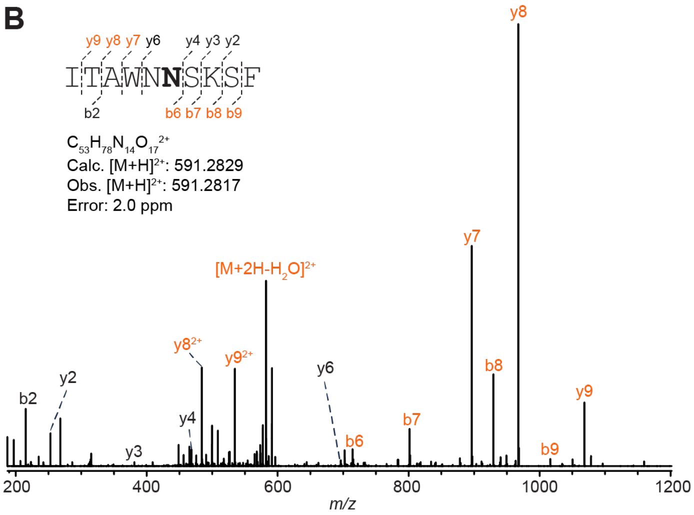

C

| Ion | Calculated Mass | Observed Mass | Error (ppm) |
|---|---|---|---|
| $[M+2H]^{2+}$ | 591.2829 | 591.2816 | 2.2 |
| $[M+2H-H_2O]^{2+}$ | 582.2777 | 582.2761 | 2.7 |
| $b9^+$ | 1016.4796 | 1016.4769 | 2.7 |
| $b9^{2+}$ | 508.7434 | 508.7421 | 2.6 |
| $y2^+$ | 253.1183 | 253.1177 | 2.4 |
| $b8^+$ | 929.4475 | 929.4452 | 2.5 |
| $b8^{2+}$ | 465.2274 | 465.2263 | 2.4 |
| $y3^+$ | 381.2133 | 381.2123 | 2.6 |
| $b7^+$ | 801.3526 | 801.3506 | 2.5 |
| $y4^+$ | 468.2453 | 468.2441 | 2.6 |
| $b6^+$ | 714.3205 | 714.3187 | 2.5 |
| $y6^+$ | 696.3312 | 696.3292 | 2.9 |
| $b3^+$ | 286.1762 | 286.1755 | 2.4 |
| $y7^+$ | 896.3897 | 896.3874 | 2.6 |
| $y7^{2+}$ | 448.6985 | 448.6974 | 2.5 |
| $b2^+$ | 215.1391 | 215.1385 | 2.8 |
| $y8^+$ | 967.4268 | 967.4240 | 2.9 |
| $y8^{2+}$ | 484.2171 | 484.2159 | 2.5 |
| $y9^+$ | 1068.4745 | 1068.4716 | 2.7 |
| $y9^{2+}$ | 534.7410 | 534.7395 | 2.8 |

**Figure S50: HR-MS/MS of the DarA-W3N variant with a β-oxotryptophan modification.** (A) Hypothetical structure of the modified species based on HR-MS/MS fragmentation. The modification is highlighted in orange. (B) CID spectrum of the +14 Da modified variant. The sequence of the GluC-digested peptide is provided with the substitution bolded. Orange colored daughter ions include the modification. (C) Observed masses and errors from CID analysis.

**A**

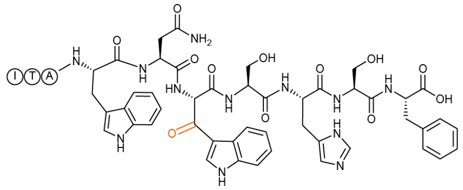

**B**

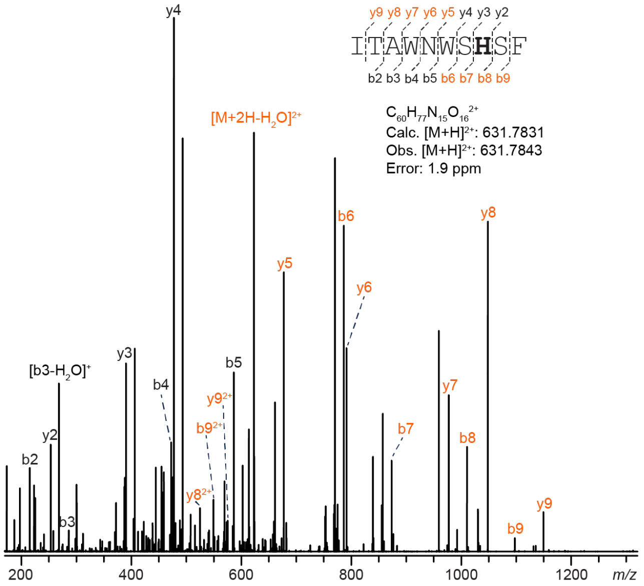

**C**

| Ion | Calculated Mass | Observed Mass | Error (ppm) |
|---|---|---|---|
| $[M+2H]^{2+}$ | 631.7831 | 631.7843 | 1.9 |
| $[M+2H-H_2O]^{2+}$ | 622.7778 | 622.7787 | 1.4 |
| $b9^{+}$ | 1097.4799 | 1097.4814 | 1.4 |
| $b9^{2+}$ | 549.2436 | 549.2444 | 1.5 |
| $y2^{+}$ | 253.1183 | 253.1187 | 1.6 |
| $b8^{+}$ | 1010.4479 | 1010.4491 | 1.2 |
| $y3^{+}$ | 390.1772 | 390.1777 | 1.3 |
| $b7^{+}$ | 873.3890 | 873.3899 | 1.0 |
| $y4^{+}$ | 477.2093 | 477.2098 | 1.0 |
| $b6^{+}$ | 786.3569 | 786.3579 | 1.3 |
| $b5^{+}$ | 586.2984 | 586.2993 | 1.5 |
| $y5^{+}$ | 677.2678 | 677.2687 | 1.3 |
| $b4^{+}$ | 472.2555 | 472.2561 | 1.3 |
| $y6^{+}$ | 791.3107 | 791.3121 | 1.8 |
| $b3^{+}$ | 286.1762 | 286.1767 | 1.7 |
| $[b3-H_2O]^{+}$ | 268.1656 | 268.1660 | 1.5 |
| $y7^{+}$ | 977.3900 | 977.3912 | 1.2 |
| $b2^{+}$ | 215.1391 | 215.1393 | 0.9 |
| $y8^{+}$ | 1048.4271 | 1048.4284 | 1.2 |
| $y8^{2+}$ | 524.7172 | 524.7178 | 1.1 |
| $y9^{+}$ | 1149.4748 | 1149.4760 | 1.0 |
| $y9^{2+}$ | 575.2411 | 575.2420 | 1.6 |

**Figure S51: HR-MS/MS of the DarA-K5H variant with a β-oxotryptophan modification.** (A) Hypothetical structure of the modified species based on HR-MS/MS fragmentation. The modification is highlighted in orange. (B) CID spectrum of the +14 Da modified variant. The sequence of the GluC-digested peptide is provided with the substitution bolded. Orange colored daughter ions include the modification. (C) Observed masses and errors from CID analysis.

A

B

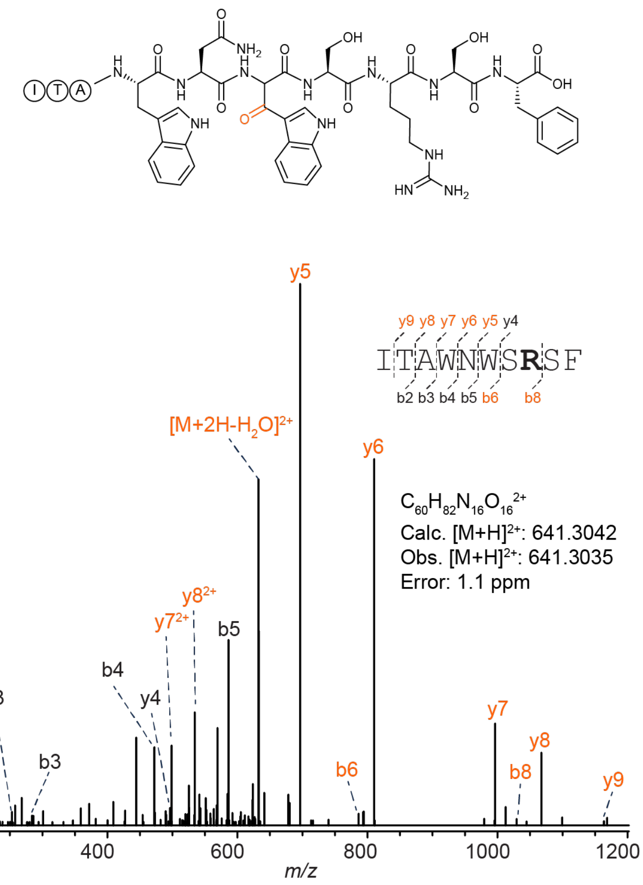

C

| Ion | Calculated Mass | Observed Mass | Error (ppm) |
|---|---|---|---|
| $[M+2H]^{2+}$ | 641.3042 | 641.3035 | 1.1 |
| $[M+2H-H_2O]^{2+}$ | 632.2989 | 632.2980 | 1.4 |
| $b8^{+}$ | 1029.4885 | 1029.4901 | 1.6 |
| $y4^{+}$ | 496.2515 | 496.2510 | 1.0 |
| $b6^{+}$ | 786.3569 | 786.3561 | 1.0 |
| $b5^{+}$ | 586.2984 | 586.2980 | 0.7 |
| $y5^{+}$ | 696.3100 | 696.3091 | 1.3 |
| $b4^{+}$ | 472.2555 | 472.2549 | 1.3 |
| $y6^{+}$ | 810.3529 | 810.3520 | 1.1 |
| $b3^{+}$ | 286.1762 | 286.1758 | 1.4 |
| $y7^{+}$ | 996.4322 | 996.4300 | 2.2 |
| $y7^{2+}$ | 498.7198 | 498.7192 | 1.2 |
| $b2^{+}$ | 215.1391 | 215.1388 | 1.4 |
| $y8^{+}$ | 1067.4693 | 1067.4675 | 1.7 |
| $y8^{2+}$ | 534.2383 | 534.2372 | 2.1 |
| $y9^{+}$ | 1168.5170 | 1168.5159 | 0.9 |

**Figure S52: HR-MS/MS of the DarA-K5R variant with a β-oxotryptophan modification.** (A) Hypothetical structure of the modified species based on HR-MS/MS fragmentation. The modification is highlighted in orange. (B) CID spectrum of the +14 Da modified variant. The sequence of the GluC-digested peptide is provided with the substitution bolded. Orange colored daughter ions include the modification. (C) Observed masses and errors from CID analysis.

**A**

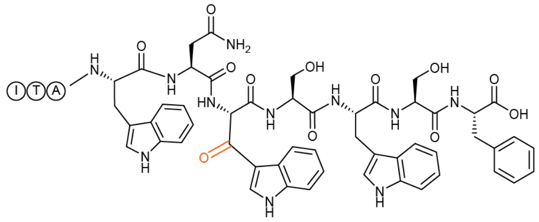

**B**

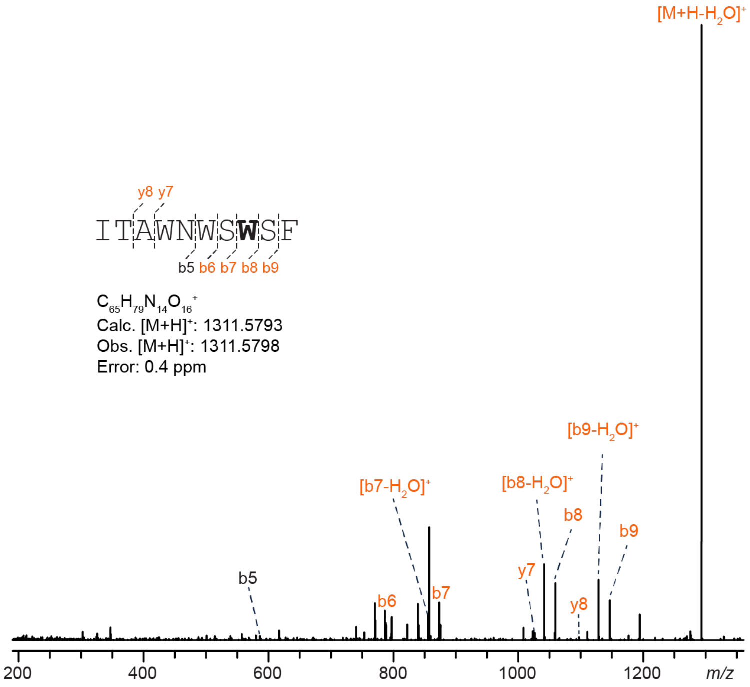

**C**

| Ion | Calculated Mass | Observed Mass | Error (ppm) |
|---|---|---|---|
| $[M+H]^+$ | 1311.5793 | 1311.5798 | 0.4 |
| $[M+H-H_2O]^+$ | 1293.5687 | 1293.5707 | 1.5 |
| $b9^+$ | 1146.5003 | 1146.4996 | 0.6 |
| $[b9-H_2O]^+$ | 1128.4898 | 1128.4904 | 0.5 |
| $b8^+$ | 1059.4683 | 1059.4682 | 0.1 |
| $[b8-H_2O]^+$ | 1041.4578 | 1041.4580 | 0.2 |
| $b7^+$ | 873.3890 | 873.3900 | 1.1 |
| $[b7-H_2O]^+$ | 855.3784 | 855.3794 | 1.2 |
| $b6^+$ | 786.3569 | 786.3583 | 1.8 |
| $b5^+$ | 586.2984 | 586.2998 | 2.4 |
| $y7^+$ | 1026.4104 | 1026.4078 | 2.5 |
| $y8^+$ | 1097.4475 | 1097.4477 | 0.2 |

**Figure S53: HR-MS/MS of the DarA-K5W variant with a β-oxotryptophan modification.** (A) Hypothetical structure of the modified species based on HR-MS/MS fragmentation. The modification is highlighted in orange. (B) CID spectrum of the +14 Da modified variant. The sequence of the GluC-digested peptide is provided with the substitution bolded. Orange colored daughter ions include the modification. (C) Observed masses and errors from CID analysis.

**A**

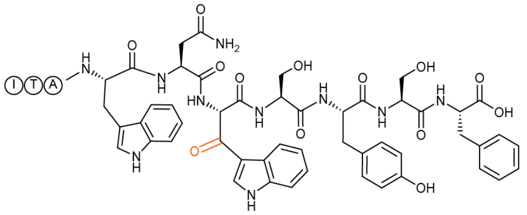

**B**

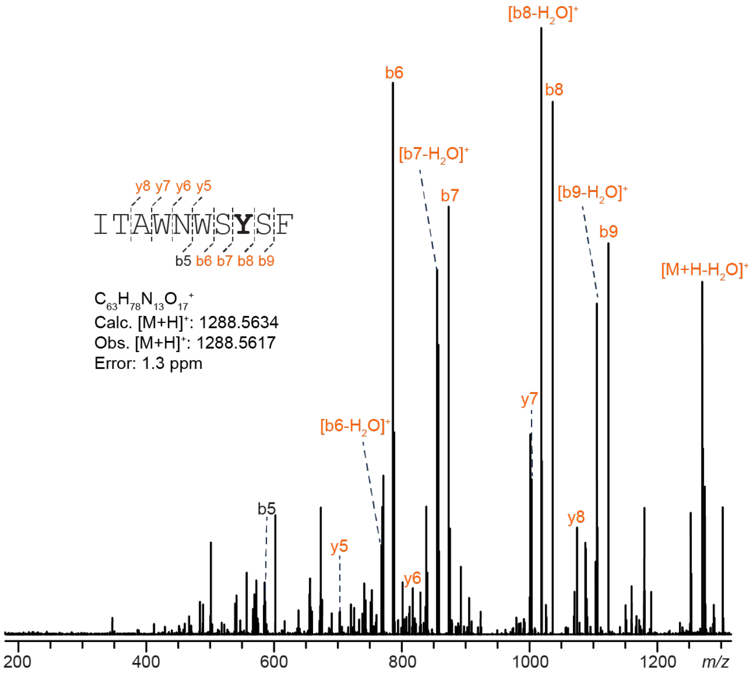

**C**

| Ion | Calculated Mass | Observed Mass | Error (ppm) |
|---|---|---|---|
| $[M+H]^+$ | 1288.5634 | 1288.5617 | 1.3 |
| $[M+H-H_2O]^+$ | 1270.5527 | 1270.5501 | 2.0 |
| $b9^+$ | 1123.4843 | 1123.4816 | 2.4 |
| $[b9-H_2O]^+$ | 1105.4738 | 1105.4710 | 2.5 |
| $b8^+$ | 1036.4523 | 1036.4496 | 2.6 |
| $[b8-H_2O]^+$ | 1018.4418 | 1018.4394 | 2.4 |
| $b7^+$ | 873.3890 | 873.3870 | 2.3 |
| $[b7-H_2O]^+$ | 855.3784 | 855.3763 | 2.5 |
| $b6^+$ | 786.3569 | 786.3553 | 2.0 |
| $[b6-H_2O]^+$ | 768.3464 | 768.3448 | 2.1 |
| $b5^+$ | 586.2984 | 586.2970 | 2.4 |
| $y5^+$ | 703.2722 | 703.2704 | 2.6 |
| $y6^+$ | 817.3151 | 817.3134 | 2.1 |
| $y7^+$ | 1003.3944 | 1003.3920 | 2.4 |
| $y8^+$ | 1074.4316 | 1074.4291 | 2.3 |

**Figure S54: HR-MS/MS of the DarA-K5Y variant with a β-oxotryptophan modification.** (A) Hypothetical structure of the modified species based on HR-MS/MS fragmentation. The modification is highlighted in orange. (B) CID spectrum of the +14 Da modified variant. The sequence of the GluC-digested peptide is provided with the substitution bolded. Orange colored daughter ions include the modification. (C) Observed masses and errors from CID analysis.

**A**

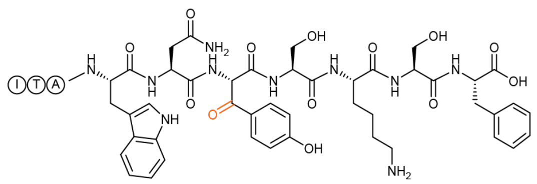

**B**

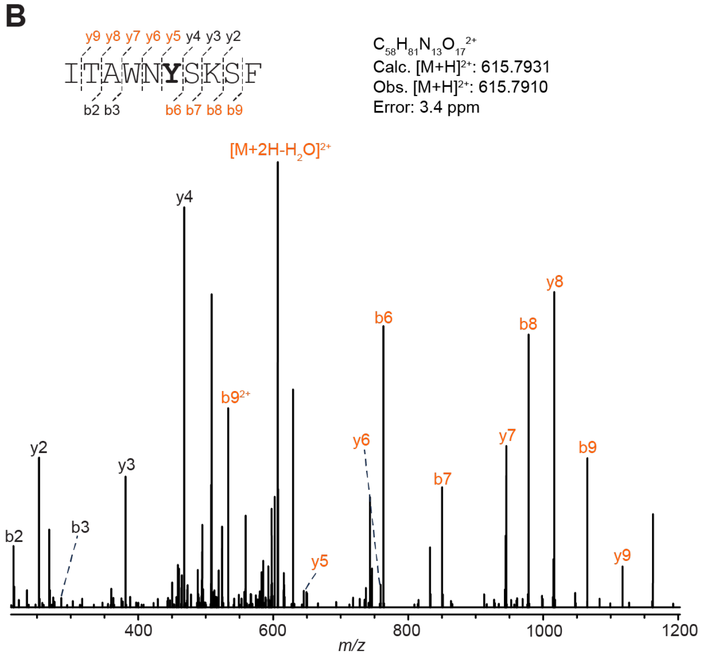

**C**

| Ion | Calculated Mass | Observed Mass | Error (ppm) |
|---|---|---|---|
| $[M+2H]^{2+}$ | 615.7931 | 615.791 | 3.4 |
| $[M+2H-H_2O]^{2+}$ | 606.7879 | 606.7853 | 4.3 |
| $b9^{+}$ | 1065.5000 | 1065.497 | 2.8 |
| $b9^{2+}$ | 533.2536 | 533.2522 | 2.6 |
| $y2^{+}$ | 253.1183 | 253.1176 | 2.8 |
| $b8^{+}$ | 978.4679 | 978.4648 | 3.2 |
| $y3^{+}$ | 381.2133 | 381.2122 | 2.9 |
| $b7^{+}$ | 850.3730 | 850.3706 | 2.8 |
| $y4^{+}$ | 468.2453 | 468.2439 | 3.0 |
| $b6^{+}$ | 763.3409 | 763.339 | 2.5 |
| $y5^{+}$ | 645.2879 | 645.2861 | 2.8 |
| $y6^{+}$ | 759.3308 | 759.329 | 2.4 |
| $y7^{+}$ | 850.3730 | 850.3709 | 2.5 |
| $b3^{+}$ | 286.1762 | 286.1753 | 3.1 |
| $y8^{+}$ | 978.4680 | 978.4651 | 3.0 |
| $b2^{+}$ | 215.1391 | 215.1384 | 3.3 |
| $y9^{+}$ | 1117.4949 | 1117.4917 | 2.9 |

**Figure S55: HR-MS/MS analysis of the DarA-W3Y variant with a β-oxotyrosine modification.** (A) Hypothetical structure of the modified species based on HR-MS/MS fragmentation. The modification is highlighted in orange. (B) CID spectrum of the +14 Da modified variant. The sequence of the GluC-digested peptide is provided with the substitution bolded. Orange colored daughter ions include the modification. (C) Observed masses and errors from CID analysis.

**Table S3: Observed daughter ions for LIFT-MS/MS of the W3Y hydrazone adduct.** Refer to **Figure 3** for the LIFT-MS/MS spectrum.

| Ion | Calculated Mass | Observed Mass |
|---|---|---|
| $[M+H]^+$ | 1410.6 | 1410.6 |
| $b9^+$ | 1245.5 | 1245.5 |
| $y2^+$ | 253.1 | 253.2 |
| $b8^+$ | 1158.5 | 1158.5 |
| $y3^+$ | 381.2 | 381.3 |
| $b7^+$ | 1030.4 | 1030.4 |
| $y4^+$ | 468.2 | 468.3 |
| $b6^+$ | 943.4 | 943.3 |
| $b5^+$ | 563.3 | 563.3 |
| $y5^+$ | 848.3 | 848.3 |
| $b4^+$ | 449.2 | 449.3 |
| $y6^+$ | 962.4 | 962.3 |
| $b3^+$ | 286.2 | 286.2 |
| $y7^+$ | 1125.4 | 1125.4 |
| $b2^+$ | 215.1 | 215.2 |
| $y8^+$ | 1196.5 | 1196.5 |
| $y9^+$ | 1297.5 | 1297.5 |

A

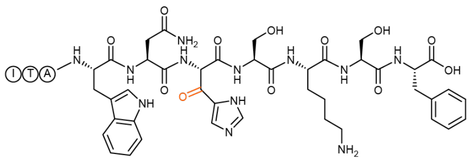

B

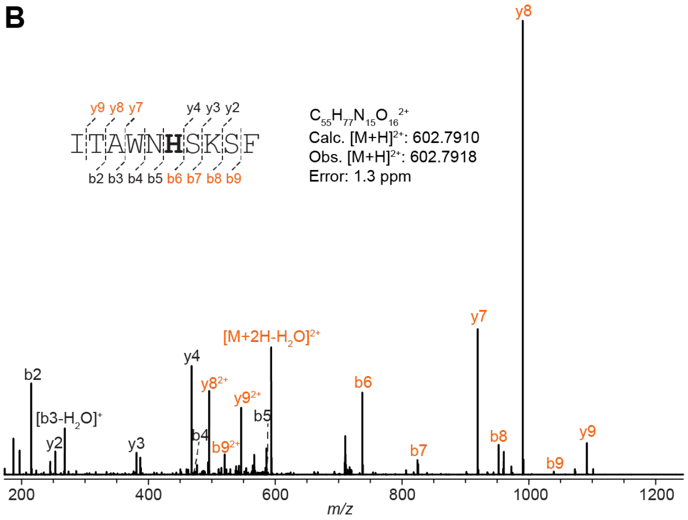

C

| Ion | Calculated Mass | Observed Mass | Error (ppm) |
|---|---|---|---|
| $[M+2H]^{2+}$ | 602.7910 | 602.7918 | 1.3 |
| $[M+2H-H_2O]^{2+}$ | 593.7857 | 593.7867 | 1.7 |
| $b9^{+}$ | 1039.4956 | 1039.4971 | 1.4 |
| $b9^{2+}$ | 520.2515 | 520.2523 | 1.5 |
| $y2^{+}$ | 253.1183 | 253.1187 | 1.6 |
| $b8^{+}$ | 952.4636 | 952.4648 | 1.3 |
| $b8^{2+}$ | 476.7355 | 476.7362 | 1.5 |
| $y3^{+}$ | 381.2133 | 381.2139 | 1.6 |
| $b7^{+}$ | 824.3686 | 824.3701 | 1.8 |
| $y4^{+}$ | 468.2453 | 468.246 | 1.5 |
| $b6^{+}$ | 737.3366 | 737.3377 | 1.5 |
| $b5^{+}$ | 586.2984 | 586.2993 | 1.5 |
| $y5^{+}$ | 619.2835 | 619.2844 | 1.5 |
| $b4^{+}$ | 472.2555 | 472.25662 | 2.4 |
| $y6^{+}$ | 733.3264 | 733.3279 | 2.0 |
| $b3^{+}$ | 286.1762 | 286.1766 | 1.4 |
| $[b3- H_2O]^{+}$ | 268.1656 | 268.1661 | 1.9 |
| $y7^{+}$ | 919.4057 | 919.4068 | 1.2 |
| $y7^{2+}$ | 460.2065 | 460.2072 | 1.5 |
| $b2^{+}$ | 215.1391 | 215.1394 | 1.4 |
| $y8^{+}$ | 990.4429 | 990.4439 | 1.0 |
| $y8^{2+}$ | 495.7251 | 495.7258 | 1.4 |
| $y9^{+}$ | 1091.4905 | 1091.4919 | 1.3 |
| $y9^{2+}$ | 546.2489 | 546.2498 | 1.6 |

**Figure S56: HR-MS/MS of the DarA-W3H variant with a β-oxohistidine modification.** (A) Hypothetical structure of the modified species based on HR-MS/MS fragmentation. The modification is highlighted in orange. (B) CID spectrum of the +14 Da modified variant. The sequence of the GluC-digested peptide is provided with the substitution bolded. Orange colored daughter ions include the modification. (C) Observed masses and errors from CID analysis.

**A**

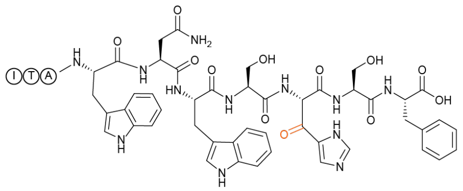

**B**

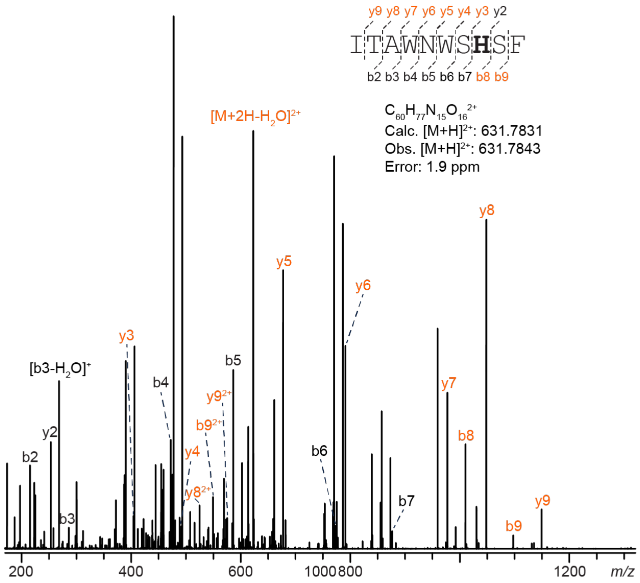

**C**

| Ion | Calculated Mass | Observed Mass | Error (ppm) |
|---|---|---|---|
| $[M+2H]^{2+}$ | 631.7831 | 631.7843 | 1.9 |
| $[M+2H-H_2O]^{2+}$ | 622.7778 | 622.7787 | 1.4 |
| $b9^{+}$ | 1097.4799 | 1097.4814 | 1.4 |
| $b9^{2+}$ | 549.2436 | 549.2444 | 1.5 |
| $y2^{+}$ | 253.1183 | 253.1187 | 1.6 |
| $b8^{+}$ | 1010.4479 | 1010.4491 | 1.2 |
| $y3^{+}$ | 404.1564 | 404.1571 | 1.7 |
| $y4^{+}$ | 491.1885 | 491.1893 | 1.6 |
| $b6^{+}$ | 772.3777 | 772.3785 | 1.0 |
| $b5^{+}$ | 586.2984 | 586.2993 | 1.5 |
| $y5^{+}$ | 677.2678 | 677.2687 | 1.3 |
| $b4^{+}$ | 472.2555 | 472.2561 | 1.3 |
| $y6^{+}$ | 791.3107 | 791.3121 | 1.8 |
| $b3^{+}$ | 286.1762 | 286.1767 | 1.7 |
| $[b3-H_2O]^{+}$ | 268.1656 | 268.1660 | 1.5 |
| $y7^{+}$ | 977.3900 | 977.3912 | 1.2 |
| $b2^{+}$ | 215.1391 | 215.1393 | 0.9 |
| $y8^{+}$ | 1048.4271 | 1048.4284 | 1.2 |
| $y8^{2+}$ | 524.7172 | 524.7178 | 1.1 |
| $y9^{+}$ | 1149.4748 | 1149.4760 | 1.0 |
| $y9^{2+}$ | 575.2411 | 575.2420 | 1.6 |

**Figure S57: HR-MS/MS of the DarA-K5H variant with a β-oxohistidine modification.** (A) Hypothetical structure of the modified species based on HR-MS/MS fragmentation. The modification is highlighted in orange. (B) CID spectrum of the +14 Da modified variant. The sequence of the GluC-digested peptide is provided with the substitution bolded. Orange colored daughter ions include the modification. (C) Observed masses and errors from CID analysis.

**A**

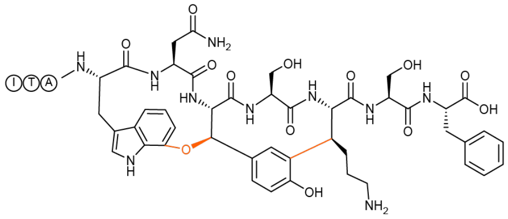

**B**

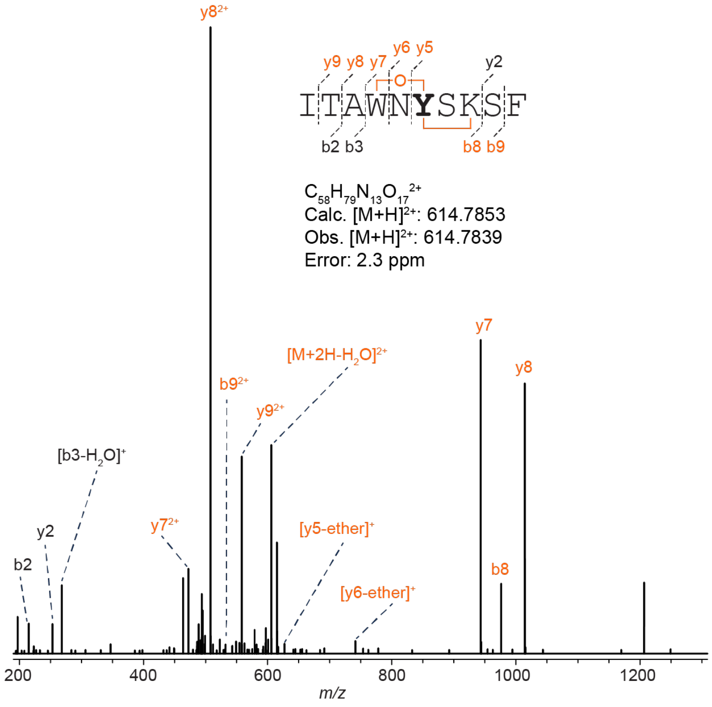

**C**

| Ion | Calculated Mass | Observed Mass | Error (ppm) |
|---|---|---|---|
| $[M+2H]^{2+}$ | 614.7853 | 614.7839 | 2.3 |
| $[M+2H-H_2O]^{2+}$ | 605.7801 | 605.7783 | 3.0 |
| $b9^{2+}$ | 532.2458 | 532.2439 | 3.6 |
| $y2^{+}$ | 253.1183 | 253.1175 | 3.2 |
| $b8^{+}$ | 976.4523 | 976.4493 | 3.1 |
| $b8^{2+}$ | 488.7293 | 488.7284 | 1.8 |
| $b3^{+}$ | 286.1762 | 286.1753 | 3.1 |
| $[y5\text{-ether}]^{+}$ | 627.2779 | 627.2754 | 4.0 |
| $y7^{+}$ | 943.3945 | 943.3916 | 3.1 |
| $y7^{2+}$ | 472.2009 | 472.1994 | 3.2 |
| $b2^{+}$ | 215.1391 | 215.1384 | 3.3 |
| $y8^{+}$ | 1014.4316 | 1014.4283 | 3.3 |
| $y8^{2+}$ | 507.7195 | 507.7179 | 3.2 |
| $y9^{2+}$ | 558.2433 | 558.2417 | 2.9 |

**Figure S58: HR-MS/MS analysis of the DarA-W3Y variant with a +12 Da modification.** (A) Structure of the modified species based on HR-MS/MS fragmentation. The modification is highlighted in orange. (B) CID spectrum of the +14 Da species associated with the GluC-digested W3Y darobactin variant. The sequence of the GluC-digested peptide is provided with the substitution bolded. Orange colored daughter ions include the modification. Distinguishing ether crosslink fragments are indicated. (C) Observed masses and errors from CID analysis.

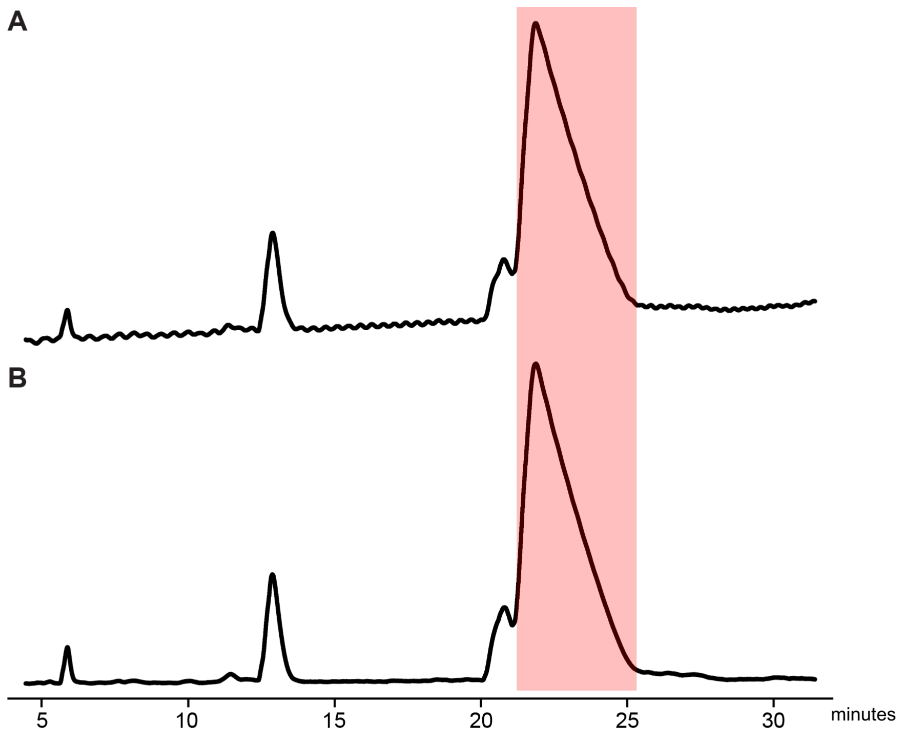

**Figure S59: UHPLC purification of darobactin W3Y.** (A) Chromatographic monitoring at 220 nm and (B) 280 nm from C18 reverse-phase purification. The light red fraction eluting at 22 min contained darobactin W3Y and was collected for NMR spectroscopic evaluation.

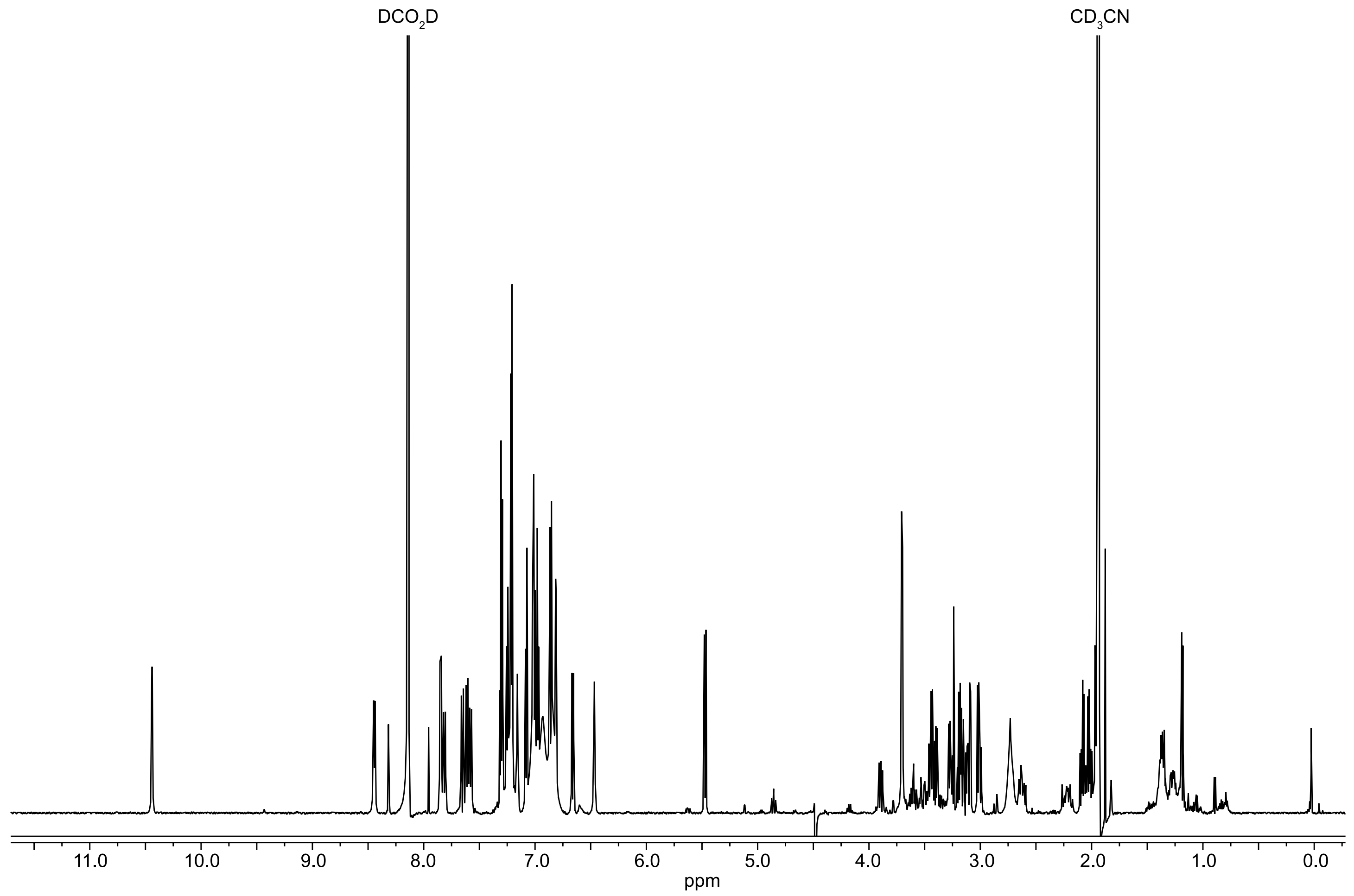

**Figure S60: $^{1}$H NMR spectrum of darobactin W3Y.** Data were acquired in 66% $H_2O$: 32% $CD_3CN$: 2% $DCO_2D$ using a water-suppression (WET) method.

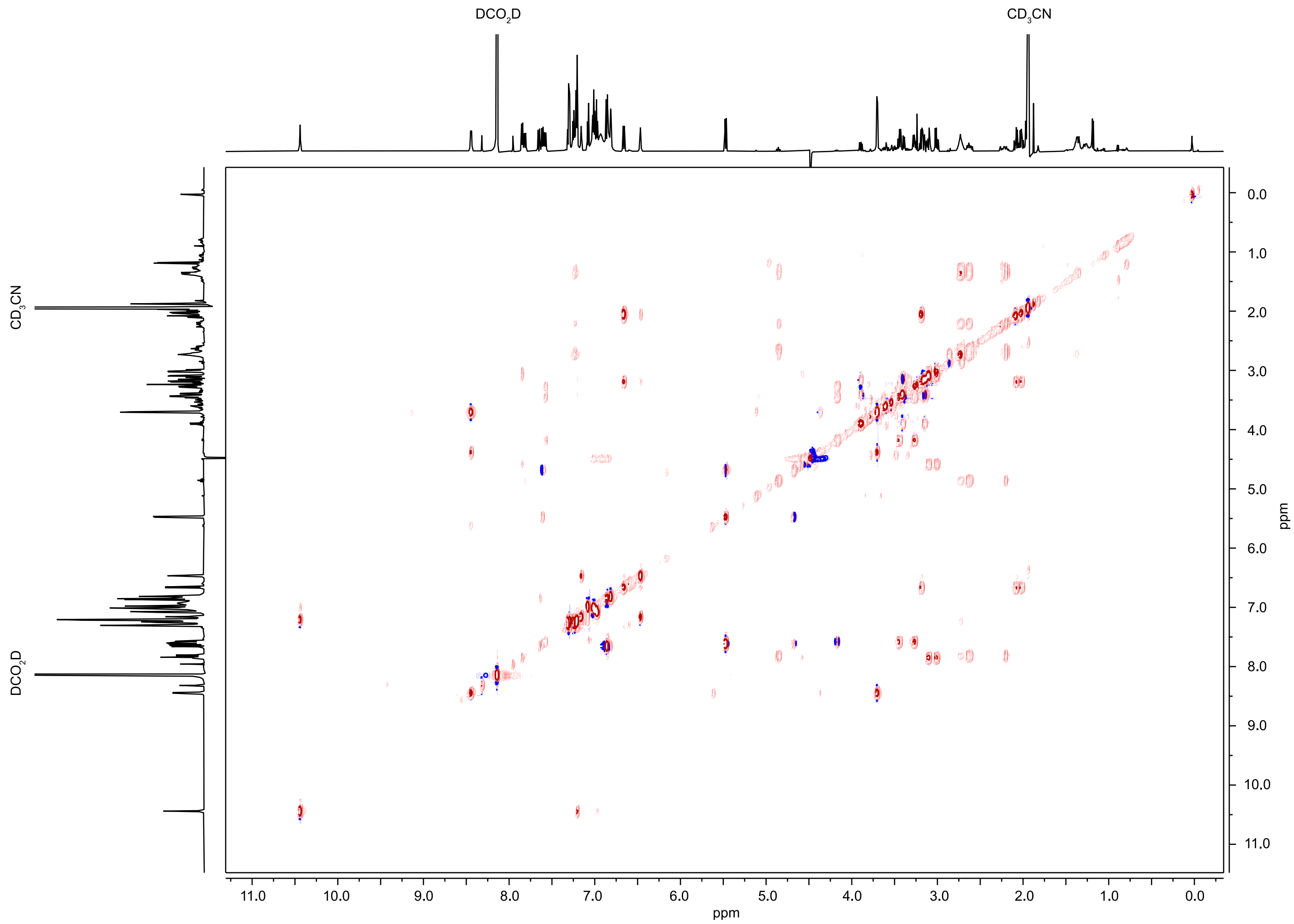

**Figure S61: [1]H-[1]H TOCSY NMR spectrum of darobactin W3Y.** Data were acquired in 66% $H_2O$: 32% $CD_3CN$: 2% $DCO_2D$ using a water-suppression (WET) method.

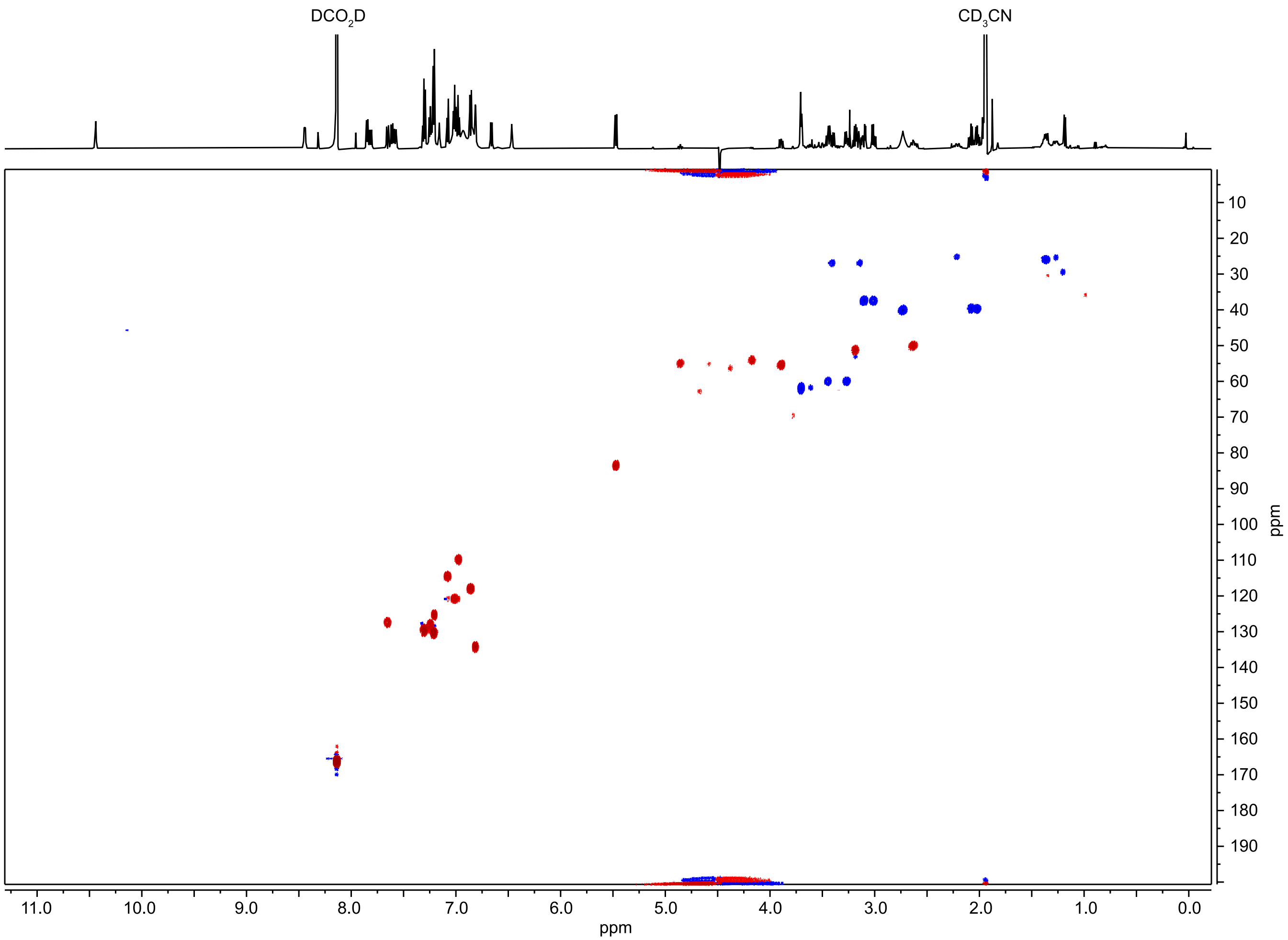

**Figure S62: $^{1}H$-$^{13}C$ HSQC NMR spectrum of darobactin W3Y.** Data were acquired in 66% $H_2O$: 32% $CD_3CN$: 2% $DCO_2D$ using a water-suppression (WET) method.

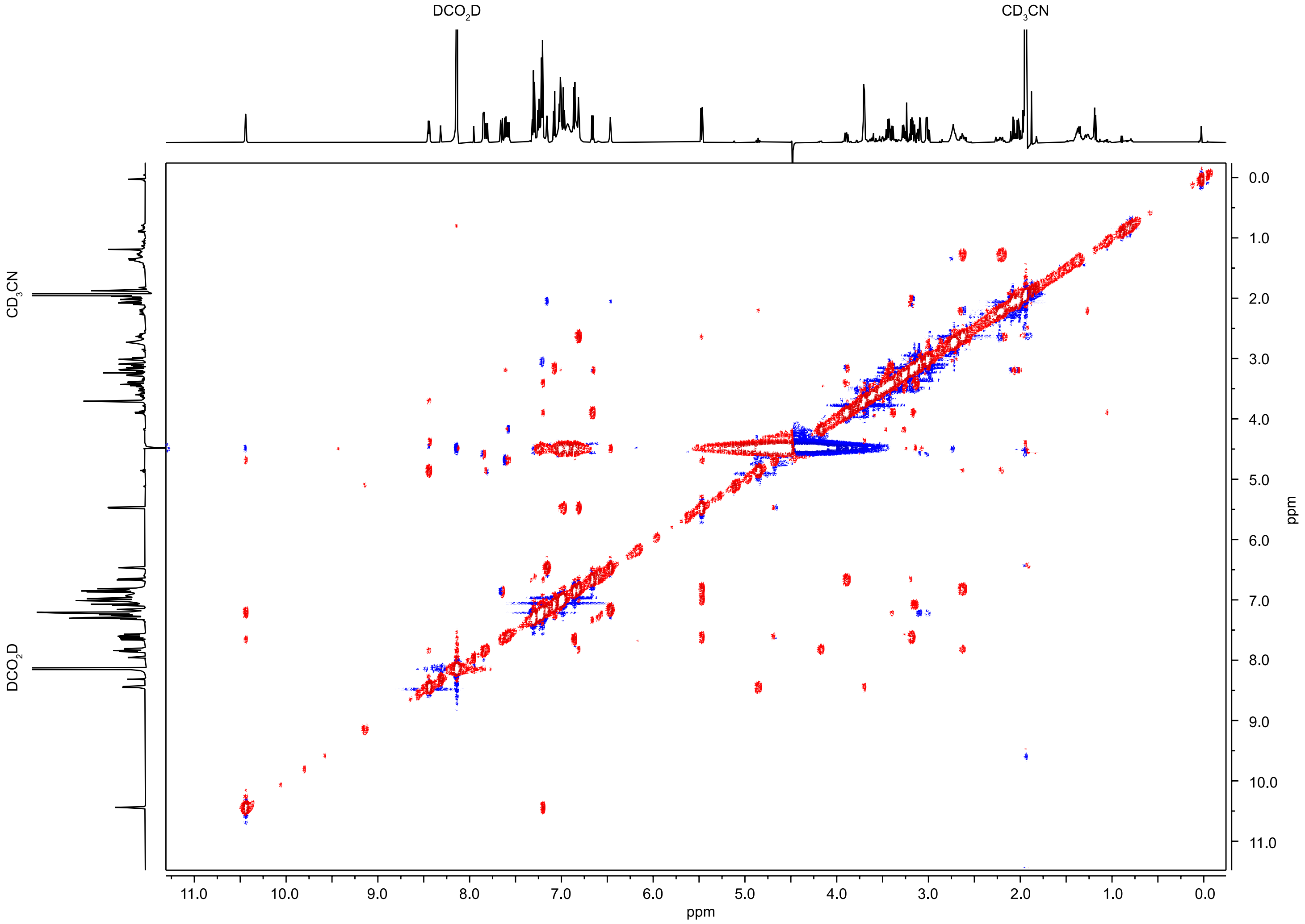

**Figure S63: $^{1}$H-$^{1}$H NOESY NMR spectrum of darobactin W3Y.** Data were acquired in 66% $H_2O$: 32% $CD_3CN$: 2% $DCO_2D$ using a water-suppression (WET) method.

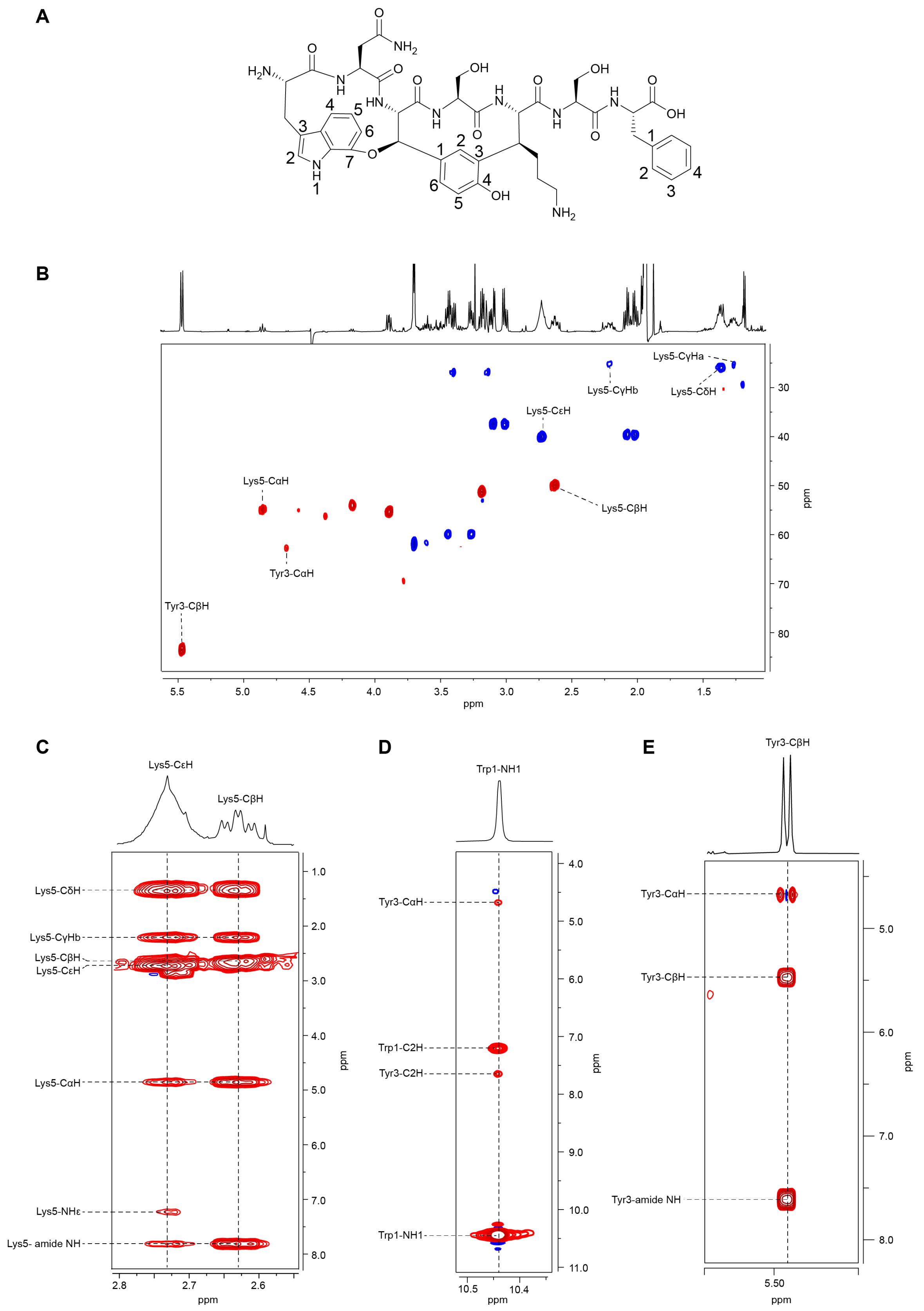

**Figure S64: Additional NMR information for Darobactin W3Y.** (A) Numbered W3R Structure**.** Numbering of atoms directly corresponds to those in **Table S3.** (B) HSQC spectrum of darobactin W3Y with a focus on the aliphatic region. (C) $^1$H-$^1$H TOCSY correlations for Lys5. (D) $^1$H-$^1$H NOESY correlations for Trp1-NH1. (E) $^1$H-$^1$H TOCSY correlations for Trp3-Hβ.

**Table S4: NMR chemical shift assignments of darobactin W3Y.** Data were acquired in 66% $H_2O$: 32% $CD_3CN$: 2% $DCO_2D$ using a water-suppression (WET) method.

| Atom | $\delta_H$, mult. ($J$ in Hz) | $\delta_C$ | | Atom | $\delta_H$, mult. ($J$ in Hz) | $\delta_C$ |
|---|---|---|---|---|---|---|
| Trp1 | | | | Ser4 | | |
| Amide NH | - | - | | Amide NH | 7.85, d (7.8) | NH |
| α | 3.89, dd (11.3, 7.6) | 55.4 (CH) | | α | 4.58, overlapped | 55.0 (CH) |
| $β_a$ | 3.14, m | 26.9 ($CH_2$) | | $β_a$ | 3.01, dd (13.9, 7.5) | 37.4 ($CH_2$) |
| $β_b$ | 3.40, m | 26.9 ($CH_2$) | | $β_b$ | 3.10, dd (13.9, 5.4) | 37.4 ($CH_2$) |
| 1 | 10.44, d (2.7) | NH | | | | |
| 2 | 7.2, m | 125.3 (CH) | | Lys5 | | |
| 3 | - | * | | Amide NH | 7.81, d (10.2) | NH |
| 4 | 7.08, d (7.8) | 114.3 (CH) | | α | 4.86, t (11.0) | 54.9 (CH) |
| 5 | 7.01, t (7.7) | 120.7 (CH) | | β | 2.63, td (11.8, 5.1) | 49.9 (CH) |
| 6 | 6.97, d (7.7) | 109.7 (CH) | | $γ_a$ | 1.27, m | 25.2 ($CH_2$) |
| 7 | - | * | | $γ_b$ | 2.21, m | 25.2 ($CH_2$) |
| | | | | δ | 1.36, m | 25.9 ($CH_2$) |
| Asn2 | | | | ε | 2.73, br s | 40.1 ($CH_2$) |
| Amide NH | 6.65, d (8.1) | NH | | NHε | 7.23, br s | NH |
| α | 3.18, m | 51.3 (CH) | | | | |
| $β_a$ | 2.02, dd (14.1, 7.3) | 39.6 ($CH_2$) | | Ser6 | | |
| $β_b$ | 2.09, dd (14.1, 7.0) | 39.6 ($CH_2$) | | Amide NH | 8.44, d (7.3) | NH |
| $NH_1$ | 6.47, br s | NH | | α | 4.38, overlapped | 56.3 (CH/CH3) |
| $NH_2$ | 7.16, br s | NH | | β | 3.7, d (5.6) | 61.2 (CH2) |
| Tyr3 | | | | Phe7 | | |
| Amide NH | 7.61, d (9.6) | NH | | Amide NH | 7.58, d (9.9) | NH |
| α | 4.67, overlapped | 62.7 (CH) | | α | 4.18, overlapped | 54.0 (CH) |
| β | 5.47, d (9.0) | 83.6 (CH) | | $β_a$ | 3.27, dd (11.7, 7.3) | 60.0 ($CH_2$) |
| 1 | | | | $β_b$ | 3.45, dd (11.7, 7.3) | 60.0 ($CH_2$) |
| 2 | 6.81, d (2.3) | 134.2 (CH) | | 1 | - | * |
| 3 | - | * | | 2 | 7.21, m | 130.2 (CH) |
| 4 | - | * | | 3 | 7.31, t (7.5) | 129.5 (CH) |
| 5 | 6.86, d (8.8) | 117.8 (CH) | | 4 | 7.24, d (7.6) | 127.8 (CH) |
| 6 | 7.65, dd (10.7, 2.1) | 127.4 (CH) | | | | |

* No $^{13}C$ spectrum was recorded, thus, no $\delta_C$ was able to be assigned for the indicated carbon atoms.

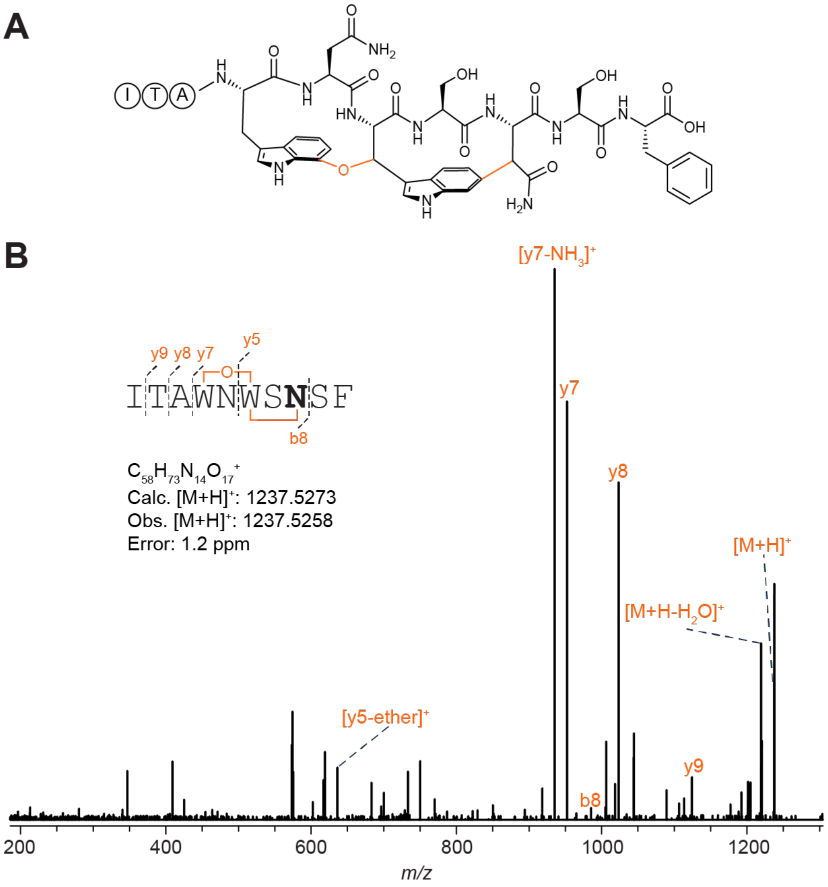

C

| Ion | Calculated Mass | Observed Mass | Error (ppm) |
|---|---|---|---|
| $[M+H]^+$ | 1237.5273 | 1237.5258 | 1.2 |
| $[M+H-H_2O]^+$ | 1219.5167 | 1219.5153 | 1.1 |
| $b8^+$ | 985.4163 | 985.4164 | 0.1 |
| $[y5\text{-ether}]^+$ | 636.2418 | 636.2398 | 3.1 |
| $y7^+$ | 952.3584 | 952.3559 | 2.6 |
| $[y7-NH_3]^+$ | 935.3319 | 935.3297 | 2.4 |
| $y8^+$ | 1023.3956 | 1023.3928 | 2.7 |
| $y9^+$ | 1124.4432 | 1124.4402 | 2.7 |

**Figure S65: HR-MS/MS analysis of darobactin K5N.** (A) Hypothetical structure of the modified species based on HR-MS/MS fragmentation. The connectivity of the linkage is presumed from the known structure of darobactin. The modification is highlighted in orange. (B) CID spectrum of the +12 Da modified variant. The sequence of the GluC-digested peptide is provided with the substitution bolded. Orange colored daughter ions include the modification. Distinguishing ether crosslink fragments are indicated. (C) Observed masses and errors from CID analysis.

**A**

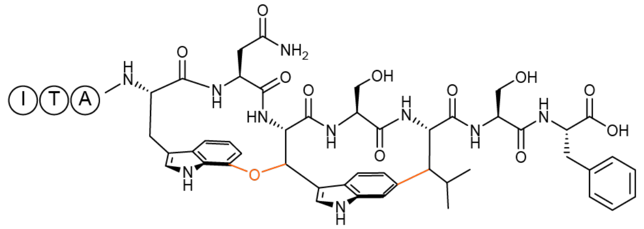

**B**

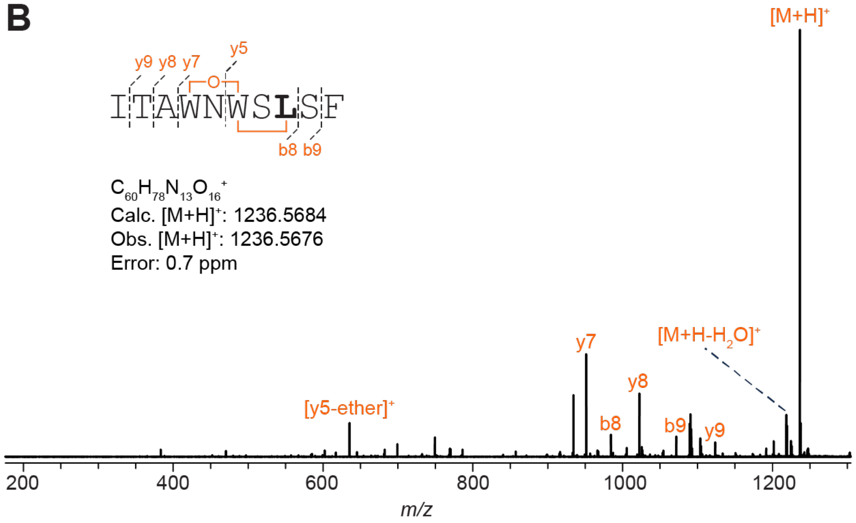

**C**

| Ion | Calculated Mass | Observed Mass | Error (ppm) |
|---|---|---|---|
| $[M+H]^+$ | 1236.5684 | 1236.5676 | 0.6 |
| $[M+H-H_2O]^+$ | 1218.5579 | 1218.5565 | 1.1 |
| $b9^+$ | 1071.4895 | 1071.4882 | 1.2 |
| $b8^+$ | 984.4574 | 984.456 | 1.4 |
| $[y5\text{-ether}]^+$ | 635.2829 | 635.2817 | 1.9 |
| $y7^+$ | 951.3996 | 951.3983 | 1.4 |
| $y8^+$ | 1022.4367 | 1022.4352 | 1.5 |
| $y9^+$ | 1123.4844 | 1123.4827 | 1.5 |

**Figure S66: HR-MS/MS analysis of darobactin K5L.** (A) Hypothetical structure of the modified species based on HR-MS/MS fragmentation. The connectivity of the linkage is presumed from the known structure of darobactin. The modification is highlighted in orange. (B) CID spectrum of the +12 Da modified variant. The sequence of the GluC-digested peptide is provided with the substitution bolded. Orange colored daughter ions include the modification. Distinguishing ether crosslink fragments are indicated. (C) Observed masses and errors from CID analysis.

**A**

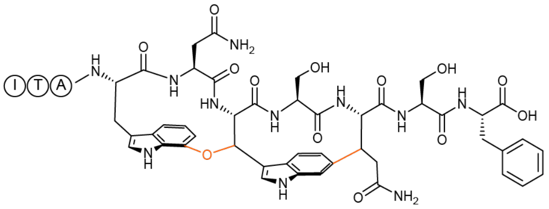

**B**

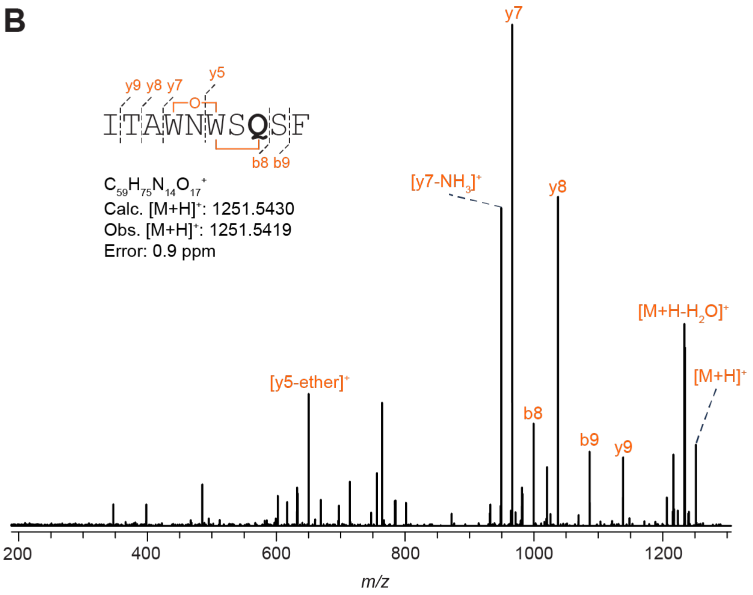

**C**

| Ion | Calculated Mass | Observed Mass | Error (ppm) |
|---|---|---|---|
| $[M+H]^+$ | 1251.5430 | 1251.5419 | 0.9 |
| $[M+H-H_2O]^+$ | 1233.5324 | 1233.5299 | 2.0 |
| $b9^+$ | 1086.4640 | 1086.4617 | 2.1 |
| $b8^+$ | 999.4320 | 999.4298 | 2.2 |
| $[y5\text{-ether}]^+$ | 650.2570 | 650.2554 | 2.5 |
| $y7^+$ | 966.3741 | 966.3719 | 2.3 |
| $[y7-NH_3]^+$ | 949.3475 | 949.3454 | 2.2 |
| $y8^+$ | 1037.4112 | 1037.4086 | 2.5 |
| $y9^+$ | 1138.4589 | 1138.4565 | 2.1 |
| $b4^+$ | 486.2348 | 486.2338 | 2.1 |

**Figure S67: HR-MS/MS analysis of darobactin K5Q.** (A) Hypothetical structure of the modified species based on HR-MS/MS fragmentation. The connectivity of the linkage is presumed from the known structure of darobactin. The modification is highlighted in orange. (B) CID spectrum of the +12 Da modified variant. The sequence of the GluC-digested peptide is provided with the substitution bolded. Orange colored daughter ions include the modification. Distinguishing ether crosslink fragments are indicated. (C) Observed masses and errors from CID analysis.

**A**

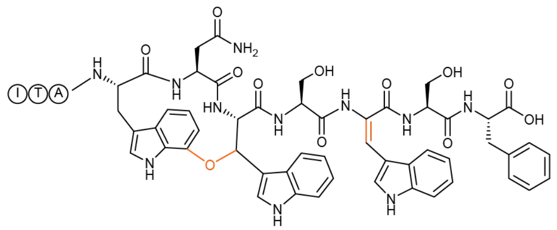

**B**

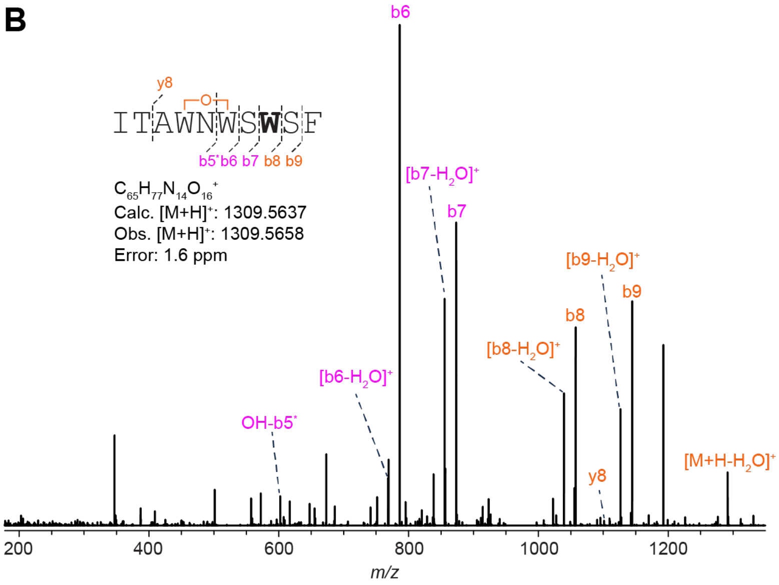

**C**

| Ion | Calculated Mass | Observed Mass | Error (ppm) |
|---|---|---|---|
| $[M+H]^+$ | 1309.5637 | 1309.5658 | 1.6 |
| $[M+H-H_2O]^+$ | 1291.5531 | 1291.5561 | 2.3 |
| $b9^+$ | 1144.4847 | 1144.486 | 1.1 |
| $[b9-H_2O]^+$ | 1226.4763 | 1226.4758 | 0.4 |
| $b8^+$ | 1057.4527 | 1057.4537 | 0.9 |
| $[b8-H_2O]^+$ | 1039.4421 | 1039.4432 | 1.1 |
| $b7^+$ | 873.3890 | 873.3904 | 1.6 |
| $[b7-H_2O]^+$ | 855.3784 | 855.3797 | 1.5 |
| $b6^+$ | 786.3569 | 786.3586 | 2.2 |
| $[b6-H_2O]^+$ | 768.3464 | 768.3481 | 2.2 |
| $b5+OH^*$ | 602.2938 | 602.2943 | 0.8 |
| $y8^+$ | 1095.4320 | 1095.4362 | 3.8 |

**Figure S68: HR-MS/MS analysis of the DarA-K5W variant with a +12 Da modification.** (A) Hypothetical structure of the modified species based on HR-MS/MS fragmentation. The connectivity of the linkage is presumed from the known structure of darobactin. The modification is highlighted in orange. (B) CID spectrum of the +12 Da modified variant. The sequence of the GluC-digested peptide is provided with the substitution bolded. Orange colored daughter ions include the +12 Da modification while those in magenta contain a +14 Da addition. Distinguishing ether crosslink fragments are indicated. (C) Observed masses and errors from CID analysis.

**A**

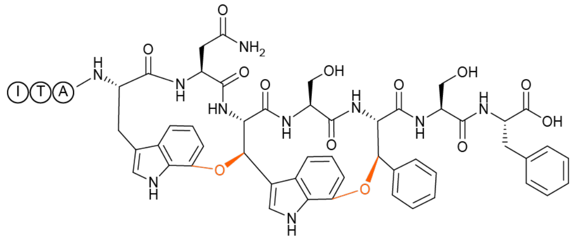

**B**

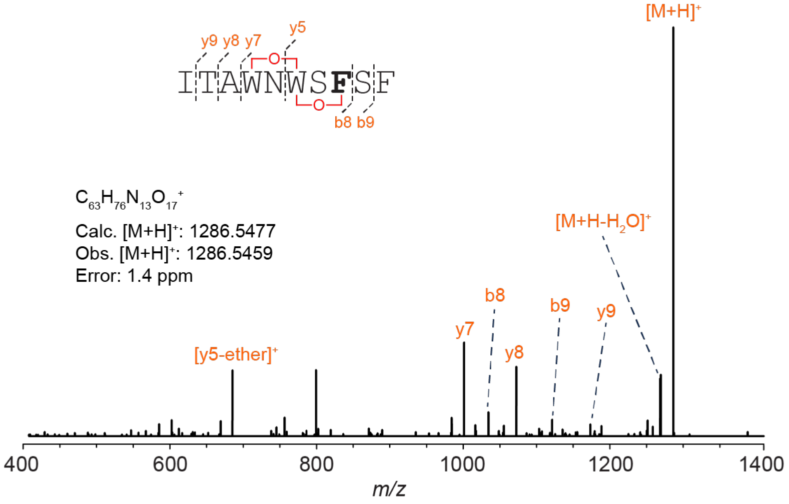

**C**

| Ion | Calculated Mass | Observed Mass | Error (ppm) |
|---|---|---|---|
| $[M+H]^+$ | 1286.5477 | 1286.5459 | 1.4 |
| $[M+H-H_2O]^+$ | 1268.5371 | 1268.5354 | 1.3 |
| $b9^+$ | 1121.4687 | 1121.4667 | 1.8 |
| $b8^+$ | 1034.4367 | 1034.4348 | 1.8 |
| $[y5\text{-ether}]^+$ | 685.2622 | 685.2604 | 2.6 |
| $y7^+$ | 1001.3788 | 1001.3771 | 1.7 |
| $y8^+$ | 1072.4159 | 1072.4139 | 1.9 |
| $y9^+$ | 1173.4636 | 1173.4616 | 1.7 |

**Figure S69: HR-MS/MS analysis of darobactin K5F. (A**) Structure of the modified species based on HR-MS/MS fragmentation (stereochemistry of linkages later determined by NMR spectroscopy). The modification is highlighted in orange. (B) CID spectrum of the +28 Da modified variant. The sequence of the GluC-digested peptide is provided with the substitution bolded. Orange colored daughter ions include the +28 Da modification. Distinguishing ether crosslink fragments are indicated. (C) Observed masses and errors from CID analysis.

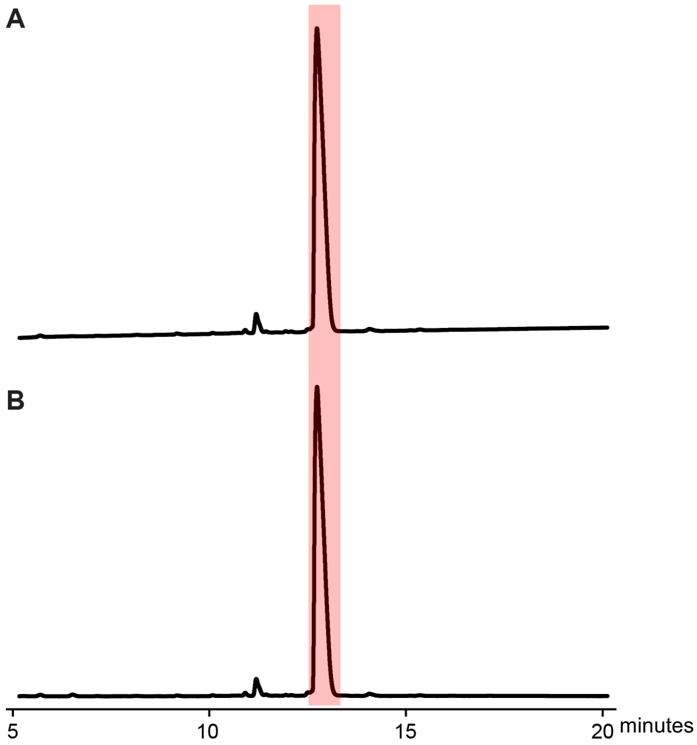

**Figure S70: UHPLC purification of darobactin K5F.** Chromatograms with the (A) 220 nm and (B) 280 nm absorbances are shown from the PFP analytical purification. The light red-colored fraction that eluted at 13 min contained darobactin K5F and was collected for NMR.

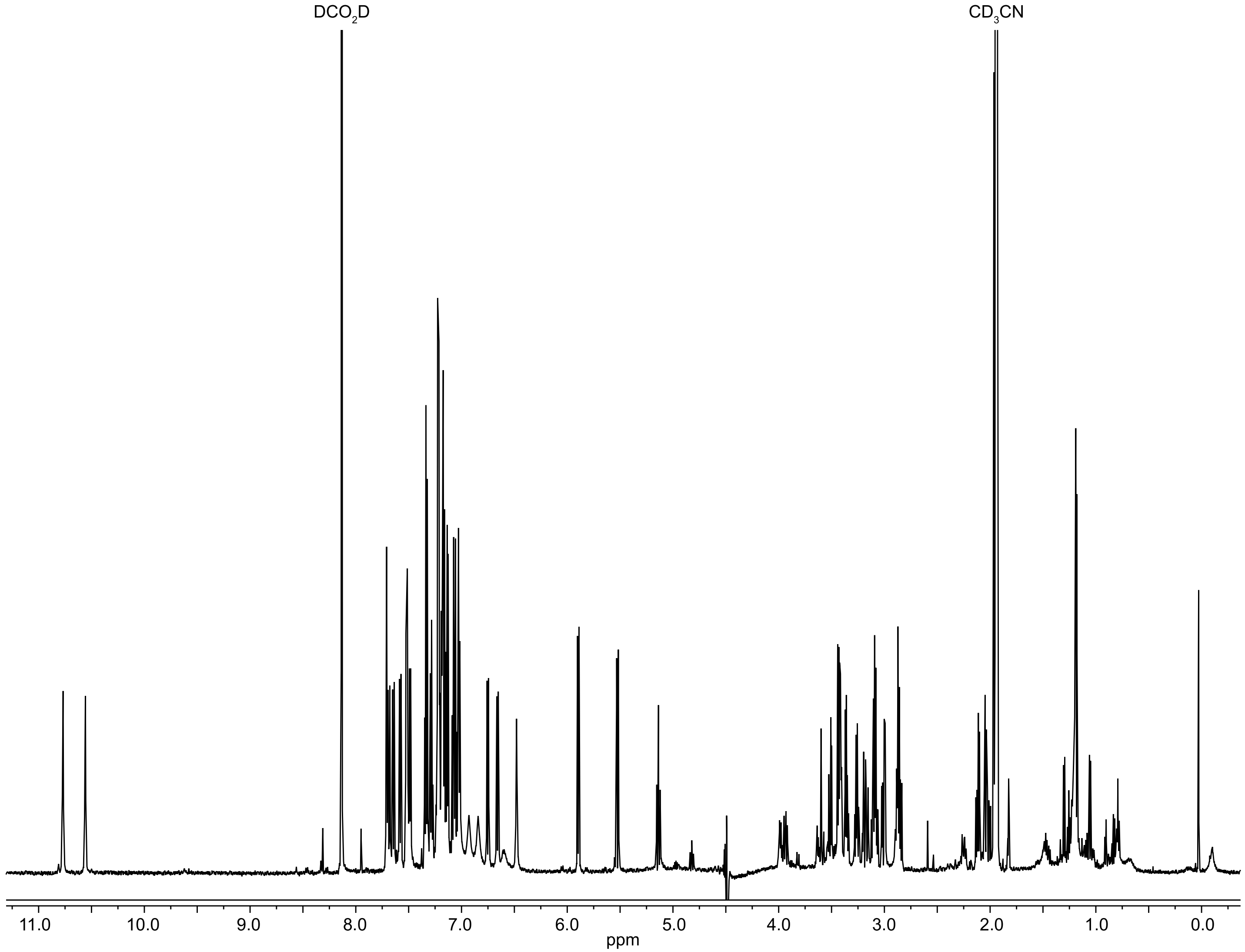

**Figure S71: $^1$H NMR spectrum of darobactin K5F.** Data were acquired in 66% $H_2O$: 32% $CD_3CN$: 2% $DCO_2D$ using a water-suppression (WET) method.

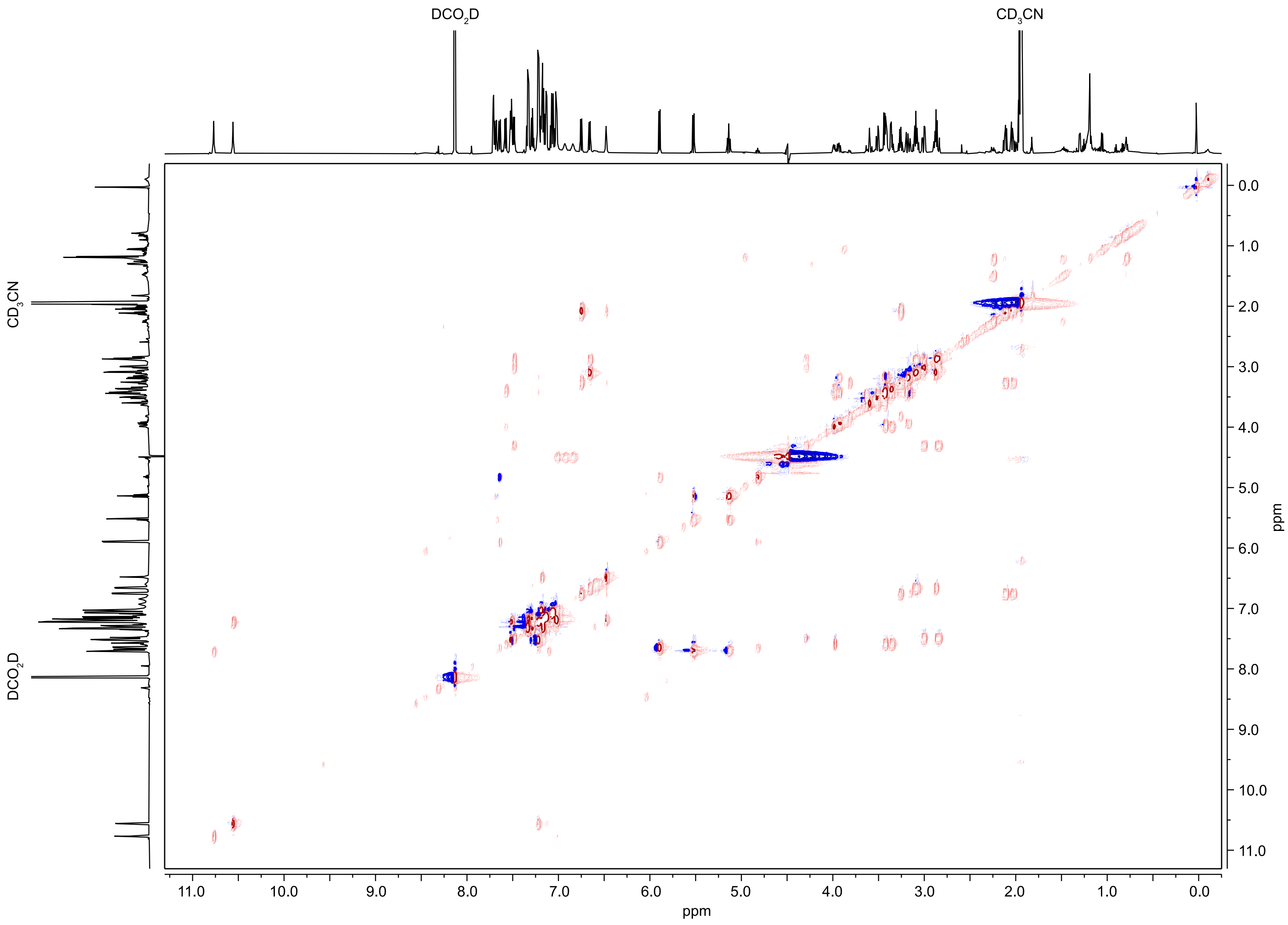

**Figure S72: [1]H-[1]H TOCSY NMR spectrum of darobactin K5F.** Data were acquired in 66% $H_2O$: 32% $CD_3CN$: 2% $DCO_2D$ using a water-suppression (WET) method.

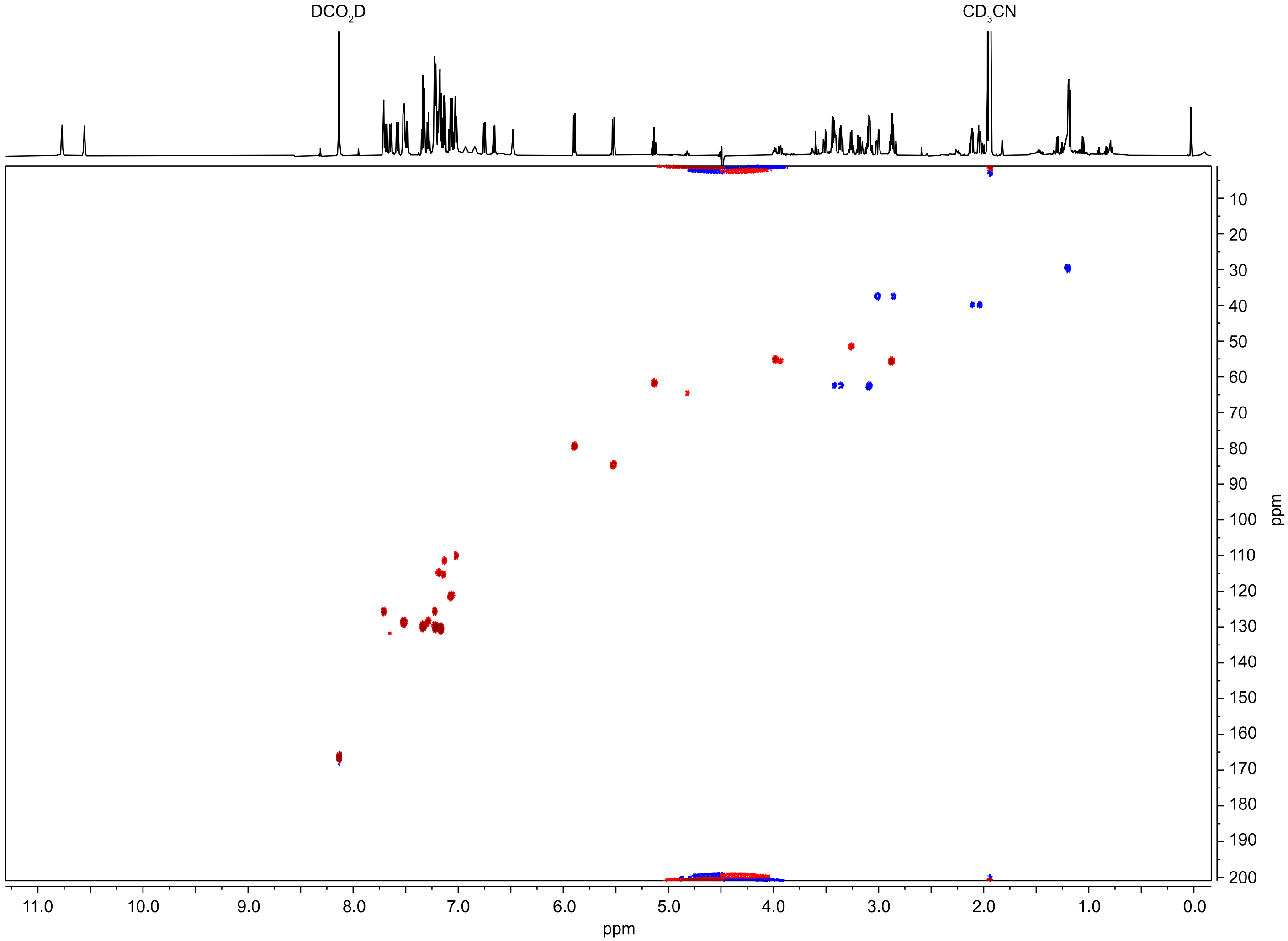

**Figure S73: $^{1}H$-$^{13}C$ HSQC NMR spectrum of darobactin K5F.** Data were acquired in 66% $H_2O$: 32% $CD_3CN$: 2% $DCO_2D$ using a water-suppression (WET) method.

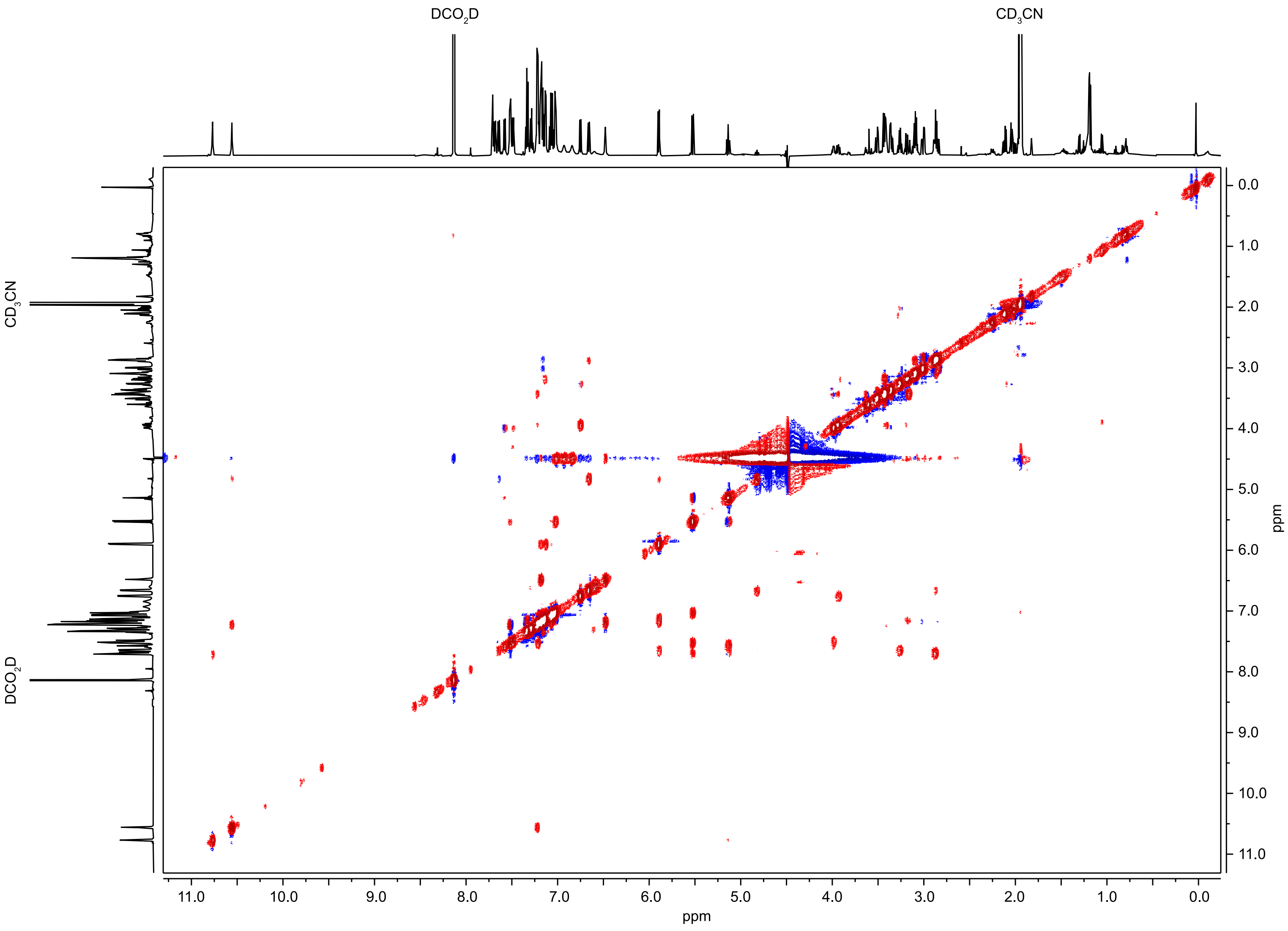

**Figure S74: [1]H-[1]H NOESY NMR spectrum of darobactin K5F.** Data were acquired in 66% $H_2O$: 32% $CD_3CN$: 2% $DCO_2D$ using a water-suppression (WET) method.

**A**

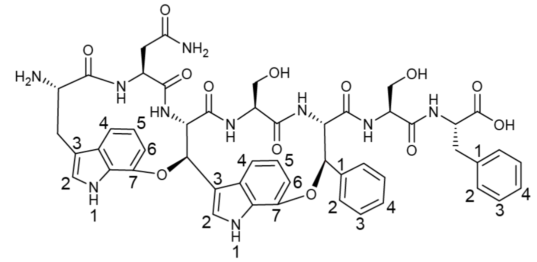

**B**

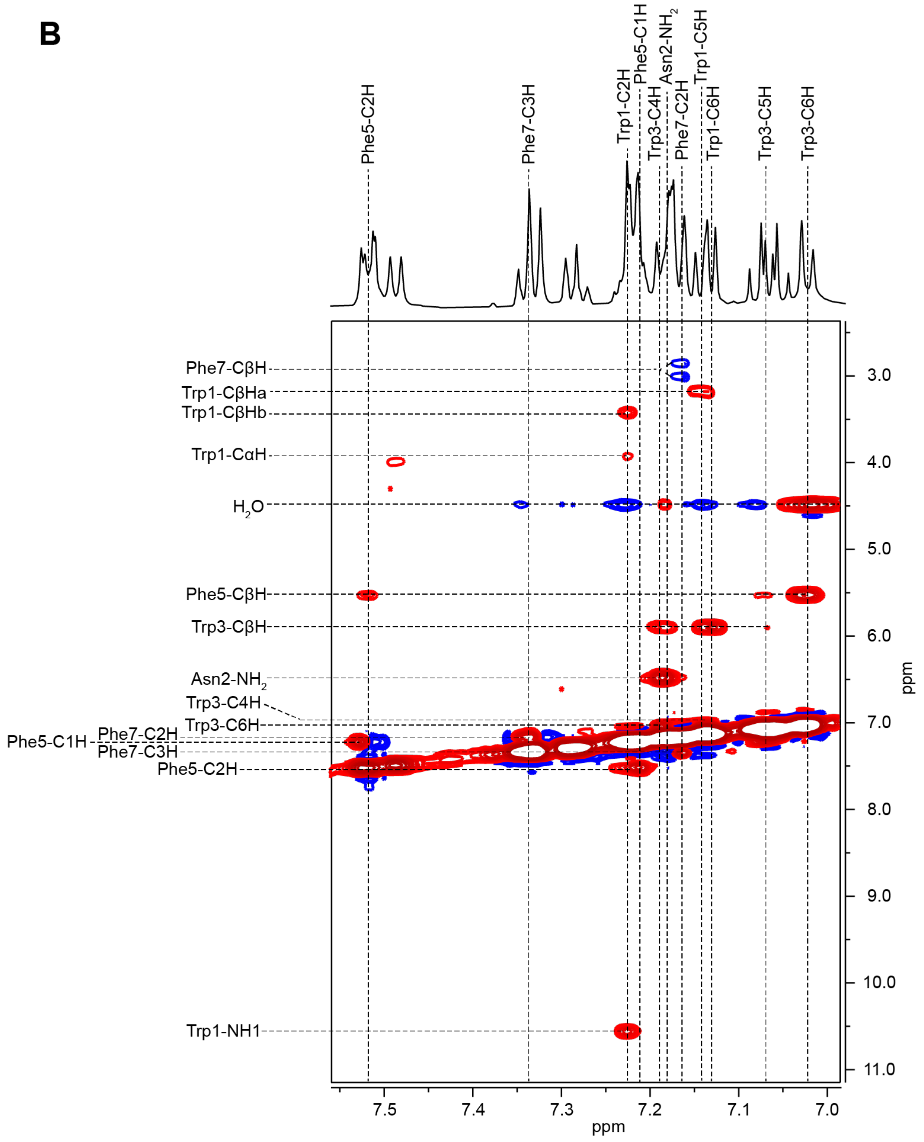

**Figure S75: Additional NMR information for darobactin K5F.**

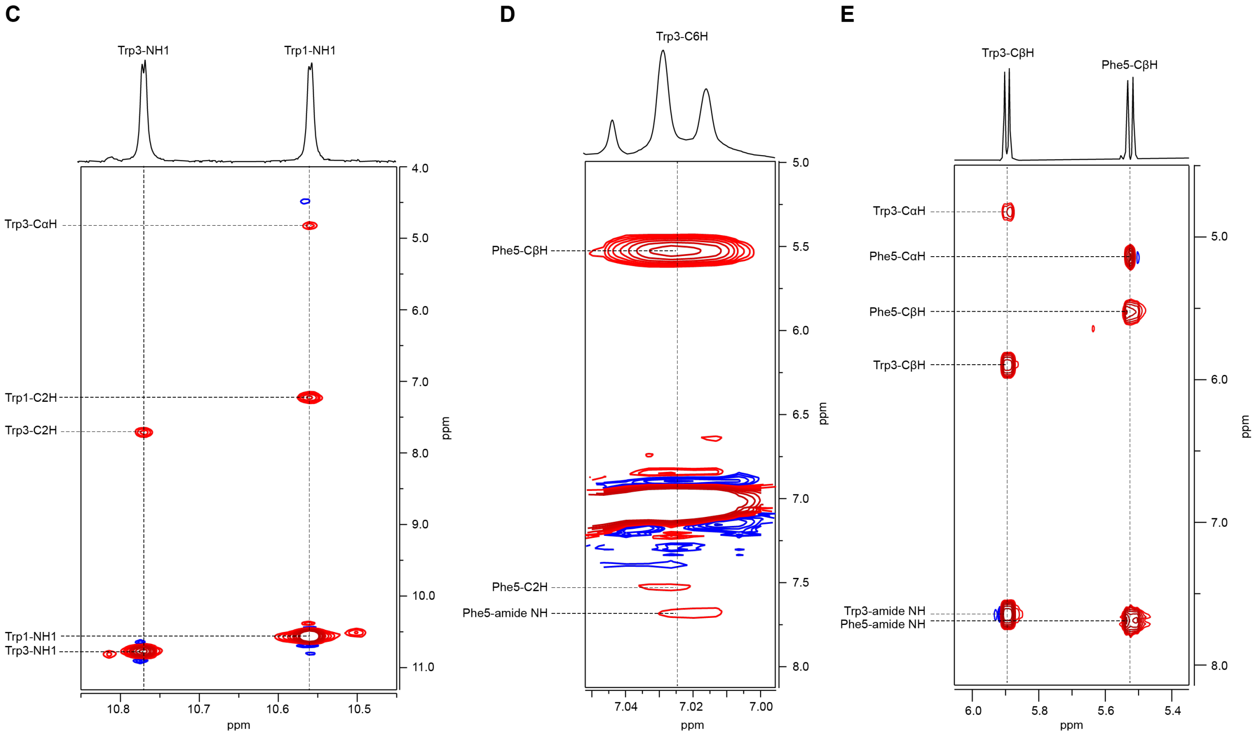

**Figure S75 cont.: Additional NMR information for darobactin K5F.** (A) Numbered darobactin K5F Structure. The numbering of atoms directly corresponds to those in **Table S4.** (B) $^{1}H$-$^{1}H$ NOESY spectrum of the aromatic region. (C) $^{1}H$-$^{1}H$ NOESY correlations for the Trp1- and Trp3-indole NHs. (D) $^{1}H$-$^{1}H$ NOSEY spectrum for Trp3-C6H to show key correlations that helped to establish the relative stereochemistry of the second macrocycle (Trp3-Phe5). (E) $^{1}H$-$^{1}H$ TOCSY correlations for Trp3 and Phe5 with a focus on the CβH for Trp3 and Phe5.

**Table S5: NMR chemical shift assignments for darobactin K5F.** Data were acquired in 66% $H_2O$: 32% $CD_3CN$: 2% $DCO_2D$ using a water-suppression (WET) method.

| Atom | $\delta_H$, mult. ($J$ in Hz) | $\delta_C$ | | Atom | $\delta_H$, mult. ($J$ in Hz) | $\delta_C$ |
|---|---|---|---|---|---|---|
| Trp 1 | | | | Ser 4 | | |
| Amide NH | - | - | | Amide NH | 6.66, d (8.0) | NH |
| α | 3.93, dd (11.3,7.7) | 55.2 (CH) | | α | 2.88, m | 55.2 (CH) |
| $\beta_a$ | 3.18, dd (13.9, 11.1) | 62.1 ($CH_2$) | | β | 3.09, t (6.7) | 62.2 ($CH_2$) |
| $\beta_b$ | 3.42, m | 62.1 ($CH_2$) | | | | |
| 1 | 10.56, d (2.6) | NH | | Phe 5 | | |
| 2 | 7.22, m | 125.2 (CH) | | Amide NH | 7.69, d (9.8) | NH |
| 3 | - | * | | α | 5.14, t (9.7) | 61.3 (CH) |
| 4 | 7.03, m | 120.9 (CH) | | β | 5.52, d (9.5) | 84.2 (CH) |
| 5 | 7.14, m | 114.9 (CH) | | 1 | - | * |
| 6 | 7.13, d (5.7) | 111.1 (CH) | | 2 | 7.52, dd (7.6, 2.1) | 128.3 (CH) |
| 7 | - | * | | 3 | 7.22, m | 129.6 (CH) |
| | | | | 4 | 7.24, m | 129.3 (CH) |
| Asn 2 | | | | | | |
| Amide NH | 6.75, d (8.1) | NH | | Ser 6 | | |
| α | 3.26, q (7.3) | 51.2 (CH) | | Amide NH | 7.58, d (8.2) | NH |
| $\beta_a$ | 2.04, m | 39.5 ($CH_2$) | | α | 3.99, m | 54.8 (CH) |
| $\beta_b$ | 2.12, dd (14.1, 6.7) | 39.5 ($CH_2$) | | $\beta_a$ | 3.36, dd (11.7, 6.1) | 62.1 ($CH_2$) |
| $NH_1$ | 6.48, br s | NH | | $\beta_b$ | 3.42, m | 62.1 ($CH_2$) |
| $NH_2$ | 7.18, br s | NH | | | | |
| | | | | Phe 7 | | |
| Trp 3 | | | | Amide NH | 7.49, d (7.4) | NH |
| Amide NH | 7.64, d (9.1) | NH | | α | 4.3, overlapped | 55.1 (CH) |
| α | 4.82, t (9.2) | 64.2 (CH) | | $\beta_a$ | 2.86, m | 37.1 ($CH_2$) |
| β | 5.9, d (8.4) | 79.2 (CH) | | $\beta_b$ | 3.01, dd (14.2, 5.5) | 37.1 ($CH_2$) |
| 1 | 10.77, d (2.8) | NH | | 1 | - | * |
| 2 | 7.71, d (2.7) | 125.3 (CH) | | 2 | 7.17, m | 130.1 (CH) |
| 3 | - | * | | 3 | 7.33, t (7.2 Hz) | 129.4 (CH) |
| 4 | 7.19, m | 114.4 (CH) | | 4 | 7.28, t (7.4 Hz) | 128.0 (CH) |
| 5 | 7.06, dd (7.9, 3.0) | 120.9 (CH) | | | | |
| 6 | 7.03, d (7.7) | 109.8 (CH) | | | | |
| 7 | - | * | | | | |

* No $^{13}C$ spectrum was recorded, thus, no $\delta_C$ was able to be assigned for the indicated carbon atoms.

A

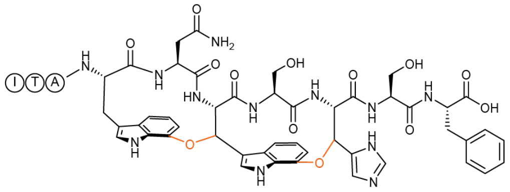

B

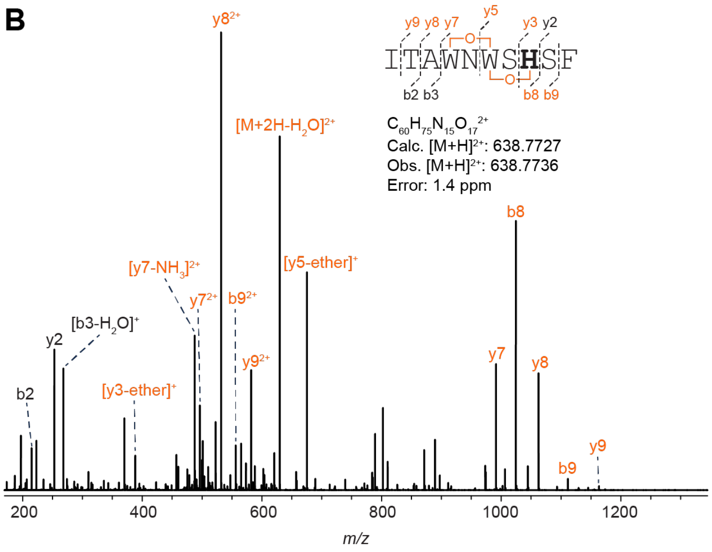

C

| Ion | Calculated Mass | Observed Mass | Error (ppm) |
|---|---|---|---|
| [M+2H]$^{2+}$ | 638.7727 | 638.7736 | 1.4 |
| [M+2H-H$_2$O]$^{2+}$ | 629.7675 | 629.7682 | 1.1 |
| b9$^+$ | 1111.4592 | 1111.4602 | 0.9 |
| b9$^{2+}$ | 556.2333 | 556.234 | 1.3 |
| y2$^+$ | 253.1183 | 253.1186 | 1.2 |
| b8$^+$ | 1024.4272 | 1024.428 | 0.8 |
| b8$^{2+}$ | 512.7173 | 512.7178 | 1.0 |
| [y3-ether]$^+$ | 388.1621 | 388.1622 | 0.3 |
| [y5-ether]$^+$ | 675.2527 | 675.2529 | 0.3 |
| b3$^+$ | 286.1762 | 286.1766 | 1.4 |
| [b3-H$_2$O]$^+$ | 268.1656 | 268.166 | 1.5 |
| y7$^+$ | 991.3692 | 991.3701 | 0.9 |
| y7$^{2+}$ | 496.1883 | 496.1890 | 1.4 |
| [y7-NH$_3$]$^{2+}$ | 487.6750 | 487.6756 | 1.2 |
| b2$^+$ | 215.1391 | 215.1393 | 0.9 |
| y8$^+$ | 1062.4063 | 1062.4074 | 1.0 |
| y8$^{2+}$ | 531.7069 | 531.7075 | 1.1 |
| y9$^+$ | 1163.4540 | 1163.4551 | 0.9 |
| y9$^{2+}$ | 582.2307 | 582.2315 | 1.4 |

**Figure S76: HR-MS/MS analysis of darobactin K5H.** (A) Hypothetical structure of the modified species based on HR-MS/MS fragmentation. The connectivity of the linkage is presumed from the known structure of darobactin. The modification is highlighted in orange. (B) CID spectrum of the +28 Da modified variant. The sequence of the GluC-digested peptide is provided with the substitution bolded. Orange colored daughter ions include the modification. Distinguishing ether crosslink fragments are indicated. (C) Observed masses and errors from CID analysis.

A

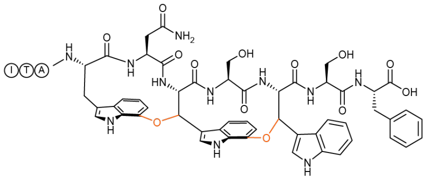

D

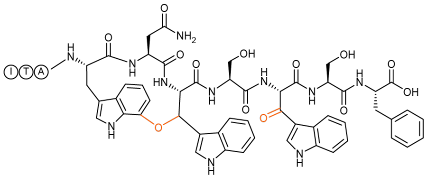

B

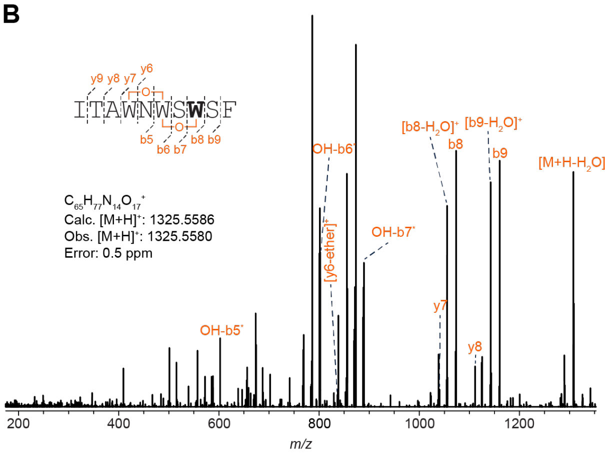

E

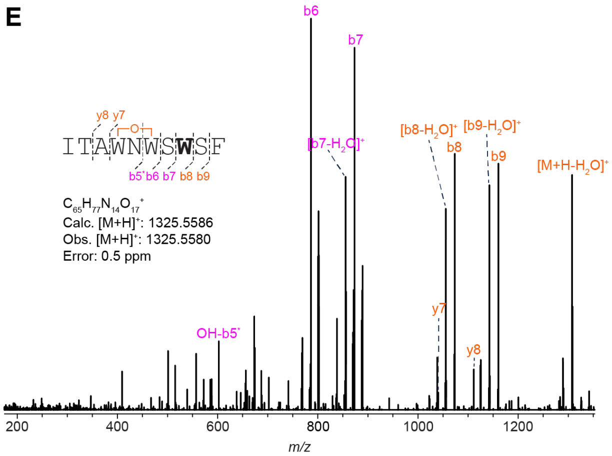

C

| Ion | Calculated Mass | Observed Mass | Error (ppm) |
|---|---|---|---|
| $[M+H]^+$ | 1325.5586 | 1325.5580 | 0.5 |
| $[M+H-H_2O]^+$ | 1307.5479 | 1307.5483 | 0.3 |
| $b9^+$ | 1160.4796 | 1160.4763 | 2.8 |
| $[b9-H_2O]^+$ | 1142.4690 | 1142.4674 | 1.4 |
| $b8^+$ | 1073.4476 | 1073.4465 | 1.0 |
| $[b8-H_2O]^+$ | 1055.4370 | 1055.4359 | 1.0 |
| $OH\text{-}b7^*$ | 889.3840 | 889.3810 | 3.4 |
| $OH\text{-}b6^*$ | 802.3519 | 802.3514 | 0.6 |
| $OH\text{-}b5^*$ | 602.2933 | 602.2943 | 1.7 |
| $[y6\text{-ether}]^+$ | 838.3156 | 838.3177 | 2.5 |
| $y7^+$ | 1040.3898 | 1040.3914 | 1.5 |
| $y8^+$ | 1111.4269 | 1111.4265 | 0.4 |

F

| Ion | Calculated Mass | Observed Mass | Error (ppm) |
|---|---|---|---|
| $[M+H]^+$ | 1325.5586 | 1325.558 | 0.5 |
| $[M+H-H_2O]^+$ | 1307.5479 | 1307.5483 | 0.3 |
| $b9^+$ | 1160.4795 | 1160.4764 | 2.7 |
| $[b9-H_2O]^+$ | 1142.4690 | 1142.4674 | 1.4 |
| $b8^+$ | 1073.4475 | 1073.4465 | 0.9 |
| $[b8-H_2O]^+$ | 1055.4370 | 1055.4359 | 1.0 |
| $b7^+$ | 873.3890 | 873.3903 | 1.5 |
| $[b7-H_2O]^+$ | 855.3784 | 855.3794 | 1.2 |
| $b6^+$ | 786.3569 | 786.3585 | 2.0 |
| $b5+OH^*$ | 602.2938 | 602.2945 | 1.2 |
| $y7^+$ | 1040.3896 | 1040.3896 | 0.0 |
| $y8^+$ | 1111.4267 | 1111.4267 | 0.0 |

**Figure S77: HR-MS/MS analysis of the DarA-K5W variant with +28 Da modifications.** (A) Hypothetical structure of GluC-digested DarA-K5W variant with a fused diether structure. The connectivity of the linkage is presumed from the known structure of darobactin. The modifications are highlighted in orange. (B) CID spectrum of the diether-containing variant. The sequence of the GluC-digested peptide is provided with the substitution bolded. Orange colored daughter ions include the modification. Distinguishing ether crosslink fragments are indicated. (C) Observed masses and errors from panel B. (D) Hypothetical structure of the DarA-K5W variant with one ether crosslink and β-oxotryptophan. (E) CID spectrum of the monoether and β-oxo-modified variant. Distinguishing ether crosslink fragments are indicated. Magenta ions are +14 Da while orange ions are +28 Da relative to the unmodified peptide. (F) Observed masses and errors from panel E.

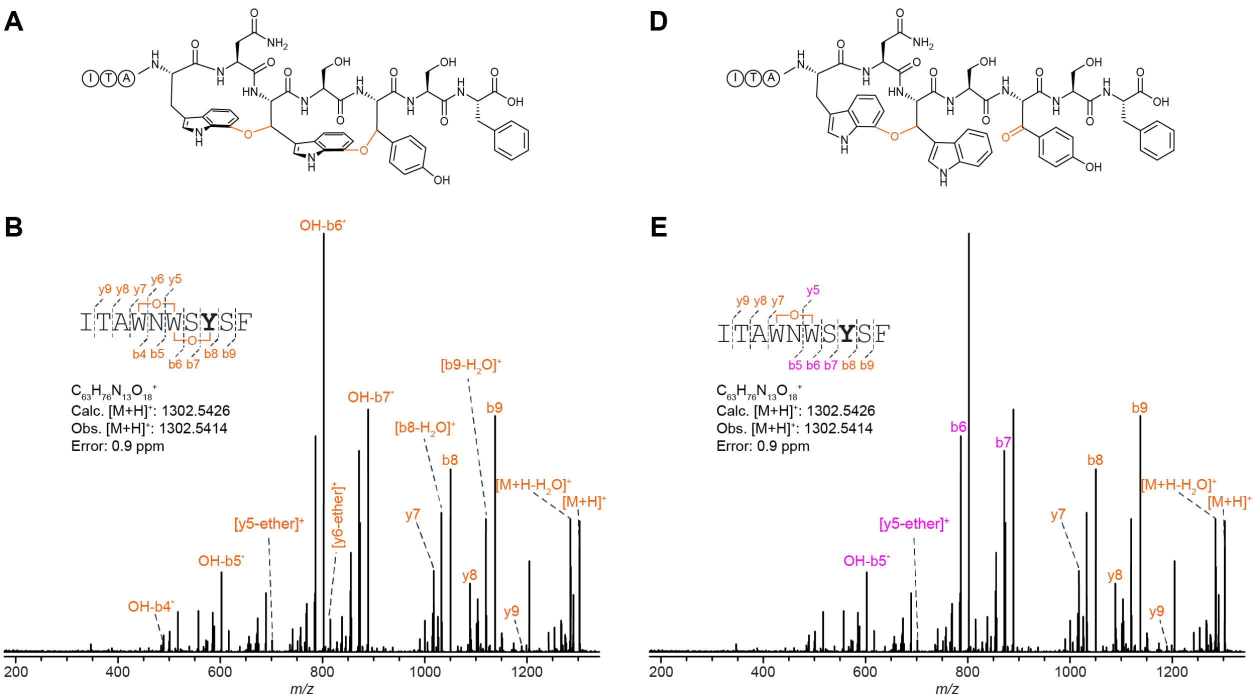

C

| Ion | Calculated Mass | Observed Mass | Error (ppm) |
|---|---|---|---|
| $[M+H]^+$ | 1302.5426 | 1302.5414 | 0.9 |
| $[M+H-H_2O]^+$ | 1284.5321 | 1284.5291 | 2.3 |
| $b9^+$ | 1137.4637 | 1137.4610 | 2.4 |
| $[b9-H_2O]^+$ | 1119.4531 | 1119.4506 | 2.2 |
| $b8^+$ | 1050.4316 | 1050.4294 | 2.1 |
| $[b8-H_2O]^+$ | 1032.4210 | 1032.4189 | 2.0 |
| $OH\text{-}b7^*$ | 889.3840 | 889.3820 | 2.2 |
| $OH\text{-}b6^*$ | 802.3519 | 802.3503 | 2.0 |
| $OH\text{-}b5^*$ | 602.2933 | 602.2920 | 2.2 |
| $[y5\text{-ether}]^+$ | 701.2566 | 701.2557 | 1.3 |
| $OH\text{-}b4^*$ | 488.2504 | 488.2495 | 1.8 |
| $[y6\text{-ether}]^+$ | 815.3000 | 815.2981 | 2.3 |
| $y7^+$ | 1017.3738 | 1017.3717 | 2.1 |
| $y8^+$ | 1088.4109 | 1088.4086 | 2.1 |
| $y9^+$ | 1189.4586 | 1189.4559 | 2.3 |

F

| Ion | Calculated Mass | Observed Mass | Error (ppm) |
|---|---|---|---|
| $[M+H]^+$ | 1302.5426 | 1302.5414 | 0.9 |
| $[M+H-H_2O]^+$ | 1284.5321 | 1284.5291 | 2.3 |
| $b9^+$ | 1137.4636 | 1137.4611 | 2.2 |
| $b8^+$ | 1050.4316 | 1050.4294 | 2.1 |
| $b7^+$ | 873.3890 | 873.3873 | 1.9 |
| $b6^+$ | 786.3570 | 786.3555 | 1.9 |
| $[y5\text{-ether}]^+$ | 701.2571 | 701.2566 | 0.7 |
| $y7^+$ | 1017.3738 | 1017.3719 | 1.9 |
| $y8^+$ | 1088.4109 | 1088.4087 | 2.0 |
| $y9^+$ | 1189.4586 | 1189.4565 | 1.8 |

**Figure S78: HR-MS/MS analysis of the DarA-K5Y variant with +28 Da modifications.** (A) Hypothetical structure of the DarA-K5Y variant with a fused diether structure. The connectivity of the linkage is presumed from the known structure of darobactin. The modifications are highlighted in orange. (B) CID spectrum of the diether-containing variant. The sequence of the GluC-digested peptide is provided with the substitution bolded. Orange colored daughter ions include the modification. Distinguishing ether crosslink fragments are indicated. (C) Observed masses and errors from panel B. (D) Hypothetical structure of the DarA-K5Y variant with one ether crosslink and β-oxotyrosine. The connectivity of the linkage is presumed from the known structure of darobactin. The modification is highlighted in orange. (E) CID spectrum of the monoether and β-oxo-modified variant. Distinguishing ether crosslink fragments are indicated. Magenta ions are +14 Da while orange ions are +28 Da relative to the unmodified peptide. (F) Observed masses and errors from panel E.

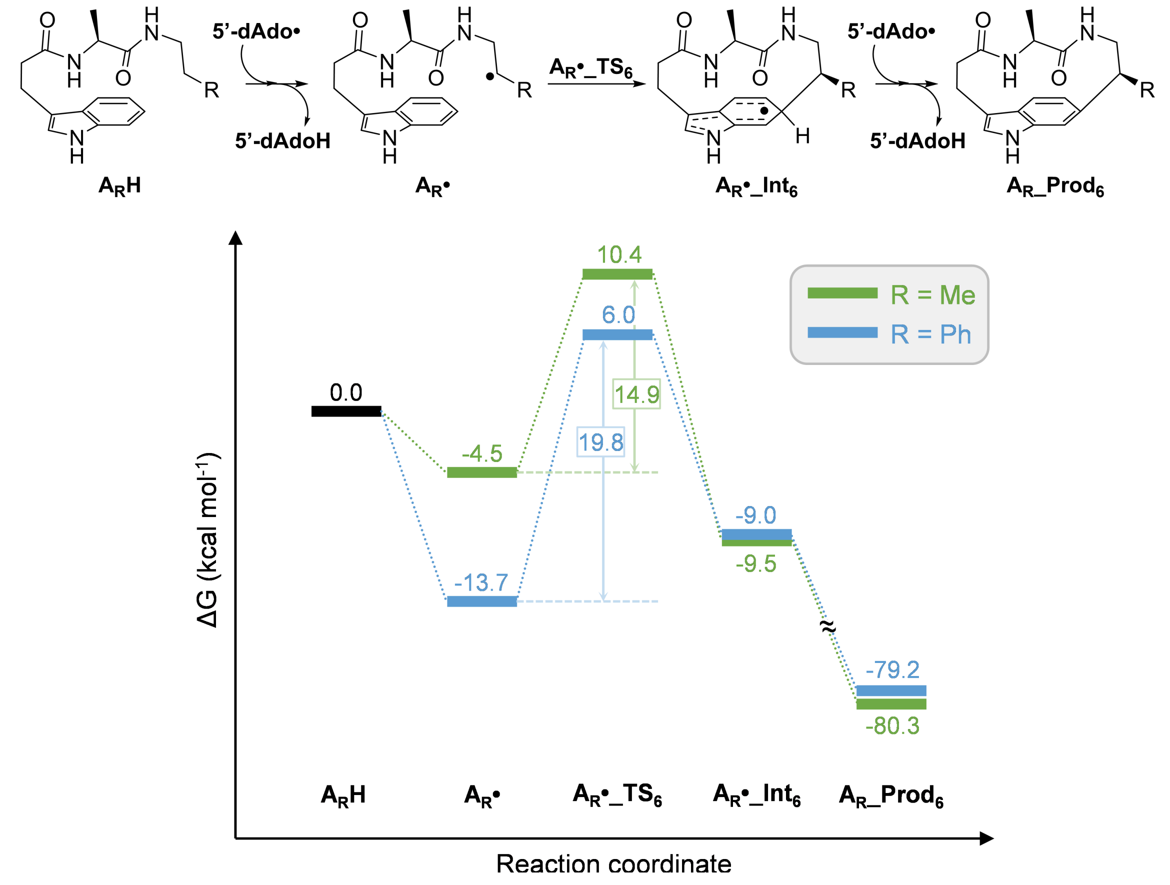

**Figure S79: Energy landscape for the formation of C-centered radicals (A•).** Proposed mechanism and minimum energy pathway calculated with PCM($H_2O$)/ωB97X-D/6-311G(d,p) for the formation of C-centered radicals (**A•**) and the subsequent addition to Trp-C6.

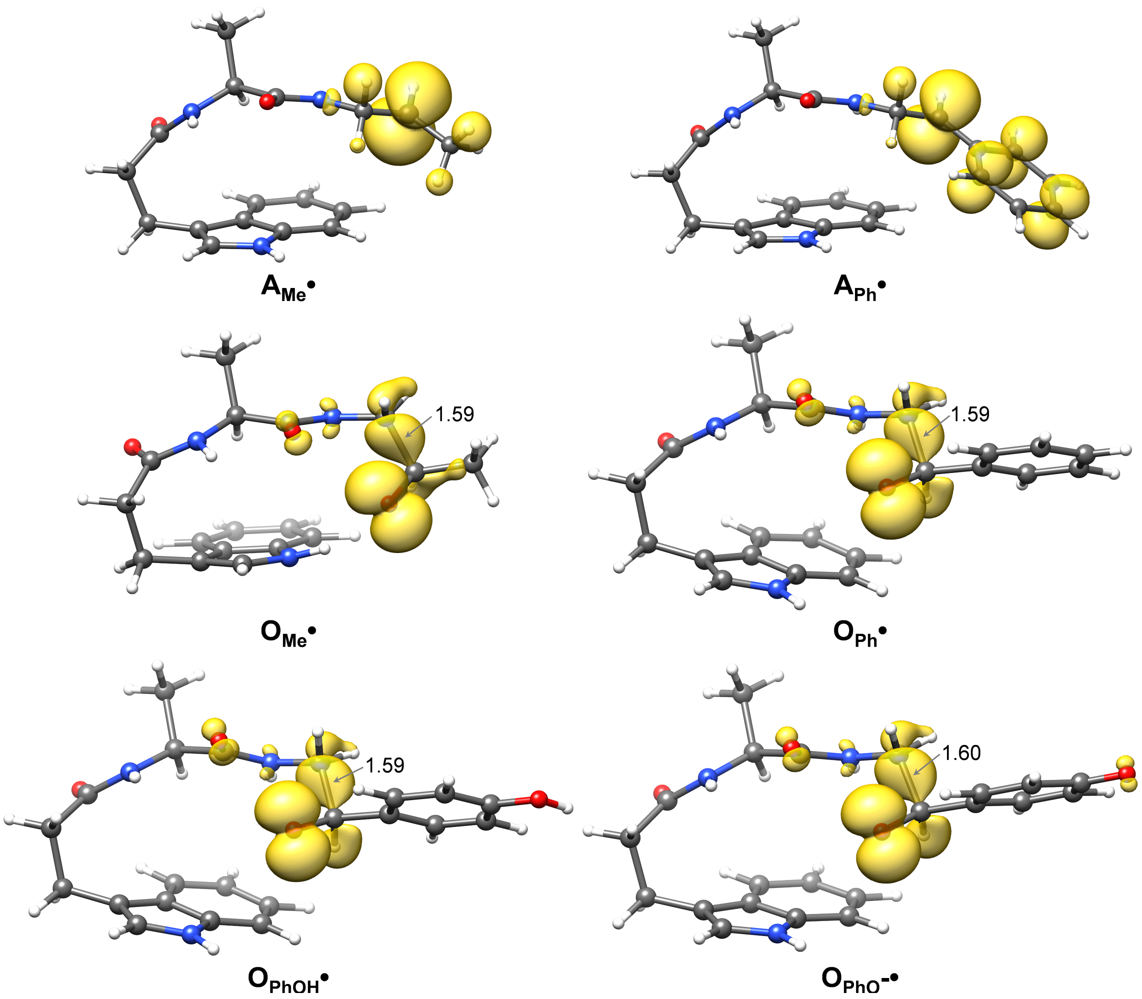

**Figure S80: Spin density isosurfaces for C-centered ($A_R$•) and O-centered radicals ($O_R$•).** Isosurfaces were plotted at an isovalue of 0.003 and calculated with PCM($H_2O$)/ωB97X-D/6-311G(d,p) theory.

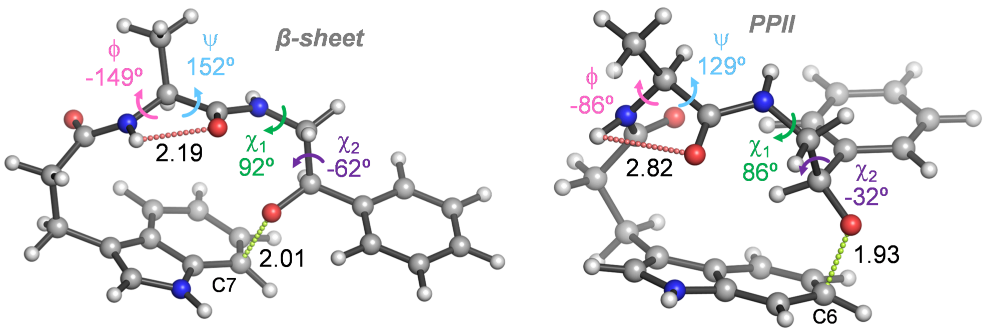

| | $O_{Ph}$•_$TS_7$ | $O_{Ph}$•_$TS_{6a}$ |
|---|---|---|
| ΔG‡ | 8.3 | 7.9 |

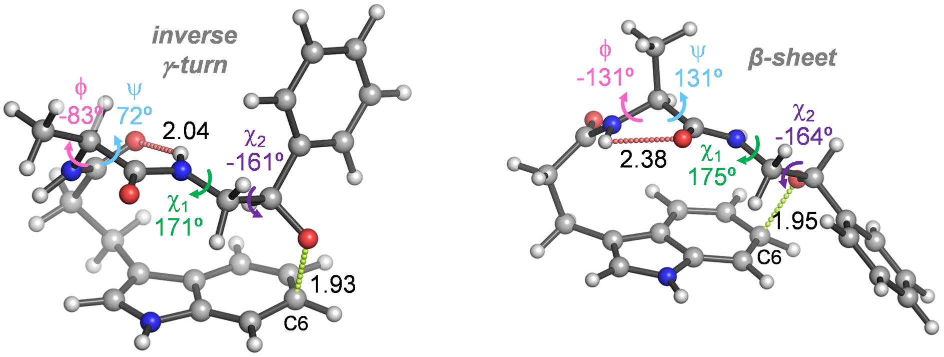

| | $O_{Ph}$•_$TS_{6b}$ | $O_{Ph}$•_$TS_{6c}$ |
|---|---|---|
| ΔG‡ | 9.7 | 9.7 |

**Figure S81: Computational analysis of indole addition by an O-centered radical.** Comparison of the transition state structures calculated with PCM($H_2O$)/ωB97X-D/6-311G(d,p) for the addition of O-centered radicals **$O_{Ph}$•** to the C7 (**$O_{Ph}$•_$TS_7$**) or C6 (**$O_{Ph}$•_$TS_{6a-c}$**) positions of indole. Attack to C6 leads to more folded conformations (as judged from the torsional ϕ and ψ angles, shown in pink and blue, respectively) and to slightly higher activation free energies (ΔG‡ in kcal $mol^{-1}$). Distances of the forming C-O bonds are shown as green dashed lines. Interactions between backbone CO and NH groups are shown as red dashed lines. Distances are given in angstroms.

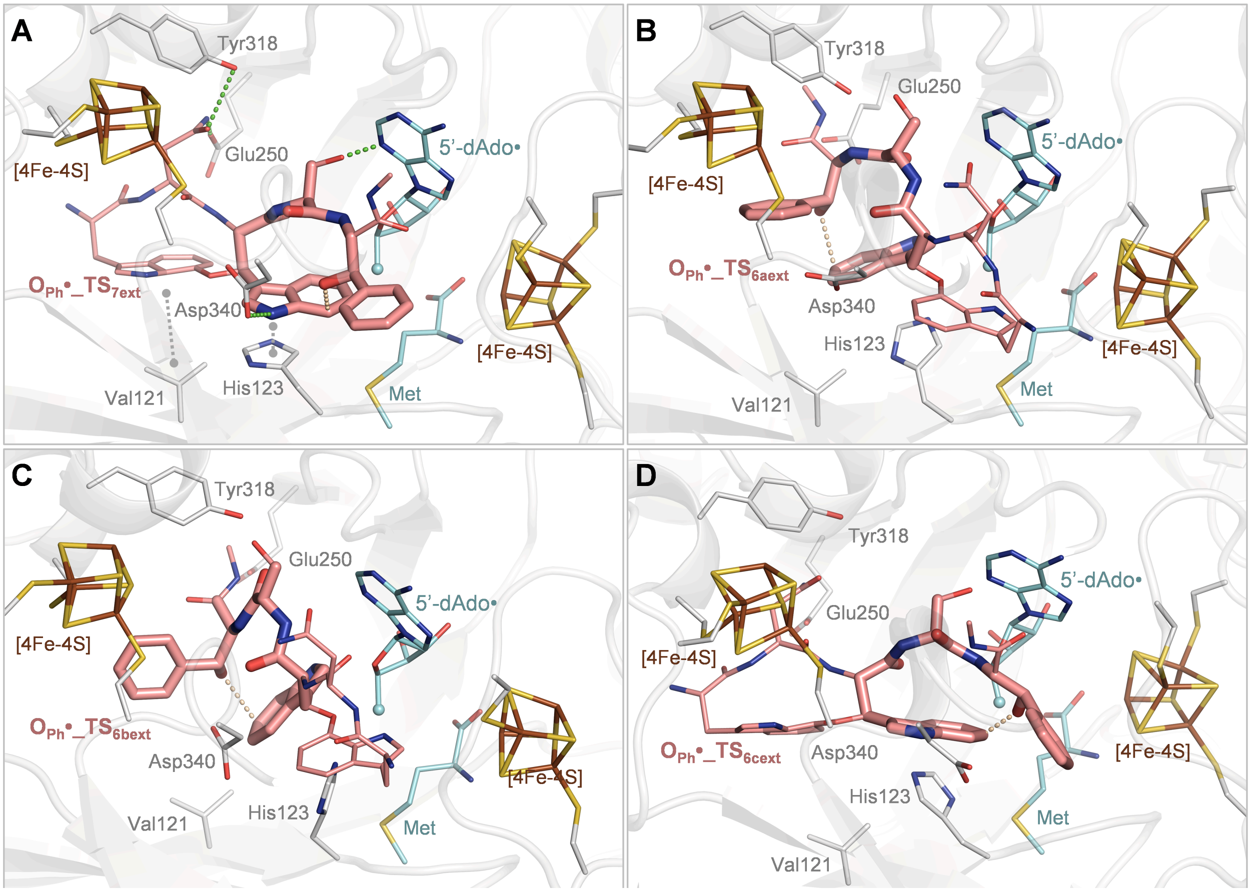

**Figure S82: Molecular docking structures of truncated models of darobactin.** Darobactin is shown as a pink wireframe. The geometries of QM-optimized transition states for ether crosslinking between residues Trp3 and Phe5 at positions C7 (A) or C6 (B-C) of the indole ring (shown as pink sticks) are featured. The activated cofactor (in the form of 5'-Ado• and methionine) is shown in cyan (the reacting radical center is shown as a sphere). Iron (in brown)-sulfur (in yellow) clusters and relevant active site residues (in grey) are shown as a wireframe.

Transition state **OPh•_$TS_{7ext}$** binds to DarE with the highest score (23.96; panel A) and places the Cβ of Trp3 at less than 5 Å from the 5'-dAdo radical. This pose involves the TS entering the active site C-terminus first and is stabilized by multiple interactions, as described in the main text. A pose with a similar arrangement is found for **OPh•_$TS_{6cext}$** but with a significantly lower score (10.10; panel D); the reduced affinity arises from steric clashes involving the NHMe capping at residue Phe5. **OPh•_$TS_{6aext}$** and **OPh•_$TS_{6bext}$** cannot be accommodated at the active site with the same orientation, and all docking poses show the transition states entering the active site N-terminus first in a non-productive arrangement (panels B and C, respectively); moreover, these poses show lower scores than that of **OPh•_$TS_{7ext}$** (20.96 and 18.73, respectively). Overall, the predicted preference for the transition state model corresponding to ether crosslink at C7 of Trp3 indole is in line with the observed regioselectivity. Hydrogen bonds and van der Waals interactions are shown as green and grey dashed lines, respectively.

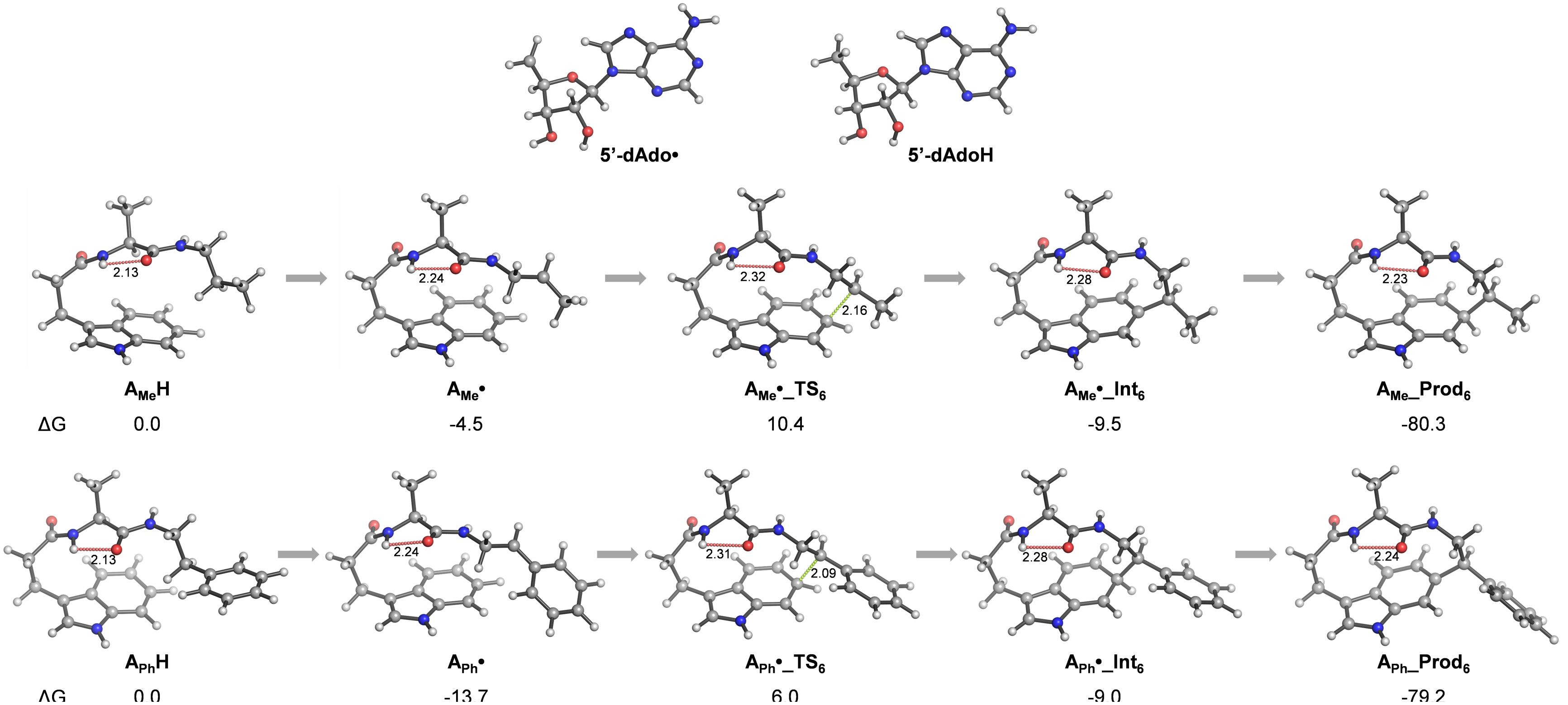

**Figure S83: Structures for the formation of C-centered radicals ($A_R$•) and addition toTrp-C6.** PCM($H_2O$)/ωB97X-D/6-311G(d,p) theory was used. Distances of the forming C-O bonds are shown as green dashed lines. Interactions between backbone CO and NH groups are shown as red dashed lines. Distances are given in angstroms. Free energies (ΔG) are given in kcal $mol^{-1}$.

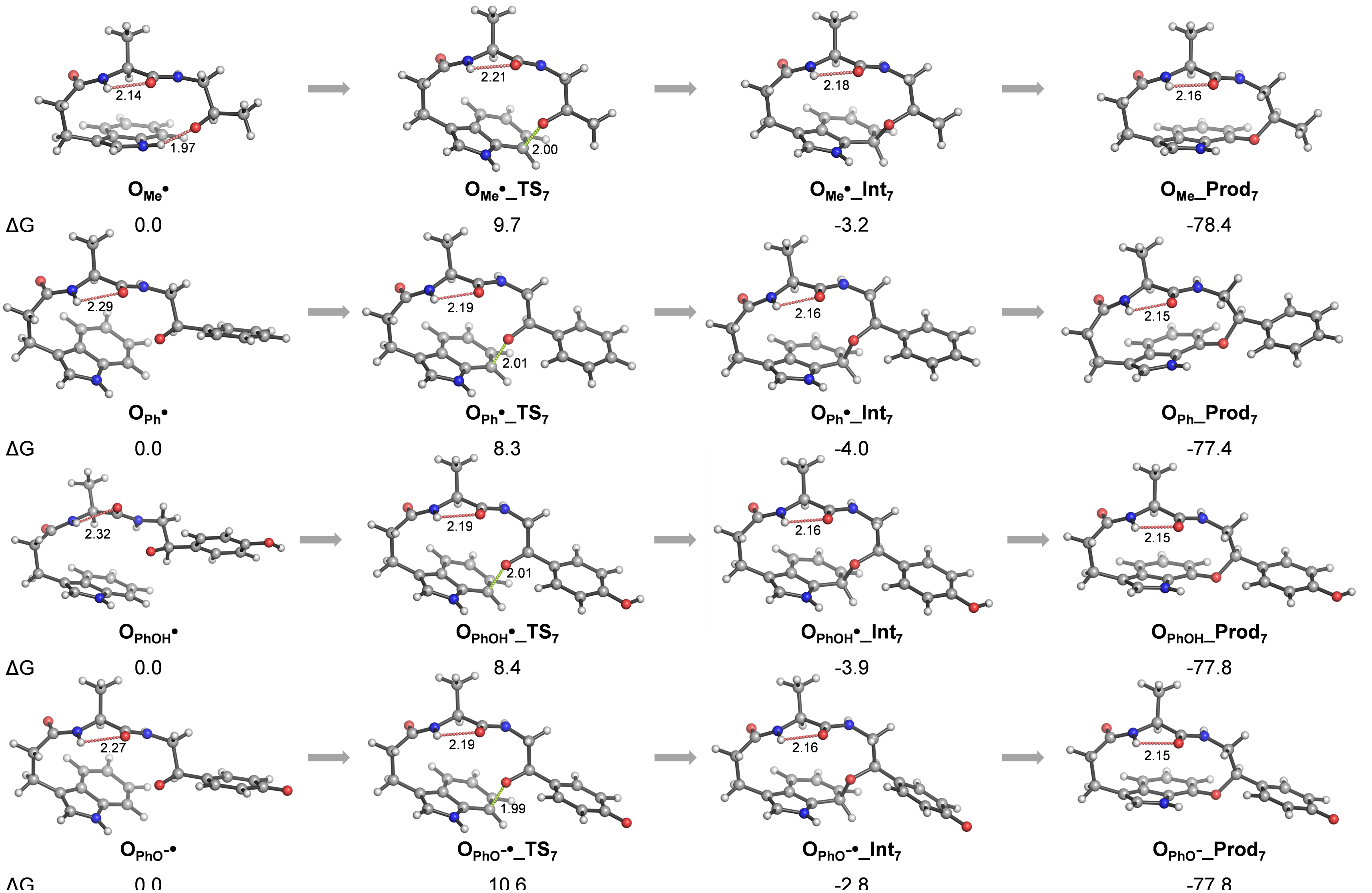

**Figure S84: Structures calculated with PCM($H_2O$)/ωB97X-D/6-311G(d,p) theory for the addition of O-centered radicals ($O_R$•) to Trp-C7.** Distances of the forming C-O bonds are shown as green dashed lines. Interactions between backbone CO and NH groups are shown as red dashed lines. Distances are given in angstroms. Free energies (ΔG) are given in kcal $mol^{-1}$.

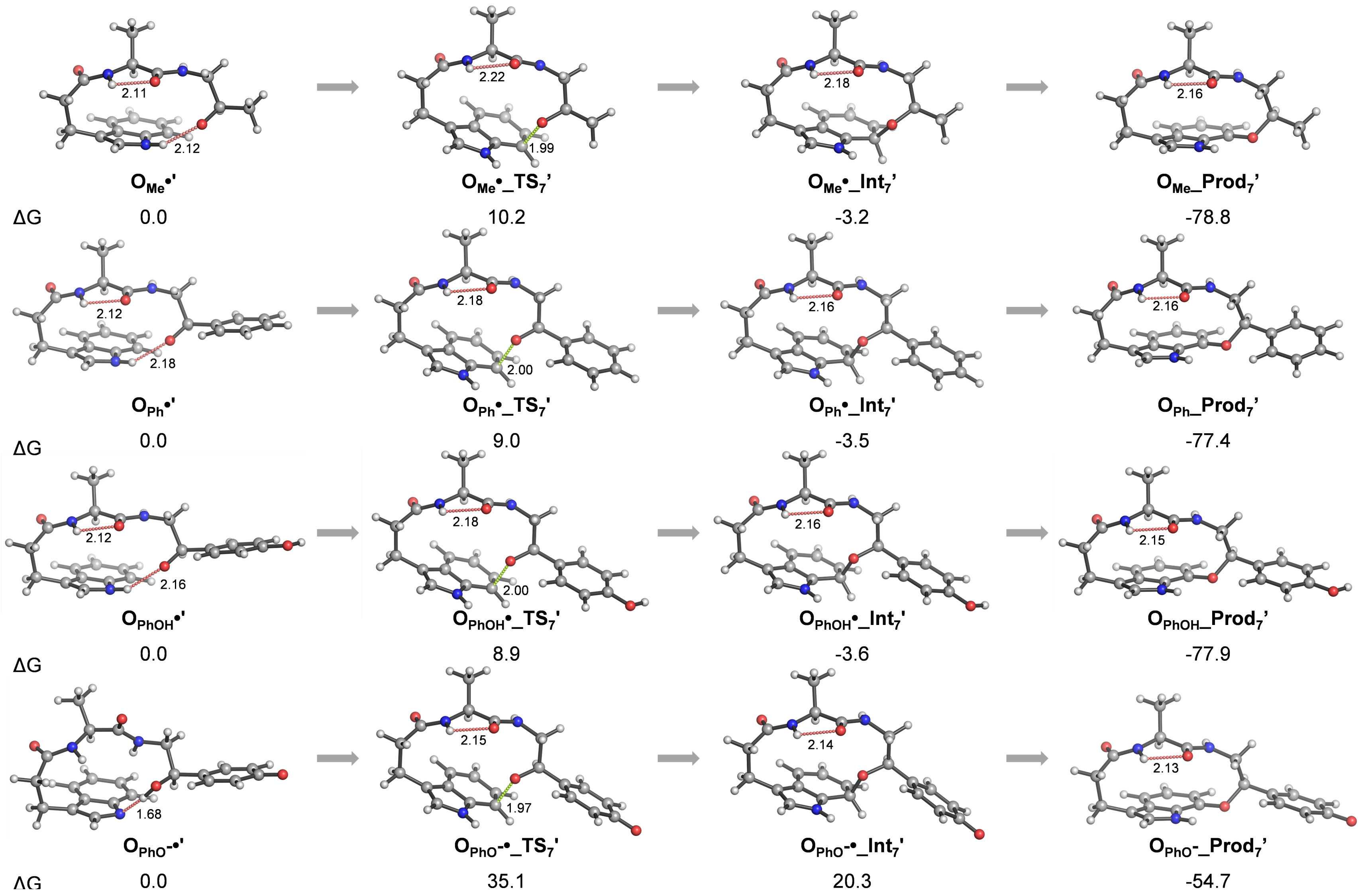

**Figure S85: Structures calculated with PCM($Et_2O$)/ωB97X-D/6-311G(d,p) theory for the addition of O-centered radicals ($O_R$•) to Trp-C7.** Distances of the forming C-O bonds are shown as green dashed lines. Interactions between backbone CO and NH groups are shown as red dashed lines. Distances are given in angstroms. Free energies (ΔG) are given in kcal $mol^{-1}$.

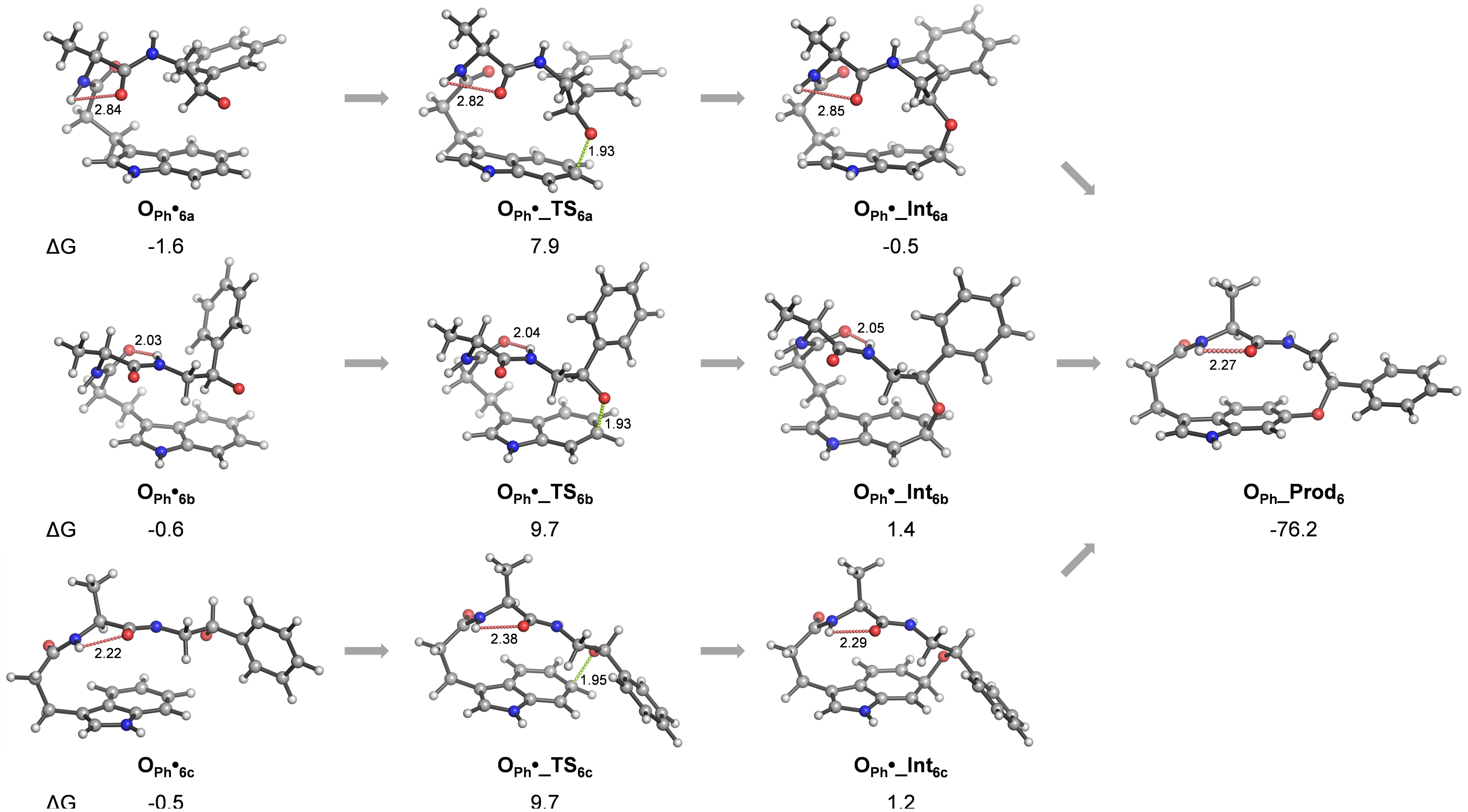

**Figure S86: Structures for the addition of O-centered radicals ($O_{Ph}•_{6a-c}$) to Trp-C6.** PCM($H_2O$)/ωB97X-D/6-311G(d,p) theory was used for the calculations. Free energies are arbitrarily referred to the oxy radical leading to addition to the C7 position of the indole (**$O_{Ph}$•**, Figure S90). Distances of the forming C-O bonds are shown as green dashed lines. Interactions between backbone CO and NH groups are shown as red dashed lines. Distances are given in angstroms. Free energies (ΔG) are given in kcal $mol^{-1}$.

**Table S6: Energies, entropies, and lowest frequencies of the lowest energy structures calculated at the PCM($H_2O$)/ωB97X-D/6-311G(d,p) level of theory.[a]**

| Structure | $E_{elec}$ (Hartree) | $E_{elec}$ + ZPE (Hartree) | H (Hartree) | S (cal $mol^{-1}$ $K^{-1}$) | G (Hartree) | Lowest freq. ($cm^{-1}$) | # imag. freq. |
|---|---|---|---|---|---|---|---|
| **5'-dAdo•** | –887.618287 | –887.385299 | –887.367957 | 134.2 | –887.427814 | 18.2 | 0 |
| **5'-dAdoH** | –888.301243 | –888.051887 | –888.035671 | 124.4 | –888.092649 | 33.9 | 0 |
| **$A_{Me}$H** | –976.390326 | –976.004198 | –975.981632 | 156.7 | –976.051350 | 15.1 | 0 |
| **$A_{Me}$•** | –975.721019 | –975.349871 | –975.327024 | 157.6 | –975.398014 | 30.4 | 0 |
| **$A_{Me}$•_TS$_6$** | –975.700322 | –975.328078 | –975.306906 | 146.4 | –975.374279 | –588.6 | 1 |
| **$A_{Me}$•_Int$_6$** | –975.735544 | –975.360381 | –975.339516 | 144.2 | –975.406007 | 37.3 | 0 |
| **$A_{Me}$_Prod$_6$** | –975.180063 | –974.813837 | –974.793645 | 138.9 | –974.858266 | 48.3 | 0 |
| **$A_{Ph}$H** | –1168.103996 | –1167.664048 | –1167.638502 | 175.7 | –1167.713994 | 9.6 | 0 |
| **$A_{Ph}$•** | –1167.451142 | –1167.025101 | –1166.999691 | 172.2 | –1167.075412 | 21.5 | 0 |
| **$A_{Ph}$•_TS$_6$** | –1167.421480 | –1166.994764 | –1166.970673 | 162.3 | –1167.043861 | –656.1 | 1 |
| **$A_{Ph}$•_Int$_6$** | –1167.447483 | –1167.018936 | –1166.994833 | 162.0 | –1167.067843 | 34.3 | 0 |
| **$A_{Ph}$_Prod$_6$** | –1166.891292 | –1166.471572 | –1166.448115 | 158.4 | –1166.519235 | 18.9 | 0 |
| **$O_{Me}$•** | –1050.933899 | –1050.556278 | –1050.533365 | 157.6 | –1050.604419 | 25.3 | 0 |
| **$O_{Me}$•_TS$_7$** | –1050.919877 | –1050.541863 | –1050.519891 | 151.4 | –1050.588955 | –418.6 | 1 |
| **$O_{Me}$•_Int$_7$** | –1050.942859 | –1050.562749 | –1050.540849 | 151.1 | –1050.609470 | 29.1 | 0 |
| **$O_{Me}$_Prod$_7$** | –1050.393663 | –1050.023206 | –1050.001896 | 144.5 | –1050.068809 | 50.2 | 0 |
| **$O_{Ph}$•** | –1242.644745 | –1242.213983 | –1242.187689 | 178.6 | –1242.265305 | 12.9 | 0 |
| **$O_{Ph}$•_TS$_7$** | –1242.633768 | –1242.201981 | –1242.176979 | 168.7 | –1242.252042 | –386.0 | 1 |
| **$O_{Ph}$•_Int$_7$** | –1242.655437 | –1242.221857 | –1242.196893 | 168.6 | –1242.271631 | 22.5 | 0 |
| **$O_{Ph}$_Prod$_7$** | –1242.103232 | –1241.679362 | –1241.654847 | 162.9 | –1241.728168 | 26.2 | 0 |
| **$O_{PhOH}$•** | –1317.870128 | –1317.435065 | –1317.407584 | 183.6 | –1317.487353 | 13.6 | 0 |
| **$O_{PhOH}$•_TS$_7$** | –1317.858784 | –1317.422924 | –1317.396618 | 175.7 | –1317.474044 | –393.4 | 1 |
| **$O_{PhOH}$•_Int$_7$** | –1317.880710 | –1317.442795 | –1317.416600 | 176.1 | –1317.493573 | 7.5 | 0 |
| **$O_{PhOH}$_Prod$_7$** | –1317.328809 | –1316.901022 | –1316.875187 | 169.7 | –1316.950899 | 21.6 | 0 |
| **$O_{PhO-}$•** | –1317.377654 | –1316.956979 | –1316.929627 | 185.7 | –1317.009226 | 11.0 | 0 |
| **$O_{PhO-}$•_TS$_7$** | –1317.363660 | –1316.941474 | –1316.915529 | 174.4 | –1316.992411 | –441.9 | 1 |
| **$O_{PhO-}$•_Int$_7$** | –1317.387638 | –1316.963109 | –1316.937394 | 171.3 | –1317.013756 | 19.7 | 0 |
| **$O_{PhO}$-Prod$_7$** | –1316.837346 | –1316.423036 | –1316.397616 | 167.2 | –1316.472751 | 29.9 | 0 |
| **$O_{Ph}$•$_{6a}$** | –1242.647729 | –1242.217097 | –1242.190864 | 176.0 | –1242.267914 | 19.5 | 0 |
| **$O_{Ph}$•_TS$_{6a}$** | –1242.635330 | –1242.203326 | –1242.178381 | 168.2 | –1242.252730 | –413.5 | 1 |
| **$O_{Ph}$•_Int$_{6a}$** | –1242.650416 | –1242.216674 | –1242.191752 | 166.2 | –1242.266117 | 24.6 | 0 |
| **$O_{Ph}$•$_{6b}$** | –1242.646689 | –1242.215484 | –1242.189427 | 174.9 | –1242.266215 | 20.3 | 0 |
| **$O_{Ph}$•_TS$_{6b}$** | –1242.632267 | –1242.200340 | –1242.175466 | 167.6 | –1242.249875 | –419.3 | 1 |
| **$O_{Ph}$•_Int$_{6b}$** | –1242.647714 | –1242.213726 | –1242.188953 | 165.7 | –1242.263100 | 20.1 | 0 |
| **$O_{Ph}$h•$_{6c}$** | –1242.645698 | –1242.215018 | –1242.188872 | 176.6 | –1242.266175 | 16.1 | 0 |
| **$O_{Ph}$•_TS$_{6c}$** | –1242.632111 | –1242.200158 | –1242.175313 | 166.9 | –1242.249920 | –388.2 | 1 |
| **$O_{Ph}$•_Int$_{6c}$** | –1242.647595 | –1242.213740 | –1242.188924 | 164.8 | –1242.263385 | 34.1 | 0 |
| **$O_{Ph}$_Prod$_6$** | –1242.100992 | –1241.677421 | –1241.652868 | 164.1 | –1241.726230 | 24.2 | 0 |

[a]1 Hartree = 627.51 kcal $mol^{-1}$. Thermal corrections at 298.15 K.

**Table S7. Energies, entropies, and lowest frequencies of the lowest energy structures calculated at the PCM($Et_2O$)/ωB97X-D/6-311G(d,p) level of theory.[a]**

| Structure | $E_{elec}$ (Hartree) | $E_{elec}$ + ZPE (Hartree) | H (Hartree) | S (cal $mol^{-1}$ $K^{-1}$) | G (Hartree) | Lowest freq. ($cm^{-1}$) | # imag. freq. |
|---|---|---|---|---|---|---|---|
| **$O_{Me}$•'** | –1050.926866 | –1050.549068 | –1050.526181 | 156.2 | –1050.597222 | 35.4 | 0 |
| **$O_{Me}$•_$TS_7$'** | –1050.912248 | –1050.533881 | –1050.512029 | 149.8 | –1050.580971 | –420.8 | 1 |
| **$O_{Me}$•_$Int_7$'** | –1050.935980 | –1050.555633 | –1050.533782 | 150.4 | –1050.602334 | 24.8 | 0 |
| **$O_{Me}$_$Prod_7$'** | –1050.387214 | –1050.016459 | –1049.995214 | 144.1 | –1050.061988 | 50.0 | 0 |
| **$O_{Ph}$•'** | –1242.638329 | –1242.206911 | –1242.180971 | 173.4 | –1242.258086 | 25.3 | 0 |
| **$O_{Ph}$•_$TS_7$'** | –1242.625393 | –1242.193686 | –1242.168615 | 169.7 | –1242.243742 | –394.5 | 1 |
| **$O_{Ph}$•_$Int_7$'** | –1242.647712 | –1242.213895 | –1242.188976 | 168.2 | –1242.263653 | 20.0 | 0 |
| **$O_{Ph}$_$Prod_7$'** | –1242.095849 | –1241.671837 | –1241.647328 | 162.7 | –1241.720639 | 27.2 | 0 |
| **$O_{PhOH}$•'** | –1317.862471 | –1317.426522 | –1317.399465 | 178.7 | –1317.478559 | 22.3 | 0 |
| **$O_{PhOH}$•_$TS_7$'** | –1317.849188 | –1317.413290 | –1317.386984 | 175.3 | –1317.464423 | –400.0 | 1 |
| **$O_{PhOH}$•_$Int_7$'** | –1317.871825 | –1317.433514 | –1317.407396 | 173.4 | –1317.484296 | 22.3 | 0 |
| **$O_{PhOH}$_$Prod_7$'** | –1317.320092 | –1316.891929 | –1316.866188 | 168.5 | –1316.941778 | 23.8 | 0 |
| **$O_{PhO^-}$•'** | –1317.391476 | –1316.968874 | –1316.942429 | 176.3 | –1317.020454 | 26.5 | 0 |
| **$O_{PhO^-}$•_$TS_7$'** | –1317.336113 | –1316.913534 | –1316.887791 | 170.9 | –1316.964494 | –463.6 | 1 |
| **$O_{PhO^-}$•_$Int_7$'** | –1317.361979 | –1316.937435 | –1316.911797 | 170.1 | –1316.988111 | 23.4 | 0 |
| **$O_{PhO^-}$_$Prod_7$'** | –1316.811761 | –1316.397108 | –1316.371821 | 165.7 | –1316.446774 | 27.8 | 0 |

[a]1 Hartree = 627.51 kcal $mol^{-1}$. Thermal corrections at 298.15 K.

**Table S8: Cartesian coordinates of the structures calculated with PCM($H_2O$)/ωB97X-D/6-311G(d,p).**

Structure **5′-dAdo•**

| | | | |
|---|---|---|---|
| C | -3.235078 | 2.203239 | -0.361745 |
| C | -3.146425 | 0.789323 | -0.806082 |
| O | -1.816591 | 0.475934 | -1.261758 |
| C | -3.433407 | -0.220782 | 0.312728 |
| O | -3.752298 | -1.505087 | -0.187219 |
| C | -2.051414 | -0.365714 | 0.945019 |
| O | -1.880635 | -1.527758 | 1.699612 |
| C | -1.140273 | -0.297978 | -0.299205 |
| N | 3.938852 | -1.042588 | -0.348358 |
| C | 2.916857 | -1.845971 | -0.665837 |
| N | 1.615162 | -1.590958 | -0.624697 |
| C | 1.374932 | -0.344571 | -0.203828 |
| C | 2.321474 | 0.603521 | 0.156361 |
| C | 3.668948 | 0.203515 | 0.071486 |
| N | 4.684059 | 1.019186 | 0.393193 |
| N | 1.718459 | 1.787507 | 0.534307 |
| C | 0.443117 | 1.546213 | 0.406342 |
| N | 0.162099 | 0.274273 | -0.032226 |
| H | -2.524249 | 2.925107 | -0.743428 |
| H | -4.110754 | 2.549539 | 0.173840 |
| H | -3.823301 | 0.598854 | -1.650202 |
| H | -4.188144 | 0.136027 | 1.016855 |
| H | -4.688091 | -1.545536 | -0.389179 |
| H | -1.852871 | 0.489438 | 1.595587 |
| H | -2.396332 | -2.209647 | 1.253222 |
| H | -0.948722 | -1.305992 | -0.677207 |
| H | 3.196924 | -2.840591 | -1.001186 |
| H | 5.630722 | 0.690288 | 0.316707 |
| H | 4.506738 | 1.956097 | 0.709377 |
| H | -0.348532 | 2.255006 | 0.603564 |

Structure **5′-dAdoH**

| | | | |
|---|---|---|---|
| C | -3.557500 | 2.092500 | -0.239600 |
| C | -3.224000 | 0.694400 | -0.727000 |
| O | -1.843500 | 0.643100 | -1.151700 |
| C | -3.363900 | -0.385100 | 0.348800 |
| O | -3.592100 | -1.673200 | -0.198700 |
| C | -1.953300 | -0.447900 | 0.931800 |
| O | -1.661600 | -1.631000 | 1.613200 |
| C | -1.109700 | -0.236000 | -0.338400 |
| N | 3.961400 | -1.032200 | -0.337200 |
| C | 2.935000 | -1.826400 | -0.664000 |
| N | 1.635800 | -1.557100 | -0.642400 |
| C | 1.403500 | -0.305100 | -0.233100 |
| C | 2.355700 | 0.634700 | 0.135300 |
| C | 3.699600 | 0.219200 | 0.071400 |
| N | 4.719100 | 1.023700 | 0.408400 |
| N | 1.761200 | 1.828100 | 0.495500 |
| C | 0.484700 | 1.601100 | 0.349300 |
| N | 0.196700 | 0.330700 | -0.087700 |
| H | -3.009000 | 2.338000 | 0.674000 |
| H | -4.626900 | 2.175100 | -0.032000 |
| H | -3.840100 | 0.435700 | -1.591500 |
| H | -4.119600 | -0.132600 | 1.096400 |
| H | -4.528200 | -1.782600 | -0.371600 |
| H | -1.790100 | 0.384700 | 1.620400 |
| H | -2.143000 | -2.325900 | 1.148800 |
| H | -0.931200 | -1.197100 | -0.831800 |
| H | 3.208800 | -2.826100 | -0.989500 |
| H | 5.664200 | 0.695500 | 0.311800 |
| H | 4.549400 | 1.976200 | 0.679200 |
| H | -0.300900 | 2.320900 | 0.525100 |
| H | -3.296000 | 2.825100 | -1.005000 |

Structure $\mathbf{A_{Me}H}$

| | | | |
|---|---|---|---|
| C | 3.509966 | 0.818515 | -0.733148 |
| C | 2.462057 | 1.576209 | 0.053744 |
| O | 2.694405 | 2.030577 | 1.170457 |
| C | 3.639591 | -0.629913 | -0.216256 |
| C | 2.394906 | -1.447104 | -0.394305 |
| C | 2.096124 | -2.263168 | -1.449283 |
| C | 1.257254 | -1.503438 | 0.487275 |
| C | 0.314986 | -2.377898 | -0.101363 |
| C | 0.947270 | -0.894385 | 1.711966 |
| N | 0.851436 | -2.822215 | -1.281949 |
| C | -0.912830 | -2.662688 | 0.500850 |
| C | -0.269554 | -1.168855 | 2.308318 |
| C | -1.189141 | -2.049084 | 1.708634 |
| H | 3.268053 | 0.804684 | -1.798856 |
| H | 3.915083 | -0.597540 | 0.842545 |
| H | 2.687060 | -2.497639 | -2.322061 |
| H | 0.417446 | -3.469731 | -1.916624 |
| H | 1.644249 | -0.201161 | 2.170620 |
| H | -1.624555 | -3.334335 | 0.034488 |
| H | -0.523784 | -0.698966 | 3.251851 |
| N | 1.251951 | 1.668879 | -0.532181 |
| C | 0.081906 | 2.142406 | 0.169964 |
| C | -1.136434 | 1.513277 | -0.512546 |
| O | -1.048622 | 0.990031 | -1.617865 |
| H | 1.063465 | 1.170801 | -1.391686 |
| H | 0.130542 | 1.787991 | 1.204418 |
| N | -2.292405 | 1.626779 | 0.162078 |
| C | -3.559600 | 1.146632 | -0.364746 |
| C | -3.820066 | -0.326091 | -0.060302 |
| H | -2.272930 | 2.000545 | 1.097026 |
| H | -3.552049 | 1.306269 | -1.445491 |
| H | -3.800657 | -0.475546 | 1.024076 |
| H | -4.352216 | 1.769826 | 0.056686 |
| H | 4.461376 | 1.338228 | -0.602730 |
| H | 4.470068 | -1.105414 | -0.743874 |
| H | -2.136924 | -2.242558 | 2.197992 |
| C | -0.010943 | 3.672199 | 0.162723 |
| H | -0.092172 | 4.041800 | -0.862328 |
| H | -0.876478 | 4.018933 | 0.730938 |
| H | 0.890294 | 4.085369 | 0.617190 |
| C | -5.151602 | -0.797678 | -0.634662 |
| H | -5.323668 | -1.852957 | -0.411193 |
| H | -5.986541 | -0.226313 | -0.217518 |
| H | -5.175036 | -0.677523 | -1.722068 |
| H | -2.997112 | -0.917345 | -0.470941 |

Structure $\mathbf{A_{Me}}$**•**

| | | | |
|---|---|---|---|
| C | 3.736761 | 0.113397 | -0.336070 |
| C | 2.778792 | 1.074579 | 0.331165 |
| O | 2.886391 | 1.388600 | 1.512034 |
| C | 3.359160 | -1.338045 | 0.035391 |
| C | 1.957286 | -1.719705 | -0.341398 |
| C | 1.544220 | -2.269269 | -1.522566 |
| C | 0.769846 | -1.560316 | 0.459614 |
| C | -0.316543 | -2.052392 | -0.300261 |
| C | 0.525139 | -1.056811 | 1.745865 |
| N | 0.183797 | -2.467877 | -1.506512 |
| C | -1.621077 | -2.081709 | 0.198225 |
| C | -0.768148 | -1.068106 | 2.236082 |
| C | -1.831216 | -1.582984 | 1.469981 |
| H | 3.717193 | 0.229496 | -1.422663 |
| H | 3.493521 | -1.464958 | 1.113839 |
| H | 2.125725 | -2.546990 | -2.388893 |
| H | -0.350457 | -2.877316 | -2.253132 |
| H | 1.337703 | -0.646968 | 2.335666 |
| H | -2.438427 | -2.477625 | -0.393877 |
| H | -0.970189 | -0.678207 | 3.227527 |
| N | 1.756699 | 1.492550 | -0.445008 |

| | | | |
|---|---|---|---|
| C | 0.557885 | 2.085745 | 0.107563 |
| C | -0.612078 | 1.580986 | -0.735938 |
| O | -0.488277 | 1.400371 | -1.942770 |
| H | 1.661576 | 1.125153 | -1.381198 |
| H | 0.457340 | 1.734821 | 1.137673 |
| N | -1.768755 | 1.410379 | -0.079364 |
| C | -2.959683 | 0.907750 | -0.748555 |
| C | -4.080243 | 0.737648 | 0.213004 |
| H | -1.745871 | 1.363463 | 0.927596 |
| H | -2.730014 | -0.044661 | -1.240998 |
| H | -4.228778 | 1.507109 | 0.964797 |
| H | -3.227751 | 1.613029 | -1.550487 |
| C | -5.144381 | -0.267894 | -0.036652 |
| H | -5.788195 | -0.396706 | 0.835465 |
| H | -5.791873 | 0.020690 | -0.879347 |
| H | -4.715122 | -1.242366 | -0.297719 |
| H | 4.746324 | 0.331525 | 0.016940 |
| H | 4.070363 | -2.009344 | -0.451997 |
| H | -2.834788 | -1.580029 | 1.880201 |
| C | 0.600598 | 3.615656 | 0.092866 |
| H | -0.311727 | 4.031065 | 0.527033 |
| H | 1.455390 | 3.959503 | 0.677427 |
| H | 0.703049 | 3.980994 | -0.931657 |

Structure **$A_{Me}$•_TS$_6$**

| | | | |
|---|---|---|---|
| C | 3.638899 | -0.618072 | -0.000222 |
| C | 2.894359 | 0.584041 | 0.532278 |
| O | 2.923933 | 0.892460 | 1.718580 |
| C | 2.865289 | -1.903963 | 0.355116 |
| C | 1.480405 | -1.980722 | -0.221438 |
| C | 1.136336 | -2.518271 | -1.434088 |
| C | 0.251776 | -1.564142 | 0.394141 |
| C | -0.801599 | -1.911938 | -0.486933 |
| C | -0.075210 | -0.983833 | 1.640978 |
| N | -0.227078 | -2.469921 | -1.602149 |
| C | -2.143934 | -1.773210 | -0.150026 |
| C | -1.391980 | -0.790728 | 1.968704 |
| C | -2.462853 | -1.129923 | 1.067731 |
| H | 3.771340 | -0.554208 | -1.083167 |
| H | 2.809696 | -1.981478 | 1.445066 |
| H | 1.766974 | -2.954128 | -2.194293 |
| H | -0.724544 | -2.832456 | -2.396707 |
| H | 0.713017 | -0.702992 | 2.331967 |
| H | -2.926423 | -2.133445 | -0.808396 |
| H | -1.649203 | -0.365329 | 2.932961 |
| N | 2.150437 | 1.245773 | -0.381651 |
| C | 1.107739 | 2.177099 | -0.004397 |
| C | -0.154193 | 1.752651 | -0.755351 |
| O | -0.097259 | 1.427045 | -1.936604 |
| H | 2.066296 | 0.860988 | -1.311787 |
| H | 0.964829 | 2.075133 | 1.073030 |
| N | -1.286322 | 1.794396 | -0.041389 |
| C | -2.568009 | 1.359632 | -0.560128 |
| C | -3.411076 | 0.726198 | 0.501912 |
| H | -1.219234 | 1.980786 | 0.946220 |
| H | -2.377951 | 0.649029 | -1.370767 |
| H | -3.375461 | 1.223757 | 1.471027 |
| H | -3.103376 | 2.213335 | -1.003977 |
| C | -4.782161 | 0.310864 | 0.057786 |
| H | -5.329283 | -0.193530 | 0.857583 |
| H | -5.377493 | 1.180326 | -0.250456 |
| H | -4.729539 | -0.365514 | -0.802013 |
| H | 4.625256 | -0.643866 | 0.467028 |
| H | 3.452292 | -2.758420 | 0.009237 |
| H | -3.430683 | -1.338111 | 1.511500 |
| C | 1.469193 | 3.623377 | -0.343466 |
| H | 0.665462 | 4.300890 | -0.045224 |
| H | 2.381196 | 3.908447 | 0.184057 |
| H | 1.636288 | 3.728642 | -1.417956 |

Structure **$A_{Me}$•_Int$_6$**

| | | | |
|---|---|---|---|
| C | 3.647172 | -0.465005 | 0.015426 |
| C | 2.841731 | 0.733787 | 0.467733 |
| O | 2.915196 | 1.166428 | 1.612865 |
| C | 2.979811 | -1.770745 | 0.499054 |
| C | 1.608984 | -1.986979 | -0.072094 |
| C | 1.316594 | -2.627141 | -1.259729 |
| C | 0.373289 | -1.534650 | 0.456293 |
| C | -0.657329 | -1.926297 | -0.442815 |
| C | 0.012790 | -0.800218 | 1.636278 |
| N | -0.038420 | -2.592626 | -1.479328 |
| C | -1.989912 | -1.655149 | -0.249435 |
| C | -1.275540 | -0.482581 | 1.862434 |
| C | -2.422388 | -0.872790 | 0.959494 |
| H | 3.745601 | -0.485834 | -1.072808 |
| H | 2.929155 | -1.751339 | 1.591678 |
| H | 1.978750 | -3.111589 | -1.960936 |
| H | -0.505722 | -2.995135 | -2.272901 |
| H | 0.787411 | -0.479460 | 2.325625 |
| H | -2.737887 | -1.991388 | -0.958446 |
| H | -1.542675 | 0.081621 | 2.751793 |
| N | 2.005589 | 1.250630 | -0.459982 |
| C | 0.941817 | 2.172823 | -0.118554 |
| C | -0.330732 | 1.658042 | -0.795246 |
| O | -0.282929 | 1.162627 | -1.914668 |
| H | 1.885067 | 0.762645 | -1.336526 |
| H | 0.829542 | 2.143867 | 0.967102 |
| N | -1.459379 | 1.824898 | -0.085510 |
| C | -2.742288 | 1.268216 | -0.468794 |
| C | -3.337031 | 0.342693 | 0.598223 |
| H | -1.370796 | 2.181208 | 0.852924 |
| H | -2.583735 | 0.721094 | -1.399613 |
| H | -3.465648 | 0.924421 | 1.520297 |
| H | -3.450539 | 2.078084 | -0.671884 |
| C | -4.722505 | -0.115945 | 0.141449 |
| H | -5.192076 | -0.748940 | 0.898268 |
| H | -5.377597 | 0.741217 | -0.035144 |
| H | -4.666800 | -0.686475 | -0.790557 |
| H | 4.644999 | -0.381541 | 0.450316 |
| H | 3.630417 | -2.605610 | 0.227162 |
| H | -3.081640 | -1.517469 | 1.568808 |
| C | 1.247586 | 3.602632 | -0.568282 |
| H | 0.433766 | 4.277173 | -0.291687 |
| H | 2.166222 | 3.949149 | -0.091528 |
| H | 1.379225 | 3.637391 | -1.652382 |

Structure **$A_{Me}$_Prod$_6$**

| | | | |
|---|---|---|---|
| C | 3.577878 | 0.085439 | -0.145941 |
| C | 2.578632 | 1.127993 | 0.324831 |
| O | 2.673999 | 1.648419 | 1.431638 |
| C | 3.322826 | -1.301881 | 0.490237 |
| C | 2.019406 | -1.890657 | 0.044425 |
| C | 1.794637 | -2.670317 | -1.056006 |
| C | 0.722583 | -1.543359 | 0.556631 |
| C | -0.238255 | -2.103006 | -0.310142 |
| C | 0.287984 | -0.748623 | 1.624684 |
| N | 0.441820 | -2.822286 | -1.259934 |
| C | -1.592481 | -1.772417 | -0.229206 |
| C | -1.052074 | -0.440070 | 1.715627 |
| C | -1.989340 | -0.883749 | 0.755359 |
| H | 3.558352 | -0.011434 | -1.234374 |
| H | 3.347375 | -1.200868 | 1.579601 |
| H | 2.502954 | -3.120917 | -1.735040 |
| H | 0.026277 | -3.311074 | -2.033823 |
| H | 0.999954 | -0.322308 | 2.322674 |
| H | -2.283957 | -2.115967 | -0.990447 |
| H | -1.391231 | 0.228103 | 2.501240 |
| N | 1.581983 | 1.428885 | -0.538316 |
| C | 0.438592 | 2.235379 | -0.156928 |
| C | -0.808669 | 1.589413 | -0.772841 |
| O | -0.729940 | 0.954770 | -1.814548 |
| H | 1.458277 | 0.869649 | -1.370724 |

| | | | |
|---|---|---|---|
| H | 0.368293 | 2.203855 | 0.932357 |
| N | -1.967683 | 1.812429 | -0.109419 |
| C | -3.162375 | 1.015885 | -0.339803 |
| C | -3.315310 | -0.155944 | 0.669059 |
| H | -1.893710 | 2.262135 | 0.790266 |
| H | -3.089311 | 0.616058 | -1.352311 |
| H | -3.521311 | 0.283570 | 1.651796 |
| H | -4.042239 | 1.662239 | -0.290259 |
| C | -4.512383 | -1.018541 | 0.274923 |
| H | -4.647345 | -1.840875 | 0.980847 |
| H | -5.428435 | -0.421709 | 0.267437 |
| H | -4.390128 | -1.444677 | -0.724837 |
| H | 4.568048 | 0.441568 | 0.147442 |
| H | 4.151143 | -1.958359 | 0.212535 |
| C | 0.579328 | 3.688357 | -0.618703 |
| H | 0.660620 | 3.733389 | -1.707439 |
| H | -0.283668 | 4.280830 | -0.306175 |
| H | 1.478860 | 4.124663 | -0.181052 |

Structure $\mathbf{A_{Ph}H}$

| | | | |
|---|---|---|---|
| C | 4.584603 | 0.169361 | -1.195131 |
| C | 3.817865 | 1.154224 | -0.338902 |
| O | 4.289559 | 1.623192 | 0.692710 |
| C | 4.508930 | -1.248910 | -0.590239 |
| C | 3.114266 | -1.792962 | -0.499690 |
| C | 2.469146 | -2.559869 | -1.429091 |
| C | 2.170086 | -1.582331 | 0.567285 |
| C | 0.980937 | -2.260077 | 0.213712 |
| C | 2.219348 | -0.882805 | 1.781668 |
| N | 1.190253 | -2.841200 | -1.010148 |
| C | -0.143558 | -2.269412 | 1.042048 |
| C | 1.104838 | -0.881542 | 2.600078 |
| C | -0.064291 | -1.574043 | 2.234482 |
| H | 4.194179 | 0.150979 | -2.215743 |
| H | 4.961636 | -1.226277 | 0.405948 |
| H | 2.830079 | -2.939393 | -2.373298 |
| H | 0.526364 | -3.403579 | -1.513615 |
| H | 3.111336 | -0.331933 | 2.060108 |
| H | -1.047249 | -2.795542 | 0.756301 |
| H | 1.127991 | -0.339455 | 3.538841 |
| N | 2.567527 | 1.431255 | -0.760386 |
| C | 1.606574 | 2.129081 | 0.061555 |
| C | 0.216748 | 1.684450 | -0.402027 |
| O | 0.055158 | 1.090466 | -1.461931 |
| H | 2.172066 | 0.920585 | -1.538463 |
| H | 1.747108 | 1.817394 | 1.101591 |
| N | -0.798494 | 2.034693 | 0.406296 |
| C | -2.183228 | 1.744573 | 0.080344 |
| C | -2.593348 | 0.314611 | 0.452862 |
| H | -0.589772 | 2.450951 | 1.299105 |
| H | -2.318202 | 1.895798 | -0.992816 |
| H | -2.462100 | 0.174295 | 1.529179 |
| H | -2.811729 | 2.468553 | 0.602540 |
| H | 5.625675 | 0.496355 | -1.230085 |
| H | 5.125544 | -1.912057 | -1.201925 |
| H | -0.922467 | -1.555978 | 2.896815 |
| C | -4.018107 | 0.031334 | 0.054879 |
| C | -5.070145 | 0.266197 | 0.940775 |
| C | -4.316736 | -0.430850 | -1.227925 |
| C | -6.387891 | 0.041270 | 0.557688 |
| H | -4.852954 | 0.624158 | 1.942552 |
| C | -5.632583 | -0.656808 | -1.615500 |
| H | -3.507384 | -0.618789 | -1.927006 |
| C | -6.672978 | -0.421429 | -0.722557 |
| H | -7.192766 | 0.224650 | 1.260859 |
| H | -5.845872 | -1.020465 | -2.614571 |
| H | -7.699540 | -0.600016 | -1.021885 |
| C | 1.762924 | 3.650291 | -0.037567 |
| H | 1.590636 | 3.983063 | -1.063942 |
| H | 1.060881 | 4.165013 | 0.621626 |
| H | 2.777337 | 3.921857 | 0.256650 |

| | | | |
|---|---|---|---|
| H | -1.913955 | -0.378778 | -0.047808 |

Structure $\mathbf{A_{Ph}\bullet}$

| | | | |
|---|---|---|---|
| C | 4.741429 | -0.626292 | -0.284724 |
| C | 4.012484 | 0.548493 | 0.327670 |
| O | 4.167748 | 0.875631 | 1.499366 |
| C | 4.049712 | -1.942099 | 0.136999 |
| C | 2.595807 | -2.015354 | -0.229632 |
| C | 2.061782 | -2.502505 | -1.389478 |
| C | 1.479230 | -1.570366 | 0.565627 |
| C | 0.304302 | -1.839904 | -0.174035 |
| C | 1.363096 | -0.979396 | 1.832477 |
| N | 0.690960 | -2.396428 | -1.365426 |
| C | -0.970066 | -1.565728 | 0.327224 |
| C | 0.103117 | -0.690535 | 2.325200 |
| C | -1.053869 | -0.988523 | 1.580373 |
| H | 4.755439 | -0.554209 | -1.375156 |
| H | 4.156570 | -2.056795 | 1.219858 |
| H | 2.560285 | -2.932684 | -2.245181 |
| H | 0.072948 | -2.709574 | -2.093837 |
| H | 2.251476 | -0.735689 | 2.404553 |
| H | -1.862220 | -1.787944 | -0.247343 |
| H | 0.000623 | -0.231507 | 3.302232 |
| N | 3.121101 | 1.151638 | -0.487517 |
| C | 2.064298 | 2.004406 | 0.013719 |
| C | 0.829938 | 1.716330 | -0.838902 |
| O | 0.931535 | 1.483532 | -2.038133 |
| H | 2.970547 | 0.781302 | -1.415074 |
| H | 1.877519 | 1.726525 | 1.054037 |
| N | -0.349809 | 1.790555 | -0.202799 |
| C | -1.594061 | 1.503717 | -0.901467 |
| C | -2.741149 | 1.547209 | 0.047167 |
| H | -0.353280 | 1.750058 | 0.804940 |
| H | -1.510664 | 0.528993 | -1.391683 |
| H | -2.689945 | 2.282936 | 0.845180 |
| H | -1.722377 | 2.245835 | -1.701308 |
| H | 5.771729 | -0.621626 | 0.075687 |
| H | 4.590987 | -2.772497 | -0.322843 |
| H | -2.029645 | -0.752891 | 1.990092 |
| C | -3.890079 | 0.730538 | -0.021736 |
| C | -4.889856 | 0.845439 | 0.980485 |
| C | -4.105019 | -0.226492 | -1.047619 |
| C | -6.017209 | 0.047799 | 0.963221 |
| H | -4.752041 | 1.572190 | 1.774383 |
| C | -5.239239 | -1.018338 | -1.054819 |
| H | -3.378652 | -0.340122 | -1.843599 |
| C | -6.202035 | -0.892554 | -0.052924 |
| H | -6.761786 | 0.153349 | 1.744293 |
| H | -5.379458 | -1.742352 | -1.849704 |
| H | -7.086949 | -1.517834 | -0.064465 |
| C | 2.421966 | 3.490653 | -0.060508 |
| H | 1.607369 | 4.102695 | 0.333462 |
| H | 3.319227 | 3.677516 | 0.531574 |
| H | 2.614792 | 3.781672 | -1.095689 |

Structure $\mathbf{A_{Ph}\bullet\_TS_6}$

| | | | |
|---|---|---|---|
| C | 4.672441 | -0.778018 | -0.034224 |
| C | 3.997738 | 0.474081 | 0.476415 |
| O | 4.061157 | 0.811851 | 1.653168 |
| C | 3.847707 | -2.014519 | 0.377453 |
| C | 2.447311 | -2.043573 | -0.165289 |
| C | 2.046659 | -2.615957 | -1.346322 |
| C | 1.259050 | -1.538033 | 0.455556 |
| C | 0.166436 | -1.867489 | -0.386928 |
| C | 0.994385 | -0.883834 | 1.682580 |
| N | 0.685664 | -2.504063 | -1.487962 |
| C | -1.155256 | -1.642717 | -0.029684 |
| C | -0.298822 | -0.602989 | 2.028403 |
| C | -1.415656 | -0.922276 | 1.166900 |
| H | 4.783752 | -0.752973 | -1.121118 |
| H | 3.814072 | -2.056615 | 1.470186 |

| | | | |
|---|---|---|---|
| H | 2.636726 | -3.117138 | -2.098757 |
| H | 0.151515 | -2.873835 | -2.255088 |
| H | 1.813817 | -0.616812 | 2.341644 |
| H | -1.974475 | -1.979182 | -0.655468 |
| H | -0.509712 | -0.122357 | 2.977907 |
| N | 3.273601 | 1.147705 | -0.445008 |
| C | 2.277058 | 2.133175 | -0.080464 |
| C | 0.983511 | 1.727513 | -0.786588 |
| O | 1.003833 | 1.334498 | -1.947899 |
| H | 3.156663 | 0.741244 | -1.362323 |
| H | 2.155554 | 2.077260 | 1.002999 |
| N | -0.134619 | 1.856595 | -0.059162 |
| C | -1.436155 | 1.449860 | -0.545751 |
| C | -2.283429 | 0.882134 | 0.557191 |
| H | -0.047943 | 2.116693 | 0.910115 |
| H | -1.266437 | 0.713376 | -1.335510 |
| H | -2.183847 | 1.399149 | 1.510294 |
| H | -1.952427 | 2.303204 | -1.007871 |
| H | 5.666547 | -0.837617 | 0.413035 |
| H | 4.385331 | -2.905991 | 0.045223 |
| H | -2.360880 | -1.114037 | 1.664664 |
| C | -3.659269 | 0.510120 | 0.236075 |
| C | -4.617373 | 0.428348 | 1.260711 |
| C | -4.056084 | 0.153239 | -1.062602 |
| C | -5.914686 | 0.015934 | 0.999081 |
| H | -4.331186 | 0.697638 | 2.272855 |
| C | -5.357571 | -0.257275 | -1.322785 |
| H | -3.345636 | 0.196168 | -1.880167 |
| C | -6.293455 | -0.329229 | -0.296495 |
| H | -6.636145 | -0.034348 | 1.807116 |
| H | -5.641560 | -0.523966 | -2.334832 |
| H | -7.308023 | -0.650135 | -0.502629 |
| C | 2.684910 | 3.550105 | -0.484636 |
| H | 1.914915 | 4.269768 | -0.196516 |
| H | 3.619059 | 3.819606 | 0.011265 |
| H | 2.831187 | 3.606840 | -1.565834 |

Structure **$A_{Ph}$•_Int$_6$**

| | | | |
|---|---|---|---|
| C | 4.655491 | -0.769331 | -0.004709 |
| C | 3.970444 | 0.507216 | 0.433405 |
| O | 4.093972 | 0.949531 | 1.570405 |
| C | 3.880370 | -1.999314 | 0.515227 |
| C | 2.486787 | -2.104544 | -0.031660 |
| C | 2.115669 | -2.754044 | -1.191992 |
| C | 1.306770 | -1.525781 | 0.499909 |
| C | 0.228240 | -1.852181 | -0.368889 |
| C | 1.036418 | -0.727648 | 1.661856 |
| N | 0.765584 | -2.604249 | -1.391982 |
| C | -1.071123 | -1.458289 | -0.161371 |
| C | -0.212852 | -0.285992 | 1.897940 |
| C | -1.404427 | -0.600127 | 1.024793 |
| H | 4.735336 | -0.820506 | -1.093488 |
| H | 3.848220 | -1.953455 | 1.607780 |
| H | 2.718445 | -3.319565 | -1.886000 |
| H | 0.248548 | -2.991905 | -2.161797 |
| H | 1.849150 | -0.461159 | 2.329863 |
| H | -1.865392 | -1.747714 | -0.840129 |
| H | -0.412826 | 0.324829 | 2.773814 |
| N | 3.179098 | 1.085748 | -0.497264 |
| C | 2.208987 | 2.109517 | -0.166180 |
| C | 0.888331 | 1.696812 | -0.819119 |
| O | 0.879471 | 1.175295 | -1.927334 |
| H | 3.007476 | 0.598392 | -1.365561 |
| H | 2.107341 | 2.115462 | 0.920912 |
| N | -0.215888 | 1.974461 | -0.104086 |
| C | -1.540229 | 1.522326 | -0.478900 |
| C | -2.206086 | 0.694667 | 0.627436 |
| H | -0.092465 | 2.349008 | 0.823256 |
| H | -1.413192 | 0.934838 | -1.388265 |
| H | -2.245365 | 1.316050 | 1.530218 |
| H | -2.178089 | 2.379277 | -0.716603 |
| H | 5.663285 | -0.769289 | 0.414997 |
| H | 4.449201 | -2.894692 | 0.252909 |
| H | -2.115521 | -1.150576 | 1.664854 |
| C | -3.639741 | 0.346605 | 0.275842 |
| C | -4.592454 | 0.281652 | 1.295682 |
| C | -4.045462 | 0.043115 | -1.025024 |
| C | -5.907584 | -0.076396 | 1.029333 |
| H | -4.296290 | 0.517393 | 2.313415 |
| C | -5.362634 | -0.314554 | -1.296553 |
| H | -3.336238 | 0.082134 | -1.843630 |
| C | -6.298379 | -0.376496 | -0.271718 |
| H | -6.629247 | -0.116700 | 1.837578 |
| H | -5.655513 | -0.544997 | -2.314776 |
| H | -7.324575 | -0.653682 | -0.484258 |
| C | 2.637977 | 3.494630 | -0.653337 |
| H | 1.892223 | 4.246749 | -0.385193 |
| H | 3.589801 | 3.766726 | -0.193832 |
| H | 2.759511 | 3.491989 | -1.739136 |

Structure **$A_{Ph}$_Prod$_6$**

| | | | |
|---|---|---|---|
| C | 4.504892 | -0.845790 | -0.124736 |
| C | 3.834376 | 0.428210 | 0.359667 |
| O | 4.057903 | 0.882249 | 1.477008 |
| C | 3.864298 | -2.115667 | 0.485042 |
| C | 2.452980 | -2.308762 | 0.020926 |
| C | 2.029441 | -2.979389 | -1.092787 |
| C | 1.301090 | -1.617175 | 0.529596 |
| C | 0.230550 | -1.872385 | -0.352725 |
| C | 1.096539 | -0.749983 | 1.609545 |
| N | 0.690935 | -2.742030 | -1.308264 |
| C | -0.976612 | -1.176362 | -0.274216 |
| C | -0.102262 | -0.075492 | 1.697011 |
| C | -1.112212 | -0.225050 | 0.721343 |
| H | 4.469764 | -0.915987 | -1.214873 |
| H | 3.904230 | -2.042464 | 1.576169 |
| H | 2.589608 | -3.602178 | -1.774106 |
| H | 0.162316 | -3.086358 | -2.090836 |
| H | 1.890396 | -0.555631 | 2.321898 |
| H | -1.738517 | -1.302428 | -1.035857 |
| H | -0.249161 | 0.648120 | 2.492610 |
| N | 2.973235 | 1.011518 | -0.504701 |
| C | 2.099264 | 2.102410 | -0.117586 |
| C | 0.730719 | 1.844986 | -0.759008 |
| O | 0.644669 | 1.237688 | -1.816000 |
| H | 2.711590 | 0.525664 | -1.350995 |
| H | 2.006963 | 2.074248 | 0.970202 |
| N | -0.332757 | 2.364501 | -0.099617 |
| C | -1.693774 | 1.936329 | -0.374600 |
| C | -2.178123 | 0.855251 | 0.631710 |
| H | -0.155620 | 2.756593 | 0.812526 |
| H | -1.688220 | 1.539580 | -1.388832 |
| H | -2.236315 | 1.339954 | 1.612219 |
| H | -2.365018 | 2.797545 | -0.344377 |
| H | 5.552280 | -0.787289 | 0.179962 |
| H | 4.478336 | -2.973679 | 0.200711 |
| C | -3.565348 | 0.330855 | 0.315019 |
| C | -4.384423 | -0.077223 | 1.371099 |
| C | -4.052233 | 0.199961 | -0.985954 |
| C | -5.647684 | -0.603335 | 1.138346 |
| H | -4.021540 | 0.018764 | 2.389800 |
| C | -5.318961 | -0.326859 | -1.223623 |
| H | -3.450118 | 0.506773 | -1.833137 |
| C | -6.120792 | -0.730803 | -0.164354 |
| H | -6.265421 | -0.911707 | 1.974405 |
| H | -5.677180 | -0.419043 | -2.242836 |
| H | -7.107523 | -1.139360 | -0.350129 |
| C | 2.652684 | 3.463140 | -0.549593 |
| H | 2.759926 | 3.500168 | -1.636347 |
| H | 1.989823 | 4.272247 | -0.234307 |
| H | 3.632476 | 3.617846 | -0.094577 |

Structure $\mathbf{O_{Me}\bullet}$

| | | | |
|---|---|---|---|
| C | -3.357863 | -1.147919 | -1.317052 |
| C | -2.657331 | -1.389972 | 0.005767 |
| O | -3.189370 | -1.161992 | 1.084494 |
| C | -3.280333 | 0.361796 | -1.674610 |
| C | -1.916897 | 0.933715 | -1.416510 |
| C | -0.781006 | 0.762440 | -2.158323 |
| C | -1.498306 | 1.563306 | -0.191497 |
| C | -0.099558 | 1.740084 | -0.272046 |
| C | -2.169552 | 1.952608 | 0.976952 |
| N | 0.311295 | 1.263243 | -1.488922 |
| C | 0.638436 | 2.296183 | 0.777212 |
| C | -1.441949 | 2.491930 | 2.020017 |
| C | -0.047163 | 2.661865 | 1.920013 |
| H | -2.886589 | -1.732420 | -2.111200 |
| H | -4.017378 | 0.902435 | -1.075267 |
| H | -0.656589 | 0.287224 | -3.120049 |
| H | 1.279676 | 1.054646 | -1.709293 |
| H | -3.239327 | 1.803879 | 1.070097 |
| H | 1.709279 | 2.439765 | 0.696560 |
| H | -1.947073 | 2.788410 | 2.932458 |
| N | -1.369317 | -1.792187 | -0.107630 |
| C | -0.391308 | -1.594527 | 0.940603 |
| C | 0.974436 | -1.524872 | 0.253583 |
| O | 1.161868 | -2.004245 | -0.854104 |
| H | -0.970558 | -1.886644 | -1.030862 |
| H | -0.596151 | -0.638619 | 1.436809 |
| N | 1.965662 | -0.957139 | 0.977601 |
| C | 3.324893 | -0.951395 | 0.531744 |
| C | 3.767221 | 0.334010 | -0.284997 |
| H | 1.734399 | -0.478889 | 1.834365 |
| H | 3.502497 | -1.822691 | -0.097029 |
| H | 3.464439 | 1.188726 | 0.339275 |
| H | 3.987735 | -0.978912 | 1.396841 |
| H | -4.399963 | -1.460538 | -1.233103 |
| H | -3.571631 | 0.483874 | -2.720399 |
| H | 0.497846 | 3.090660 | 2.753537 |
| O | 3.080587 | 0.304665 | -1.449111 |
| C | -0.406034 | -2.720995 | 1.979133 |
| H | -0.180890 | -3.677620 | 1.501864 |
| H | 0.325784 | -2.538071 | 2.768757 |
| H | -1.398975 | -2.772614 | 2.426263 |
| C | 5.279711 | 0.294156 | -0.484536 |
| H | 5.592529 | 1.145309 | -1.091300 |
| H | 5.795144 | 0.348555 | 0.477311 |
| H | 5.573242 | -0.625781 | -0.995941 |

Structure $\mathbf{O_{Me}\bullet\_TS_7}$

| | | | |
|---|---|---|---|
| C | 3.652881 | -0.067435 | -0.747240 |
| C | 2.875484 | 0.920444 | 0.094995 |
| O | 3.172070 | 1.148312 | 1.264509 |
| C | 3.317945 | -1.510578 | -0.315162 |
| C | 1.871440 | -1.876908 | -0.473424 |
| C | 1.294516 | -2.496441 | -1.545841 |
| C | 0.806544 | -1.651351 | 0.472696 |
| C | -0.370167 | -2.176606 | -0.092252 |
| C | 0.748660 | -1.069223 | 1.746902 |
| N | -0.054051 | -2.670729 | -1.316135 |
| C | -1.636484 | -2.095185 | 0.545284 |
| C | -0.470823 | -1.047103 | 2.427522 |
| C | -1.619160 | -1.591990 | 1.868737 |
| H | 3.433177 | 0.058410 | -1.810176 |
| H | 3.614770 | -1.635265 | 0.730410 |
| H | 1.742317 | -2.847642 | -2.462719 |
| H | -0.700395 | -3.097080 | -1.958470 |
| H | 1.636001 | -0.634260 | 2.192638 |
| H | -2.446365 | -2.737373 | 0.228064 |
| H | -0.514906 | -0.622627 | 3.423828 |
| N | 1.800026 | 1.467198 | -0.503100 |
| C | 0.784473 | 2.199874 | 0.222557 |
| C | -0.525941 | 1.988799 | -0.538567 |
| O | -0.520694 | 1.891793 | -1.759515 |
| H | 1.559619 | 1.214292 | -1.450718 |
| H | 0.712244 | 1.768718 | 1.224727 |
| N | -1.636475 | 1.983045 | 0.218539 |
| C | -2.944545 | 1.660356 | -0.311067 |
| C | -3.326621 | 0.196950 | -0.022812 |
| H | -1.521623 | 1.957124 | 1.219780 |
| H | -2.914330 | 1.828472 | -1.388299 |
| H | -3.316825 | 0.085367 | 1.069980 |
| H | -3.688097 | 2.332814 | 0.125395 |
| C | -4.732230 | -0.117974 | -0.530919 |
| H | -4.994321 | -1.148650 | -0.281216 |
| H | -5.479821 | 0.545444 | -0.085593 |
| H | -4.773636 | -0.007474 | -1.618650 |
| H | 4.717623 | 0.118353 | -0.591849 |
| H | 3.935012 | -2.194066 | -0.903070 |
| H | -2.540069 | -1.597944 | 2.439617 |
| O | -2.374160 | -0.635508 | -0.602231 |
| C | 1.105695 | 3.692592 | 0.331976 |
| H | 1.178920 | 4.135421 | -0.664004 |
| H | 0.328257 | 4.214279 | 0.894722 |
| H | 2.058210 | 3.820968 | 0.848064 |

Structure $\mathbf{O_{Me}\bullet\_Int_7}$

| | | | |
|---|---|---|---|
| C | 3.700014 | -0.139606 | -0.732171 |
| C | 2.915632 | 0.884027 | 0.061729 |
| O | 3.223592 | 1.190552 | 1.209356 |
| C | 3.317416 | -1.569114 | -0.284286 |
| C | 1.855479 | -1.864940 | -0.443397 |
| C | 1.230559 | -2.378986 | -1.548858 |
| C | 0.816326 | -1.580588 | 0.509637 |
| C | -0.390431 | -1.944360 | -0.075444 |
| C | 0.833540 | -1.037092 | 1.821265 |
| N | -0.125725 | -2.423922 | -1.316221 |
| C | -1.739820 | -1.690289 | 0.498307 |
| C | -0.394971 | -0.899970 | 2.496210 |
| C | -1.596238 | -1.227214 | 1.924172 |
| H | 3.512156 | -0.032271 | -1.803445 |
| H | 3.608734 | -1.694843 | 0.762700 |
| H | 1.641566 | -2.725476 | -2.484188 |
| H | -0.816598 | -2.729081 | -1.981168 |
| H | 1.761133 | -0.729122 | 2.286698 |
| H | -2.363726 | -2.595861 | 0.474605 |
| H | -0.388369 | -0.546029 | 3.522242 |
| N | 1.821302 | 1.374882 | -0.555446 |
| C | 0.776135 | 2.080259 | 0.155116 |
| C | -0.529067 | 1.807696 | -0.597487 |
| O | -0.519043 | 1.585166 | -1.801161 |
| H | 1.557936 | 1.024133 | -1.465623 |
| H | 0.713236 | 1.663770 | 1.164847 |
| N | -1.648336 | 1.901637 | 0.145562 |
| C | -2.957667 | 1.552888 | -0.367999 |
| C | -3.368641 | 0.137500 | 0.048497 |
| H | -1.542329 | 1.968780 | 1.145912 |
| H | -2.908306 | 1.614229 | -1.455804 |
| H | -3.434285 | 0.115522 | 1.139729 |
| H | -3.695615 | 2.275932 | -0.011759 |
| C | -4.711071 | -0.260155 | -0.545169 |
| H | -4.984583 | -1.266167 | -0.219902 |
| H | -5.498415 | 0.429304 | -0.228311 |
| H | -4.658788 | -0.251331 | -1.637333 |
| H | 4.763092 | 0.028890 | -0.549522 |
| H | 3.910704 | -2.278486 | -0.866318 |
| H | -2.497732 | -1.167764 | 2.522012 |
| O | -2.348243 | -0.734051 | -0.406852 |
| C | 1.042638 | 3.585726 | 0.241075 |
| H | 1.095794 | 4.017234 | -0.761295 |
| H | 0.251265 | 4.088879 | 0.801121 |
| H | 1.992993 | 3.753558 | 0.749605 |

Structure **$O_{Me}$•_Prod$_7$**

| | | | |
|---|---|---|---|
| C | 3.758968 | -0.098959 | -0.736705 |
| C | 2.937132 | 0.810119 | 0.157739 |
| O | 3.192187 | 0.961086 | 1.346651 |
| C | 3.311517 | -1.573278 | -0.563326 |
| C | 1.825888 | -1.719053 | -0.705645 |
| C | 1.100119 | -1.728361 | -1.866269 |
| C | 0.871235 | -1.612573 | 0.365412 |
| C | -0.400465 | -1.534773 | -0.232320 |
| C | 0.979120 | -1.519896 | 1.764021 |
| N | -0.240569 | -1.634320 | -1.586415 |
| C | -1.558415 | -1.289134 | 0.512954 |
| C | -0.173300 | -1.347931 | 2.501956 |
| C | -1.437419 | -1.229624 | 1.887649 |
| H | 3.643394 | 0.189369 | -1.784676 |
| H | 3.623881 | -1.924223 | 0.423944 |
| H | 1.441971 | -1.776364 | -2.889193 |
| H | -0.978135 | -1.536184 | -2.261772 |
| H | 1.948165 | -1.553066 | 2.247325 |
| H | -0.115768 | -1.279825 | 3.582493 |
| N | 1.868736 | 1.386700 | -0.439861 |
| C | 0.759498 | 1.932326 | 0.315109 |
| C | -0.503815 | 1.690854 | -0.518831 |
| O | -0.443228 | 1.562850 | -1.734513 |
| H | 1.640001 | 1.130712 | -1.390839 |
| H | 0.681741 | 1.374476 | 1.253869 |
| N | -1.654623 | 1.658146 | 0.181596 |
| C | -2.936482 | 1.320153 | -0.407324 |
| C | -3.537674 | 0.023400 | 0.142176 |
| H | -1.596435 | 1.704379 | 1.187062 |
| H | -2.783108 | 1.225131 | -1.482881 |
| H | -3.629828 | 0.099386 | 1.230311 |
| H | -3.649355 | 2.131384 | -0.232152 |
| C | -4.902701 | -0.240526 | -0.461459 |
| H | -5.314057 | -1.169825 | -0.065194 |
| H | -5.588763 | 0.574341 | -0.222330 |
| H | -4.827601 | -0.326346 | -1.548558 |
| H | 4.810766 | 0.006306 | -0.464682 |
| H | 3.841130 | -2.181842 | -1.300167 |
| H | -2.313139 | -1.059886 | 2.502365 |
| O | -2.720898 | -1.111033 | -0.187099 |
| C | 0.935237 | 3.422663 | 0.618773 |
| H | 1.002412 | 3.992959 | -0.310753 |
| H | 0.097785 | 3.804260 | 1.207220 |
| H | 1.855002 | 3.565508 | 1.187593 |

Structure **$O_{Ph}$•**

| | | | |
|---|---|---|---|
| C | 4.607047 | 0.325896 | -1.270820 |
| C | 3.875513 | 1.184622 | -0.263452 |
| O | 4.362476 | 1.459725 | 0.828789 |
| C | 4.551918 | -1.154384 | -0.832798 |
| C | 3.158570 | -1.698536 | -0.717466 |
| C | 2.447486 | -2.343074 | -1.690732 |
| C | 2.282634 | -1.612494 | 0.423739 |
| C | 1.065584 | -2.234485 | 0.063731 |
| C | 2.411740 | -1.068521 | 1.710898 |
| N | 1.193395 | -2.666231 | -1.230164 |
| C | -0.009116 | -2.336191 | 0.949512 |
| C | 1.343612 | -1.151124 | 2.584860 |
| C | 0.143722 | -1.784201 | 2.207353 |
| H | 4.173875 | 0.428472 | -2.268920 |
| H | 5.065142 | -1.248530 | 0.129003 |
| H | 2.744723 | -2.605509 | -2.695055 |
| H | 0.486502 | -3.145130 | -1.760579 |
| H | 3.327018 | -0.565374 | 2.001692 |
| H | -0.933187 | -2.818695 | 0.653225 |
| H | 1.429228 | -0.727584 | 3.579409 |
| N | 2.626639 | 1.553634 | -0.614445 |
| C | 1.685285 | 2.101887 | 0.338180 |
| C | 0.286450 | 1.827527 | -0.202831 |
| O | 0.047611 | 1.881544 | -1.399340 |
| H | 2.250853 | 1.277652 | -1.509120 |
| H | 1.828643 | 1.585817 | 1.291394 |
| N | -0.665272 | 1.602816 | 0.730170 |
| C | -2.044566 | 1.454046 | 0.384193 |
| C | -2.481244 | -0.039659 | 0.061743 |
| H | -0.384648 | 1.437366 | 1.684359 |
| H | -2.267385 | 2.059220 | -0.493944 |
| H | -2.162072 | -0.608085 | 0.951216 |
| H | -2.667517 | 1.774506 | 1.218527 |
| H | 5.646735 | 0.657194 | -1.309624 |
| H | 5.121622 | -1.744337 | -1.554870 |
| H | -0.674755 | -1.840214 | 2.916768 |
| C | -3.990116 | -0.072986 | -0.097113 |
| C | -4.805429 | -0.271125 | 1.014452 |
| C | -4.570869 | 0.122311 | -1.346917 |
| C | -6.189251 | -0.272289 | 0.879190 |
| H | -4.356279 | -0.430197 | 1.990475 |
| C | -5.954057 | 0.121354 | -1.483895 |
| H | -3.931517 | 0.264475 | -2.210992 |
| C | -6.765986 | -0.075053 | -0.371160 |
| H | -6.816747 | -0.431488 | 1.748768 |
| H | -6.399462 | 0.272606 | -2.460745 |
| H | -7.844687 | -0.077634 | -0.478618 |
| O | -1.780607 | -0.425157 | -1.017569 |
| C | 1.866955 | 3.609559 | 0.543103 |
| H | 1.717013 | 4.138142 | -0.400905 |
| H | 1.154083 | 3.987343 | 1.279562 |
| H | 2.878367 | 3.801075 | 0.902401 |

Structure **$O_{Ph}$•_TS$_7$**

| | | | |
|---|---|---|---|
| C | 4.590512 | -0.343038 | -0.877421 |
| C | 3.915519 | 0.739643 | -0.063056 |
| O | 4.268408 | 1.001105 | 1.083355 |
| C | 4.196225 | -1.736545 | -0.346668 |
| C | 2.724182 | -2.016289 | -0.419432 |
| C | 2.050468 | -2.610853 | -1.448715 |
| C | 1.729288 | -1.701375 | 0.575004 |
| C | 0.490660 | -2.150187 | 0.080877 |
| C | 1.778382 | -1.092106 | 1.836404 |
| N | 0.706608 | -2.683981 | -1.147776 |
| C | -0.730921 | -1.947933 | 0.773876 |
| C | 0.601609 | -0.976914 | 2.581209 |
| C | -0.611206 | -1.440570 | 2.089882 |
| H | 4.327297 | -0.264841 | -1.935034 |
| H | 4.534887 | -1.821611 | 0.689886 |
| H | 2.424066 | -3.007773 | -2.380010 |
| H | 0.000497 | -3.078435 | -1.746227 |
| H | 2.715571 | -0.709416 | 2.224441 |
| H | -1.620823 | -2.489874 | 0.482974 |
| H | 0.640821 | -0.537200 | 3.570964 |
| N | 2.869784 | 1.344666 | -0.658637 |
| C | 1.957767 | 2.213277 | 0.052865 |
| C | 0.592430 | 2.056207 | -0.621510 |
| O | 0.512974 | 1.791987 | -1.812858 |
| H | 2.564616 | 1.053399 | -1.576331 |
| H | 1.903412 | 1.864252 | 1.087840 |
| N | -0.468999 | 2.283580 | 0.177314 |
| C | -1.818923 | 1.935911 | -0.208803 |
| C | -2.174507 | 0.513374 | 0.290324 |
| H | -0.291424 | 2.385828 | 1.164640 |
| H | -1.883206 | 1.969913 | -1.296465 |
| H | -2.014896 | 0.535376 | 1.378980 |
| H | -2.518444 | 2.663465 | 0.208392 |
| H | 5.670532 | -0.211694 | -0.782421 |
| H | 4.742711 | -2.487956 | -0.921374 |
| H | -1.502903 | -1.363287 | 2.700598 |
| C | -3.640445 | 0.201344 | 0.035391 |
| C | -4.635290 | 0.763657 | 0.835671 |
| C | -4.010621 | -0.637373 | -1.012036 |
| C | -5.976892 | 0.497639 | 0.589351 |
| H | -4.357285 | 1.409695 | 1.663745 |

| | | | |
|---|---|---|---|
| C | -5.352659 | -0.907258 | -1.260403 |
| H | -3.231655 | -1.082220 | -1.620185 |
| C | -6.339567 | -0.339353 | -0.462076 |
| H | -6.740184 | 0.938326 | 1.221149 |
| H | -5.628317 | -1.564311 | -2.078121 |
| H | -7.385547 | -0.550852 | -0.653741 |
| O | -1.299148 | -0.361509 | -0.320077 |
| C | 2.408725 | 3.676398 | 0.031813 |
| H | 2.464314 | 4.038111 | -0.997604 |
| H | 1.711924 | 4.305746 | 0.589915 |
| H | 3.396005 | 3.756903 | 0.489020 |

Structure **$O_{Ph}$•_Int$_7$**

| | | | |
|---|---|---|---|
| C | 4.597238 | -0.497439 | -0.893074 |
| C | 3.953911 | 0.646191 | -0.136418 |
| O | 4.346901 | 0.990485 | 0.973871 |
| C | 4.138907 | -1.852907 | -0.309474 |
| C | 2.651441 | -2.038000 | -0.360174 |
| C | 1.918049 | -2.565269 | -1.390645 |
| C | 1.701903 | -1.612959 | 0.632419 |
| C | 0.435110 | -1.910717 | 0.144490 |
| C | 1.844695 | -1.007336 | 1.908754 |
| N | 0.580039 | -2.481425 | -1.077259 |
| C | -0.836873 | -1.458004 | 0.766482 |
| C | 0.679307 | -0.771219 | 2.665369 |
| C | -0.577733 | -1.032473 | 2.186312 |
| H | 4.345269 | -0.455321 | -1.955667 |
| H | 4.486009 | -1.921776 | 0.725580 |
| H | 2.239943 | -3.005602 | -2.321375 |
| H | -0.172937 | -2.775324 | -1.676806 |
| H | 2.821399 | -0.744249 | 2.294755 |
| H | -1.616605 | -2.230814 | 0.735466 |
| H | 0.783773 | -0.397629 | 3.678973 |
| N | 2.887947 | 1.211664 | -0.739568 |
| C | 1.971099 | 2.084583 | -0.038911 |
| C | 0.601274 | 1.909553 | -0.702348 |
| O | 0.509437 | 1.527591 | -1.859911 |
| H | 2.535046 | 0.827924 | -1.605321 |
| H | 1.914799 | 1.749045 | 1.001522 |
| N | -0.452900 | 2.259586 | 0.065765 |
| C | -1.814095 | 1.901394 | -0.279103 |
| C | -2.200736 | 0.569446 | 0.383135 |
| H | -0.269365 | 2.460014 | 1.037010 |
| H | -1.873512 | 1.801786 | -1.363279 |
| H | -2.094255 | 0.702536 | 1.465712 |
| H | -2.498201 | 2.690284 | 0.039382 |
| H | 5.680396 | -0.403252 | -0.792173 |
| H | 4.637840 | -2.649998 | -0.865941 |
| H | -1.436189 | -0.897140 | 2.833630 |
| C | -3.633113 | 0.185916 | 0.075030 |
| C | -4.677346 | 0.707631 | 0.837389 |
| C | -3.927218 | -0.660288 | -0.991535 |
| C | -5.997103 | 0.393357 | 0.535223 |
| H | -4.455975 | 1.359824 | 1.677088 |
| C | -5.246810 | -0.979817 | -1.292070 |
| H | -3.112964 | -1.072764 | -1.575900 |
| C | -6.284855 | -0.452396 | -0.531445 |
| H | -6.800910 | 0.802735 | 1.136740 |
| H | -5.464844 | -1.643000 | -2.121804 |
| H | -7.313419 | -0.702744 | -0.765248 |
| O | -1.268226 | -0.368186 | -0.095430 |
| C | 2.414322 | 3.549916 | -0.078071 |
| H | 2.463281 | 3.901516 | -1.111396 |
| H | 1.719097 | 4.183338 | 0.477281 |
| H | 3.403568 | 3.637736 | 0.372946 |

Structure **$O_{Ph}$_Prod$_7$**

| | | | |
|---|---|---|---|
| C | 4.710536 | -0.412808 | -1.027970 |
| C | 4.033310 | 0.688081 | -0.233702 |
| O | 4.428599 | 1.031823 | 0.873759 |
| C | 4.228505 | -1.803001 | -0.539073 |
| C | 2.731725 | -1.884255 | -0.495653 |
| C | 1.879036 | -2.060232 | -1.551807 |
| C | 1.909496 | -1.530892 | 0.630841 |
| C | 0.581951 | -1.488042 | 0.164917 |
| C | 2.179001 | -1.194020 | 1.968635 |
| N | 0.583038 | -1.839930 | -1.155930 |
| C | -0.471126 | -1.045053 | 0.970925 |
| C | 1.126079 | -0.824241 | 2.779218 |
| C | -0.194168 | -0.744943 | 2.289814 |
| H | 4.486831 | -0.310214 | -2.092917 |
| H | 4.637234 | -1.988604 | 0.457862 |
| H | 2.100751 | -2.311534 | -2.578046 |
| H | -0.220595 | -1.818760 | -1.758822 |
| H | 3.194665 | -1.196809 | 2.345425 |
| H | 1.309180 | -0.565792 | 3.816011 |
| N | 2.926139 | 1.211651 | -0.809724 |
| C | 1.931087 | 1.941650 | -0.051550 |
| C | 0.572153 | 1.625064 | -0.685948 |
| O | 0.485742 | 1.266005 | -1.852196 |
| H | 2.578960 | 0.803556 | -1.667145 |
| H | 1.942728 | 1.563215 | 0.975946 |
| N | -0.491603 | 1.796205 | 0.125110 |
| C | -1.843625 | 1.437978 | -0.253267 |
| C | -2.428847 | 0.284217 | 0.577635 |
| H | -0.317329 | 2.032503 | 1.089516 |
| H | -1.823345 | 1.151663 | -1.305373 |
| H | -2.409770 | 0.555874 | 1.637360 |
| H | -2.504250 | 2.302768 | -0.144788 |
| H | 5.789892 | -0.323615 | -0.892624 |
| H | 4.646712 | -2.564230 | -1.201905 |
| H | -0.987592 | -0.416932 | 2.950505 |
| C | -3.859298 | 0.038665 | 0.163469 |
| C | -4.902066 | 0.627683 | 0.874014 |
| C | -4.148559 | -0.730943 | -0.962470 |
| C | -6.219804 | 0.451170 | 0.466316 |
| H | -4.683994 | 1.225260 | 1.753467 |
| C | -5.465471 | -0.913161 | -1.366087 |
| H | -3.340022 | -1.195503 | -1.514555 |
| C | -6.504001 | -0.321096 | -0.654112 |
| H | -7.024022 | 0.911784 | 1.028551 |
| H | -5.681737 | -1.518050 | -2.239379 |
| H | -7.531000 | -0.463706 | -0.970355 |
| O | -1.700745 | -0.930579 | 0.377070 |
| C | 2.190858 | 3.450509 | -0.043093 |
| H | 2.172664 | 3.843595 | -1.062345 |
| H | 1.438591 | 3.974716 | 0.550705 |
| H | 3.173026 | 3.641354 | 0.391340 |

Structure **$O_{PhOH}$•**

| | | | |
|---|---|---|---|
| C | 4.820294 | 0.243699 | -1.399845 |
| C | 4.169895 | 1.123746 | -0.356274 |
| O | 4.716810 | 1.371692 | 0.713999 |
| C | 4.750691 | -1.233182 | -0.954082 |
| C | 3.353621 | -1.751281 | -0.777159 |
| C | 2.600922 | -2.406368 | -1.711468 |
| C | 2.523624 | -1.642826 | 0.396437 |
| C | 1.290652 | -2.265080 | 0.094436 |
| C | 2.706341 | -1.083586 | 1.670511 |
| N | 1.364649 | -2.715884 | -1.196793 |
| C | 0.254240 | -2.355260 | 1.025652 |
| C | 1.675132 | -1.153529 | 2.589154 |
| C | 0.460000 | -1.789455 | 2.269872 |
| H | 4.336319 | 0.356281 | -2.373133 |
| H | 5.303101 | -1.335201 | -0.015038 |
| H | 2.857405 | -2.687007 | -2.722093 |
| H | 0.634390 | -3.197449 | -1.691951 |
| H | 3.634537 | -0.581111 | 1.918384 |
| H | -0.680788 | -2.842986 | 0.775131 |
| H | 1.803252 | -0.719331 | 3.574512 |
| N | 2.921949 | 1.543641 | -0.647007 |
| C | 2.049817 | 2.129016 | 0.349752 |

| | | | |
|---|---|---|---|
| C | 0.618369 | 1.908896 | -0.126023 |
| O | 0.317203 | 2.046894 | -1.301945 |
| H | 2.499968 | 1.299836 | -1.530080 |
| H | 2.220837 | 1.606787 | 1.294737 |
| N | -0.284446 | 1.627702 | 0.838826 |
| C | -1.677363 | 1.486114 | 0.545576 |
| C | -2.108571 | 0.014087 | 0.132972 |
| H | 0.045317 | 1.389352 | 1.761384 |
| H | -1.943062 | 2.148960 | -0.277291 |
| H | -1.783335 | -0.601099 | 0.989198 |
| H | -2.265900 | 1.740206 | 1.426288 |
| H | 5.864983 | 0.546312 | -1.496807 |
| H | 5.276128 | -1.837523 | -1.697533 |
| H | -0.327989 | -1.836727 | 3.013563 |
| C | -3.614231 | -0.028189 | -0.030834 |
| C | -4.439226 | -0.284625 | 1.059361 |
| C | -4.199217 | 0.214160 | -1.271670 |
| C | -5.821823 | -0.301312 | 0.922102 |
| H | -4.001889 | -0.480782 | 2.033849 |
| C | -5.576632 | 0.199437 | -1.424735 |
| H | -3.562555 | 0.404870 | -2.128458 |
| C | -6.394561 | -0.059176 | -0.325132 |
| H | -6.453410 | -0.507476 | 1.780335 |
| H | -6.034378 | 0.383621 | -2.389601 |
| O | -1.397201 | -0.316167 | -0.958168 |
| C | 2.299902 | 3.627579 | 0.544095 |
| H | 2.129757 | 4.161452 | -0.393329 |
| H | 1.634054 | 4.032122 | 1.309955 |
| H | 3.332364 | 3.779602 | 0.859879 |
| O | -7.736195 | -0.065740 | -0.530800 |
| H | -8.190368 | -0.261531 | 0.292059 |

Structure **$O_{PhOH}$•_TS$_7$**

| | | | |
|---|---|---|---|
| C | 4.871064 | -0.414988 | -0.927894 |
| C | 4.229414 | 0.684104 | -0.108707 |
| O | 4.604277 | 0.943840 | 1.031108 |
| C | 4.456593 | -1.797962 | -0.385151 |
| C | 2.978470 | -2.049333 | -0.436051 |
| C | 2.279441 | -2.635449 | -1.453219 |
| C | 2.003710 | -1.712640 | 0.571238 |
| C | 0.750356 | -2.141335 | 0.096789 |
| C | 2.081934 | -1.100672 | 1.830013 |
| N | 0.938962 | -2.683281 | -1.132868 |
| C | -0.457468 | -1.916499 | 0.807161 |
| C | 0.918465 | -0.963533 | 2.591712 |
| C | -0.309172 | -1.408324 | 2.119962 |
| H | 4.594063 | -0.335888 | -1.981926 |
| H | 4.808018 | -1.885006 | 0.646990 |
| H | 2.632523 | -3.042776 | -2.388034 |
| H | 0.217206 | -3.067029 | -1.719503 |
| H | 3.031243 | -0.733487 | 2.203264 |
| H | -1.359627 | -2.446633 | 0.532897 |
| H | 0.979821 | -0.521968 | 3.579543 |
| N | 3.186636 | 1.306190 | -0.691753 |
| C | 2.300195 | 2.193011 | 0.029517 |
| C | 0.922343 | 2.056373 | -0.623778 |
| O | 0.820789 | 1.789807 | -1.813063 |
| H | 2.862529 | 1.016199 | -1.603345 |
| H | 2.255329 | 1.848912 | 1.066620 |
| N | -0.122366 | 2.303609 | 0.190467 |
| C | -1.485253 | 1.979593 | -0.171393 |
| C | -1.862391 | 0.568683 | 0.343369 |
| H | 0.073104 | 2.406096 | 1.174366 |
| H | -1.567710 | 2.009007 | -1.258038 |
| H | -1.675789 | 0.593489 | 1.427594 |
| H | -2.163457 | 2.722909 | 0.253582 |
| H | 5.954748 | -0.304937 | -0.849006 |
| H | 4.980100 | -2.562645 | -0.963750 |
| H | -1.190243 | -1.314414 | 2.743663 |
| C | -3.338193 | 0.285151 | 0.122664 |
| C | -4.304350 | 0.808191 | 0.978615 |
| C | -3.759873 | -0.487716 | -0.956702 |
| C | -5.657627 | 0.576246 | 0.765527 |
| H | -4.000141 | 1.404413 | 1.834220 |
| C | -5.107321 | -0.730467 | -1.183004 |
| H | -3.012070 | -0.910663 | -1.617519 |
| C | -6.063313 | -0.195449 | -0.321962 |
| H | -6.396453 | 0.987911 | 1.446023 |
| H | -5.433720 | -1.334631 | -2.021802 |
| O | -1.018914 | -0.328182 | -0.282563 |
| C | 2.775411 | 3.648141 | -0.004353 |
| H | 2.821107 | 4.005014 | -1.035953 |
| H | 2.098019 | 4.291234 | 0.561931 |
| H | 3.770903 | 3.713696 | 0.437182 |
| O | -7.369990 | -0.462619 | -0.586354 |
| H | -7.926027 | -0.037955 | 0.070947 |

Structure **$O_{PhOH}$•_Int$_7$**

| | | | |
|---|---|---|---|
| C | 4.843689 | -0.613205 | -0.972126 |
| C | 4.255394 | 0.556627 | -0.210632 |
| O | 4.689552 | 0.900940 | 0.884383 |
| C | 4.364570 | -1.949264 | -0.361055 |
| C | 2.871869 | -2.094291 | -0.370504 |
| C | 2.097957 | -2.615054 | -1.374344 |
| C | 1.960263 | -1.629789 | 0.639709 |
| C | 0.673468 | -1.898156 | 0.188280 |
| C | 2.152902 | -1.011757 | 1.903560 |
| N | 0.771212 | -2.489370 | -1.028572 |
| C | -0.567728 | -1.394155 | 0.833142 |
| C | 1.014656 | -0.732420 | 2.686277 |
| C | -0.261050 | -0.962721 | 2.242051 |
| H | 4.562473 | -0.574903 | -2.027498 |
| H | 4.737479 | -2.016635 | 0.665092 |
| H | 2.383506 | -3.077497 | -2.306224 |
| H | -0.004396 | -2.770742 | -1.604829 |
| H | 3.146360 | -0.772794 | 2.261430 |
| H | -1.375539 | -2.138150 | 0.828435 |
| H | 1.155821 | -0.349853 | 3.692078 |
| N | 3.190832 | 1.147508 | -0.791027 |
| C | 2.320988 | 2.058178 | -0.078458 |
| C | 0.931177 | 1.922847 | -0.708958 |
| O | 0.802013 | 1.538334 | -1.862271 |
| H | 2.804674 | 0.767835 | -1.644300 |
| H | 2.277656 | 1.734870 | 0.966420 |
| N | -0.094442 | 2.308420 | 0.080280 |
| C | -1.472604 | 1.980979 | -0.228940 |
| C | -1.879324 | 0.667651 | 0.456744 |
| H | 0.117478 | 2.505506 | 1.046552 |
| H | -1.559920 | 1.872464 | -1.310425 |
| H | -1.747083 | 0.810936 | 1.535242 |
| H | -2.129140 | 2.790085 | 0.097067 |
| H | 5.931562 | -0.549026 | -0.903269 |
| H | 4.826711 | -2.765437 | -0.921654 |
| H | -1.098866 | -0.793522 | 2.908172 |
| C | -3.322812 | 0.307865 | 0.180980 |
| C | -4.343485 | 0.779400 | 1.001353 |
| C | -3.663212 | -0.471907 | -0.923633 |
| C | -5.675123 | 0.489728 | 0.729685 |
| H | -4.100950 | 1.380984 | 1.872169 |
| C | -4.986779 | -0.774233 | -1.204360 |
| H | -2.874641 | -0.854206 | -1.561626 |
| C | -6.000143 | -0.290572 | -0.378043 |
| H | -6.458913 | 0.863096 | 1.381205 |
| H | -5.251375 | -1.385023 | -2.059689 |
| O | -0.976407 | -0.297400 | -0.029316 |
| C | 2.812511 | 3.506935 | -0.144306 |
| H | 2.848335 | 3.845830 | -1.182417 |
| H | 2.152671 | 4.169426 | 0.420253 |
| H | 3.814848 | 3.565620 | 0.281862 |
| O | -7.279316 | -0.616060 | -0.697567 |
| H | -7.881819 | -0.227393 | -0.059174 |

Structure **O$_{PhOH}$_Prod$_7$**

| | | | |
|---|---|---|---|
| C | 4.982083 | -0.491524 | -1.107441 |
| C | 4.346628 | 0.627357 | -0.303892 |
| O | 4.776279 | 0.970183 | 0.791049 |
| C | 4.483345 | -1.868894 | -0.599328 |
| C | 2.986655 | -1.919394 | -0.520643 |
| C | 2.106169 | -2.084135 | -1.555650 |
| C | 2.198300 | -1.542624 | 0.622424 |
| C | 0.861441 | -1.475348 | 0.187232 |
| C | 2.505695 | -1.203416 | 1.951489 |
| N | 0.824681 | -1.834539 | -1.131226 |
| C | -0.164062 | -1.006229 | 1.014111 |
| C | 1.479526 | -0.807392 | 2.783694 |
| C | 0.150132 | -0.704106 | 2.324380 |
| H | 4.735342 | -0.390004 | -2.167398 |
| H | 4.911393 | -2.057470 | 0.388912 |
| H | 2.298725 | -2.346189 | -2.585093 |
| H | 0.006864 | -1.803679 | -1.714238 |
| H | 3.529565 | -1.224814 | 2.304777 |
| H | 1.691853 | -0.546657 | 3.814349 |
| N | 3.235962 | 1.169080 | -0.855767 |
| C | 2.273953 | 1.922441 | -0.078056 |
| C | 0.894868 | 1.633095 | -0.681526 |
| O | 0.775752 | 1.273546 | -1.844919 |
| H | 2.859496 | 0.763349 | -1.701928 |
| H | 2.300193 | 1.546510 | 0.950108 |
| N | -0.146513 | 1.828096 | 0.152327 |
| C | -1.514769 | 1.496788 | -0.192453 |
| C | -2.106057 | 0.362159 | 0.658534 |
| H | 0.055007 | 2.060286 | 1.112341 |
| H | -1.524618 | 1.207804 | -1.243921 |
| H | -2.053483 | 0.640484 | 1.715399 |
| H | -2.154007 | 2.376017 | -0.072202 |
| H | 6.065945 | -0.423873 | -0.998055 |
| H | 4.870427 | -2.642061 | -1.267233 |
| H | -0.620897 | -0.356242 | 3.001268 |
| C | -3.548404 | 0.133099 | 0.287367 |
| C | -4.574899 | 0.573425 | 1.115097 |
| C | -3.881353 | -0.492123 | -0.915717 |
| C | -5.907339 | 0.401263 | 0.758722 |
| H | -4.339179 | 1.058056 | 2.056986 |
| C | -5.204419 | -0.677448 | -1.279904 |
| H | -3.094558 | -0.849096 | -1.570739 |
| C | -6.225654 | -0.227402 | -0.442271 |
| H | -6.696444 | 0.748413 | 1.417662 |
| H | -5.464135 | -1.168105 | -2.210465 |
| O | -1.404332 | -0.870866 | 0.448723 |
| C | 2.566234 | 3.425310 | -0.080280 |
| H | 2.534271 | 3.815742 | -1.100218 |
| H | 1.838527 | 3.967332 | 0.527913 |
| H | 3.561513 | 3.596137 | 0.332069 |
| O | -7.503436 | -0.432607 | -0.848376 |
| H | -8.113253 | -0.081860 | -0.194775 |

Structure **O$_{PhO-}$•**

| | | | |
|---|---|---|---|
| C | 4.892802 | 0.352688 | -1.324787 |
| C | 4.162379 | 1.192604 | -0.300309 |
| O | 4.657485 | 1.459724 | 0.790296 |
| C | 4.845687 | -1.134192 | -0.906359 |
| C | 3.452170 | -1.670043 | -0.759403 |
| C | 2.693881 | -2.265722 | -1.728243 |
| C | 2.617680 | -1.602941 | 0.413655 |
| C | 1.374573 | -2.183345 | 0.075493 |
| C | 2.802482 | -1.102592 | 1.711741 |
| N | 1.449525 | -2.577738 | -1.234312 |
| C | 0.324993 | -2.282126 | 0.991169 |
| C | 1.760398 | -1.183818 | 2.616594 |
| C | 0.531586 | -1.771599 | 2.258456 |
| H | 4.451176 | 0.466595 | -2.317953 |
| H | 5.383829 | -1.243715 | 0.039974 |
| H | 2.949255 | -2.498037 | -2.751328 |
| H | 0.708364 | -3.008912 | -1.758746 |
| H | 3.738857 | -0.630414 | 1.986378 |
| H | -0.621499 | -2.727389 | 0.709042 |
| H | 1.888226 | -0.792535 | 3.619806 |
| N | 2.906040 | 1.550643 | -0.635478 |
| C | 1.959141 | 2.059744 | 0.333554 |
| C | 0.562075 | 1.801777 | -0.224096 |
| O | 0.336041 | 1.882929 | -1.421924 |
| H | 2.525687 | 1.279324 | -1.529777 |
| H | 2.099143 | 1.508381 | 1.267697 |
| N | -0.393283 | 1.565555 | 0.700532 |
| C | -1.777127 | 1.426340 | 0.363760 |
| C | -2.252268 | -0.078536 | 0.084213 |
| H | -0.109627 | 1.383395 | 1.650765 |
| H | -1.993144 | 2.005624 | -0.532726 |
| H | -1.916723 | -0.604080 | 0.999017 |
| H | -2.390290 | 1.781393 | 1.190558 |
| H | 5.930625 | 0.689289 | -1.367103 |
| H | 5.393309 | -1.715839 | -1.651894 |
| H | -0.266777 | -1.824704 | 2.990572 |
| C | -3.753107 | -0.095689 | -0.040497 |
| C | -4.568772 | -0.173803 | 1.093745 |
| C | -4.384507 | 0.013033 | -1.283170 |
| C | -5.948347 | -0.154757 | 1.001666 |
| H | -4.103252 | -0.258219 | 2.075415 |
| C | -5.762158 | 0.032671 | -1.397485 |
| H | -3.767509 | 0.072784 | -2.176406 |
| C | -6.636603 | -0.048328 | -0.257574 |
| H | -6.556800 | -0.226903 | 1.899995 |
| H | -6.227486 | 0.112222 | -2.377140 |
| O | -1.542820 | -0.495079 | -0.971331 |
| C | 2.135602 | 3.559252 | 0.595241 |
| H | 1.985492 | 4.122182 | -0.328778 |
| H | 1.419553 | 3.906796 | 1.343420 |
| H | 3.145478 | 3.741038 | 0.963672 |
| O | -7.900359 | -0.028975 | -0.355022 |

Structure **O$_{PhO-}$•_TS$_7$**

| | | | |
|---|---|---|---|
| C | 4.828569 | -0.418201 | -0.944188 |
| C | 4.194952 | 0.663379 | -0.095434 |
| O | 4.577589 | 0.894583 | 1.048090 |
| C | 4.409894 | -1.813161 | -0.435834 |
| C | 2.930092 | -2.054001 | -0.486928 |
| C | 2.221486 | -2.602131 | -1.518913 |
| C | 1.962650 | -1.734537 | 0.532546 |
| C | 0.703213 | -2.132114 | 0.049699 |
| C | 2.052736 | -1.157167 | 1.808455 |
| N | 0.881701 | -2.642441 | -1.196098 |
| C | -0.502391 | -1.900724 | 0.763888 |
| C | 0.892996 | -1.021860 | 2.573403 |
| C | -0.343486 | -1.433234 | 2.090399 |
| H | 4.547756 | -0.311274 | -1.994792 |
| H | 4.765541 | -1.928721 | 0.592119 |
| H | 2.567308 | -2.985824 | -2.466437 |
| H | 0.152577 | -2.994411 | -1.793183 |
| H | 3.008134 | -0.813926 | 2.188860 |
| H | -1.411308 | -2.414605 | 0.480643 |
| H | 0.961266 | -0.607661 | 3.572782 |
| N | 3.152285 | 1.303583 | -0.658354 |
| C | 2.271751 | 2.177137 | 0.086256 |
| C | 0.895306 | 2.076928 | -0.578532 |
| O | 0.799735 | 1.841601 | -1.775632 |
| H | 2.822137 | 1.036616 | -1.574903 |
| H | 2.216511 | 1.800894 | 1.111666 |
| N | -0.150520 | 2.319962 | 0.233935 |
| C | -1.515494 | 1.999274 | -0.133594 |
| C | -1.903294 | 0.582517 | 0.359058 |
| H | 0.043374 | 2.387843 | 1.221302 |
| H | -1.598729 | 2.050621 | -1.219536 |
| H | -1.718586 | 0.597432 | 1.444556 |
| H | -2.193370 | 2.733955 | 0.306535 |

| | | | |
|---|---|---|---|
| H | 5.913057 | -0.314414 | -0.866908 |
| H | 4.927450 | -2.564768 | -1.036678 |
| H | -1.223101 | -1.338017 | 2.716033 |
| C | -3.367052 | 0.288087 | 0.127452 |
| C | -4.323975 | 0.497964 | 1.124697 |
| C | -3.828049 | -0.175912 | -1.109501 |
| C | -5.672874 | 0.265575 | 0.909482 |
| H | -3.996530 | 0.852197 | 2.101962 |
| C | -5.169927 | -0.419034 | -1.344621 |
| H | -3.101655 | -0.361022 | -1.897643 |
| C | -6.183856 | -0.210937 | -0.345977 |
| H | -6.389780 | 0.433317 | 1.710180 |
| H | -5.495733 | -0.786358 | -2.315507 |
| O | -1.028910 | -0.292312 | -0.273995 |
| C | 2.763931 | 3.626949 | 0.100762 |
| H | 2.817683 | 4.015880 | -0.918845 |
| H | 2.091509 | 4.259522 | 0.684555 |
| H | 3.758351 | 3.667336 | 0.547729 |
| O | -7.417314 | -0.432764 | -0.554251 |

Structure **$O_{PhO-}$•_Int$_7$**

| | | | |
|---|---|---|---|
| C | 4.776526 | -0.654944 | -0.984461 |
| C | 4.205356 | 0.475422 | -0.152557 |
| O | 4.631743 | 0.730848 | 0.969530 |
| C | 4.265465 | -2.018252 | -0.465990 |
| C | 2.769765 | -2.122006 | -0.493429 |
| C | 1.988110 | -2.525309 | -1.544500 |
| C | 1.865629 | -1.705788 | 0.544042 |
| C | 0.574790 | -1.879615 | 0.059372 |
| C | 2.067514 | -1.195627 | 1.854547 |
| N | 0.663605 | -2.372631 | -1.201256 |
| C | -0.649513 | -1.350752 | 0.719007 |
| C | 0.931329 | -0.919430 | 2.641817 |
| C | -0.346546 | -1.049160 | 2.163956 |
| H | 4.504994 | -0.542281 | -2.037077 |
| H | 4.630064 | -2.162188 | 0.555193 |
| H | 2.266251 | -2.918612 | -2.509818 |
| H | -0.116099 | -2.563446 | -1.808160 |
| H | 3.065483 | -1.030652 | 2.240563 |
| H | -1.501529 | -2.041036 | 0.644274 |
| H | 1.075037 | -0.619787 | 3.675230 |
| N | 3.166880 | 1.135218 | -0.705607 |
| C | 2.314171 | 2.022795 | 0.055758 |
| C | 0.937189 | 2.003888 | -0.618142 |
| O | 0.821562 | 1.698774 | -1.796658 |
| H | 2.783304 | 0.825583 | -1.587725 |
| H | 2.222832 | 1.616127 | 1.068079 |
| N | -0.087405 | 2.397297 | 0.168960 |
| C | -1.471551 | 2.099620 | -0.160823 |
| C | -1.899209 | 0.762624 | 0.460599 |
| H | 0.116829 | 2.511583 | 1.150507 |
| H | -1.555247 | 2.052650 | -1.246890 |
| H | -1.771690 | 0.859932 | 1.546410 |
| H | -2.117733 | 2.899608 | 0.205285 |
| H | 5.864964 | -0.616757 | -0.903961 |
| H | 4.712602 | -2.805568 | -1.077826 |
| H | -1.187107 | -0.878402 | 2.826386 |
| C | -3.325722 | 0.386389 | 0.173042 |
| C | -4.275040 | 0.280493 | 1.193757 |
| C | -3.757793 | 0.114187 | -1.132637 |
| C | -5.591475 | -0.065482 | 0.938385 |
| H | -3.967095 | 0.476063 | 2.220117 |
| C | -5.065200 | -0.236065 | -1.409012 |
| H | -3.038371 | 0.168631 | -1.947230 |
| C | -6.072240 | -0.347621 | -0.385616 |
| H | -6.304519 | -0.140725 | 1.756014 |
| H | -5.369777 | -0.443490 | -2.432405 |
| O | -0.958241 | -0.170535 | -0.053586 |
| C | 2.870791 | 3.447682 | 0.127325 |
| H | 2.953973 | 3.870507 | -0.876784 |
| H | 2.223110 | 4.090552 | 0.727537 |
| H | 3.860600 | 3.424034 | 0.584930 |
| O | -7.273377 | -0.670315 | -0.633529 |

Structure **$O_{PhO-}$-Prod$_7$**

| | | | |
|---|---|---|---|
| C | 4.958637 | -0.451132 | -1.111549 |
| C | 4.324841 | 0.617085 | -0.240301 |
| O | 4.751315 | 0.885572 | 0.876622 |
| C | 4.448595 | -1.855578 | -0.697392 |
| C | 2.951054 | -1.899865 | -0.633450 |
| C | 2.079494 | -1.986370 | -1.685864 |
| C | 2.155291 | -1.588662 | 0.524270 |
| C | 0.824661 | -1.477453 | 0.081620 |
| C | 2.451511 | -1.336489 | 1.875804 |
| N | 0.797161 | -1.750058 | -1.258021 |
| C | -0.209011 | -1.045791 | 0.920868 |
| C | 1.419280 | -0.979257 | 2.717871 |
| C | 0.096149 | -0.829725 | 2.251365 |
| H | 4.718124 | -0.279358 | -2.163884 |
| H | 4.868246 | -2.110743 | 0.279455 |
| H | 2.280275 | -2.182361 | -2.728415 |
| H | -0.015241 | -1.668610 | -1.843706 |
| H | 3.470733 | -1.391611 | 2.239006 |
| H | 1.622267 | -0.784958 | 3.765172 |
| N | 3.220760 | 1.200322 | -0.761709 |
| C | 2.258495 | 1.908366 | 0.057503 |
| C | 0.882034 | 1.678339 | -0.578155 |
| O | 0.773549 | 1.403332 | -1.766299 |
| H | 2.845650 | 0.850562 | -1.633211 |
| H | 2.270335 | 1.461840 | 1.057241 |
| N | -0.164350 | 1.828291 | 0.256222 |
| C | -1.537575 | 1.542066 | -0.118331 |
| C | -2.157550 | 0.357729 | 0.637704 |
| H | 0.032890 | 1.979250 | 1.233261 |
| H | -1.544168 | 1.342358 | -1.189852 |
| H | -2.101950 | 0.555345 | 1.712918 |
| H | -2.161525 | 2.420011 | 0.069920 |
| H | 6.042282 | -0.397042 | -0.992380 |
| H | 4.836179 | -2.586043 | -1.411596 |
| H | -0.677390 | -0.512353 | 2.940272 |
| C | -3.593408 | 0.155007 | 0.259469 |
| C | -4.614053 | 0.243081 | 1.211068 |
| C | -3.970311 | -0.133764 | -1.059921 |
| C | -5.944860 | 0.065827 | 0.876145 |
| H | -4.352413 | 0.458972 | 2.245602 |
| C | -5.291712 | -0.323393 | -1.412973 |
| H | -3.202555 | -0.224644 | -1.825290 |
| C | -6.371087 | -0.230565 | -0.464197 |
| H | -6.714167 | 0.144038 | 1.640531 |
| H | -5.552007 | -0.552339 | -2.443590 |
| O | -1.433404 | -0.863210 | 0.345968 |
| C | 2.566772 | 3.404311 | 0.163268 |
| H | 2.546900 | 3.866217 | -0.826730 |
| H | 1.839396 | 3.909492 | 0.802854 |
| H | 3.560321 | 3.535382 | 0.594069 |
| O | -7.585291 | -0.398932 | -0.784546 |

Structure **$O_{Ph}$•$_{6a}$**

| | | | |
|---|---|---|---|
| C | -2.881810 | 0.405808 | 2.335702 |
| C | -2.495276 | -0.805081 | 1.512841 |
| O | -1.646509 | -1.596897 | 1.905357 |
| C | -1.687274 | 1.365363 | 2.450225 |
| C | -1.261420 | 1.964893 | 1.139729 |
| C | -2.060487 | 2.407833 | 0.121407 |
| C | 0.090782 | 2.213235 | 0.716394 |
| C | 0.027727 | 2.806927 | -0.567908 |
| C | 1.347290 | 1.989173 | 1.299724 |
| N | -1.296612 | 2.920659 | -0.897123 |
| C | 1.174130 | 3.168039 | -1.278419 |
| C | 2.483294 | 2.339993 | 0.597877 |
| C | 2.397166 | 2.921399 | -0.681917 |
| H | -3.752115 | 0.915838 | 1.915694 |

| | | | |
|---|---|---|---|
| H | -0.847449 | 0.812165 | 2.879690 |
| H | -3.136317 | 2.406967 | 0.034375 |
| H | -1.654758 | 3.243048 | -1.779132 |
| H | 1.425451 | 1.529677 | 2.278899 |
| H | 1.107391 | 3.610836 | -2.265292 |
| H | 3.460217 | 2.155839 | 1.030031 |
| N | -3.124086 | -0.947770 | 0.321655 |
| C | -2.684489 | -1.915380 | -0.667133 |
| C | -1.598767 | -1.263698 | -1.532886 |
| O | -1.819621 | -0.227684 | -2.140862 |
| H | -3.695910 | -0.187346 | -0.010695 |
| H | -2.273127 | -2.762273 | -0.117292 |
| N | -0.411855 | -1.910118 | -1.582118 |
| C | 0.715985 | -1.361395 | -2.287605 |
| C | 1.566121 | -0.383034 | -1.394015 |
| H | -0.266209 | -2.707328 | -0.983861 |
| H | 0.354633 | -0.794891 | -3.143830 |
| H | 0.822241 | 0.356446 | -1.036013 |
| H | 1.358530 | -2.173133 | -2.628940 |
| H | -3.157419 | 0.051023 | 3.331446 |
| H | -1.940188 | 2.151495 | 3.168629 |
| O | 2.421666 | 0.271794 | -2.209093 |
| H | 3.307836 | 3.179827 | -1.209538 |
| C | -3.850410 | -2.363694 | -1.539371 |
| H | -3.514564 | -3.103665 | -2.268513 |
| H | -4.630855 | -2.808758 | -0.920354 |
| H | -4.267815 | -1.511693 | -2.081387 |
| C | 2.206959 | -1.092330 | -0.218602 |
| C | 3.539858 | -1.489751 | -0.283367 |
| C | 1.469621 | -1.366141 | 0.931776 |
| C | 4.129715 | -2.152497 | 0.787077 |
| H | 4.116663 | -1.263779 | -1.173278 |
| C | 2.056494 | -2.033393 | 2.000654 |
| H | 0.433368 | -1.055602 | 1.006591 |
| C | 3.388872 | -2.427359 | 1.932330 |
| H | 5.169642 | -2.454174 | 0.727687 |
| H | 1.470935 | -2.237836 | 2.890175 |
| H | 3.848918 | -2.943233 | 2.767712 |

Structure **$O_{Ph}$•_$TS_{6a}$**

| | | | |
|---|---|---|---|
| C | -3.138661 | -0.357685 | 2.162068 |
| C | -2.231756 | -1.302737 | 1.397661 |
| O | -1.142406 | -1.640523 | 1.845853 |
| C | -2.433118 | 0.985225 | 2.412391 |
| C | -2.044081 | 1.679693 | 1.138537 |
| C | -2.873278 | 1.964107 | 0.072283 |
| C | -0.747167 | 2.112643 | 0.753476 |
| C | -0.843687 | 2.630358 | -0.569347 |
| C | 0.517675 | 2.127817 | 1.401967 |
| N | -2.165201 | 2.545454 | -0.936830 |
| C | 0.249739 | 3.093326 | -1.270603 |
| C | 1.613357 | 2.562647 | 0.716913 |
| C | 1.530547 | 2.931454 | -0.674461 |
| H | -4.082688 | -0.201511 | 1.634552 |
| H | -1.536827 | 0.798666 | 3.007774 |
| H | -3.931579 | 1.783072 | -0.039654 |
| H | -2.535257 | 2.776504 | -1.842784 |
| H | 0.606521 | 1.796417 | 2.430520 |
| H | 0.161676 | 3.484180 | -2.276325 |
| H | 2.587231 | 2.568232 | 1.190594 |
| N | -2.699810 | -1.746364 | 0.207930 |
| C | -1.882481 | -2.516486 | -0.713128 |
| C | -1.084548 | -1.538758 | -1.587999 |
| O | -1.654638 | -0.628052 | -2.171019 |
| H | -3.526714 | -1.312509 | -0.171246 |
| H | -1.205415 | -3.120421 | -0.107564 |
| N | 0.245525 | -1.762753 | -1.669565 |
| C | 1.143726 | -0.749601 | -2.194881 |
| C | 1.582907 | 0.221838 | -1.072864 |
| H | 0.641450 | -2.463273 | -1.061829 |
| H | 0.603617 | -0.179461 | -2.948985 |
| H | 0.649697 | 0.455927 | -0.539589 |
| H | 2.006325 | -1.230694 | -2.660293 |
| H | -3.373014 | -0.830403 | 3.119546 |
| H | -3.089628 | 1.616983 | 3.017145 |
| O | 2.127260 | 1.363389 | -1.632517 |
| H | 2.357155 | 3.487510 | -1.094991 |
| C | -2.757211 | -3.411125 | -1.583463 |
| H | -2.136578 | -4.002332 | -2.259518 |
| H | -3.341561 | -4.088904 | -0.959164 |
| H | -3.439084 | -2.804989 | -2.185196 |
| C | 2.519379 | -0.465379 | -0.090726 |
| C | 3.899075 | -0.387183 | -0.273695 |
| C | 2.021301 | -1.203301 | 0.983206 |
| C | 4.767207 | -1.039041 | 0.594230 |
| H | 4.284096 | 0.204381 | -1.096669 |
| C | 2.889347 | -1.859506 | 1.851107 |
| H | 0.950126 | -1.253959 | 1.156299 |
| C | 4.264499 | -1.780911 | 1.659431 |
| H | 5.838754 | -0.967094 | 0.441447 |
| H | 2.488842 | -2.425699 | 2.685350 |
| H | 4.940901 | -2.287860 | 2.338694 |

Structure **$O_{Ph}$•_$Int_{6a}$**

| | | | |
|---|---|---|---|
| C | 3.152309 | 0.741129 | 2.117116 |
| C | 2.066679 | 1.481530 | 1.357356 |
| O | 0.929643 | 1.584020 | 1.802472 |
| C | 2.746852 | -0.721764 | 2.373639 |
| C | 2.401786 | -1.437783 | 1.099658 |
| C | 3.205884 | -1.540809 | -0.023691 |
| C | 1.157549 | -2.008535 | 0.752185 |
| C | 1.227679 | -2.405397 | -0.616229 |
| C | -0.058821 | -2.252634 | 1.478483 |
| N | 2.518475 | -2.154596 | -1.032118 |
| C | 0.153950 | -2.888751 | -1.316516 |
| C | -1.149270 | -2.705142 | 0.828376 |
| C | -1.196449 | -2.851558 | -0.670692 |
| H | 4.102174 | 0.780935 | 1.578559 |
| H | 1.880108 | -0.733460 | 3.037590 |
| H | 4.219941 | -1.204509 | -0.176281 |
| H | 2.829141 | -2.196669 | -1.987238 |
| H | -0.089435 | -2.079424 | 2.549456 |
| H | 0.221351 | -3.157661 | -2.364041 |
| H | -2.074397 | -2.888397 | 1.363737 |
| N | 2.438130 | 2.024303 | 0.173654 |
| C | 1.488551 | 2.627874 | -0.744044 |
| C | 0.851391 | 1.517005 | -1.593405 |
| O | 1.552357 | 0.682903 | -2.143843 |
| H | 3.334837 | 1.765298 | -0.206340 |
| H | 0.726921 | 3.120420 | -0.137609 |
| N | -0.499198 | 1.544420 | -1.689905 |
| C | -1.240761 | 0.366307 | -2.107773 |
| C | -1.591519 | -0.473666 | -0.867234 |
| H | -0.991190 | 2.209829 | -1.112740 |
| H | -0.595411 | -0.221188 | -2.758882 |
| H | -0.670639 | -0.522924 | -0.283804 |
| H | -2.135274 | 0.661341 | -2.660334 |
| H | 3.296133 | 1.255114 | 3.071442 |
| H | 3.562660 | -1.220035 | 2.904478 |
| O | -2.000690 | -1.772173 | -1.257387 |
| H | -1.778333 | -3.734516 | -0.945642 |
| C | 2.189150 | 3.641548 | -1.641502 |
| H | 1.469807 | 4.107860 | -2.317315 |
| H | 2.659295 | 4.419119 | -1.037395 |
| H | 2.955576 | 3.147278 | -2.244010 |
| C | -2.649470 | 0.197078 | -0.016618 |
| C | -4.003675 | -0.009919 | -0.277644 |
| C | -2.279639 | 1.062234 | 1.012959 |
| C | -4.973518 | 0.640445 | 0.474883 |
| H | -4.291416 | -0.695838 | -1.066136 |
| C | -3.252113 | 1.718279 | 1.762851 |

| | | | |
|---|---|---|---|
| H | -1.227258 | 1.213393 | 1.238458 |
| C | -4.600217 | 1.510066 | 1.496150 |
| H | -6.023716 | 0.468637 | 0.265733 |
| H | -2.953635 | 2.386425 | 2.563365 |
| H | -5.357536 | 2.016852 | 2.083740 |

Structure **$O_{Ph}$•$_{6b}$**

| | | | |
|---|---|---|---|
| C | -2.903645 | 0.606829 | 2.214870 |
| C | -2.364508 | -0.606523 | 1.488819 |
| O | -1.302317 | -1.131513 | 1.820914 |
| C | -1.832719 | 1.702029 | 2.269004 |
| C | -1.355282 | 2.137411 | 0.913116 |
| C | -2.077958 | 2.238161 | -0.244070 |
| C | -0.013555 | 2.526141 | 0.576014 |
| C | -0.001916 | 2.855833 | -0.800636 |
| C | 1.185459 | 2.602248 | 1.304866 |
| N | -1.274060 | 2.667278 | -1.272925 |
| C | 1.160247 | 3.263553 | -1.457395 |
| C | 2.339190 | 2.998122 | 0.655392 |
| C | 2.325164 | 3.327115 | -0.713591 |
| H | -3.817078 | 0.976827 | 1.743009 |
| H | -0.980685 | 1.318626 | 2.835975 |
| H | -3.121433 | 2.034456 | -0.429033 |
| H | -1.574583 | 2.808181 | -2.221990 |
| H | 1.206047 | 2.347309 | 2.358963 |
| H | 1.153465 | 3.503746 | -2.513876 |
| H | 3.272875 | 3.052471 | 1.203079 |
| N | -3.091653 | -1.058703 | 0.448276 |
| C | -2.673531 | -2.176108 | -0.390575 |
| C | -1.689223 | -1.680589 | -1.471057 |
| O | -1.963228 | -1.676125 | -2.662411 |
| H | -3.922925 | -0.557560 | 0.177280 |
| H | -2.133615 | -2.870555 | 0.258079 |
| N | -0.511454 | -1.246724 | -0.983703 |
| C | 0.503807 | -0.683179 | -1.847345 |
| C | 1.695941 | -0.145986 | -1.034042 |
| H | -0.378480 | -1.271734 | 0.021874 |
| H | 0.090959 | 0.147148 | -2.426661 |
| H | 1.301846 | 0.680569 | -0.406039 |
| H | 0.863974 | -1.440767 | -2.551928 |
| H | -3.165790 | 0.299270 | 3.230660 |
| H | -2.226541 | 2.552345 | 2.834355 |
| O | 2.588902 | 0.466932 | -1.856342 |
| H | 3.246994 | 3.630235 | -1.195811 |
| C | -3.882842 | -2.860975 | -0.999635 |
| H | -3.569127 | -3.716898 | -1.597576 |
| H | -4.549888 | -3.208964 | -0.209056 |
| H | -4.428046 | -2.177107 | -1.653793 |
| C | 2.323972 | -1.164667 | -0.101532 |
| C | 2.001903 | -1.186683 | 1.253247 |
| C | 3.226112 | -2.101469 | -0.605526 |
| C | 2.567290 | -2.139476 | 2.094686 |
| H | 1.295912 | -0.465226 | 1.652404 |
| C | 3.797153 | -3.047721 | 0.235208 |
| H | 3.483804 | -2.085413 | -1.659510 |
| C | 3.466979 | -3.069465 | 1.587623 |
| H | 2.306969 | -2.151020 | 3.146947 |
| H | 4.499528 | -3.770731 | -0.163855 |
| H | 3.910877 | -3.810124 | 2.243021 |

Structure **$O_{Ph}$•_TS$_{6b}$**

| | | | |
|---|---|---|---|
| C | 3.236889 | 0.097920 | 2.229557 |
| C | 2.273252 | 1.068109 | 1.576936 |
| O | 1.112222 | 1.182834 | 1.966751 |
| C | 2.659670 | -1.326223 | 2.213493 |
| C | 2.261659 | -1.762038 | 0.832619 |
| C | 2.995924 | -1.614314 | -0.326790 |
| C | 1.010092 | -2.304634 | 0.439243 |
| C | 1.028103 | -2.442939 | -0.973371 |
| C | -0.155525 | -2.700277 | 1.152802 |
| N | 2.265843 | -2.027414 | -1.402709 |
| C | -0.065386 | -2.882400 | -1.692358 |
| C | -1.249560 | -3.126446 | 0.458401 |
| C | -1.278048 | -3.091273 | -0.979597 |
| H | 4.210160 | 0.122346 | 1.733711 |
| H | 1.778507 | -1.351559 | 2.857705 |
| H | 3.995283 | -1.230534 | -0.463819 |
| H | 2.571357 | -1.988144 | -2.360242 |
| H | -0.165521 | -2.673749 | 2.236832 |
| H | -0.043129 | -2.973625 | -2.770924 |
| H | -2.146814 | -3.431148 | 0.983053 |
| N | 2.757261 | 1.774626 | 0.535153 |
| C | 1.970178 | 2.749881 | -0.212441 |
| C | 1.120876 | 2.030726 | -1.282467 |
| O | 1.371375 | 2.094042 | -2.477872 |
| H | 3.698790 | 1.598346 | 0.222232 |
| H | 1.286402 | 3.216285 | 0.501279 |
| N | 0.091554 | 1.329738 | -0.774571 |
| C | -0.762078 | 0.483654 | -1.590247 |
| C | -1.688380 | -0.348396 | -0.695482 |
| H | 0.037078 | 1.261590 | 0.236989 |
| H | -0.149027 | -0.204889 | -2.174692 |
| H | -1.066421 | -0.694231 | 0.147990 |
| H | -1.337732 | 1.090974 | -2.297370 |
| H | 3.390454 | 0.422563 | 3.262028 |
| H | 3.393571 | -2.008193 | 2.651679 |
| O | -2.211060 | -1.441953 | -1.370256 |
| H | -2.069445 | -3.631642 | -1.480317 |
| C | 2.880147 | 3.797648 | -0.825987 |
| H | 2.288820 | 4.550149 | -1.348611 |
| H | 3.463257 | 4.288965 | -0.045099 |
| H | 3.559523 | 3.342864 | -1.549928 |
| C | -2.837112 | 0.436738 | -0.060055 |
| C | -3.800057 | -0.275973 | 0.658423 |
| C | -2.961378 | 1.821666 | -0.147887 |
| C | -4.854669 | 0.377210 | 1.279514 |
| H | -3.715336 | -1.354911 | 0.714859 |
| C | -4.021861 | 2.479892 | 0.472114 |
| H | -2.226939 | 2.405025 | -0.690789 |
| C | -4.970624 | 1.762393 | 1.187455 |
| H | -5.590869 | -0.193692 | 1.834784 |
| H | -4.102299 | 3.558379 | 0.391747 |
| H | -5.795247 | 2.275265 | 1.669553 |

Structure **$O_{Ph}$•_Int$_{6b}$**

| | | | |
|---|---|---|---|
| C | -3.549626 | 0.217327 | -2.009278 |
| C | -2.422492 | 1.090861 | -1.496157 |
| O | -1.304134 | 1.072123 | -2.007886 |
| C | -3.149056 | -1.267669 | -1.986088 |
| C | -2.628340 | -1.677720 | -0.639366 |
| C | -3.261453 | -1.492667 | 0.578967 |
| C | -1.342508 | -2.182671 | -0.349045 |
| C | -1.213675 | -2.262350 | 1.066294 |
| C | -0.249975 | -2.629599 | -1.169138 |
| N | -2.421961 | -1.857494 | 1.594417 |
| C | -0.054400 | -2.634309 | 1.697303 |
| C | 0.921369 | -2.971043 | -0.597441 |
| C | 1.186857 | -2.791356 | 0.873371 |
| H | -4.456823 | 0.372065 | -1.420662 |
| H | -2.371866 | -1.435208 | -2.734145 |
| H | -4.247688 | -1.110308 | 0.792704 |
| H | -2.630537 | -1.775203 | 2.574653 |
| H | -0.380663 | -2.705694 | -2.243788 |
| H | 0.029314 | -2.675309 | 2.776853 |
| H | 1.754132 | -3.310329 | -1.203956 |
| N | -2.716940 | 1.876414 | -0.439552 |
| C | -1.776135 | 2.824506 | 0.147362 |
| C | -0.862935 | 2.114261 | 1.168878 |
| O | -0.951987 | 2.294916 | 2.374696 |
| H | -3.630765 | 1.807026 | -0.020608 |
| H | -1.144180 | 3.184285 | -0.668806 |
| N | 0.038330 | 1.286108 | 0.608238 |

| | | | |
|---|---|---|---|
| C | 0.922587 | 0.449735 | 1.401832 |
| C | 1.763646 | -0.441617 | 0.474713 |
| H | -0.049470 | 1.119540 | -0.389826 |
| H | 0.333961 | -0.188486 | 2.061805 |
| H | 1.148607 | -0.653814 | -0.408911 |
| H | 1.560932 | 1.076009 | 2.032591 |
| H | -3.769640 | 0.527773 | -3.034360 |
| H | -4.013716 | -1.867035 | -2.283277 |
| O | 2.106783 | -1.667247 | 1.092331 |
| H | 1.784013 | -3.626063 | 1.248556 |
| C | -2.526174 | 3.983988 | 0.777249 |
| H | -1.822494 | 4.716663 | 1.173451 |
| H | -3.155852 | 4.469881 | 0.029864 |
| H | -3.150850 | 3.637973 | 1.603598 |
| C | 3.039464 | 0.214720 | -0.029616 |
| C | 4.106753 | -0.580846 | -0.447559 |
| C | 3.154863 | 1.600896 | -0.142268 |
| C | 5.262408 | -0.005847 | -0.962162 |
| H | 4.027554 | -1.656713 | -0.355170 |
| C | 4.310796 | 2.177117 | -0.658829 |
| H | 2.337356 | 2.243081 | 0.165525 |
| C | 5.369838 | 1.376717 | -1.069924 |
| H | 6.082906 | -0.640637 | -1.278277 |
| H | 4.381957 | 3.256270 | -0.737169 |
| H | 6.271646 | 1.826090 | -1.470093 |

Structure $\mathbf{O_{Ph}\bullet_{6c}}$

| | | | |
|---|---|---|---|
| C | 4.941552 | -0.015139 | -0.694957 |
| C | 4.087136 | 0.978804 | 0.059071 |
| O | 4.323434 | 1.295229 | 1.220186 |
| C | 4.562420 | -1.452493 | -0.274402 |
| C | 3.120468 | -1.800558 | -0.504871 |
| C | 2.579126 | -2.363282 | -1.626854 |
| C | 2.023845 | -1.601785 | 0.408086 |
| C | 0.856814 | -2.085836 | -0.227292 |
| C | 1.921906 | -1.069159 | 1.702247 |
| N | 1.224093 | -2.533184 | -1.469365 |
| C | -0.389514 | -2.084361 | 0.404496 |
| C | 0.686710 | -1.046561 | 2.322465 |
| C | -0.457084 | -1.560115 | 1.682143 |
| H | 4.812860 | 0.096047 | -1.774528 |
| H | 4.800671 | -1.575776 | 0.786423 |
| H | 3.064599 | -2.670576 | -2.541004 |
| H | 0.609524 | -2.946151 | -2.149348 |
| H | 2.797815 | -0.663765 | 2.196737 |
| H | -1.270667 | -2.479529 | -0.088691 |
| H | 0.592876 | -0.633184 | 3.320358 |
| N | 3.005784 | 1.425426 | -0.614574 |
| C | 1.885196 | 2.055411 | 0.048938 |
| C | 0.624166 | 1.568939 | -0.662545 |
| O | 0.629053 | 1.315275 | -1.860968 |
| H | 2.804328 | 1.050139 | -1.530761 |
| H | 1.878967 | 1.718278 | 1.088589 |
| N | -0.480201 | 1.499500 | 0.100364 |
| C | -1.724261 | 0.999132 | -0.431204 |
| C | -2.849390 | 1.173643 | 0.614278 |
| H | -0.384257 | 1.481269 | 1.105821 |
| H | -1.643372 | -0.065128 | -0.673727 |
| H | -3.026653 | 2.268661 | 0.699103 |
| H | -1.987153 | 1.535469 | -1.343959 |
| H | 5.988313 | 0.173795 | -0.450033 |
| H | 5.202974 | -2.146649 | -0.823753 |
| H | -1.408225 | -1.541042 | 2.201574 |
| C | -4.122824 | 0.492799 | 0.157674 |
| C | -4.455307 | -0.767140 | 0.648529 |
| C | -4.944934 | 1.099546 | -0.789264 |
| C | -5.599935 | -1.413782 | 0.197846 |
| H | -3.818055 | -1.238311 | 1.388608 |
| C | -6.089873 | 0.453226 | -1.240377 |
| H | -4.692609 | 2.083478 | -1.172598 |
| C | -6.418891 | -0.805154 | -0.747609 |
| H | -5.852288 | -2.394552 | 0.584552 |
| H | -6.727144 | 0.934076 | -1.973594 |
| H | -7.311695 | -1.309685 | -1.098718 |
| O | -2.432200 | 0.847749 | 1.864628 |
| C | 1.962544 | 3.583887 | 0.009962 |
| H | 2.880168 | 3.911539 | 0.500961 |
| H | 1.970421 | 3.936174 | -1.024199 |
| H | 1.109355 | 4.027068 | 0.528285 |

Structure $\mathbf{O_{Ph}\bullet\_TS_{6c}}$

| | | | |
|---|---|---|---|
| C | 4.638066 | -0.931365 | -0.477965 |
| C | 4.047714 | 0.217577 | 0.307839 |
| O | 4.153107 | 0.290971 | 1.526940 |
| C | 3.788685 | -2.196148 | -0.238556 |
| C | 2.335909 | -2.035085 | -0.584030 |
| C | 1.759728 | -2.222869 | -1.822971 |
| C | 1.272144 | -1.690549 | 0.295493 |
| C | 0.068583 | -1.717933 | -0.456709 |
| C | 1.202859 | -1.407447 | 1.686741 |
| N | 0.410626 | -2.026841 | -1.751919 |
| C | -1.177100 | -1.544926 | 0.117656 |
| C | -0.012096 | -1.196343 | 2.264458 |
| C | -1.236281 | -1.210385 | 1.496390 |
| H | 4.674003 | -0.704374 | -1.546199 |
| H | 3.876996 | -2.477502 | 0.814113 |
| H | 2.225684 | -2.499757 | -2.756567 |
| H | -0.232580 | -2.135676 | -2.517637 |
| H | 2.112945 | -1.366764 | 2.274116 |
| H | -2.089352 | -1.655279 | -0.455933 |
| H | -0.084780 | -0.986464 | 3.324621 |
| N | 3.334895 | 1.101752 | -0.422862 |
| C | 2.404881 | 2.041170 | 0.173641 |
| C | 1.108908 | 1.929746 | -0.628164 |
| O | 1.126990 | 1.991091 | -1.853797 |
| H | 3.222900 | 0.945164 | -1.413616 |
| H | 2.246816 | 1.722009 | 1.205342 |
| N | -0.008200 | 1.781846 | 0.092781 |
| C | -1.296754 | 1.546796 | -0.516395 |
| C | -2.309498 | 1.296392 | 0.619920 |
| H | 0.046100 | 1.510849 | 1.067372 |
| H | -1.237561 | 0.685162 | -1.188464 |
| H | -2.587314 | 2.289165 | 1.015198 |
| H | -1.594086 | 2.412044 | -1.114966 |
| H | 5.657688 | -1.105489 | -0.129115 |
| H | 4.217753 | -3.014309 | -0.821638 |
| H | -2.153329 | -1.448052 | 2.017593 |
| C | -3.598853 | 0.642646 | 0.141246 |
| C | -4.407063 | -0.012113 | 1.071318 |
| C | -4.019805 | 0.697287 | -1.187251 |
| C | -5.596676 | -0.613943 | 0.682743 |
| H | -4.085386 | -0.051550 | 2.105523 |
| C | -5.212897 | 0.097055 | -1.578875 |
| H | -3.419976 | 1.203657 | -1.934889 |
| C | -6.003746 | -0.563637 | -0.647033 |
| H | -6.207747 | -1.123787 | 1.419329 |
| H | -5.520993 | 0.144599 | -2.617401 |
| H | -6.930516 | -1.035246 | -0.953440 |
| O | -1.729165 | 0.667774 | 1.716822 |
| C | 2.928984 | 3.475862 | 0.150988 |
| H | 3.866376 | 3.534699 | 0.706948 |
| H | 3.107405 | 3.795057 | -0.878237 |
| H | 2.206111 | 4.155132 | 0.609308 |

Structure $\mathbf{O_{Ph}\bullet\_Int_{6c}}$

| | | | |
|---|---|---|---|
| C | -4.662218 | -0.944233 | 0.429018 |
| C | -4.064735 | 0.258832 | -0.266598 |
| O | -4.228449 | 0.463574 | -1.464143 |
| C | -3.837180 | -2.200984 | 0.076341 |
| C | -2.385748 | -2.082008 | 0.440100 |
| C | -1.823032 | -2.379944 | 1.668077 |
| C | -1.331296 | -1.611030 | -0.377784 |

| | | | |
|---|---|---|---|
| C | -0.132145 | -1.653308 | 0.385941 |
| C | -1.256307 | -1.184558 | -1.747163 |
| N | -0.478501 | -2.125917 | 1.634351 |
| C | 1.103755 | -1.316502 | -0.107248 |
| C | -0.072020 | -0.830313 | -2.279242 |
| C | 1.220305 | -0.800000 | -1.506854 |
| H | -4.678546 | -0.805895 | 1.512942 |
| H | -3.930591 | -2.387365 | -0.996985 |
| H | -2.291265 | -2.763323 | 2.561673 |
| H | 0.158013 | -2.270985 | 2.399057 |
| H | -2.161486 | -1.143513 | -2.343413 |
| H | 2.002631 | -1.443083 | 0.482617 |
| H | -0.007083 | -0.502030 | -3.310754 |
| N | -3.281431 | 1.039358 | 0.509613 |
| C | -2.340154 | 1.989076 | -0.048189 |
| C | -1.033432 | 1.820154 | 0.727048 |
| O | -1.040321 | 1.646808 | 1.940357 |
| H | -3.099288 | 0.761184 | 1.463236 |
| H | -2.195560 | 1.723539 | -1.097841 |
| N | 0.087332 | 1.911121 | -0.005729 |
| C | 1.388924 | 1.615270 | 0.557575 |
| C | 2.351330 | 1.255606 | -0.589071 |
| H | 0.010535 | 1.813772 | -1.008392 |
| H | 1.284053 | 0.798450 | 1.274321 |
| H | 2.646356 | 2.197806 | -1.064021 |
| H | 1.780170 | 2.483627 | 1.096425 |
| H | -5.689297 | -1.070884 | 0.081868 |
| H | -4.281130 | -3.058342 | 0.587642 |
| H | 1.977368 | -1.396583 | -2.034364 |
| C | 3.626009 | 0.564280 | -0.136354 |
| C | 4.407997 | -0.089848 | -1.089772 |
| C | 4.062530 | 0.587282 | 1.187247 |
| C | 5.589653 | -0.721700 | -0.726699 |
| H | 4.076686 | -0.106908 | -2.121951 |
| C | 5.249884 | -0.040501 | 1.551485 |
| H | 3.480987 | 1.089507 | 1.951465 |
| C | 6.014965 | -0.700167 | 0.598022 |
| H | 6.180396 | -1.231345 | -1.479504 |
| H | 5.573007 | -0.015556 | 2.586047 |
| H | 6.936532 | -1.194082 | 0.883895 |
| O | 1.690864 | 0.587766 | -1.650914 |
| C | -2.836394 | 3.432395 | 0.051175 |
| H | -3.778275 | 3.533321 | -0.490737 |
| H | -3.000061 | 3.704213 | 1.096567 |
| H | -2.106838 | 4.120554 | -0.382181 |

Structure **O$_{Ph}$_Prod$_6$**

| | | | |
|---|---|---|---|
| C | 4.948785 | 0.334944 | 0.026813 |
| C | 3.849018 | 1.329153 | 0.340745 |
| O | 3.830804 | 1.950335 | 1.398513 |
| C | 4.698182 | -1.021120 | 0.724588 |
| C | 3.474250 | -1.709613 | 0.202790 |
| C | 3.409580 | -2.565271 | -0.860513 |
| C | 2.113802 | -1.469893 | 0.606809 |
| C | 1.283967 | -2.192944 | -0.275171 |
| C | 1.519516 | -0.681369 | 1.603235 |
| N | 2.096900 | -2.868424 | -1.144213 |
| C | -0.107450 | -2.081833 | -0.247460 |
| C | 0.144610 | -0.589622 | 1.660532 |
| C | -0.654288 | -1.252269 | 0.707842 |
| H | 5.036544 | 0.174346 | -1.050630 |
| H | 4.613341 | -0.853146 | 1.802598 |
| H | 4.207841 | -2.982202 | -1.455873 |
| H | 1.788449 | -3.463055 | -1.893643 |
| H | 2.132865 | -0.120421 | 2.300030 |
| H | -0.744199 | -2.574290 | -0.972054 |
| H | -0.331766 | 0.030479 | 2.410932 |
| N | 2.882505 | 1.460331 | -0.595849 |
| C | 1.664188 | 2.196681 | -0.335267 |
| C | 0.493618 | 1.419112 | -0.944178 |
| O | 0.645280 | 0.694794 | -1.916843 |
| H | 2.850161 | 0.817123 | -1.374495 |
| H | 1.543198 | 2.247725 | 0.749298 |
| N | -0.691833 | 1.643319 | -0.342223 |
| C | -1.931800 | 0.969921 | -0.674056 |
| C | -2.551350 | 0.210721 | 0.519059 |
| H | -0.696364 | 2.236390 | 0.472746 |
| H | -1.734054 | 0.278316 | -1.494618 |
| H | -2.386476 | 0.801952 | 1.429056 |
| H | -2.662255 | 1.706115 | -1.018838 |
| H | 5.884939 | 0.765115 | 0.388504 |
| H | 5.577771 | -1.650637 | 0.569372 |
| O | -2.023191 | -1.103629 | 0.702979 |
| C | 1.720102 | 3.616651 | -0.907819 |
| H | 1.848732 | 3.582670 | -1.992362 |
| H | 0.804764 | 4.167509 | -0.679811 |
| H | 2.565499 | 4.150333 | -0.470392 |
| C | -4.039890 | 0.053255 | 0.308872 |
| C | -4.926117 | 0.964403 | 0.878028 |
| C | -4.536723 | -0.966604 | -0.500962 |
| C | -6.292414 | 0.861148 | 0.640685 |
| H | -4.547036 | 1.758794 | 1.513218 |
| C | -5.902211 | -1.072097 | -0.736445 |
| H | -3.850086 | -1.682350 | -0.937110 |
| C | -6.783465 | -0.158338 | -0.167113 |
| H | -6.973255 | 1.573589 | 1.092425 |
| H | -6.279592 | -1.870354 | -1.365613 |
| H | -7.848640 | -0.242666 | -0.350011 |

**Table S9: Cartesian coordinates of the structures calculated with PCM($Et_2O$)/ωB97X-D/6-311G(d,p).**

Structure **$O_{Me}$•'**

| | | | |
|---|---|---|---|
| C | -3.496227 | -0.866232 | -1.278823 |
| C | -2.737206 | -1.304168 | -0.040890 |
| O | -3.209568 | -1.222566 | 1.083255 |
| C | -3.387420 | 0.673887 | -1.434007 |
| C | -1.984841 | 1.167224 | -1.232048 |
| C | -0.943359 | 1.125021 | -2.116397 |
| C | -1.420196 | 1.578960 | 0.026765 |
| C | -0.037533 | 1.769966 | -0.181405 |
| C | -1.948172 | 1.762175 | 1.312984 |
| N | 0.224619 | 1.510166 | -1.500268 |
| C | 0.823278 | 2.148282 | 0.852346 |
| C | -1.098112 | 2.120857 | 2.340999 |
| C | 0.277586 | 2.315164 | 2.110877 |
| H | -3.087283 | -1.350652 | -2.169633 |
| H | -4.045374 | 1.145774 | -0.699606 |
| H | -0.937223 | 0.827167 | -3.154314 |
| H | 1.157281 | 1.347709 | -1.850008 |
| H | -3.000751 | 1.588697 | 1.503402 |
| H | 1.877131 | 2.312280 | 0.665677 |
| H | -1.492368 | 2.256246 | 3.341786 |
| N | -1.465333 | -1.708433 | -0.279420 |
| C | -0.438953 | -1.688618 | 0.738246 |
| C | 0.896769 | -1.545766 | 0.003790 |
| O | 1.003498 | -1.755395 | -1.192745 |
| H | -1.106274 | -1.651144 | -1.222211 |
| H | -0.591011 | -0.809841 | 1.377143 |
| N | 1.960211 | -1.235260 | 0.785209 |
| C | 3.291953 | -1.178250 | 0.266745 |
| C | 3.786863 | 0.258615 | -0.173802 |
| H | 1.795228 | -0.944974 | 1.735720 |
| H | 3.364804 | -1.837880 | -0.596800 |
| H | 3.563584 | 0.911998 | 0.685960 |
| H | 3.996054 | -1.501264 | 1.035269 |
| H | -4.541911 | -1.160386 | -1.175681 |
| H | -3.764850 | 0.951281 | -2.421229 |
| H | 0.918235 | 2.608296 | 2.935468 |
| O | 3.051282 | 0.602548 | -1.253167 |
| C | -0.450396 | -2.952765 | 1.604106 |
| H | -0.268279 | -3.836014 | 0.987333 |
| H | 0.310320 | -2.905095 | 2.386678 |
| H | -1.429583 | -3.043817 | 2.074301 |
| C | 5.285023 | 0.205118 | -0.461312 |
| H | 5.625493 | 1.176731 | -0.822710 |
| H | 5.845028 | -0.040259 | 0.444786 |
| H | 5.499230 | -0.543683 | -1.227760 |

Structure **$O_{Me}$•_TS$_7$'**

| | | | |
|---|---|---|---|
| C | 3.654029 | -0.053190 | -0.781656 |
| C | 2.874473 | 0.927538 | 0.068040 |
| O | 3.179339 | 1.157695 | 1.233014 |
| C | 3.328987 | -1.497035 | -0.344703 |
| C | 1.880165 | -1.861181 | -0.483813 |
| C | 1.284633 | -2.464153 | -1.554389 |
| C | 0.829665 | -1.638248 | 0.479104 |
| C | -0.356843 | -2.152256 | -0.074153 |
| C | 0.790971 | -1.060698 | 1.755360 |
| N | -0.060874 | -2.638153 | -1.305581 |
| C | -1.613743 | -2.075511 | 0.580887 |
| C | -0.422128 | -1.027403 | 2.446780 |
| C | -1.580233 | -1.564704 | 1.900371 |
| H | 3.424298 | 0.071278 | -1.842982 |
| H | 3.639104 | -1.618948 | 0.697124 |
| H | 1.717195 | -2.803140 | -2.483134 |
| H | -0.728132 | -3.014453 | -1.956869 |
| H | 1.686242 | -0.632958 | 2.192143 |
| H | -2.423248 | -2.726471 | 0.280945 |
| H | -0.451354 | -0.605844 | 3.445115 |
| N | 1.784583 | 1.459705 | -0.519721 |
| C | 0.768461 | 2.170833 | 0.224794 |
| C | -0.544632 | 1.970647 | -0.533595 |
| O | -0.550429 | 1.904169 | -1.753302 |
| H | 1.528648 | 1.196415 | -1.460458 |
| H | 0.704032 | 1.719282 | 1.219004 |
| N | -1.651889 | 1.948769 | 0.235129 |
| C | -2.959659 | 1.634723 | -0.298643 |
| C | -3.347081 | 0.169745 | -0.028395 |
| H | -1.522849 | 1.861756 | 1.230973 |
| H | -2.921891 | 1.810960 | -1.374260 |
| H | -3.360977 | 0.046490 | 1.064106 |
| H | -3.704271 | 2.306721 | 0.138476 |
| C | -4.742109 | -0.141670 | -0.568234 |
| H | -5.011446 | -1.173190 | -0.330339 |
| H | -5.499066 | 0.521047 | -0.137360 |
| H | -4.757879 | -0.026610 | -1.655782 |
| H | 4.718294 | 0.140375 | -0.633161 |
| H | 3.938603 | -2.182205 | -0.938914 |
| H | -2.493715 | -1.568928 | 2.483527 |
| O | -2.386376 | -0.660323 | -0.593852 |
| C | 1.084629 | 3.662206 | 0.364009 |
| H | 1.152147 | 4.124433 | -0.623410 |
| H | 0.307214 | 4.170630 | 0.939344 |
| H | 2.038526 | 3.780929 | 0.879642 |

Structure **$O_{Me}$•_Int$_7$'**

| | | | |
|---|---|---|---|
| C | 3.739527 | -0.182180 | -0.721628 |
| C | 2.960635 | 0.847776 | 0.069549 |
| O | 3.252519 | 1.139086 | 1.222685 |
| C | 3.302987 | -1.603684 | -0.298471 |
| C | 1.830568 | -1.840007 | -0.462790 |
| C | 1.183855 | -2.279797 | -1.586584 |
| C | 0.805249 | -1.560876 | 0.507271 |
| C | -0.414170 | -1.860001 | -0.086273 |
| C | 0.841425 | -1.057517 | 1.834175 |
| N | -0.171910 | -2.296870 | -1.346121 |
| C | -1.757496 | -1.657795 | 0.521526 |
| C | -0.383648 | -0.880882 | 2.505737 |
| C | -1.597375 | -1.155104 | 1.930860 |
| H | 3.577508 | -0.055175 | -1.795336 |
| H | 3.586126 | -1.755704 | 0.746946 |
| H | 1.579003 | -2.588767 | -2.541676 |
| H | -0.880610 | -2.504401 | -2.029035 |
| H | 1.777821 | -0.790181 | 2.306817 |
| H | -2.312523 | -2.608933 | 0.555413 |
| H | -0.365807 | -0.534908 | 3.534423 |
| N | 1.879328 | 1.355164 | -0.561937 |
| C | 0.821012 | 2.033756 | 0.152240 |
| C | -0.478412 | 1.742911 | -0.602295 |
| O | -0.469945 | 1.519773 | -1.803351 |
| H | 1.615479 | 0.999296 | -1.470295 |
| H | 0.766926 | 1.609012 | 1.159353 |
| N | -1.598497 | 1.826514 | 0.145925 |
| C | -2.916213 | 1.533278 | -0.376806 |
| C | -3.416049 | 0.152881 | 0.057539 |
| H | -1.489103 | 1.880202 | 1.146348 |
| H | -2.845477 | 1.565409 | -1.464766 |
| H | -3.480765 | 0.141841 | 1.149253 |
| H | -3.620568 | 2.304107 | -0.050534 |
| C | -4.786263 | -0.152114 | -0.528148 |
| H | -5.132368 | -1.130692 | -0.189646 |
| H | -5.520060 | 0.598715 | -0.222109 |
| H | -4.734463 | -0.163272 | -1.620088 |
| H | 4.801947 | -0.045052 | -0.512255 |
| H | 3.869525 | -2.327993 | -0.889093 |

```
H      -2.493669 -1.072350  2.534105
O      -2.482367 -0.807730 -0.400301
C       1.059081  3.543089  0.251503
H       1.108948  3.983232 -0.747141
H       0.255871  4.028103  0.811263
H       2.003877  3.722430  0.766295
```

Structure **O$_{Me}$•_Prod$_{7}$'**

```
C       3.761835 -0.099300 -0.740169
C       2.937896  0.812615  0.150740
O       3.185971  0.963077  1.338503
C       3.312869 -1.571813 -0.557936
C       1.826986 -1.715717 -0.698568
C       1.100809 -1.732677 -1.858130
C       0.872873 -1.603713  0.372481
C      -0.398895 -1.531192 -0.225086
C       0.981848 -1.500392  1.769767
N      -0.240373 -1.639495 -1.578448
C      -1.556515 -1.284687  0.518962
C      -0.170461 -1.327735  2.507265
C      -1.435229 -1.217454  1.893028
H       3.647208  0.184679 -1.789846
H       3.624274 -1.915249  0.432106
H       1.442566 -1.788212 -2.880796
H      -0.978136 -1.532309 -2.251631
H       1.951239 -1.523305  2.252500
H      -0.111417 -1.254140  3.587293
N       1.870408  1.390227 -0.453500
C       0.758732  1.922790  0.305352
C      -0.504215  1.682007 -0.528351
O      -0.448768  1.541707 -1.740889
H       1.638499  1.126382 -1.401394
H       0.685304  1.356032  1.239787
N      -1.658291  1.666220  0.174677
C      -2.937202  1.317890 -0.413417
C      -3.535961  0.021152  0.140081
H      -1.592865  1.687213  1.180359
H      -2.779295  1.218229 -1.488091
H      -3.629664  0.102467  1.228545
H      -3.654917  2.127526 -0.247006
C      -4.901145 -0.246360 -0.462630
H      -5.309960 -1.176051 -0.065009
H      -5.589882  0.567721 -0.227485
H      -4.824848 -0.336445 -1.549234
H       4.812941  0.008121 -0.466473
H       3.840798 -2.188540 -1.289570
H      -2.312885 -1.054722  2.507244
O      -2.720693 -1.112613 -0.181998
C       0.930358  3.410230  0.624941
H       0.996856  3.990608 -0.298321
H       0.092255  3.784793  1.217591
H       1.850140  3.545307  1.195427
```

Structure **O$_{Ph}$•'**

```
C       4.529694  0.981957 -1.391933
C       3.853079  1.314768 -0.076051
O       4.404695  1.162006  1.003834
C       4.427008 -0.543338 -1.655992
C       3.041856 -1.066777 -1.415235
C       1.956405 -0.980169 -2.241331
C       2.546305 -1.566803 -0.159293
C       1.156342 -1.760617 -0.308863
C       3.141739 -1.827203  1.083427
N       0.825244 -1.418862 -1.592999
C       0.353728 -2.217225  0.739935
C       2.348998 -2.262725  2.127594
C       0.965112 -2.458973  1.955367
H       4.055503  1.524704 -2.214080
H       5.129485 -1.060438 -0.996991
H       1.894844 -0.617255 -3.256482
H      -0.123458 -1.251472 -1.889882
H       4.201321 -1.654401  1.230667
H      -0.706564 -2.382063  0.595775
H       2.795707 -2.458698  3.095782
N       2.561962  1.711232 -0.194247
C       1.617627  1.611034  0.895476
C       0.228662  1.518521  0.261051
O       0.025050  1.810128 -0.904352
H       2.134411  1.711738 -1.109879
H       1.816490  0.687256  1.453202
N      -0.772023  1.154348  1.103412
C      -2.136058  1.135699  0.691191
C      -2.657154 -0.268443  0.120885
H      -0.532861  0.795177  2.013718
H      -2.291937  1.876787 -0.091180
H      -2.407102 -0.989840  0.917738
H      -2.783782  1.349815  1.541281
H       5.576768  1.284222 -1.337293
H       4.750142 -0.741135 -2.680921
H       0.370731 -2.814485  2.790005
C      -4.159414 -0.157233 -0.063467
C      -5.016426 -0.506960  0.976014
C      -4.686853  0.321901 -1.258591
C      -6.392053 -0.376341  0.824279
H      -4.608398 -0.890213  1.906901
C      -6.061605  0.452570 -1.410918
H      -4.013931  0.579039 -2.068655
C      -6.916454  0.105016 -0.370020
H      -7.053972 -0.655416  1.636123
H      -6.467214  0.824207 -2.345057
H      -7.989009  0.204990 -0.490970
O      -1.965680 -0.506925 -1.000287
C       1.700161  2.806835  1.850416
H       1.473367  3.733342  1.317716
H       1.002964  2.698621  2.684341
H       2.713517  2.863842  2.247773
```

Structure **O$_{Ph}$•_TS$_{7}$'**

```
C       4.589327 -0.343170 -0.895583
C       3.910331  0.733267 -0.075126
O       4.262294  0.989089  1.070820
C       4.197451 -1.737754 -0.366542
C       2.723556 -2.009015 -0.430446
C       2.037315 -2.578338 -1.464667
C       1.738327 -1.698903  0.575183
C       0.492728 -2.127314  0.082533
C       1.800808 -1.104204  1.842376
N       0.695200 -2.646873 -1.154124
C      -0.723177 -1.924210  0.784218
C       0.628121 -0.981522  2.592012
C      -0.592254 -1.427198  2.102298
H       4.323204 -0.261900 -1.952578
H       4.542720 -1.824712  0.667463
H       2.400989 -2.960251 -2.406173
H      -0.025453 -2.988344 -1.766548
H       2.743935 -0.735215  2.229309
H      -1.617490 -2.461087  0.497255
H       0.677758 -0.554440  3.587040
N       2.860203  1.336679 -0.667621
C       1.943751  2.184263  0.061343
C       0.582411  2.049420 -0.625686
O       0.503320  1.785830 -1.813691
H       2.537145  1.033162 -1.575496
H       1.880587  1.804564  1.085702
N      -0.481565  2.303418  0.170010
C      -1.829225  1.945112 -0.214344
C      -2.176070  0.523743  0.294259
H      -0.300419  2.373843  1.159411
H      -1.890980  1.969338 -1.302308
H      -2.017292  0.551950  1.383849
H      -2.536333  2.670578  0.195002
H       5.668849 -0.207823 -0.800791
```

| | | | |
|---|---|---|---|
| H | 4.737437 | -2.490895 | -0.945642 |
| H | -1.480040 | -1.349129 | 2.718877 |
| C | -3.638117 | 0.198564 | 0.038890 |
| C | -4.632907 | 0.675068 | 0.891502 |
| C | -4.004536 | -0.563134 | -1.067177 |
| C | -5.971959 | 0.399495 | 0.641754 |
| H | -4.356702 | 1.261694 | 1.763512 |
| C | -5.343098 | -0.841709 | -1.319656 |
| H | -3.224282 | -0.938973 | -1.718837 |
| C | -6.330569 | -0.360505 | -0.466874 |
| H | -6.735877 | 0.772428 | 1.315068 |
| H | -5.616772 | -1.436840 | -2.184107 |
| H | -7.374320 | -0.579489 | -0.662195 |
| O | -1.291790 | -0.344454 | -0.310948 |
| C | 2.399317 | 3.645997 | 0.089159 |
| H | 2.464173 | 4.038219 | -0.928387 |
| H | 1.699851 | 4.261894 | 0.659325 |
| H | 3.382669 | 3.707413 | 0.557380 |

Structure **$O_{Ph}$•_Int$_7$'**

| | | | |
|---|---|---|---|
| C | 4.588080 | -0.506791 | -0.909863 |
| C | 3.943154 | 0.629532 | -0.142260 |
| O | 4.333275 | 0.960423 | 0.970809 |
| C | 4.131103 | -1.865451 | -0.333143 |
| C | 2.642579 | -2.040612 | -0.378419 |
| C | 1.900141 | -2.536559 | -1.417003 |
| C | 1.701432 | -1.622193 | 0.624682 |
| C | 0.430201 | -1.892464 | 0.134134 |
| C | 1.855216 | -1.038183 | 1.909286 |
| N | 0.564107 | -2.443249 | -1.097475 |
| C | -0.835056 | -1.433647 | 0.764223 |
| C | 0.693885 | -0.792683 | 2.668964 |
| C | -0.567759 | -1.025635 | 2.187580 |
| H | 4.333367 | -0.457016 | -1.971794 |
| H | 4.482689 | -1.940282 | 0.699715 |
| H | 2.214350 | -2.959990 | -2.358177 |
| H | -0.195140 | -2.686999 | -1.710336 |
| H | 2.835935 | -0.793135 | 2.296673 |
| H | -1.622587 | -2.199138 | 0.729249 |
| H | 0.805907 | -0.436607 | 3.688150 |
| N | 2.876396 | 1.201994 | -0.741644 |
| C | 1.962282 | 2.060094 | -0.021150 |
| C | 0.594201 | 1.917811 | -0.695148 |
| O | 0.496023 | 1.546941 | -1.852446 |
| H | 2.511306 | 0.818786 | -1.602816 |
| H | 1.896992 | 1.692828 | 1.008684 |
| N | -0.457903 | 2.289223 | 0.074781 |
| C | -1.818986 | 1.927939 | -0.267111 |
| C | -2.198881 | 0.592870 | 0.392124 |
| H | -0.267837 | 2.448782 | 1.052245 |
| H | -1.877815 | 1.830227 | -1.351539 |
| H | -2.095882 | 0.723254 | 1.476232 |
| H | -2.507411 | 2.713299 | 0.052421 |
| H | 5.671005 | -0.410378 | -0.809031 |
| H | 4.623828 | -2.662363 | -0.895822 |
| H | -1.423357 | -0.888560 | 2.838597 |
| C | -3.624817 | 0.193852 | 0.078791 |
| C | -4.664958 | 0.552959 | 0.932710 |
| C | -3.916019 | -0.507965 | -1.089402 |
| C | -5.979183 | 0.220692 | 0.623625 |
| H | -4.445212 | 1.091815 | 1.849745 |
| C | -5.228064 | -0.844123 | -1.398372 |
| H | -3.103078 | -0.793763 | -1.747080 |
| C | -6.263133 | -0.479439 | -0.543342 |
| H | -6.780543 | 0.502173 | 1.297487 |
| H | -5.444452 | -1.392295 | -2.308570 |
| H | -7.286666 | -0.743519 | -0.783900 |
| O | -1.256265 | -0.333892 | -0.086908 |
| C | 2.420700 | 3.521254 | -0.009660 |
| H | 2.482006 | 3.904595 | -1.030874 |
| H | 1.727044 | 4.145131 | 0.558899 |
| H | 3.406383 | 3.582076 | 0.453104 |

Structure **$O_{Ph}$_Prod$_7$'**

| | | | |
|---|---|---|---|
| C | 4.715728 | -0.413210 | -1.023191 |
| C | 4.030698 | 0.690991 | -0.238851 |
| O | 4.413375 | 1.037829 | 0.869202 |
| C | 4.234672 | -1.799715 | -0.523963 |
| C | 2.738065 | -1.880700 | -0.481187 |
| C | 1.886005 | -2.069700 | -1.534695 |
| C | 1.915291 | -1.517865 | 0.641948 |
| C | 0.587857 | -1.484888 | 0.176570 |
| C | 2.185130 | -1.163248 | 1.974550 |
| N | 0.588789 | -1.851223 | -1.140198 |
| C | -0.466289 | -1.037324 | 0.977368 |
| C | 1.131052 | -0.791047 | 2.781973 |
| C | -0.189982 | -0.723553 | 2.293057 |
| H | 4.496576 | -0.318622 | -2.090250 |
| H | 4.641449 | -1.974849 | 0.475440 |
| H | 2.108006 | -2.332536 | -2.558032 |
| H | -0.213166 | -1.821324 | -1.744100 |
| H | 3.201576 | -1.151363 | 2.348344 |
| H | 1.314552 | -0.522359 | 3.816030 |
| N | 2.926288 | 1.211713 | -0.828602 |
| C | 1.926713 | 1.936365 | -0.073246 |
| C | 0.568514 | 1.609587 | -0.702771 |
| O | 0.476923 | 1.230632 | -1.860322 |
| H | 2.580075 | 0.793501 | -1.681303 |
| H | 1.943563 | 1.561475 | 0.955912 |
| N | -0.495983 | 1.800252 | 0.108606 |
| C | -1.847635 | 1.433002 | -0.261591 |
| C | -2.427682 | 0.279459 | 0.574660 |
| H | -0.312360 | 2.022238 | 1.074449 |
| H | -1.828672 | 1.142648 | -1.312706 |
| H | -2.409635 | 0.555982 | 1.633927 |
| H | -2.512709 | 2.295469 | -0.157999 |
| H | 5.793907 | -0.318617 | -0.882508 |
| H | 4.653787 | -2.568608 | -1.177693 |
| H | -0.986109 | -0.400395 | 2.953089 |
| C | -3.859371 | 0.031641 | 0.165231 |
| C | -4.895484 | 0.672759 | 0.839206 |
| C | -4.156941 | -0.788929 | -0.920837 |
| C | -6.213751 | 0.499031 | 0.434205 |
| H | -4.672135 | 1.308328 | 1.690591 |
| C | -5.474765 | -0.968206 | -1.321898 |
| H | -3.354168 | -1.296427 | -1.441862 |
| C | -6.505950 | -0.323350 | -0.647366 |
| H | -7.012543 | 1.000150 | 0.968860 |
| H | -5.697428 | -1.613720 | -2.163918 |
| H | -7.533771 | -0.464514 | -0.961475 |
| O | -1.695771 | -0.930512 | 0.380286 |
| C | 2.179005 | 3.446695 | -0.068653 |
| H | 2.160683 | 3.836452 | -1.089092 |
| H | 1.423617 | 3.970533 | 0.522074 |
| H | 3.159758 | 3.641001 | 0.367146 |

Structure **$O_{PhOH}$•'**

| | | | |
|---|---|---|---|
| C | 4.808726 | 1.074537 | -1.415708 |
| C | 4.159843 | 1.343565 | -0.071686 |
| O | 4.738495 | 1.149847 | 0.987208 |
| C | 4.717529 | -0.440299 | -1.738650 |
| C | 3.343677 | -0.988978 | -1.488955 |
| C | 2.238827 | -0.875959 | -2.285706 |
| C | 2.880505 | -1.545629 | -0.244409 |
| C | 1.488681 | -1.743126 | -0.370859 |
| C | 3.504934 | -1.852515 | 0.973378 |
| N | 1.125791 | -1.350384 | -1.631501 |
| C | 0.712350 | -2.246809 | 0.676113 |
| C | 2.738343 | -2.335328 | 2.016207 |
| C | 1.351995 | -2.533321 | 1.866989 |
| H | 4.308444 | 1.643828 | -2.203738 |
| H | 5.440441 | -0.974380 | -1.116337 |

| | | | |
|---|---|---|---|
| H | 2.151562 | -0.469961 | -3.282506 |
| H | 0.168913 | -1.175731 | -1.897634 |
| H | 4.566520 | -1.678972 | 1.104230 |
| H | -0.350050 | -2.412073 | 0.548921 |
| H | 3.207816 | -2.567385 | 2.965468 |
| N | 2.862262 | 1.729826 | -0.142549 |
| C | 1.942013 | 1.565687 | 0.960090 |
| C | 0.541216 | 1.488133 | 0.350402 |
| O | 0.308470 | 1.848694 | -0.790085 |
| H | 2.415854 | 1.774367 | -1.047850 |
| H | 2.163388 | 0.619086 | 1.468632 |
| N | -0.435819 | 1.056994 | 1.188873 |
| C | -1.808621 | 1.053004 | 0.805977 |
| C | -2.339994 | -0.316366 | 0.160597 |
| H | -0.172094 | 0.641136 | 2.067938 |
| H | -1.984683 | 1.839672 | 0.074114 |
| H | -2.080178 | -1.081548 | 0.912634 |
| H | -2.439345 | 1.209826 | 1.680939 |
| H | 5.853222 | 1.387519 | -1.373296 |
| H | 5.019833 | -0.593011 | -2.777559 |
| H | 0.778494 | -2.925984 | 2.699592 |
| C | -3.841834 | -0.196859 | -0.002237 |
| C | -4.698404 | -0.631742 | 1.002025 |
| C | -4.385215 | 0.377930 | -1.149120 |
| C | -6.075352 | -0.497154 | 0.872800 |
| H | -4.293445 | -1.090460 | 1.899316 |
| C | -5.755447 | 0.518229 | -1.291401 |
| H | -3.722187 | 0.706089 | -1.941661 |
| C | -6.607520 | 0.081039 | -0.277486 |
| H | -6.733988 | -0.847146 | 1.661588 |
| H | -6.184034 | 0.961132 | -2.182422 |
| O | -1.658116 | -0.487660 | -0.978538 |
| C | 2.029511 | 2.715125 | 1.969948 |
| H | 1.782723 | 3.663400 | 1.486729 |
| H | 1.348742 | 2.558214 | 2.809705 |
| H | 3.049437 | 2.764939 | 2.351193 |
| O | -7.940701 | 0.240753 | -0.469783 |
| H | -8.420790 | -0.099672 | 0.287975 |

Structure **$O_{PhOH}•\_TS_7'$**

| | | | |
|---|---|---|---|
| C | 4.854835 | -0.435253 | -0.960843 |
| C | 4.219694 | 0.661940 | -0.132730 |
| O | 4.600707 | 0.913897 | 1.004804 |
| C | 4.442150 | -1.817121 | -0.414442 |
| C | 2.961507 | -2.055332 | -0.446359 |
| C | 2.241509 | -2.617879 | -1.461238 |
| C | 2.004293 | -1.713958 | 0.576140 |
| C | 0.739471 | -2.117267 | 0.112274 |
| C | 2.106211 | -1.110518 | 1.836793 |
| N | 0.904726 | -2.652618 | -1.123165 |
| C | -0.457272 | -1.877585 | 0.835157 |
| C | 0.951965 | -0.953590 | 2.608164 |
| C | -0.288335 | -1.373322 | 2.146079 |
| H | 4.567725 | -0.353952 | -2.012331 |
| H | 4.806517 | -1.905030 | 0.612931 |
| H | 2.577020 | -3.016679 | -2.406218 |
| H | 0.164001 | -2.982881 | -1.717553 |
| H | 3.065381 | -0.760937 | 2.201673 |
| H | -1.370127 | -2.394257 | 0.570441 |
| H | 1.031477 | -0.519648 | 3.598309 |
| N | 3.173956 | 1.289203 | -0.707817 |
| C | 2.291851 | 2.161706 | 0.034082 |
| C | 0.916269 | 2.058966 | -0.630052 |
| O | 0.812177 | 1.796886 | -1.816620 |
| H | 2.826989 | 0.991193 | -1.608664 |
| H | 2.236542 | 1.786173 | 1.060407 |
| N | -0.128281 | 2.338316 | 0.182301 |
| C | -1.490045 | 2.005788 | -0.176548 |
| C | -1.860067 | 0.597804 | 0.351649 |
| H | 0.071499 | 2.405192 | 1.168400 |
| H | -1.570075 | 2.022646 | -1.263509 |
| H | -1.678192 | 0.633127 | 1.437478 |
| H | -2.174448 | 2.749670 | 0.238545 |
| H | 5.939044 | -0.324955 | -0.890342 |
| H | 4.953044 | -2.586069 | -0.999159 |
| H | -1.161830 | -1.268047 | 2.778827 |
| C | -3.331089 | 0.297156 | 0.127304 |
| C | -4.302597 | 0.754860 | 1.012194 |
| C | -3.742145 | -0.427521 | -0.989508 |
| C | -5.651967 | 0.504487 | 0.793244 |
| H | -4.006467 | 1.313667 | 1.895676 |
| C | -5.084118 | -0.687439 | -1.222104 |
| H | -2.987861 | -0.796627 | -1.674841 |
| C | -6.046543 | -0.219170 | -0.329789 |
| H | -6.395438 | 0.865249 | 1.497618 |
| H | -5.404323 | -1.252570 | -2.089746 |
| O | -1.004583 | -0.294334 | -0.262473 |
| C | 2.783941 | 3.611680 | 0.049458 |
| H | 2.840613 | 3.999322 | -0.970338 |
| H | 2.109816 | 4.246140 | 0.629739 |
| H | 3.776555 | 3.650017 | 0.500279 |
| O | -7.348138 | -0.502791 | -0.601863 |
| H | -7.908686 | -0.124904 | 0.078822 |

Structure **$O_{PhOH}•\_Int_7'$**

| | | | |
|---|---|---|---|
| C | 4.833500 | -0.625224 | -0.990902 |
| C | 4.245211 | 0.533967 | -0.211754 |
| O | 4.675062 | 0.856758 | 0.889069 |
| C | 4.351058 | -1.967514 | -0.396887 |
| C | 2.857303 | -2.097536 | -0.403241 |
| C | 2.073637 | -2.574143 | -1.420421 |
| C | 1.955498 | -1.645786 | 0.621231 |
| C | 0.664392 | -1.877941 | 0.164194 |
| C | 2.159993 | -1.060097 | 1.898044 |
| N | 0.749717 | -2.437599 | -1.067734 |
| C | -0.568940 | -1.371639 | 0.821089 |
| C | 1.026539 | -0.773574 | 2.684950 |
| C | -0.253472 | -0.968097 | 2.236153 |
| H | 4.551958 | -0.571436 | -2.045846 |
| H | 4.726952 | -2.049911 | 0.626794 |
| H | 2.350338 | -3.011174 | -2.367133 |
| H | -0.032188 | -2.657686 | -1.660826 |
| H | 3.157354 | -0.844594 | 2.259635 |
| H | -1.384454 | -2.108052 | 0.807848 |
| H | 1.175492 | -0.416104 | 3.698949 |
| N | 3.182327 | 1.137909 | -0.786324 |
| C | 2.317280 | 2.032892 | -0.050103 |
| C | 0.928340 | 1.934107 | -0.688524 |
| O | 0.790108 | 1.568491 | -1.843540 |
| H | 2.785104 | 0.765103 | -1.637634 |
| H | 2.265980 | 1.676073 | 0.984086 |
| N | -0.092244 | 2.336600 | 0.107296 |
| C | -1.471906 | 2.014767 | -0.200477 |
| C | -1.877707 | 0.693860 | 0.470530 |
| H | 0.126856 | 2.488114 | 1.080003 |
| H | -1.559693 | 1.918696 | -1.283137 |
| H | -1.749787 | 0.824406 | 1.552061 |
| H | -2.128606 | 2.820812 | 0.134010 |
| H | 5.921220 | -0.561707 | -0.919784 |
| H | 4.804841 | -2.780655 | -0.969050 |
| H | -1.087854 | -0.799184 | 2.906950 |
| C | -3.316026 | 0.325458 | 0.187511 |
| C | -4.322727 | 0.594886 | 1.107613 |
| C | -3.664970 | -0.268886 | -1.026044 |
| C | -5.650312 | 0.288644 | 0.830642 |
| H | -4.072975 | 1.048403 | 2.062132 |
| C | -4.982049 | -0.583597 | -1.314908 |
| H | -2.885997 | -0.493838 | -1.746000 |
| C | -5.983115 | -0.302479 | -0.385166 |
| H | -6.423386 | 0.502146 | 1.562424 |
| H | -5.255069 | -1.047309 | -2.255448 |
| O | -0.967218 | -0.260006 | -0.023497 |

```
C       2.826778  3.477099 -0.063959
H       2.874391  3.849110 -1.090104
H       2.170742  4.130057  0.516535
H       3.826026  3.507006  0.372016
O      -7.257263 -0.631994 -0.718628
H      -7.851194 -0.387640 -0.005818
```

Structure **$O_{PhOH}$_Prod$_7$'**

```
C       4.984054 -0.496211 -1.107224
C       4.345246  0.624758 -0.307980
O       4.767134  0.967134  0.787262
C       4.484464 -1.870609 -0.592885
C       2.987881 -1.918721 -0.514691
C       2.107475 -2.090874 -1.547711
C       2.199820 -1.535135  0.626325
C       0.863062 -1.472732  0.191734
C       2.508471 -1.184413  1.951618
N       0.825461 -1.840872 -1.124043
C      -0.162529 -0.999099  1.014884
C       1.481912 -0.787271  2.782218
C       0.151886 -0.690365  2.323473
H       4.739833 -0.398198 -2.168528
H       4.910905 -2.052226  0.397153
H       2.299625 -2.361647 -2.575038
H       0.008593 -1.802970 -1.707060
H       3.533278 -1.195220  2.301935
H       1.695172 -0.521393  3.811313
N       3.235881  1.166983 -0.867984
C       2.273325  1.914306 -0.087032
C       0.892707  1.623786 -0.685026
O       0.764965  1.251972 -1.841686
H       2.856665  0.754964 -1.709645
H       2.305202  1.535832  0.940487
N      -0.147030  1.837364  0.151774
C      -1.516535  1.505014 -0.186179
C      -2.105830  0.364386  0.659176
H       0.065083  2.048530  1.114132
H      -1.529292  1.222939 -1.239562
H      -2.055740  0.638578  1.717910
H      -2.157292  2.383002 -0.061579
H       6.067312 -0.426077 -0.994243
H       4.871103 -2.649932 -1.254242
H      -0.621082 -0.348134  3.001200
C      -3.547862  0.135128  0.287538
C      -4.575281  0.542742  1.129421
C      -3.879695 -0.461229 -0.930352
C      -5.907189  0.368231  0.772251
H      -4.341463  1.001855  2.084672
C      -5.201137 -0.648326 -1.296040
H      -3.091383 -0.795159 -1.595446
C      -6.223624 -0.230510 -0.443825
H      -6.697043  0.689637  1.443722
H      -5.461065 -1.116909 -2.237603
O      -1.401871 -0.864249  0.446068
C       2.563828  3.417647 -0.084722
H       2.532121  3.810626 -1.103597
H       1.835536  3.958647  0.524226
H       3.558848  3.585579  0.329096
O      -7.500430 -0.436591 -0.852049
H      -8.110168 -0.115797 -0.184116
```

Structure **$O_{PhO\text{-}}$•'**

```
C       4.218766  0.745985 -2.024344
C       3.925466  1.225231 -0.616940
O       4.766646  1.187328  0.269997
C       4.107788 -0.798330 -2.075838
C       2.819773 -1.305395 -1.499694
C       1.568339 -1.340174 -2.092516
C       2.585080 -1.680415 -0.144429
C       1.180825 -1.936877 -0.054269
C       3.392198 -1.786782  0.999317
N       0.562981 -1.718795 -1.251128
C       0.621646 -2.335201  1.174061
C       2.817418 -2.149673  2.203654
C       1.437671 -2.428934  2.288423
H       3.514035  1.184003 -2.736859
H       4.952123 -1.220707 -1.521665
H       1.340010 -1.088098 -3.124223
H      -0.773423 -0.695908 -1.310687
H       4.450505 -1.550362  0.946577
H      -0.435093 -2.576614  1.244681
H       3.431375 -2.222398  3.095559
N       2.647835  1.641304 -0.415528
C       2.060607  1.728921  0.917763
C       0.550339  1.901863  0.722415
O       0.101814  2.917843  0.213197
H       2.003150  1.449586 -1.167152
H       2.273904  0.795209  1.451766
N      -0.213503  0.884377  1.174698
C      -1.649408  0.868676  1.012150
C      -2.117645 -0.201567 -0.004211
H       0.247107  0.026927  1.453286
H      -1.940847  1.853994  0.646205
H      -1.861131 -1.188848  0.406277
H      -2.129110  0.700789  1.981356
H       5.227372  1.067284 -2.294685
H       4.226518 -1.113343 -3.117295
H       1.012606 -2.731103  3.240502
C      -3.622476 -0.138737 -0.112058
C      -4.429418 -0.558829  0.969610
C      -4.232470  0.363871 -1.277588
C      -5.791926 -0.478040  0.904620
H      -3.952840 -0.962563  1.857770
C      -5.594977  0.458473 -1.369547
H      -3.589027  0.661994 -2.095441
C      -6.453360  0.043494 -0.277435
H      -6.422848 -0.803490  1.723864
H      -6.079080  0.844386 -2.259644
O      -1.509240 -0.020853 -1.241182
C       2.605892  2.911229  1.715101
H       2.375130  3.842615  1.195288
H       2.145845  2.937592  2.706766
H       3.684248  2.809657  1.826992
O      -7.694531  0.125490 -0.348888
```

Structure **$O_{PhO\text{-}}$•_TS$_7$'**

```
C       4.804855 -0.475338 -0.966324
C       4.189581  0.608583 -0.105079
O       4.588284  0.830663  1.033088
C       4.366246 -1.867344 -0.466026
C       2.881276 -2.075143 -0.505042
C       2.145995 -2.575332 -1.542537
C       1.934528 -1.746909  0.530636
C       0.659897 -2.092196  0.051509
C       2.053786 -1.187903  1.812899
N       0.809675 -2.585750 -1.204980
C      -0.533760 -1.824101  0.772028
C       0.903274 -1.013017  2.580903
C      -0.351910 -1.370691  2.098923
H       4.519704 -0.355489 -2.014769
H       4.728797 -1.995441  0.557989
H       2.470632 -2.942703 -2.504073
H       0.056248 -2.846733 -1.817510
H       3.023064 -0.882676  2.190899
H      -1.465501 -2.296237  0.489474
H       0.991053 -0.611053  3.584180
N       3.146266  1.260110 -0.655424
C       2.270139  2.116275  0.112795
C       0.907700  2.085837 -0.589155
O       0.828843  1.849407 -1.784058
H       2.781654  0.977815 -1.554846
H       2.175642  1.688801  1.116071
```

| | | | |
|---|---|---|---|
| N | -0.144845 | 2.397607 | 0.198083 |
| C | -1.507760 | 2.066379 | -0.177972 |
| C | -1.891076 | 0.657720 | 0.341711 |
| H | 0.037817 | 2.428570 | 1.189751 |
| H | -1.579928 | 2.096535 | -1.265086 |
| H | -1.710233 | 0.697137 | 1.428565 |
| H | -2.194456 | 2.804547 | 0.241266 |
| H | 5.890967 | -0.386251 | -0.892120 |
| H | 4.864644 | -2.625364 | -1.075833 |
| H | -1.226181 | -1.244750 | 2.726850 |
| C | -3.344093 | 0.332678 | 0.115515 |
| C | -4.280927 | 0.408927 | 1.150757 |
| C | -3.820620 | -0.043472 | -1.147388 |
| C | -5.621965 | 0.132055 | 0.952186 |
| H | -3.939988 | 0.691540 | 2.147380 |
| C | -5.153961 | -0.329019 | -1.368465 |
| H | -3.111365 | -0.121505 | -1.969376 |
| C | -6.149689 | -0.259837 | -0.328138 |
| H | -6.324223 | 0.192947 | 1.780296 |
| H | -5.497348 | -0.620737 | -2.358640 |
| O | -0.990291 | -0.209693 | -0.268228 |
| C | 2.801160 | 3.548576 | 0.216895 |
| H | 2.889488 | 3.988536 | -0.779370 |
| H | 2.131335 | 4.169524 | 0.816424 |
| H | 3.784641 | 3.534923 | 0.688889 |
| O | -7.370611 | -0.523638 | -0.522544 |

Structure **$O_{PhO-}$•_Int$_7$'**

| | | | |
|---|---|---|---|
| C | 4.742338 | -0.709264 | -0.990940 |
| C | 4.182363 | 0.427027 | -0.157203 |
| O | 4.613253 | 0.675416 | 0.962821 |
| C | 4.220269 | -2.066819 | -0.468346 |
| C | 2.723235 | -2.146881 | -0.486316 |
| C | 1.927683 | -2.511616 | -1.540985 |
| C | 1.834718 | -1.721671 | 0.560530 |
| C | 0.539189 | -1.852084 | 0.077025 |
| C | 2.053287 | -1.223485 | 1.873206 |
| N | 0.609208 | -2.331054 | -1.189292 |
| C | -0.666059 | -1.280331 | 0.735038 |
| C | 0.926586 | -0.909646 | 2.660109 |
| C | -0.354851 | -0.991356 | 2.181132 |
| H | 4.467624 | -0.595135 | -2.042956 |
| H | 4.589434 | -2.212179 | 0.550850 |
| H | 2.192594 | -2.894810 | -2.514152 |
| H | -0.178004 | -2.452916 | -1.803685 |
| H | 3.055987 | -1.086510 | 2.257922 |
| H | -1.547237 | -1.932339 | 0.655718 |
| H | 1.081204 | -0.616659 | 3.694115 |
| N | 3.150603 | 1.098536 | -0.711291 |
| C | 2.302341 | 1.984997 | 0.055723 |
| C | 0.944964 | 2.028918 | -0.657911 |
| O | 0.846192 | 1.724367 | -1.835095 |
| H | 2.750338 | 0.782274 | -1.584090 |
| H | 2.166466 | 1.547359 | 1.051047 |
| N | -0.083333 | 2.476106 | 0.103086 |
| C | -1.465297 | 2.156786 | -0.229035 |
| C | -1.883928 | 0.840119 | 0.439982 |
| H | 0.110148 | 2.556960 | 1.090479 |
| H | -1.535929 | 2.066799 | -1.313167 |
| H | -1.762988 | 0.979848 | 1.524102 |
| H | -2.122712 | 2.963819 | 0.099033 |
| H | 5.831347 | -0.678171 | -0.912803 |
| H | 4.653747 | -2.861831 | -1.080822 |
| H | -1.191098 | -0.789449 | 2.840421 |
| C | -3.298160 | 0.423751 | 0.164241 |
| C | -4.247451 | 0.330066 | 1.187384 |
| C | -3.720385 | 0.091854 | -1.131805 |
| C | -5.552319 | -0.057871 | 0.944815 |
| H | -3.945848 | 0.570847 | 2.206859 |
| C | -5.015068 | -0.304181 | -1.396526 |
| H | -2.998486 | 0.137400 | -1.945192 |
| C | -6.025164 | -0.404039 | -0.370434 |
| H | -6.267191 | -0.122066 | 1.761615 |
| H | -5.314683 | -0.558371 | -2.410660 |
| O | -0.915629 | -0.088987 | -0.032779 |
| C | 2.900333 | 3.387905 | 0.193890 |
| H | 3.029895 | 3.839417 | -0.792668 |
| H | 2.251980 | 4.032736 | 0.791780 |
| H | 3.871938 | 3.316090 | 0.684150 |
| O | -7.210770 | -0.767209 | -0.605803 |

Structure **$O_{PhO-}$_Prod$_7$'**

| | | | |
|---|---|---|---|
| C | 4.970353 | -0.434201 | -1.107452 |
| C | 4.330507 | 0.627109 | -0.230961 |
| O | 4.748232 | 0.884156 | 0.889627 |
| C | 4.458977 | -1.840550 | -0.701853 |
| C | 2.961540 | -1.886202 | -0.638511 |
| C | 2.089754 | -1.978525 | -1.690386 |
| C | 2.164843 | -1.578328 | 0.519531 |
| C | 0.834180 | -1.477483 | 0.077429 |
| C | 2.458959 | -1.317699 | 1.870057 |
| N | 0.807022 | -1.749866 | -1.261867 |
| C | -0.206170 | -1.051180 | 0.911550 |
| C | 1.422122 | -0.968018 | 2.709191 |
| C | 0.097524 | -0.830900 | 2.242094 |
| H | 4.733944 | -0.255703 | -2.160093 |
| H | 4.877231 | -2.098882 | 0.274694 |
| H | 2.290804 | -2.177139 | -2.732506 |
| H | -0.009345 | -1.671252 | -1.842043 |
| H | 3.478421 | -1.357733 | 2.234163 |
| H | 1.623259 | -0.771041 | 3.756495 |
| N | 3.228365 | 1.213401 | -0.757018 |
| C | 2.252949 | 1.893481 | 0.069193 |
| C | 0.884294 | 1.667552 | -0.586631 |
| O | 0.790760 | 1.395002 | -1.775300 |
| H | 2.852469 | 0.861575 | -1.627518 |
| H | 2.258705 | 1.420973 | 1.057540 |
| N | -0.172385 | 1.819770 | 0.236506 |
| C | -1.542396 | 1.520641 | -0.146951 |
| C | -2.162300 | 0.340606 | 0.616011 |
| H | 0.020481 | 1.935462 | 1.219049 |
| H | -1.538376 | 1.311163 | -1.216464 |
| H | -2.101624 | 0.543363 | 1.690845 |
| H | -2.173484 | 2.396703 | 0.025240 |
| H | 6.053285 | -0.378298 | -0.982073 |
| H | 4.847090 | -2.570020 | -1.417472 |
| H | -0.681739 | -0.527143 | 2.930844 |
| C | -3.597895 | 0.139096 | 0.249204 |
| C | -4.606964 | 0.202348 | 1.216505 |
| C | -3.994433 | -0.115341 | -1.072626 |
| C | -5.941089 | 0.040191 | 0.896838 |
| H | -4.329840 | 0.387773 | 2.253792 |
| C | -5.319387 | -0.288713 | -1.413322 |
| H | -3.236711 | -0.190269 | -1.850458 |
| C | -6.389456 | -0.216269 | -0.447693 |
| H | -6.701787 | 0.099760 | 1.671233 |
| H | -5.597468 | -0.489065 | -2.445186 |
| O | -1.425641 | -0.877965 | 0.331302 |
| C | 2.553203 | 3.387319 | 0.216603 |
| H | 2.538802 | 3.874747 | -0.761204 |
| H | 1.818628 | 3.873907 | 0.862844 |
| H | 3.542936 | 3.508819 | 0.658863 |
| O | -7.603523 | -0.366248 | -0.754 |

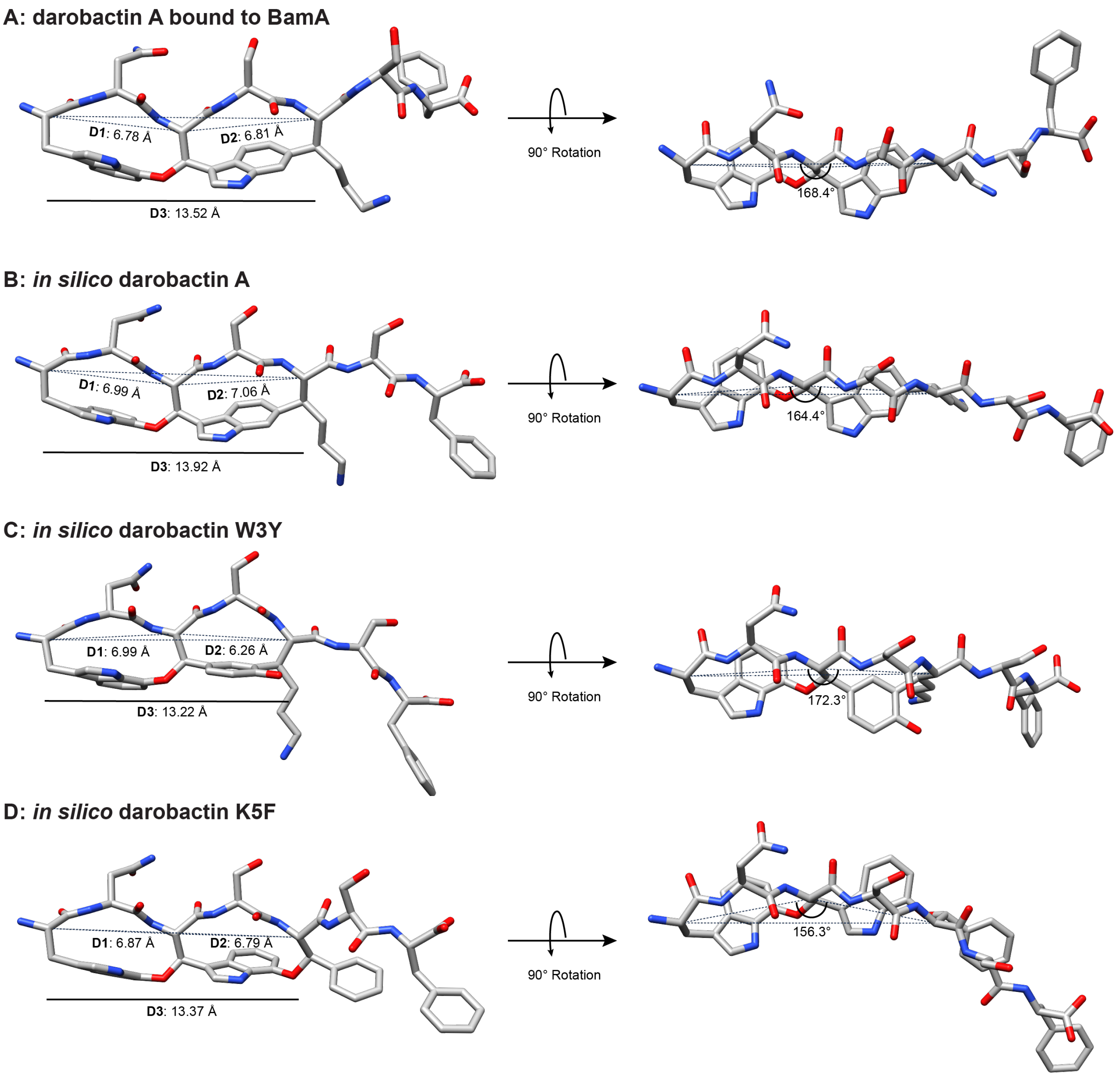

**Figure S87: Macrocycle measurements of darobactins**. (A) From PDB code: 7NRE, darobactin A bound to BamA.[28] *Left view*, dashed lines are distances measured from the Cα of each modified residue. **D1**, **D2**, and **D3** correspond to the distances between core position 1-3, 3-5, and 1-5. *Right view*, rotated edge view of the molecules. The angle formed by D1 and D2 at the Cα of position 3 is given. (B) *In silico* drawn and energy minimized darobactin A using MOE. (C) Same as panel B but for darobactin W3Y. (D) Same for darobactin K5F. The distances and angles are tabulated in **Table S10**, along with other crystallographically determined darobactin:BamA structures.

**Table S10: Distances and angles of modeled and crystallographic darobactins. D1, D2,** and **D3** correspond to distances indicated in **Figure S87**. The angle produced by D1 and D2 at the Cα of position 3 is also given.

| **Molecule** | **D1** | **D2** | **D3** | **∠ Position 3 Cα** |
|---|---|---|---|---|
| Darobactin A (PDB: 7NRE) | 6.78 Å | 6.81 Å | 13.52 Å | 168.4° |
| Darobactin A (PDB: 7NRF) | 6.74 Å | 6.78 Å | 13.46 Å | 169.2° |
| Darobactin B (PDB: 7P1C) | 6.78 Å | 7.08 Å | 13.77 Å | 166.9° |
| *In silico* darobactin A | 6.99 Å | 7.06 Å | 13.92 Å | 164.4° |
| *In silico* darobactin W3Y | 6.99 Å | 6.26 Å | 13.22 Å | 172.3° |
| *In silico* darobactin K5F | 6.87 Å | 6.79 Å | 13.37 Å | 156.3° |

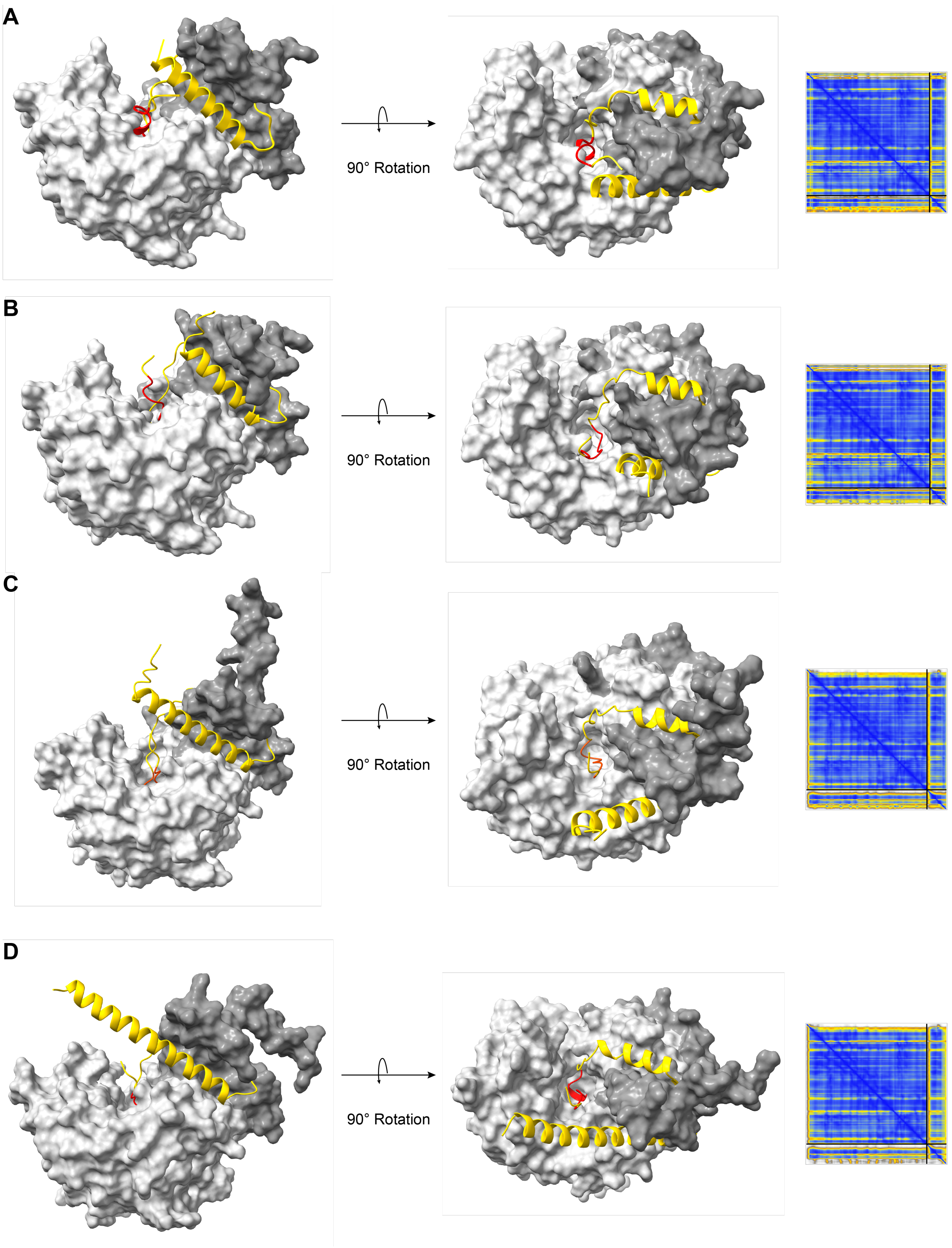
A
90° Rotation
B
90° Rotation
C
90° Rotation
D
90° Rotation

**Figure S88 (previous page): AlphaFold multimer structures of darobactin precursors and their cognate rSAM maturases.** AlphaFold multimer predicted structures and PAE plots of each respective precursor peptide (gold cartoon) and darobactin maturase (gray surface) for (A) darobactin A, (B) darobactin B, and (C) darobactin C, and (D) darobactin D.[15] The darobactin cores are shown in red. Dark gray regions of each darobactin maturase is the N-terminal ~60 residue extension (domain architecture shown in **Figure S89**). *Left*: side on view of precursor peptide and enzyme. *Right*: top-down view. The coordinate files for these predicted structures are available in **Supplementary Dataset 2**.

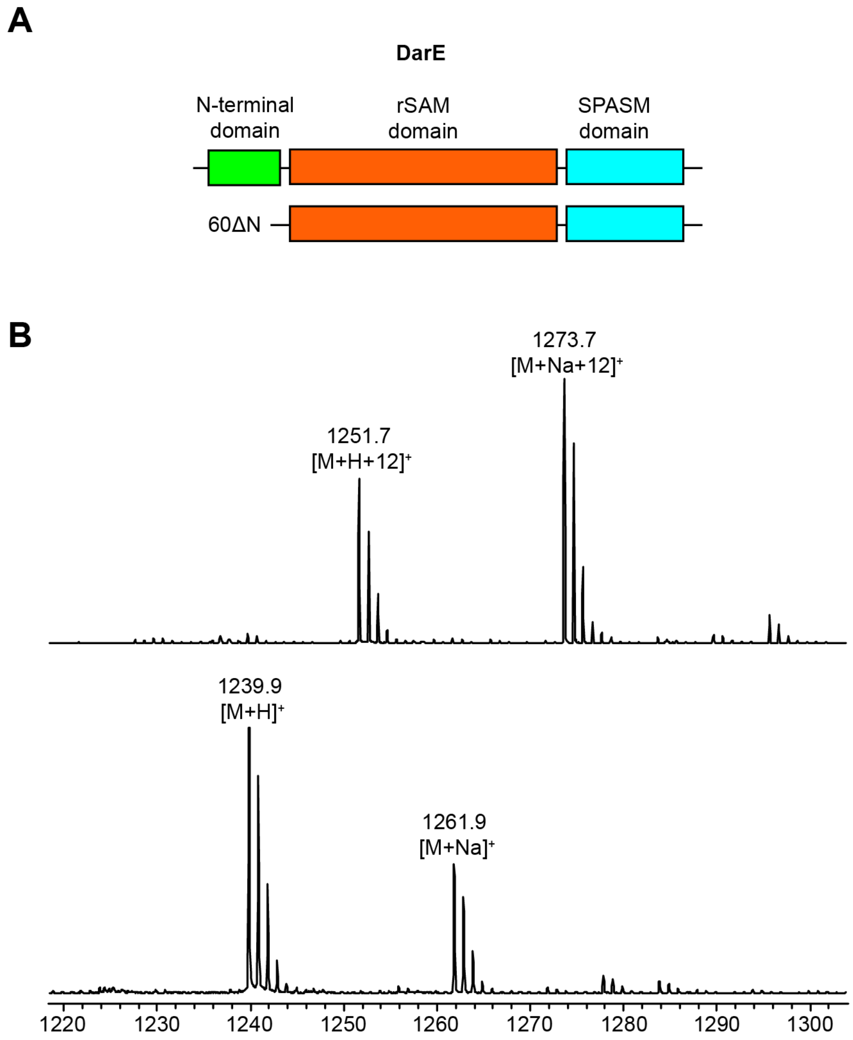

**Figure S89: DarE domain architecture and N-terminal truncation.** (A) Protein domain architecture of DarE. *Top*: full length DarE. *Bottom*: DarE 60ΔN. (B) MALDI-TOF mass spectrum of GluC-digested DarA peptide co-expressed with DarE (*top*) or DarE 60ΔN (*bottom*).

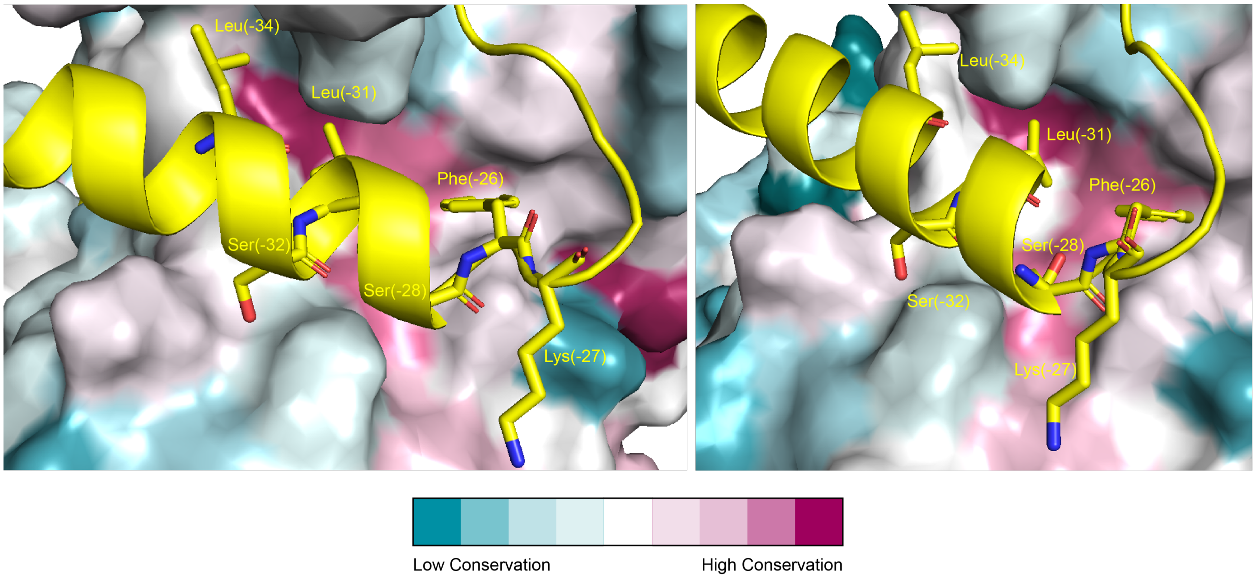

**Figure S90: Conservation of DarE residues.** Zoomed views of the AlphaFold multimer prediction of DarA bound to DarE (**Supplementary Dataset 2**). The DarA leader residues substituted with Ala have the side chain shown in yellow stick. The left and right panels are slight rotations of the same predicted structure to enable a clearer view of the binding pockets of different residues. Conservation of DarE residues was calculated by ConSurf and ranges from teal to purple as indicated.[29]

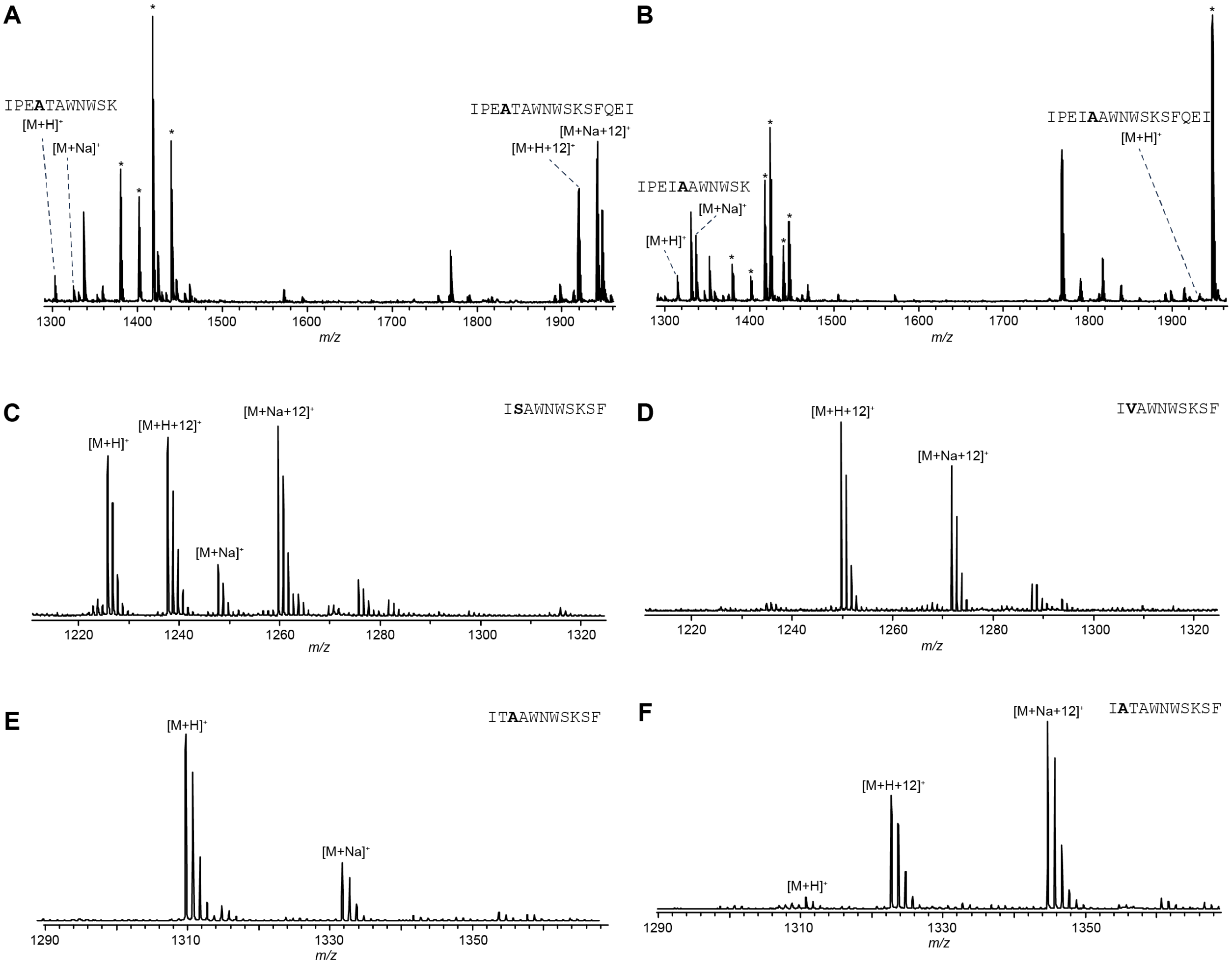

**Figure S91: MALDI-TOF-MS spectra of Thr(-2) Substitutions.** (A) Trypsin digests of DarA variant I(-3)A and (B) DarA variant T(-3)A. The digested peptide sequence is shown with the corresponding observed ions. As trypsin does not cut after crosslinked Lys, unmodified and modified peptides appear in different mass ranges. The region for spanning both the missed trypsin cleavage and proper trypsin cleavage is shown. When both an ether and C-C crosslink are formed on a DarA variant, trypsin misses the Lys5 cut site and the peptide displays a +12 Da mass increase. * indicates unrelated tryptic peptides. (C) GluC digests of DarA variant T(-2)S, (D) DarA variant T(-2)V, (E) the Ala(-2) insertion, and (F) Ala(-3) insertion. Altered residues are bolded.

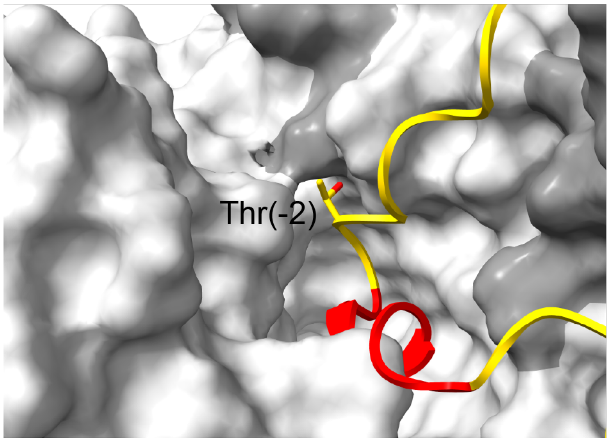

**Figure S92: Hypothetical placement of DarA Thr(-2) with DarE.** AlphaFold multimer predicted structure of DarA and DarE was used (**Supplementary Dataset 2**). DarE is colored in light gray (rSAM and SPASM domains) and dark gray (N-terminal domain). DarA leader and follower regions are yellow while the core region is red. Thr(-2) is shown in stick.

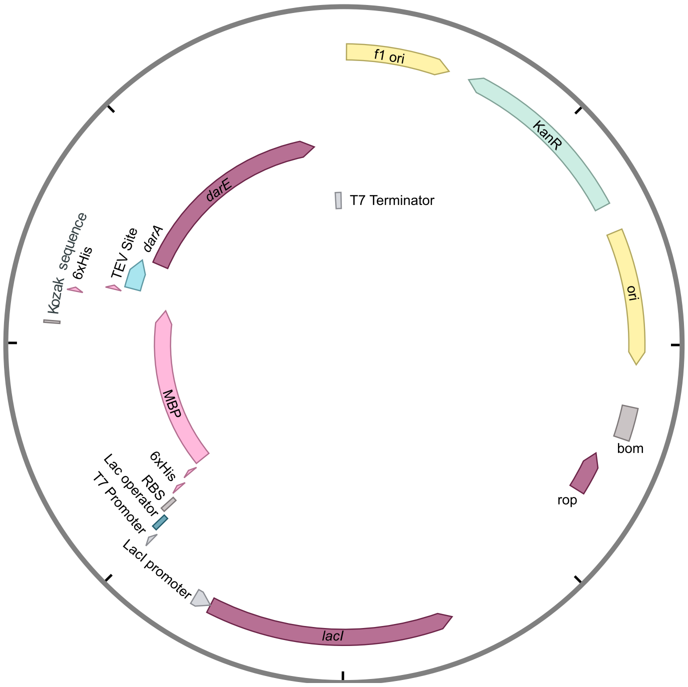

**Figure S93: pET28MBP-DarA/DarE Plasmid Map**. Shown is the plasmid used for all DarA-DarE co-expressions. All mutations described in this work were derived from this plasmid.

*darA:*
ATGCATAACACATTGAATGAAACTGTTAAAACTCAGGAGGCCTTAAATTCCCTTGCTGCCTCGTTCAAGG
AAACAGAGTTATCAATTACGGACAAAGCATTAAACGAATTGTCCAATAAGCCAAAGATCCCAGAAATTAC
TGCTTGGAATTGGTCTAAGTCCTTTCAGGAAATCTAA

*darE:*
ATGGATACGATTATCCCTATTAAGTATTTGGACTCGGACGAATCATCTATCCTGAAGAAGAGCTCCAAGA
TCAACTATCGCCAATTGGCATGTCGTATTATTGGCGAGATCAGCGCAGAAAAAATCCTGGACGACGATGA
ATTGGCACTGTATAATAAGGAAATCAGTATCCATTTCTCACCCGAAATTATCAATGCTAATAAGTTAGTA
GTGGTCGTCAAAGCAACGCGTTTATGCAATCTGCGTTGCACGTATTGCCACTCCTGGGCCGAGGGTAAAG
GCAATACCTTAACTTTCTTCAATTTGATGCGTAGCATCCACCGTTTCTTATCTTTACCGAATATTAAACG
TTTCGAATTTGTGTGGCATGGGGGGGAGGTTACGCTTTTGAACGTTAATTATTTTAAAAAATTAATTTGG
TTACAGGAACAATTCAAGAAACCTGACCAGGTGATTACCAACTCTGTCCAGACGAACGCCGTTAATATCC
CGGAAGATTGGCTTGTTTTCCTTAAGGGAATTGGCATGGGGGTTGGCATTAGTGTAGACGGAATTCCAGA
AATTCACGACAGCCGTCGCCTGGACTACCGCGGCCGTCCTACTTCGCATAAGGTTGCCGCTAGCATGAAG
AAGCTTCGCTCCTACGGAATCCCTTACGGAGCCTTGATTGTCGTAGACCGTGATGTGTATGAGTCTAATA
TCGAAAAAATGCTTTCGTATTTCTATGAAATCGGCCTGACTGATATTGAATTCCTGAACATCGTACCGGA
CAACCGCTGCCAACCAGGGGACGATCCGGGCGGTTCATATATCACCTATCACAACTATATTAATTTCCTT
TCGAAAGTGTTCCGCGTCTGGTGGAATGGCTATCAGGGAAAGATTAACATTCGTTTGTTTGACGGTTTCA
TCGACTCTATCAAGAGTTCTCAAAAAAAGATGAGCGACTGCTATTGGGCAGGGAATTGTTCGCAAGAAAT
TATTACTCTTGAACCAAACGGAACAGTAAGTGCGTGCGACAAGTATGTTGGCGCCGAAGGAAACAATTAT
GGATCCATTATCGATAACGACTTGGGCAACTTATTGTCGAAGTCTAACACCAACAAAGATCATTTAAAGG
AGGAAATGGAAAGTTACGAGAAGATGCACCAGTGCAAATGGTTTCATCTTTGCAATGGCGGCTGCCCGCA
TGACCGCGTGACCAACCGCAAACATAACCCCAATTACGACGGAAGCTGTTGCGGCACTGGGGGCTTATTG
GAGACAATCAAGCAAACCATTGCTGCCTAA

**Figure S94: *E. coli* optimized gene sequences.** The sequences of the commercially obtained synthetic genes shown were cloned into a pET28-MBP plasmid with primers in **Table S1**.

(5) Gaussian 16, Revision C.01, Frisch, M. J.; Trucks, G. W.; Schlegel, H. B.; Scuseria, G. E.; Robb, M. A.; Cheeseman, J. R.; Scalmani, G.; Barone, V.; Petersson, G. A.; Nakatsuji, H.; Li, X.; Caricato, M.; Marenich, A. V.; Bloino, J.; Janesko, B. G.; Gomperts, R.; Mennucci, B.; Hratchian, H. P.; Ortiz, J. V.; Izmaylov, A. F.; Sonnenberg, J. L.; Williams-Young, D.; Ding, F.; Lipparini, F.; Egidi, F.; Goings, J.; Peng, B.; Petrone, A.; Henderson, T.; Ranasinghe, D.; Zakrzewski, V. G.; Gao, J.; Rega, N.; Zheng, G.; Liang, W.; Hada, M.; Ehara, M.; Toyota, K.; Fukuda, R.; Hasegawa, J.; Ishida, M.; Nakajima, T.; Honda, Y.; Kitao, O.; Nakai, H.; Vreven, T.; Throssell, K.; Montgomery, J. A., Jr.; Peralta, J. E.; Ogliaro, F.; Bearpark, M. J.; Heyd, J. J.; Brothers, E. N.; Kudin, K. N.; Staroverov, V. N.; Keith, T. A.; Kobayashi, R.; Normand, J.; Raghavachari, K.; Rendell, A. P.; Burant, J. C.; Iyengar, S. S.; Tomasi, J.; Cossi, M.; Millam, J. M.; Klene, M.; Adamo, C.; Cammi, R.; Ochterski, J. W.; Martin, R. L.; Morokuma, K.; Farkas, O.; Foresman, J. B.; Fox, D. J. Gaussian, Inc., Wallingford CT, 2016.

(13) Jumper, J.; Evans, R.; Pritzel, A.; Green, T.; Figurnov, M.; Ronneberger, O.; Tunyasuvunakool, K.; Bates, R.; Žídek, A.; Potapenko, A.; Bridgland, A.; Meyer, C.; Kohl, S. A. A.; Ballard, A. J.; Cowie, A.; Romera-Paredes, B.; Nikolov, S.; Jain, R.; Adler, J.; Back, T.; Petersen, S.; Reiman, D.; Clancy, E.; Zielinski, M.; Steinegger, M.; Pacholska, M.; Berghammer, T.; Bodenstein, S.; Silver, D.; Vinyals, O.; Senior, A. W.; Kavukcuoglu, K.; Kohli, P.; Hassabis, D. Highly Accurate Protein Structure Prediction with AlphaFold. *Nature* **2021**, *596* (7873), 583–589. https://doi.org/10.1038/s41586-021-03819-2.

(14) Hekkelman, M. L.; de Vries, I.; Joosten, R. P.; Perrakis, A. AlphaFill: Enriching AlphaFold Models with Ligands and Cofactors. *Nat Methods* **2023**, *20* (2), 205–213. https://doi.org/10.1038/s41592-022-01685-y.
(15) Evans, R.; O'Neill, M.; Pritzel, A.; Antropova, N.; Senior, A.; Green, T.; Žídek, A.; Bates, R.; Blackwell, S.; Yim, J.; Ronneberger, O.; Bodenstein, S.; Zielinski, M.; Bridgland, A.; Potapenko, A.; Cowie, A.; Tunyasuvunakool, K.; Jain, R.; Clancy, E.; Kohli, P.; Jumper, J.; Hassabis, D. Protein Complex Prediction with AlphaFold-Multimer. bioRxiv March 10, 2022, p 2021.10.04.463034. https://doi.org/10.1101/2021.10.04.463034.
(16) Case, D.A.; Aktulga, H.M.; Belfon, K.; Ben-Shalom, I.Y.; Berryman, J.T.; Brozell, S.R.; Cerutti, D.S.; Cheatham, III, T.E.; Cisneros, G.A.; Cruzeiro, V.W.D.; Darden, T.A.; Duke, R.E.; Giambasu, G.; Gilson, M.K.; Gohlke, H.; Goetz, A.W.; Harris, R.; Izadi, S.; Izmailov, S.A.; Kasavajhala, K.; Kaymak, M.C.; King, E.; Kovalenko, A.; Kurtzman, T.; Lee, T.S.; LeGrand, S.; Li, P.; Lin, C.; Liu, J.; Luchko, T.; Luo, R.; Machado, M.; Man, V.; Manathunga, M.; Merz, K.M.; Miao, Y.; Mikhailovskii, O.; Monard, G.; Nguyen, H.; O'Hearn, K.A.; Onufriev, A.; Pan, F.; Pantano, S.; Qi, R.; Rahnamoun, A.; Roe, D.R.; Roitberg, A.; Sagui, C.; Schott-Verdugo, S.; Shajan, A.; Shen, J.; Simmerling, C.L.; Skrynnikov, N.R.; Smith, J.; Swails, J.; Walker, R.C.; Wang, J.; Wang, J.; Wei, H.; Wolf, R.M.; Wu, X.; Xiong, Y.; Xue, Y.; York, D.M.; Zhao, S.; Kollman, P.A. Amber 2022, 2022, University of California, San Francisco.
(17) Tian, C.; Kasavajhala, K.; Belfon, K. A. A.; Raguette, L.; Huang, H.; Migues, A. N.; Bickel, J.; Wang, Y.; Pincay, J.; Wu, Q.; Simmerling, C. ff19SB: Amino-Acid-Specific Protein Backbone Parameters Trained against Quantum Mechanics Energy Surfaces in Solution. *J. Chem. Theory Comput.* **2020**, *16* (1), 528–552. https://doi.org/10.1021/acs.jctc.9b00591.
(18) Wang, J.; Wolf, R. M.; Caldwell, J. W.; Kollman, P. A.; Case, D. A. Development and Testing of a General Amber Force Field. *Journal of Computational Chemistry* **2004**, *25* (9), 1157–1174. https://doi.org/10.1002/jcc.20035.
(19) Carvalho, A. T. P.; Swart, M. Electronic Structure Investigation and Parametrization of Biologically Relevant Iron–Sulfur Clusters. *J. Chem. Inf. Model.* **2014**, *54* (2), 613–620. https://doi.org/10.1021/ci400718m.
(20) Izadi, S.; Onufriev, A. V. Accuracy Limit of Rigid 3-Point Water Models. *The Journal of Chemical Physics* **2016**, *145* (7), 074501. https://doi.org/10.1063/1.4960175.
(21) Andersen, H. C. Molecular Dynamics Simulations at Constant Pressure and/or Temperature. *The Journal of Chemical Physics* **2008**, *72* (4), 2384–2393. https://doi.org/10.1063/1.439486.
(22) Andrea, T. A.; Swope, W. C.; Andersen, H. C. The Role of Long Ranged Forces in Determining the Structure and Properties of Liquid Water. *The Journal of Chemical Physics* **1983**, *79* (9), 4576–4584. https://doi.org/10.1063/1.446373.
(23) Miyamoto, S.; Kollman, P. A. Settle: An Analytical Version of the SHAKE and RATTLE Algorithm for Rigid Water Models. *Journal of Computational Chemistry* **1992**, *13* (8), 952–962. https://doi.org/10.1002/jcc.540130805.
(24) Darden, T.; York, D.; Pedersen, L. Particle Mesh Ewald: An N·log(N) Method for Ewald Sums in Large Systems. *The Journal of Chemical Physics* **1993**, *98* (12), 10089–10092. https://doi.org/10.1063/1.464397.
(25) Jones, G.; Willett, P.; Glen, R. C.; Leach, A. R.; Taylor, R. Development and Validation of a Genetic Algorithm for Flexible docking11Edited by F. E. Cohen. *Journal of Molecular Biology* **1997**, *267* (3), 727–748. https://doi.org/10.1006/jmbi.1996.0897.
(26) Verdonk, M. L.; Cole, J. C.; Hartshorn, M. J.; Murray, C. W.; Taylor, R. D. Improved Protein–Ligand Docking Using GOLD. *Proteins: Structure, Function, and Bioinformatics* **2003**, *52* (4), 609–623. https://doi.org/10.1002/prot.10465.
(27) Ma, S.; Xi, W.; Wang, S.; Chen, H.; Guo, S.; Mo, T.; Chen, W.; Deng, Z.; Chen, F.; Ding, W.; Zhang, Q. Substrate-Controlled Catalysis in the Ether Cross-Link-Forming Radical SAM Enzymes. *J. Am. Chem. Soc.* **2023**. https://doi.org/10.1021/jacs.3c04355.
(28) Kaur, H.; Jakob, R. P.; Marzinek, J. K.; Green, R.; Imai, Y.; Bolla, J. R.; Agustoni, E.; Robinson, C. V.; Bond, P. J.; Lewis, K.; Maier, T.; Hiller, S. The Antibiotic Darobactin Mimics a β-Strand to Inhibit Outer Membrane Insertase. *Nature* **2021**, *593* (7857), 125–129. https://doi.org/10.1038/s41586-021-03455-w.
(29) Ashkenazy, H.; Abadi, S.; Martz, E.; Chay, O.; Mayrose, I.; Pupko, T.; Ben-Tal, N. ConSurf 2016: An Improved Methodology to Estimate and Visualize Evolutionary Conservation in Macromolecules. *Nucleic Acids Res* **2016**, *44* (Web Server issue), W344–W350. https://doi.org/10.1093/nar/gkw408.
